# Single-component self-assembling protein nanoparticles displaying stabilized prefusion-closed hemagglutinin trimers for influenza vaccine development

**DOI:** 10.1101/2025.10.25.684546

**Authors:** Yi-Nan Zhang, Xueyong Zhu, Keegan Braz Gomes, Yi-Zong Lee, Connor DesRoberts, Linling He, Ian A. Wilson, Jiang Zhu

**Author notes:** Correspondence and requests for materials should be addressed to: IAW, JZ. These authors contributed equally to this work (Yi-Nan Zhang, Xueyong Zhu, Keegan Braz Gomes).

## Abstract

Current influenza vaccines primarily target hemagglutinin (HA), the major surface glycoprotein and principal determinant of neutralizing antibody (NAb) responses. However, antigenic drift and shift, together with HA’s intrinsic metastability and low-pH sensitivity, pose major challenges to the development of broadly protective influenza vaccines. Here, we stabilize HA in the prefusion-closed conformation through rationally selected amino acid substitutions. One approach targets a conserved residue across multiple influenza A subtypes (H1, H3, H5, and H7) and both influenza B lineages, providing a unified and broadly applicable framework for HA stabilization to support vaccine development. Using H1 and H3 as test cases, stabilized HA trimers were displayed on 24- and 60-mer single-component self-assembling protein nanoparticles (SApNPs) to enhance lymph node trafficking and immunogenicity. Compared with soluble trimers, HA-presenting SApNPs exhibited prolonged retention in lymph node follicles and elicited robust germinal center (GC) responses, key hallmarks of effective virus-like particle (VLP) vaccines. In mice, these SApNPs induced strong humoral immunity and conferred protection against homologous viral challenge. Furthermore, glycan engineering to enrich oligomannose content augments vaccine-induced NAb responses. Together, these findings provide mechanistic insights and establish design principles for next-generation HA-based influenza vaccines targeting both seasonal and pandemic strains.

**ONE-SENTENCE SUMMARY:** Rational HA design and nanoparticle display guide next-generation strategies for influenza vaccine development

## INTRODUCTION

Each year, seasonal influenza causes an estimated 3 to 5 million cases of severe illness and 290,000 to 650,000 deaths worldwide (*1*). Sporadic influenza pandemics have also claimed millions of lives (*2–4*). Influenza is a contagious respiratory disease caused by viruses of the *Orthomyxoviridae* family, which possess negative-sense, single-stranded RNA genomes and are classified into four types: A, B, C, and D (*2, 5*). Among them, influenza A and B viruses pose the greatest threat to human health (*6–8*). Hemagglutinin (HA), the most abundant surface glycoprotein, binds to sialic acid receptors on host cells to mediate viral entry (*9, 10*). Neuraminidase (NA), the second major surface protein, facilitates virion release by cleaving sialic acids from the host cell surface (*10, 11*). Their interplay has been extensively studied in the context of viral entry and immune evasion, revealing a dynamic balance between the two (*12–15*). Matrix protein 1 (M1) forms the viral endoskeleton and plays a critical role in assembly, RNA interaction, and budding (*16*), while matrix protein 2 (M2) functions as a proton channel required for pH regulation during entry and replication (*17, 18*). Influenza A viruses (IAVs) are classified into subtypes based on the antigenic properties of HA (H1–H19) and NA (N1–N11) (*19, 20*). In contrast, influenza B viruses (IBVs) consist of a single subtype divided into two lineages, Victoria and Yamagata (*21*), although the Yamagata lineage appears to have become extinct since the COVID-19 pandemic (*22*).

Given its global burden, seasonal vaccination remains the primary means of preventing severe influenza in the general population (*23–25*). Until the 2024–2025 season, influenza vaccines were quadrivalent, targeting the H1N1 and H3N2 subtypes of influenza A and the Victoria and Yamagata lineages of influenza B; current formulations in the United States for 2025–2026 are trivalent (*26*). Most vaccines are produced in embryonated chicken eggs, in which HA and NA genes from selected strains are inserted into the genome of a donor virus optimized for replication and yield. The resulting recombinant viruses are harvested, inactivated, and formulated into one of three types: (1) whole-inactivated virus vaccines; (2) split-virus vaccines, in which inactivated virions are disrupted with detergent into protein-containing fragments; or (3) subunit vaccines containing purified HA and NA (*27–30*). Despite vaccination efforts, influenza viruses continue to evade immune protection through two main mechanisms (*31–33*). Antigenic drift involves the gradual accumulation of point mutations in HA and NA, producing variants less recognizable to the host immune system and necessitating frequent vaccine updates. Antigenic shift results from gene reassortment between distinct IAVs, leading to the sudden emergence of strains that differ substantially from circulating and vaccine strains. This shift has caused four pandemics since 1918, claiming tens of millions of deaths worldwide. More recently, reassorted highly pathogenic avian influenza (HPAI) strains have raised concern about sustained human-to-human transmission. In addition, the six-month lead time required for egg-based production and frequent mismatches between predicted and circulating strains limit vaccine effectiveness, which has historically ranged from 19% to 60%, although cell culture– based and recombinant vaccines are now available (*26*). These limitations underscore the urgent need for improved influenza vaccine platforms (*34*).

HA on influenza virions forms a homotrimer (*35*). It is synthesized as a precursor, HA0, which is cleaved into HA1, responsible for receptor binding, and HA2, the membrane-anchored subunit that drives membrane fusion. Exposure to low pH triggers a conformational transition from the prefusion to postfusion state, characterized by HA1 displacement and HA2 refolding. This transition releases the N-terminal fusion peptide (FP) from its hydrophobic pocket within HA2 and enables its insertion into the host membrane, initiating fusion and viral entry (*35*). Given its central role in attachment and membrane fusion, HA is the primary target of host immune responses (*36, 37*). Over the past two decades, hundreds of broadly neutralizing antibodies (bNAbs) recognizing diverse HA epitopes have been identified and characterized, providing key insights into how the humoral immune system engages HA and blocks viral entry (*38–43*). These discoveries have catalyzed the development of recombinant HA-based vaccines (*44–47*), with a focus on conserved epitopes to achieve broad protection across subtypes and strains (*48–50*). Two major immunogen design strategies have emerged: one based on full-length HA ectodomains and the other targeting the stalk region. In the former, chimeric HAs have been engineered to redirect antibody responses toward the stalk (*51–53*), while the COBRA (computationally optimized broadly reactive antigen) strategy has been used to resurface HA and broaden neutralizing antibody (NAb) responses (*54–56*). In stalk-based designs, “headless” HAs—lacking the immunodominant globular head—were an intuitive starting point (*57*), later refined through multiple design iterations (*58–60*). To further expand the breadth of vaccine-elicited NAb responses, protein nanoparticles (NPs) have been used to display HAs from multiple subtypes (mosaic HA-NPs) or conserved stem antigens (*61–64*). Several ferritin-based HA-NP vaccines have advanced to clinical trials (*65–68*). Although HA is highly sensitive to low pH (*69–71*), structure-guided stabilization remains largely underexplored (*72–74*), representing a critical gap in the design of next-generation HA-based influenza vaccines.

Previously, we established a rational vaccine design strategy for class I viral glycoproteins (*75–77*), integrating antigen optimization based on metastability analysis, multivalent display on self-assembling protein nanoparticles (SApNPs), and glycan remodeling. This approach has been applied to human immunodeficiency virus type 1 (HIV-1) (*78–81*), filoviruses (*82, 83*), severe acute respiratory syndrome coronaviruses (SARS-CoV-1/2) (*84, 85*), respiratory syncytial virus (RSV), and human metapneumovirus (hMPV) (*86*), with immunogens extensively evaluated in vivo. Building on these advances, we extended this strategy to influenza virus. Sequence–structure analysis identified a conserved inward-facing residue in the HA2 central triple helix—N95 in IAV and Q95 in IBV—as a potential determinant of HA dissociation. This site likely contributes to HA metastability and facilitates the low-pH-triggered transition from the prefusion to postfusion state. We hypothesized that substituting N95/Q95 with leucine could stabilize HA in a prefusion-closed conformation and, when combined with pH-switch mutations, further enhance stability. This strategy was validated using six HAs from the H1, H3, H5, and H7 subtypes of IAV and both IBV lineages, yielding stabilized prefusion-closed HA trimers. Antigenicity analyses of H1 and H3 HAs with distinct glycan modifications revealed native-like antibody binding profiles that were subtype-specific and glycan-dependent. N95L-stabilized H1 and H3 trimers were then displayed on 24-meric ferritin (FR) and multilayered 60-meric E2p and I3-01v9 SApNPs to generate virus-like particles (VLPs). These SApNPs showed prolonged retention in lymph node follicles and induced stronger germinal center (GC) reactions than soluble trimers in mice. Stabilized HA trimers and HA-presenting SApNPs elicited robust binding (bAb), hemagglutination inhibition (HAI), and NAb responses in mice, exhibiting diverse cross-reactivity. Unexpectedly, enrichment of oligomannose glycans positively correlated with NAb induction and protection. Together, these findings establish a rational design platform for HA-based influenza vaccines and may inform future strategies for both seasonal and pandemic preparedness.

## RESULTS

### Rational design and in vitro characterization of HA trimers for group 1 IAVs

We previously investigated the molecular basis of glycoprotein metastability across diverse class I viruses (*75–77*). Triggers of metastability are often localized within specific regions of the heptad repeat 1 (HR1) and/or heptad repeat 2 (HR2) stalk in the fusion domain (*79, 80, 82, 83, 85, 86*). Metastability can manifest as conformational disorder or trimer dissociation, facilitating the rapid transition from the prefusion to postfusion state. Byrd-Leotis et al. reported that K58I and D112G mutations in HA2 shifted the pH threshold for membrane fusion across multiple IAV strains (*74*). Milder et al. identified two histidine-containing pH-switch regions in HA2 and showed that H26W (pHS1) and K51I/E103I (pHS2) mutations improved HA stability across diverse IAV subtypes (*73*). Lee et al. introduced an interprotomer disulfide bond between HA1 residue 30 and HA2 residue 47, enhancing the structural and thermal stability of PR8/H1 and HK68/H3 HA trimers (*72*). While promising, these design strategies remain limited by subtype or strain specificity and have not been extended to IBVs. A generalizable approach is therefore needed to stabilize prefusion-closed HA trimers across group 1 and 2 IAVs as well as IBVs.

To identify conserved features underlying HA metastability, we analyzed HA sequences from human (47,554), avian (17,672), and swine (10,191) IAVs, as well as IBVs (16,646), with a focus on residues 38–125 in HA2, which encompass HR1 and the HR1–HR2 linker. The central helix (residues 75–116) exhibited a distinct sequence conservation pattern (**Fig. 1a**, left). Mapping this region onto an H1 HA structure (PDB 3LZG) revealed that inward-facing residues 77, 80, 84, and 91, located near the top of the central coiled coil, are predominantly hydrophobic aliphatic residues, whereas residue 88 is more variable (**Fig. 1a**, right). Notably, a hydrophilic residue—N95 in IAVs and Q95 in IBVs—is consistently present across all sequences analyzed (**Fig. 1a**, right). N95/Q95 is followed by a conserved hydrophobic stretch (residues 96 and 98– 102) and a region enriched in inward-facing charged or polar residues. This sequence–structure pattern likely enables HA2 to maintain the prefusion trimer while remaining primed for conformational change during fusion. N95/Q95 in HA is reminiscent of W615 in ebolavirus GP (*82, 83*) and the acidic patch in RSV F (*86*), both creating suboptimal interactions that promote metastability. Based on these observations, we hypothesized that a hydrophobic substitution at position 95 (e.g., to leucine) could reduce sensitivity to low pH, prevent trimer dissociation, and complement other stabilizing mutations. To test this hypothesis, we designed a panel of soluble HA constructs from diverse subtypes (**Fig. S1a**), all truncated at residue 175 in HA2 for IAV (and the equivalent position for IBV), including wildtype (WT), N95L (or Q95L), and N95L (or Q95L) combined with pHS1/2 mutations such as H26W/K51I/E103I (HKE) in IAVs (*73*). All constructs were appended with a foldon trimerization motif and a C-terminal His_6_ tag to enable purification by immobilized metal affinity chromatography (IMAC) using a nickel column.

**Fig. 1.**
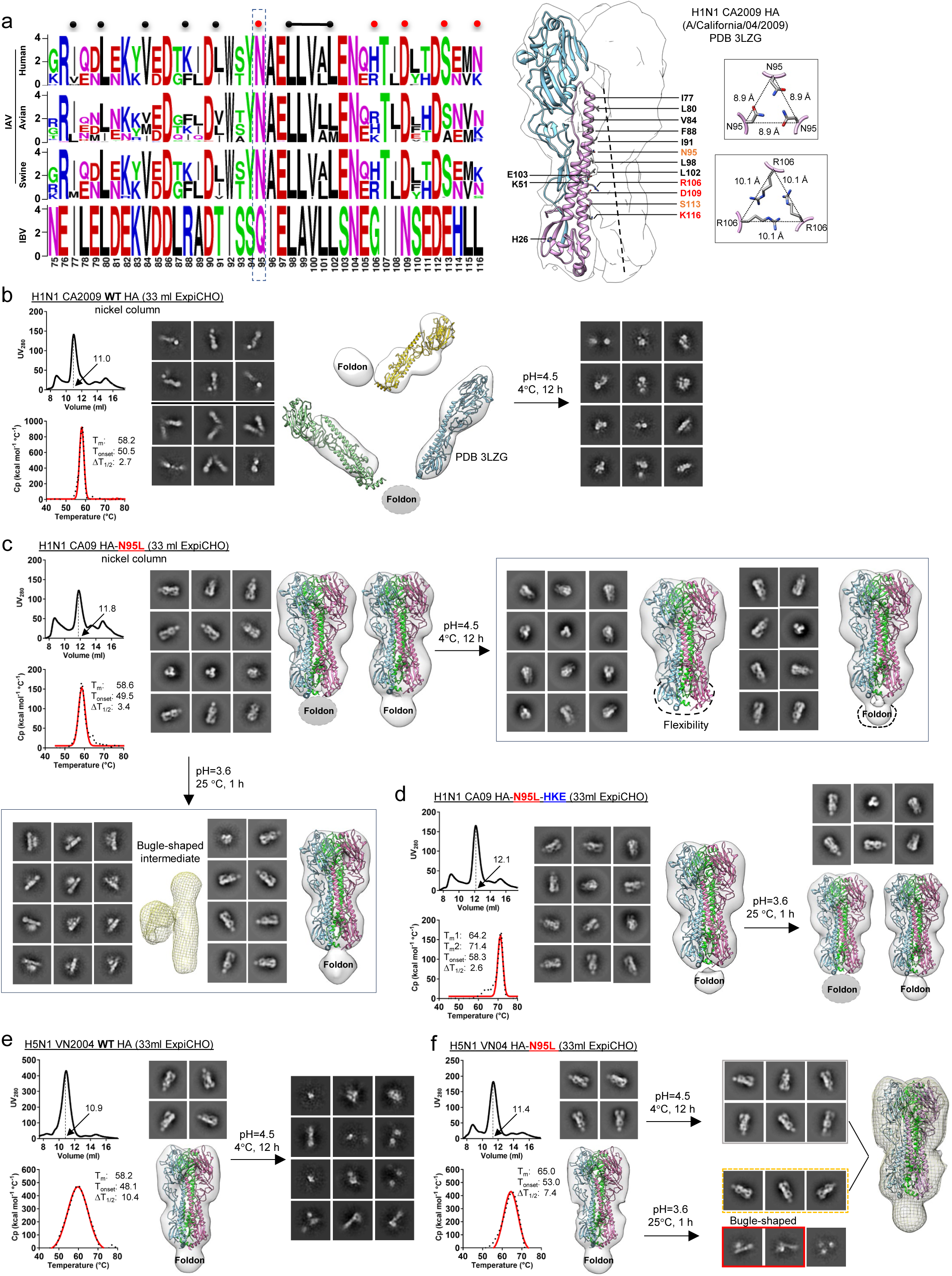
Rational design and in vitro characterization of stable prefusion-closed HA trimers for group 1 influenza A viruses. (**a**) Structure-guided selection of HA-stabilizing mutations. Left: Sequence logo analysis of HA2 residues 75–116, corresponding to the central helix, from IAVs of human, avian, and swine origin, as well as IBVs. Numbering follows IAV HA2 convention. Right: Ribbon representation of the HA ectodomain protomer within the trimeric molecular surface, with HA1 shown in light blue and HA2 in plum. The crystal structure of an H1N1 HA (PDB 3LZG) is used to depict the architecture. Inward-facing residues targeted for stabilization are shown as stick models. Boxed insets display the top-down views of residues N95 and R106 in the trimeric context. (**b**–**d**) In vitro characterization of A/California/04/2009(H1N1) (CA09) WT HA (**b**), HA-N95L (**c**), and HA-N95L-HKE (**d**) constructs. Shown are size exclusion chromatography (SEC) profiles, differential scanning calorimetry (DSC) thermograms, representative 2D class averages, and 3D reconstructions from negative-stain EM (nsEM) analysis. Trimer peak positions are indicated on SEC traces; major thermal parameters are annotated on DSC curves. nsEM images are also shown after relatively mild (pH 4.5, 4 °C, 12 h) and harsh (pH 3.6, 25 °C, 1 h) acidic challenges. The 3D reconstruction of a bugle-shaped HA intermediate identified in (**c**) is shown as a yellow mesh. The CA09 HA structure (PDB 3LZG) was fitted into EM density maps. (**e**,**f**) In vitro characterization of A/Vietnam/1203/2004 (VN04) WT HA (**e**) and HA-N95L (**f**) constructs. SEC profiles, DSC thermograms, and nsEM 2D/3D analyses are shown as in (**b–d**). All group 1 HA trimers were transiently expressed in ExpiCHO cells and purified using a nickel column.

We first evaluated three constructs derived from A/California/04/2009 (CA09), a pandemic H1N1 strain (*87*). HAs were transiently expressed in ExpiCHO cells and purified using a nickel column, with yield, thermostability, and conformation assessed by size-exclusion chromatography (SEC), differential scanning calorimetry (DSC), and negative-stain electron microscopy (nsEM), respectively. The SEC-purified HA was subjected to a two-phase acidic challenge: prolonged mild treatment (pH 4.5, 4 °C, 12 h), followed, if stable, by brief exposure to harsher conditions (pH 3.6, 25 °C, 1 h). In SEC, WT HA showed a dominant trimer peak at 11.0 ml, with minor aggregates (8.8 ml) and a broad shoulder (13.8–15.1 ml) corresponding to dimers and monomers (**Fig. 1b**, top left). The purified trimer displayed a distinct DSC peak with a melting temperature (T_m_) of 58.2 °C (**Fig. 1b**, bottom left). nsEM revealed single- and double-chain particles tethered to the foldon motif in 2D class averages, consistent with a mixture of open trimers in various conformations (**Fig. 1b**, middle). Fitting the 3D density map with the H1 HA structure (PDB 3LZG) confirmed the tendency of the CA09 HA trimer to open, as previously reported (*88–90*). Moreover, WT HA failed to withstand pH 4.5 and displayed multiple postfusion species (**Fig. 1b**, right). HA-N95L showed a similar SEC profile, with the trimer peak shifted to 11.8 ml, and a comparable DSC thermogram, though with a slightly broader melting transition (ΔT_1/2_ = 3.4 °C vs. 2.7 °C for WT) (**Fig. 1c**, top left). However, nsEM revealed a striking difference: 2D class averages showed prefusion-closed trimers with discernible head and stem regions, and the crystal structure (PDB 3LZG) fit the 3D density map with near-perfect agreement (**Fig. 1c**, top left). These results indicate that N95L effectively prevents trimer opening. After prolonged acidic treatment (pH 4.5, 4 °C, 12 h), most trimers remained in a prefusion-closed conformation, although some showed reduced stem density (**Fig. 1c**, top right). A brief, more acidic exposure (pH 3.6, 25 °C, 1 h) produced a bugle-shaped intermediate, while a small fraction of trimers remained prefusion-closed (**Fig. 1c**, bottom left). Recent studies have elucidated stepwise HA conformational transitions induced by endosomal acidification (pH ∼5.0) (*91, 92*). The bugle-shaped HA observed here resembled the intermediate state IV defined by cryo-EM (*91*). We next examined the combined effect of N95L and the previously reported HKE mutation (*73*). For HA-N95L-HKE, the trimer peak shifted further to 12.1 ml, suggesting tighter packing (**Fig. 1d**, top left). The HKE mutation enhanced thermostability, increasing the T_m_ to 71.4 °C (**Fig. 1d**, bottom left), with nsEM uniformly showing prefusion-closed trimers (**Fig. 1d**, middle). Notably, HA-N95L-HKE trimers remained fully intact even after brief pH 3.6 treatment, with no detectable conformational change (**Fig. 1d**, right).

Recent outbreaks of avian influenza A (H5N1) in poultry and dairy cattle in the United States have renewed concern about zoonotic spillover (*93–95*). This underscores the need for a general strategy to stabilize H5 HA for vaccine development. Here, we evaluated N95L alone and in combination with pHS1/2 mutations using the A/Vietnam/1203/2004(H5N1) (VN04) strain as a test case (*96–98*). Following ExpiCHO expression and nickel affinity purification, WT H5 HA exhibited a prominent trimer peak at 10.0 ml, with minimal signals corresponding to aggregates, dimers, or monomers (**Fig. 1e**, top left). Its melting point (58.2 °C) was identical to that of WT H1 HA (**Fig. 1e**, bottom left), but the peak was notably broader (ΔT_1/2_ = 10.4 °C). The crystal structure of this H5 HA trimer (PDB 3GBM) fit well into the nsEM-derived 3D density map (**Fig. 1e**, middle). As expected, mild acidic treatment at pH 4.5 overnight at 4 °C triggered a conformational shift to the postfusion state (**Fig. 1e**, right). The N95L mutation improved biophysical properties: it shifted the trimer peak to 11.4 ml in SEC, increased the T_m_ by 6.8 °C, narrowed the melting peak (7.4 °C for N95L vs. 10.4 °C for WT) in DSC, and preserved the prefusion-closed conformation in nsEM (**Fig. 1f**, left and middle). Most HA-N95L trimers remained in a prefusion-closed state after mild acidic challenge at pH 4.5, but both bugle-shaped fusion intermediates and postfusion species were observed following a brief exposure to pH 3.6 at 25 °C (**Fig. 1f**, right). HA-N95L-HKE showed improved thermostability over N95L alone while retaining the prefusion-closed state (**Fig. S1b**, top left). Mild low-pH treatment had no adverse effects on HA structure, as shown by nsEM (**Fig. S1b**, top right). Further lowering the pH to 3.6, even briefly at room temperature, induced visible structural disruption: some particles adopted extended postfusion shapes and irregular morphology, although most trimers remained stable in a prefusion-closed conformation (**Fig. S1b**, bottom left). These results demonstrate the stabilizing effect of the N95L mutation on H5 HA and the potential advantage of combined use with the pHS1/2 mutation.

Other stabilizing mutations and factors critical to HA purification and nsEM analysis were examined using group 1 HAs as examples. We first investigated the inward-facing residue R106, located just downstream of the hydrophobic stretch (residues 98–102) in the central triple helix, which forms a repulsive cluster around the trimer axis (**Fig. 1a**, right). To determine whether R106 affects HA metastability similarly to N95, we generated three HA constructs based on CA09 H1: R106M (R), E103L/R106M (ER), and N95L/E103L/R106M (N95L-ER) (**Fig. S1c**). R106M alone improved HA expression and purity but had little effect on thermostability, with open HA trimers still observed by nsEM (**Fig. S1d**). The HA-ER construct produced mixed results—reducing HA purity during expression but markedly enhancing thermostability (T_m_ increased by <7 °C) and preventing trimer opening (**Fig. S1e**). HA-N95L-ER showed further reduced expression, likely due to increased hydrophobicity, but improved trimer thermostability and preserved the prefusion-closed conformation (**Fig. S1f**). We next evaluated immunoaffinity chromatography (IAC) for HA purification, as used for other viral glycoproteins (*78–86, 99–101*). For CA09 H1 HA, we prepared an IAC column using bNAb 2D1, which targets an epitope near the receptor-binding site (RBS) on the globular head (*102*) (**Fig. S1g**). To prevent low-pH-induced conformational changes, we used a high-salt elution buffer (3M MgCl_2_, pH 7.2). 2D1 purification reduced non-trimer species and modestly improved thermostability for WT HA, HA-N95L, and HA-N95L-HKE (**Fig. S1h-j**), preserving the prefusion-closed conformation once N95L was introduced. For VN04 H5 HA, we prepared an IAC column using the stem-specific bNAb CR6261 (*97*). For WT HA and HA-N95L, CR6261 purification led to substantial aggregation (**Fig. S1k, l**) but did not compromise trimer structural integrity, as confirmed for HA-N95L by nsEM (**Fig. S1l**). Finally, nsEM micrographs of H1 and H5 HA samples were examined for anisotropy (**Fig. S1m**). CA09 HA constructs—particularly those containing multiple mutations—showed clumping that increased orientation bias and reduced stem density. In contrast, VN04 constructs exhibited no such anisotropy.

Our results highlight the importance of N95 in HA metastability and demonstrate that hydrophobic substitution at this site can enhance structural and thermal stability in both H1 and H5 strains. The intrinsic tendency of the CA09 H1 HA trimer to splay open suggests that a broadly applicable stabilizing mutation such as N95L could be incorporated into HA-based vaccines to mitigate potentially suboptimal biophysical, structural, and immunogenic properties in pandemic strains. The pHS1/2 mutations (HKE) may further enhance the acid resistance of HA-N95L.

### In vitro and structural characterization of stabilized group 2 influenza A HA trimers

Group 2 IAVs include HA subtypes H3, H4, H7, H10, H14, and H15 (*103, 104*). Among these, H3N2 caused the 1968 Hong Kong pandemic (*105*) and remains one of the two IAV subtypes included in current seasonal vaccines (*106*). The avian-origin H7N9 has also been associated with at least five outbreaks since 2013 (*107, 108*). Notably, the N38 glycan in HA1—a defining feature of group 2 HAs—is absent from most group 1 HAs and contributes to differences in HA stability and NAb recognition between the two groups (*109*). Since the fusion pH of IAV HAs falls within a relatively narrow range of 4.8–6.2 (*110–112*), it is important to determine whether group 2 HAs exhibit similar sensitivity to acidic conditions. To this end, we designed a series of soluble HA constructs based on A/Hong Kong/1/1968 (H3N2) (HK68), the original pandemic H3N2 strain (*113*), and A/Guangdong/Th005/2017 (GD17), a highly pathogenic avian H7N9 strain (*114*), and evaluated them using a validation pipeline similar to that used for the H1 and H5 HAs.

Three HK68 H3 HA constructs—WT, N95L, and N95L-HKE—were transiently expressed in ExpiCHO cells and purified using a nickel column for in vitro characterization and a two-phase acidic challenge. WT HA exhibited a trimer peak at 11.9 ml in SEC, with reduced aggregation and fewer monomer/dimer species compared with CA09 HA (**Fig. 2a**, left top). DSC revealed a T_m_ of 55.1 °C, with unfolding onset at 46.2 °C, both slightly lower than those of CA09 HA (**Fig. 2a**, left bottom). nsEM confirmed that WT HA maintained a prefusion-closed trimer, suggesting that trimer dissociation is not intrinsic to this pandemic strain (**Fig. 2a**, middle). However, exposure to mild acidity produced heterogeneous particles, with some 2D classes showing a single globular head domain loosely attached to the central helical coiled coil (**Fig. 2a**, right). The N95L mutation did not alter the SEC profile but increased the T_m_ by 2.4 °C and the T_onset_ by 4.6 °C (**Fig. 2b**, left). Previously, Milder et al. reported a crystal structure of HK68 HA containing the pHS1/2 mutations (HKE) (*73*). Here, to assess the structural impact of N95L, we determined a 2.3-Å resolution crystal structure of HK68 HA-N95L (**Fig. 2c**, **Fig. S2b**, **Table S1**), which showed a C_α_ root mean square deviation (RMSD) of 0.3 Å relative to the WT HA structure (PDB 4FNK) (*115*). In WT HA, N95 is sandwiched between L91 and L98, forming three triangles around the trimer axis with C_β_–C_β_ distances of 5.7, 8.4, and 8.6 Å for L91, N95, and L98, respectively. Upon N95L substitution, these distances shifted to 5.9, 7.6, and 9.1 Å, suggesting that the hydrophobic leucine (L95) promotes closer packing and slightly displaces inward-facing contacts beyond position 95. In the crystal structure, L95 forms hydrophobic interactions with surrounding HA2 residues L91, W92, and L98 (**Fig. 2c**, **Fig. S2b**), providing a structural basis for enhanced trimer stability in the HA-N95L construct. No major conformational changes were observed around L95, except for the side chains of HA2 Y94, which rotated slightly away from the three-fold axis at the center of the trimer. Incorporating the HKE mutations (*73*) into the HA-N95L construct improved trimer yield and further reduced monomer/dimer content (**Fig. 2d**, top left). DSC revealed a substantial increase in thermostability, with a >12 °C increase in T_m_ (**Fig. 2d**, bottom left). nsEM confirmed that HA-N95L-HKE retained a stable prefusion-closed trimer conformation (**Fig. 2d**, middle). Notably, HA-N95L-HKE exhibited enhanced resistance to acidic challenge: a subpopulation of trimers maintained the prefusion-closed trimeric state upon mild acidic challenge and even after exposure to pH 3.6 at ambient temperature, although a large portion of 2D classes showed signs of structural rearrangement and particle heterogeneity (**Fig. 2d**, right).

**Fig. 2.**
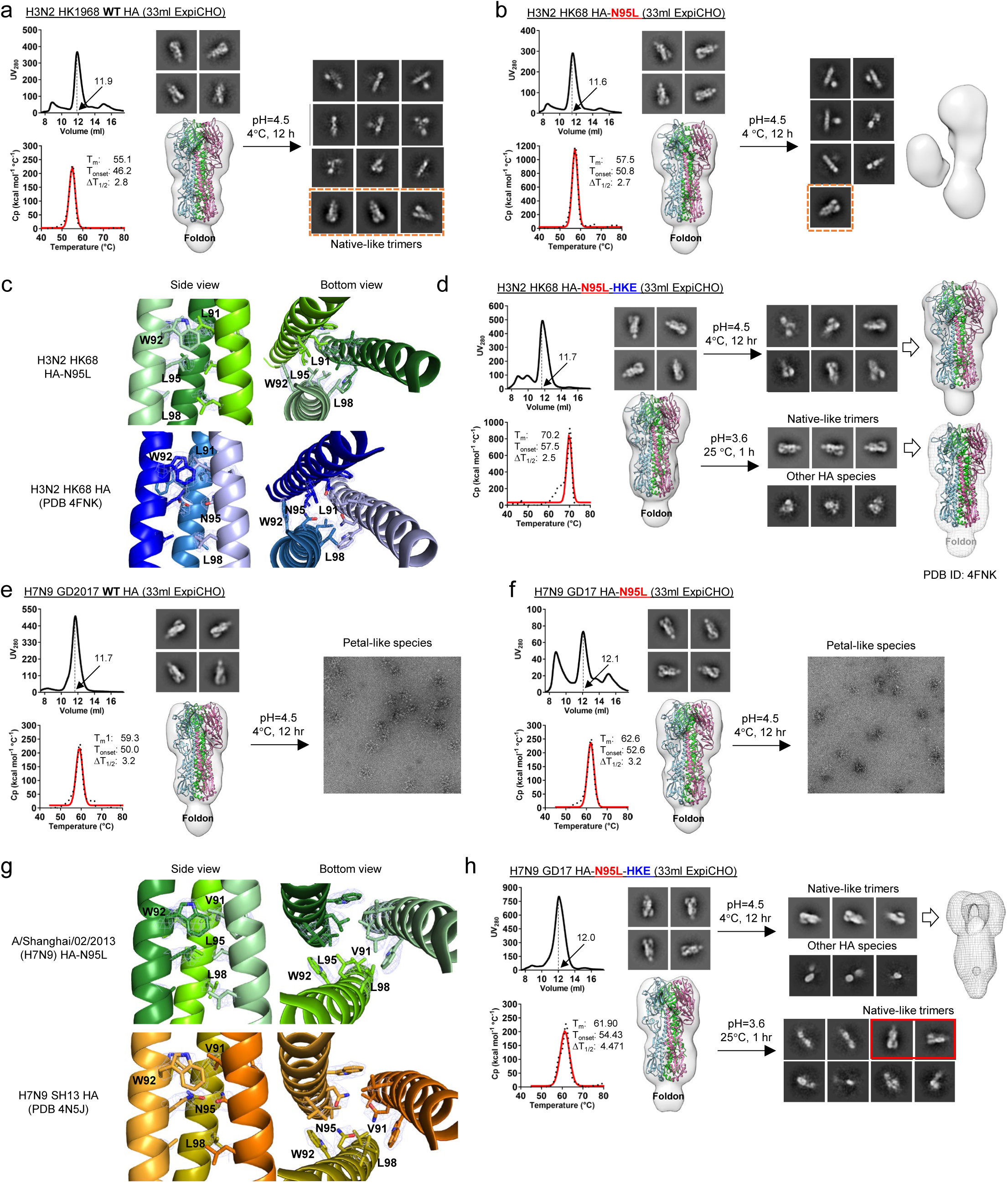
Rational design and in vitro characterization of stable prefusion-closed HA trimers for group 2 influenza A viruses. (**a-d**) In vitro characterization of A/Hong Kong/1/1968(H3N2) (HK68) WT HA (**a**), HA-N95L (**b**), and HA-N95L-HKE (**d**) constructs, and structural comparison between WT HA and HA-N95L (**c**). SEC profiles, DSC thermograms, and nsEM 2D/3D analyses are shown as in Fig. 1b**-**d. In the structural comparison, central helix regions surrounding residue 95 are shown in ribbon models for HA-N95L (top) and WT HA (PDB 4FNK; bottom), with side and top views. Side chains for L91, L95/N95, and L98 are shown in density maps. (**e-h**) In vitro characterization of A/Guangdong/Th005/2017 (GD17) WT HA (**e**), HA-N95L (**f**), and HA-N95L-HKE (**h**) constructs, and structural comparison between SH13 WT HA and SH13 HA-N95L (**g**). SEC, DSC, and nsEM analyses are shown as in Fig. 1b–d. In (**g**), ribbon models of the central helix region surrounding residue 95 are shown for HA-N95L (top) and WT HA (PDB 4N5J; bottom), in both side (left) and top (right) views. Density maps are shown to highlight side chains of L91, L95/N95, and L98. In (**c**) and (**g**), HA-N95L is shown in shades of green, while WT HA is shown in shades of blue for H3 and orange for H7. All group 2 HA trimers were transiently expressed in ExpiCHO cells and purified using a nickel column.

Three GD17 H7 HA constructs—WT, N95L, and N95L-HKE—were characterized using the same approach. Side-by-side comparisons of WT HA and HA-N95L revealed similarities to H3 and features unique to H7 (**Fig. 2e, f**). In both subtypes, the N95L mutation slightly improved the T_m_ while preserving the prefusion-closed conformation. Unique to H7 HA, however, the leucine substitution at position 95 markedly reduced expression yield and increased aggregation, accompanied by additional dimer and monomer peaks. nsEM analysis after mild acidic treatment (pH 4.5, 4 °C, overnight) revealed a previously unreported low-pH-induced species for both WT HA and HA-N95L, exhibiting a symmetric, petal-like architecture (**Fig. 2e, f**, right). To examine the structural impact of the N95L mutation on H7 HA, we selected A/Shanghai/2/2013 (H7N9) (SH13), a strain isolated during an earlier H7N9 outbreak, and determined the crystal structure of SH13 HA-N95L at 2.9 Å resolution (**Fig. 2g**). Compared with WT HA (PDB 4N5J) (*116*), which displayed interprotomer C_β_–C_β_ distances of 5.9, 7.8, and 9.1 Å for V91, N95, and L98, respectively, HA-N95L yielded 6.0, 7.8, and 9.6 Å—suggesting only modest displacement—and an overall C_α_ RMSD of 0.4 Å. Notably, incorporation of the HKE mutations reversed the negative impact of the N95L mutation on H7 HA expression but slightly reduced the T_m_ and structural cooperativity during thermal unfolding, as reflected by an increased ΔT_1/2_ (**Fig. 2h**, left). H7 HA-N95L-HKE retained the prefusion-closed trimer (**Fig. 2h**, middle) and, similar to the H3 counterpart, exhibited partial resistance to low pH, showing a mixture of prefusion-closed trimers and bugle-shaped—but not petal-shaped—particles in nsEM analysis (**Fig. 2h**, right).

Based on the results from IAC purification of H1 CA09 and H5 VN04 HA, we evaluated bNAb-based IAC methods for the H3 and H7 constructs. We first examined the utility of the bNAb C05, which targets the RBS using a 24-amino-acid heavy-chain complementarity-determining region 3 (HCDR3) and neutralizes both group 1 (H1, H2, H9) and group 2 (H3) IAVs (*115*) (**Fig. S2d**). A crystal structure of C05 bound to HK68 HA has been reported (*115*). Here, we prepared a C05-based IAC column to purify all three HK68 HA constructs—WT, N95L, and N95L-HKE. Similar to 2D1-purified CA09 HA (**Fig. S1c-f**), C05-purified HK68 HAs exhibited a prominent trimer peak, a small aggregation peak, and negligible dimer/monomer species. nsEM confirmed that C05-purified HAs maintained a prefusion-closed trimer. We next evaluated the bNAb F045-092, which has a 23-amino-acid HCDR3 loop and neutralizes H3N2 strains spanning 1968-2011 (*117*) (**Fig. S2e**). Comparable SEC profiles were observed for C05 and F045-092-purified samples, and WT HK68 HA purified via F045-092 retained the prefusion-closed trimer conformation. Lastly, we tested bNAbs CR9114 (*118*) and FluA-20 (*119*), which recognize the conserved stem and the trimer interface of the HA head domain, respectively, for IAC purification of GD17 H7 HA-N95L. IAC with CR9114 produced a low but detectable trimer peak in SEC (**Fig. S2f**), whereas FluA-20 failed to capture any HA protein, suggesting a tightly closed pre-fusion trimer that does not readily expose the trimer interface (**Fig. S2g)**. These results indicate that head-directed antibodies, rather than stem- or trimer interface-targeting NAbs, are more effective for IAC purification of the prefusion-closed HA trimers.

### Rational design and stabilization of influenza B HA trimers

Before the COVID-19 pandemic, IBVs co-circulated annually with IAVs, contributing to seasonal epidemics (*120*). Since the pandemic, the Yamagata lineage has not been detected in global surveillance (*22*), likely due to widespread mask use, reduced travel, and other public health measures that suppressed transmission of human-specific IBVs. Although the Yamagata lineage may currently be extinct, its potential re-emergence cannot be ruled out and should be considered in vaccine design. Here, we examined HA design and stabilization strategies for two IBV strains: B/Brisbane/60/2008 (BB08; Victoria lineage) and B/Florida/4/2006 (FL06; Yamagata lineage). Sequence analysis revealed that residue Q95 is conserved across all IBV HA sequences, analogous to N95 in IAVs (**Fig. 1a**, left). We therefore hypothesized that Q95L could stabilize IBV HA, either alone or in combination with a double mutation (N51L/S103L, or NS) inspired by the pHS2 mutation, which was shown to stabilize HAs across IAV subtypes and strains (*73*).

To test this hypothesis, we designed six soluble IBV HA constructs—WT, Q95L, and Q95L-NS—for both BB08 and FL06, expressed them transiently in ExpiCHO cells, and purified the proteins using a nickel column. In vitro characterization revealed similar biophysical properties between the two IBV lineages. WT HAs from BB08 and FL06 produced comparable SEC profiles, each with a trimer peak at 11.5–11.7 ml flanked by aggregation and monomer/dimer peaks, resembling those of IAV HAs (**Fig. 3a,b**, left). Both WT HAs exhibited T_m_ values around 60 °C, T_onset_ values of 55–56 °C, and prefusion-closed trimer conformations by nsEM (**Fig. 3a,b**, left). The Q95L mutation had mixed effects (**Fig. 3a,b**, middle): it increased the T_m_ by ∼5 °C for BB08 but reduced HA yield, particularly for FL06. Nonetheless, both constructs retained the prefusion-closed trimer. Neither construct, however, withstood mild acidic challenge, exhibiting both bugle-shaped intermediates and postfusion species by nsEM (**Fig. S3b**). The NS mutation appeared to compensate for the reduced protein expression introduced by Q95L while maintaining enhanced thermostability and structural integrity (**Fig. 3a,b**, right). However, HA-Q95L-NS constructs still transitioned to postfusion forms under mild acidic conditions, indicating that these stabilizing mutations provide only limited resistance to low-pH-induced conformational changes (**Fig. 3a,b**, rightmost). As with IAV HAs (**Fig. S1g-l**; **Fig. S2d-g**), we evaluated the utility of bNAbs, in this case CR8071 and CR8033, for IAC purification of BB08 and FL06 HAs, respectively (**Fig. 3c,d**). Although both antibodies target the HA head, CR8071 binds the conserved vestigial esterase domain at the base of the head with a perpendicular angle of approach, whereas CR8033 recognizes an epitope overlapping the RBS (*118*). IAC-purified samples showed SEC profiles similar to those obtained by nickel purification but yielded significantly less protein, likely due to weak antibody binding. Nonetheless, nsEM confirmed the exclusive presence of prefusion-closed HA trimers.

**Fig. 3.**
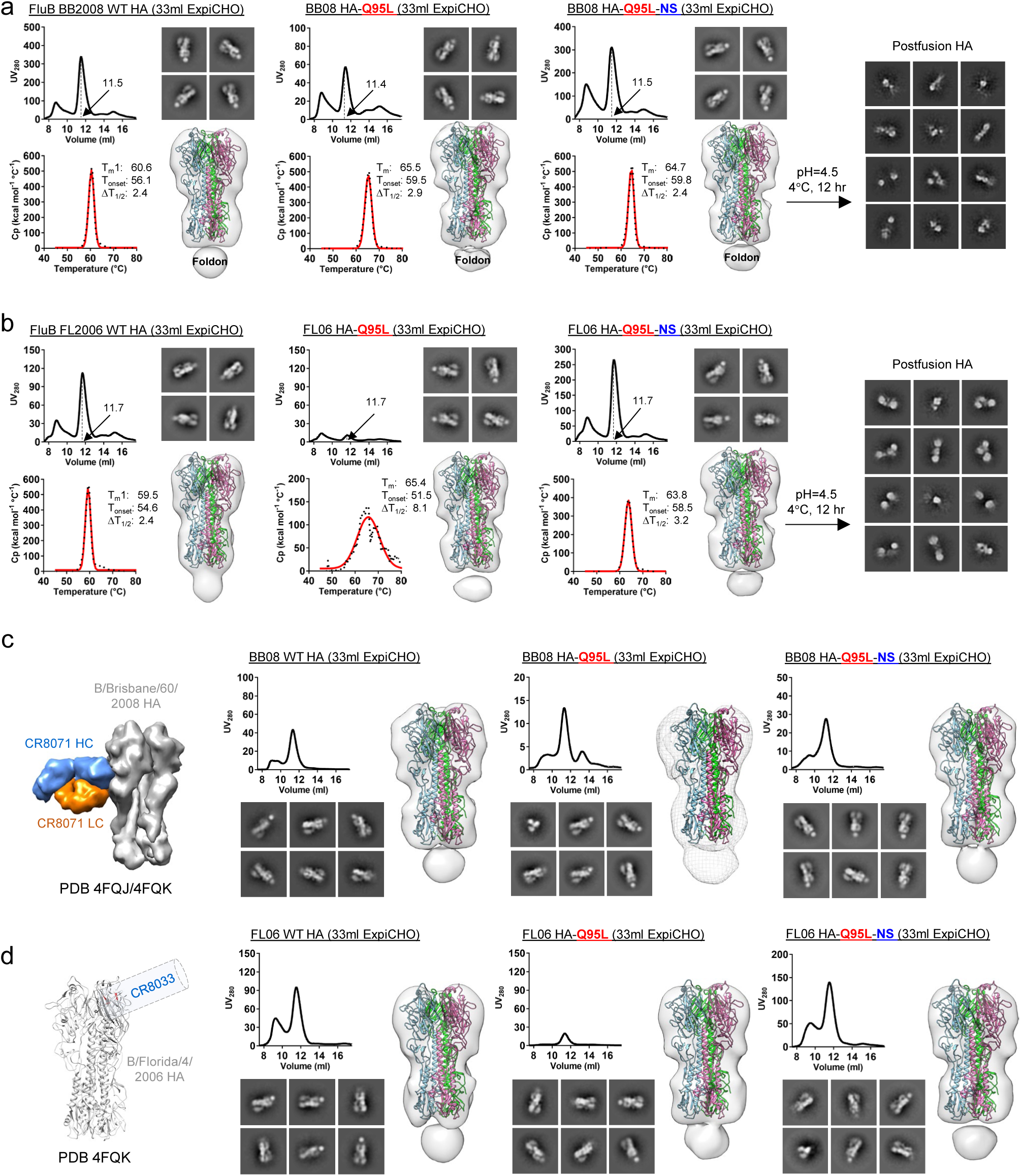
In vitro characterization of stable prefusion-closed HA trimers for influenza B viruses. (**a**) Characterization of B/Brisbane/60/2008 (Victoria lineage; BB08) WT HA (left), HA-Q95L (middle), and HA-Q95L-NS (right). (**b**) Characterization of B/Florida/4/2006 (Yamagata lineage; FL06) WT HA (left), HA-Q95L (middle), and HA-Q95L-NS (right). In (**a**,**b**), SEC profiles, DSC thermograms, and nsEM 2D class averages and 3D reconstructions are shown as in Fig. 1b**-**d. For HA-Q95L-NS, nsEM analysis after relatively mild acidic challenge (pH 4.5, 4 °C, 12 h) is included. (**c**) Characterization of CR8071-purified BB08 WT HA (left), HA-Q95L (middle), and HA-Q95L-NS (right). (**d**) Characterization of CR8033-purified FL06 WT HA (left), HA-Q95L (middle), and HA-Q95L-NS (right). In (**c**,**d**), each panel includes a structural model depicting the angle of approach for the antibody used in IAC purification (left), followed by SEC profiles and nsEM 2D/3D analyses. Acidic challenge was not performed in these panels.

In summary, although N95 in IAV and Q95 in IBV are both conserved, their mutations exert distinct effects on HA expression, stability, and structure. These results also demonstrate that increased thermostability does not necessarily confer greater acid resistance, suggesting that the fusion trigger may differ between IAV and IBV. Finally, unlike HIV-1 (*78*), filoviruses (*82*), or HCV (*101*), for which one or two bNAbs are often sufficient for broad IAC purification, none of the antibodies tested here showed comparable utility across influenza HAs.

### Antigenicity of rationally designed HA trimers carrying wildtype and modified glycans

In our previous studies, enzyme-linked immunosorbent assay (ELISA) and biolayer interferometry (BLI) were used with panels of representative antibodies to evaluate the antigenicity of rationally designed immunogens (*78–83, 85, 86, 99–101, 121*). Here, we adopted a similar approach to assess the antigenic profiles of stabilized HA constructs derived from CA09 and HK68, using a panel of antibodies targeting either the head or the stem. Head-directed antibodies included F045-092 (*117*), C05 (*115*), S139/1 (*122*), 2D1 (*102*), and FluA-20 (*119*), all known for their broad reactivity across diverse IAV strains and/or subtypes. H5M9, an H5-specific head-directed NAb, served as a negative control (*123*). Stem-directed antibodies included cross-group bNAbs 16.a.26 (*124*), 429 B01 (*125*), MEDI8852 (*126*), and CR9114 (*118*), as well as group-restricted bNAbs CR6261 (group 1) (*97*) and CR8020 (group 2) (*127*). In total, 12 antibodies were assayed.

Half-maximal effective concentration (EC_50_) values were derived from ELISA binding curves for comparison. For the four CA09 HA constructs—WT, HA-ER, HA-N95L, and HA-N95L-HKE (*73*) (**Fig. 4a**, **Fig. S4a**)—head antibody binding varied by construct. F045-092 bound weakly to all, whereas C05 and S139/1 showed no reactivity except for S139/1, which recognized HA-N95L-HKE. 2D1 bound all constructs with high affinity, consistent with its reported potency against A/California/04/2009 and A/South Carolina/1/1918 H1N1 (*127*). FluA-20 (*119*) also bound robustly, while H5M9 showed no reactivity, as expected. Among stem-directed antibodies, 429 B01, MEDI8852, and CR9114 bound strongly to all four CA09 constructs, consistent with their reported neutralization of A/California/04/2009 (*125, 126, 128*). 16.a.26 bound moderately to WT HA and HA-N95L-HKE but weakly to HA-ER and HA-N95L. CR6261, a group 1-specific stem bNAb, also bound strongly, consistent with previous reports using CA09-derived HA (*129*). In contrast, CR8020, which targets group 2 stem, showed no binding to any CA09 construct. For HK68 HA variants—WT, HA-N95L, and HA-N95L-HKE (**Fig.4b**, **Fig. S4b**)—head-directed antibodies F045-092, C05, and S139/1 bound to all constructs, consistent with prior reports of their broad reactivity and potent neutralization across the H3 subtype (*115, 117, 119, 122*). FluA-20, a broadly reactive non-NAb (*119*), also bound robustly to all HK68 constructs, whereas 2D1, which recognizes a pandemic H1 epitope, and H5M9 showed minimal reactivity. Stem-directed bNAbs—16.a.26, 429 B01, MEDI8852, and CR9114—bound strongly to all HK68 constructs, consistent with their broad reactivity against group 2 HAs (*118, 124–126*). As expected, CR6261, a group 1 stem bNAb (*127*), showed no binding to HK68 HAs, as H3 belongs to group 2. Conversely, CR8020, which targets the group 2 stem, bound strongly to all three HK68 constructs.

**Fig. 4.**
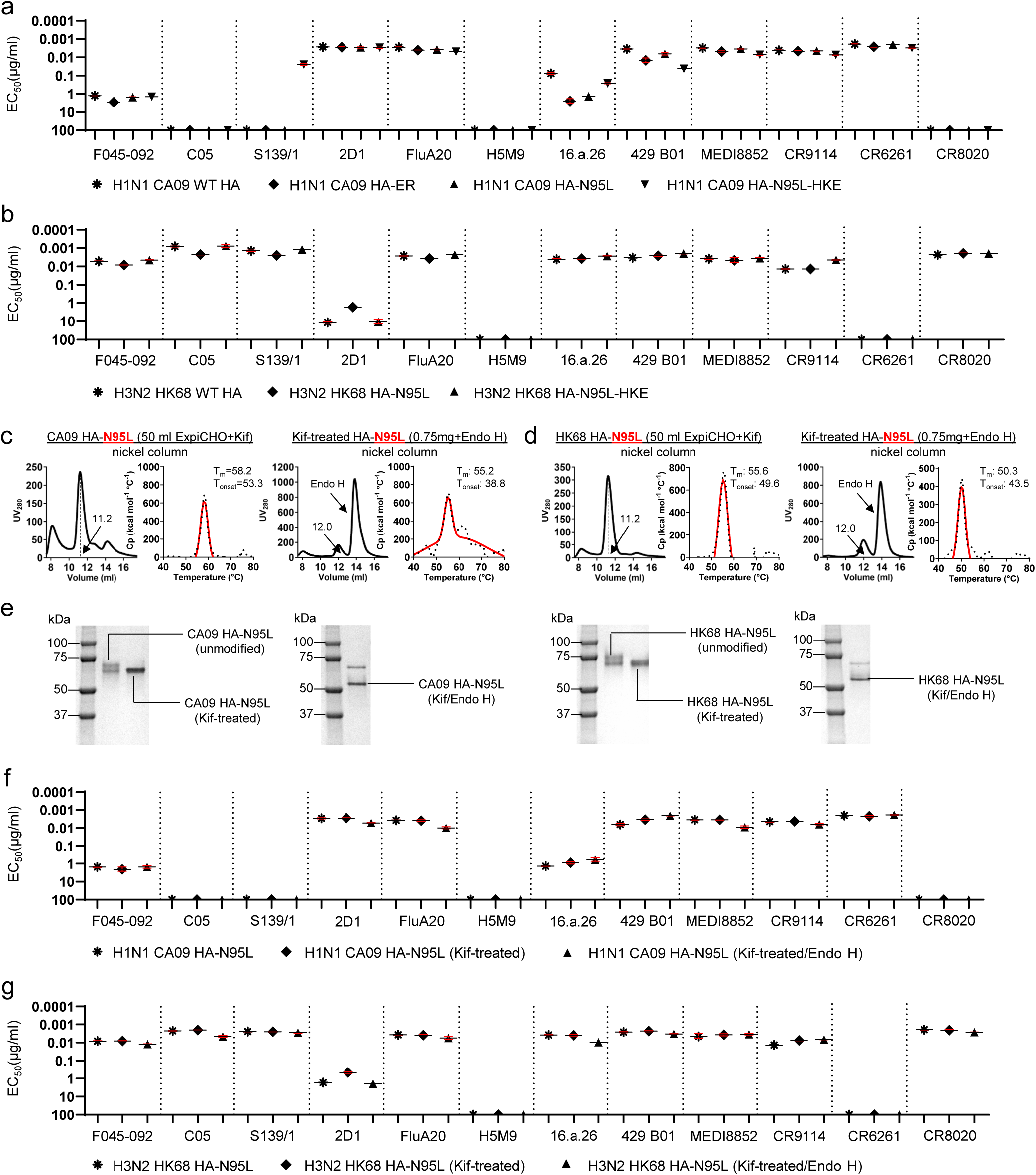
Antigenic and glycan analyses of wildtype and glycan-modified CA09 and HK68 HA trimers. (**a**) ELISA-derived EC_50_ (µg/ml) values of four CA09 HA constructs — WT HA, HA-ER, HA-N95L, and HA-N95L-HKE — binding to twelve antibodies targeting diverse antigenic sites on HA. (**b**) ELISA-derived EC_50_ (µg/ml) values of four HK68 HA constructs—WT HA, HA-N95L, and HA-N95L-HKE — binding to the same panel of twelve antibodies. (**c**) SEC profiles and DSC thermograms of Kif-treated (left) and Kif/endo H-treated (right) CA09 HA-N95L constructs. (**d**) SEC profiles and DSC thermograms of Kif-treated (left) and Kif/endo H-treated (right) HK68 HA-N95L constructs. (**e**) SDS-PAGE gels of unmodified, Kif-treated, and Kif/Endo H-treated HA-N95L derived from CA09 (left) and HK68 (right). Bands are labeled on the gels. (**f**) ELISA-derived EC_50_ (µg/ml) values for three CA09 HA-N95L constructs with distinct glycan statuses—unmodified, Kif-treated, and Kif/endo H-treated—against the same antibody panel as in (a). (**g**) ELISA-derived EC_50_ (µg/ml) values for three HK68 HA-N95L constructs with different glycan modifications, as in (f). In (**a, b, f, g**), the antibody panel includes head-specific antibodies F045-092, C05, S139/1, 2D1, FluA-20, and H5M9 (H5-specific); stem-directed antibodies 16.a.26, 429 B01, MEDI8852, and CR9114; and group-restricted antibodies CR6261 (group 1) and CR8020 (group 2). If the optical density at 450 nm (OD) is <0.5 at the starting and highest antibody concentration (10 μg/ml), binding is considered negligible, and the EC_50_ was assigned a value of 100 μg/ml to enable plotting and comparison. Error bars indicate the difference between duplicates for each concentration tested. All HA proteins were transiently expressed in ExpiCHO cells and purified using a nickel column.

Glycans are essential to immune recognition of HA and contribute to the emergence of pandemic influenza strains by modulating antigenicity and host adaptation (*130*). We previously demonstrated that glycan modifications exert distinct effects on antigenicity and immunogenicity depending on the virus (*78, 82, 121*). Here, we used kifunensine (Kif) to enrich oligomannose-type glycans and endoglycosidase H (Endo H) to trim them to a single N-acetylglucosamine (GlcNAc) residue at each N-glycan site. For CA09 HA-N95L, Kif treatment increased the trimer-to-non-trimer ratio in SEC while maintaining overall thermostability (**Fig. 4c**, left). Subsequent Endo H digestion shifted the trimer peak in SEC and reduced the T_m_ (**Fig. 4c**, right). Similar effects were observed for HK68 HA-N95L, although the loss of thermostability was more pronounced (**Fig. 4d**). Sodium dodecyl sulfate–polyacrylamide gel electrophoresis (SDS-PAGE) analysis confirmed molecular weight changes following glycan processing. For both CA09 and HK68 HA-N95L (**Fig. 4e**), Kif-treated samples produced slightly lower but sharper bands than unmodified samples, indicative of increased glycan uniformity, whereas subsequent Endo H trimming yielded a band just above 50 kDa, close to the theoretical molecular weight (60.6 kDa) predicted from the amino acid sequence. Notably, for both constructs, a fraction of HA proteins remained untrimmed at room temperature, as indicated by a lighter band below 75 kDa. Overall, the antigenicity of Kif-treated and Endo H-trimmed HA-N95L samples was largely comparable to that of unmodified proteins, as assessed for CA09 (**Fig. 4f, Fig. S4c**) and HK68 (**Fig. 4g, Fig. S4d**), suggesting that glycan modifications had little adverse effect on antibody recognition.

### Rational design and in vitro characterization of HA trimer-presenting 1c-SApNPs

Multivalent display of full HA ectodomains, receptor-binding domains (RBDs), or stem domains presents a promising strategy to elicit broad and durable NAb responses (*131–133*). Both the 24-mer ferritin (FR) (*59–62*) and the two-component I53-dn5 nanoparticle (2c-NP) system (*63, 64*) have been used to present HA antigens, with several NP-based vaccine candidates advancing to clinical trials (*65–68*). Previously, we employed FR and two 60-mer 1c-SApNPs, E2p and I3-01v9, to display stabilized HIV-1 Env (*78*), filovirus GP (*82, 83*), and SARS-CoV-2 spike (*85*) as VLP vaccine candidates. These 1c-SApNPs are multilayered by design, incorporating an internal layer of locking domains (LDs) and a hydrophobic core formed by 60 PADRE epitopes (*85*), designated E2p-LD4-PADRE (L4P) and I3-01v9-LD7-PADRE (L7P). To improve presentation of trimeric antigens with narrow stalks, I3-01v9 was structurally optimized to generate I3-01v9b/c (*82*). Here, we evaluated stabilized HA trimers displayed on FR, E2p-L4P, and I3-01v9b/c-L7P for influenza vaccine development. To this end, CA09 and HK68 HA-N95L were selected as representative full HA antigens for 1c-SApNP display. HA was fused via its C terminus to the N terminus of each NP subunit, with no linker for FR and a single glycine (G) linker for E2p and I3-01v9b/c (**Fig. S5a**). Structural modeling estimated particle diameters of 38.4, 49.7, and 51.8 nm for FR, E2p, and I3-01v9b/c, respectively, measured at HA1-N159 (**Fig. 5a**). In total, six constructs—three based on CA09 and three on HK68—were generated for experimental evaluation.

**Fig. 5.**
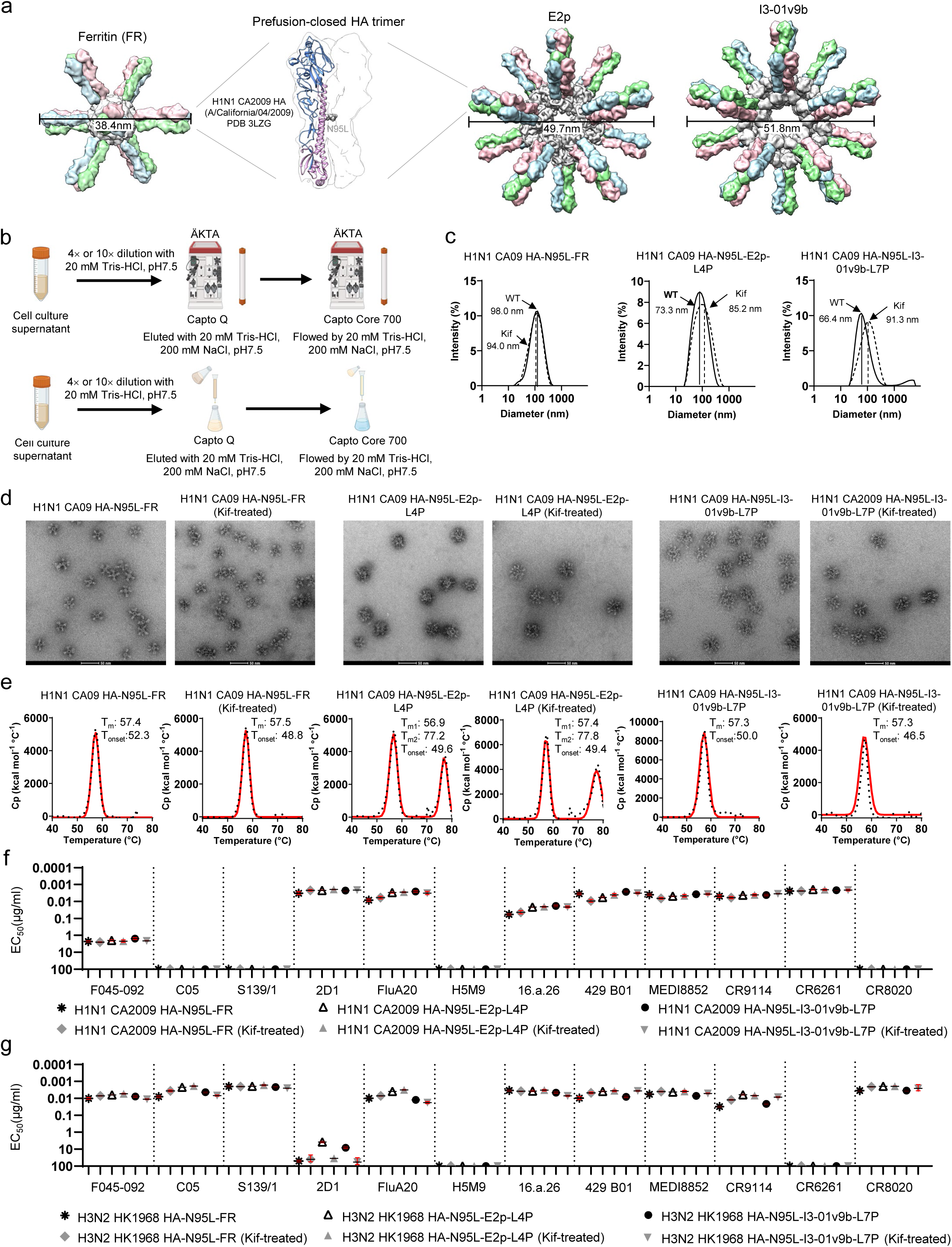
Rational design and characterization of HA trimer-presenting self-assembling protein nanoparticles. (**a**) Structural representation of a soluble HA trimer (center) and three HA trimer-presenting SApNPs—FR (left), E2p and I3-01v9b (right). Ribbon diagrams of the CA09 HA trimer (PDB 3LZG) are overlaid with transparent molecular surfaces, with HA1 in light blue and HA2 in plum. The N95L mutation is shown in gray. Surface representations are shown for the FR 24-mer and E2p/I3-01v9b 60-mers, displaying 8 and 20 HA trimers, respectively. The SApNP scaffold is shown in gray, with individual HA protomers colored distinctly. Particle diameters (nm) are indicated. (**b**) Schematic overview of the two-resin IEX purification process for HA-presenting SApNPs. The process can be implemented in either analytic mode (on an ÄKTA Pure system) or fast mode (via gravity). Key parameters are labeled. (**c**) Particle size distributions of unmodified (solid lines) and Kif-treated (dotted lines) CA09 HA-N95L FR, E2p-L4P, and I3-01v9b-L7P SApNPs. Hydrodynamic diameters (D_h_) were measured by DLS, with average particle sizes indicated on each plot. (**d**) Representative nsEM micrographs of unmodified and Kif-treated CA09 HA-N95L FR, E2p-L4P, and I3-01v9b-L7P SApNPs. Scale bar = 50 nm. (**e**) DSC thermograms of unmodified and Kif-treated CA09 HA-N95L FR, E2p-L4P, and I3-01v9-L7P SApNPs. Key thermal parameters are indicated on the thermograms. (**f**) ELISA-derived EC_50_ (µg/ml) values for unmodified and Kif-treated CA09 HA-N95L FR, E2p-L4P, and I3-01v9b-L7P SApNPs (six total) binding to twelve antibodies targeting diverse antigenic sites on HA. (**g**) ELISA-derived EC_50_ (µg/ml) values for unmodified and Kif-treated HK68 HA-N95L FR, E2p-L4P, and I3-01v9b-L7P SApNPs (six total) binding to the same panel of twelve antibodies. In (f) and (g), if the optical density at 450 nm (OD_450_) is <0.5 at the starting and highest antibody concentration (10 μg/ml), binding is considered negligible, and the EC_50_ was assigned a value of 100 μg/ml to enable plotting and comparison. Error bars indicate the difference between duplicates for each concentration tested. All HA proteins were transiently expressed in ExpiCHO cells and purified using a nickel column.

We first tested the feasibility of IAC for purifying HA-presenting SApNPs (**Fig. S5b**, left). The head-directed bNAb 2D1 efficiently purified CA09 HA-N95L displayed on FR, and to a lesser extent on E2p, but failed to capture HA-N95L on I3-01v9b SApNPs, as shown by SEC (Superose 6 10/300 GL) and nsEM (**Fig. S5b**). In contrast, C05, despite its success with the HK68 HA-N95L trimer, did not recover any of the corresponding SApNPs (**Fig. S5c**), suggesting that the repetitive HA display on the nanoparticle surface may hinder antibody-based IAC purification. To address this limitation, we developed an ion-exchange chromatography (IEX) method combining Capto Q resin for SApNP capture with Capto Core 700 (CC700) for removal of smaller impurities, operable in either pressure- or gravity-driven mode (**Fig. 5b**). Applied first to CA09 SApNPs, CC700 flow-through fractions showed high recovery for FR and comparable yields for E2p and I3-01v9b (**Fig. S5d**). IEX-purified samples were analyzed by dynamic light scattering (DLS), which showed hydrodynamic diameters (Dh) of 98.0/94.0 nm (unmodified/Kif-treated) for FR, 73.3/85.2 nm for E2p, and 66.4/91.3 nm for I3-01v9b (**Fig. 5c**), all larger than the diameters predicted by structural modeling. nsEM confirmed the purity and expected morphology of all three IEX-purified SApNPs: well-formed spherical cores decorated with evenly distributed prefusion-closed HA trimers (**Fig. 5d**). DSC analysis showed that particulate display and Kif treatment had minimal effects on HA thermostability, with < 2 °C reductions in T_m_ across constructs (**Fig. 5e**). The secondary melting peak at ∼77–78 °C in E2p thermograms likely corresponded to denaturation of the E2p core. For HK68 SApNPs, lower recovery was observed after the CC700 step (**Fig. S5e**). Nevertheless, nsEM verified the structural integrity of these particles (**Fig. S5f**). SDS-PAGE confirmed the expected molecular weight order (E2p-L4P > I3-01v9b-L7P > FR) for both CA09 and HK68 SApNPs, with Kif-treated samples consistently migrating above their unmodified counterparts (**Fig. S5g**).

The antigenicity of IEX-purified CA09 and HK68 HA-N95L SApNPs was evaluated by ELISA using the same antibody panel (**Fig. 5f, g; Fig. S5h, i**). Multivalent display produced similar effects on HA–antibody interactions across constructs. For CA09 HA-N95L, three head-directed NAbs—F045-092 (*117*), C05 (*115*), and S139/1 (*122*)—showed weak or no binding when HA was displayed on SApNPs, as did the group 2 stem-directed bNAb CR8020 (*127*). Likewise, HK68 HA-N95L SApNPs showed minimal recognition by the head-directed NAb 2D1 (*102*) and no binding to the group 1 stem-directed bNAb, CR6261 (*97*). Kif treatment modestly enhanced the antigenicity of CA09 SApNPs, reflected by ∼1.1–1.4-fold reductions in EC_50_ values for most NAbs. An exception was observed for the stem-directed bNAbs 429 B01 and MEDI8852 binding to FR, where EC_50_ values increased by ∼1.6–2.7-fold. Similarly, HK68 SApNPs exhibited greater binding affinities for nearly all NAbs, with EC_50_ values reduced by ∼1.0–2.6-fold compared with their unmodified counterparts. Overall, all neutralizing HA epitopes—though subtype specific in some cases—were well preserved on SApNPs, regardless of glycan modification.

In summary, stabilized HA trimers from diverse subtypes were successfully displayed on FR-, E2p-, and I3-01v9b/c-based SApNP platforms. Although IAC proved unsuitable for HA-presenting SApNPs, likely due to altered epitope accessibility, the IEX workflow provided a robust and scalable downstream purification process. Together, these findings demonstrate that N95L-stabilized HA trimers displayed on SApNPs possess biochemical, biophysical, structural, and antigenic properties that support their further evaluation in vivo.

### Distribution, trafficking, and retention of HA trimers and SApNPs in lymph nodes

Building on our previous analyses of 1c-SApNPs presenting HIV-1 Env, influenza M2e, SARS-CoV-2 spike, and Sudan virus (SUDV) GP (*78, 82, 84, 100*), we investigated the in vivo behavior of influenza HA trimers and HA-presenting FR and E2p SApNPs. Our goal was to understand how these vaccine antigens interact with cellular components in lymph nodes and initiate adaptive immune responses. To elicit robust humoral immunity, antigens must be transported through the lymphatic system and accumulate in sentinel lymph nodes, where they encounter B cells and activate them by engaging B cell receptors (BCRs) within lymphoid follicles (*134–136*).

We first examined the accumulation and distribution of HA-presenting E2p SApNPs in lymph nodes. Mice were intradermally injected via four footpads (10 μg per footpad), and sentinel lymph nodes were collected at 12 and 48 h after a single-dose immunization. To detect HA antigens, lymph node sections at 48 h were stained with a panel of human NAbs: FluA-20, which targets the HA head (*119*), and two stem-directed bNAbs, MEDI8852 (*118*) and CR9114 (*126*) (**Fig. S6a**). Among these antibodies, MEDI8852 produced the highest signal-to-noise ratio in immunohistological staining and was therefore selected for subsequent studies of HA-based immunogen trafficking and retention. Consistent with previous findings with SApNPs displaying SARS-CoV-2 spike, HIV-1 Env, influenza M2e, and SUDV GP (*78, 82, 84, 100*), HA-presenting SApNPs localized predominantly to the centers of lymph node follicles at both 12 and 48 h post-injection (**Figs. 6a,b**; images on the left, schematics on the right).

**Fig. 6.**
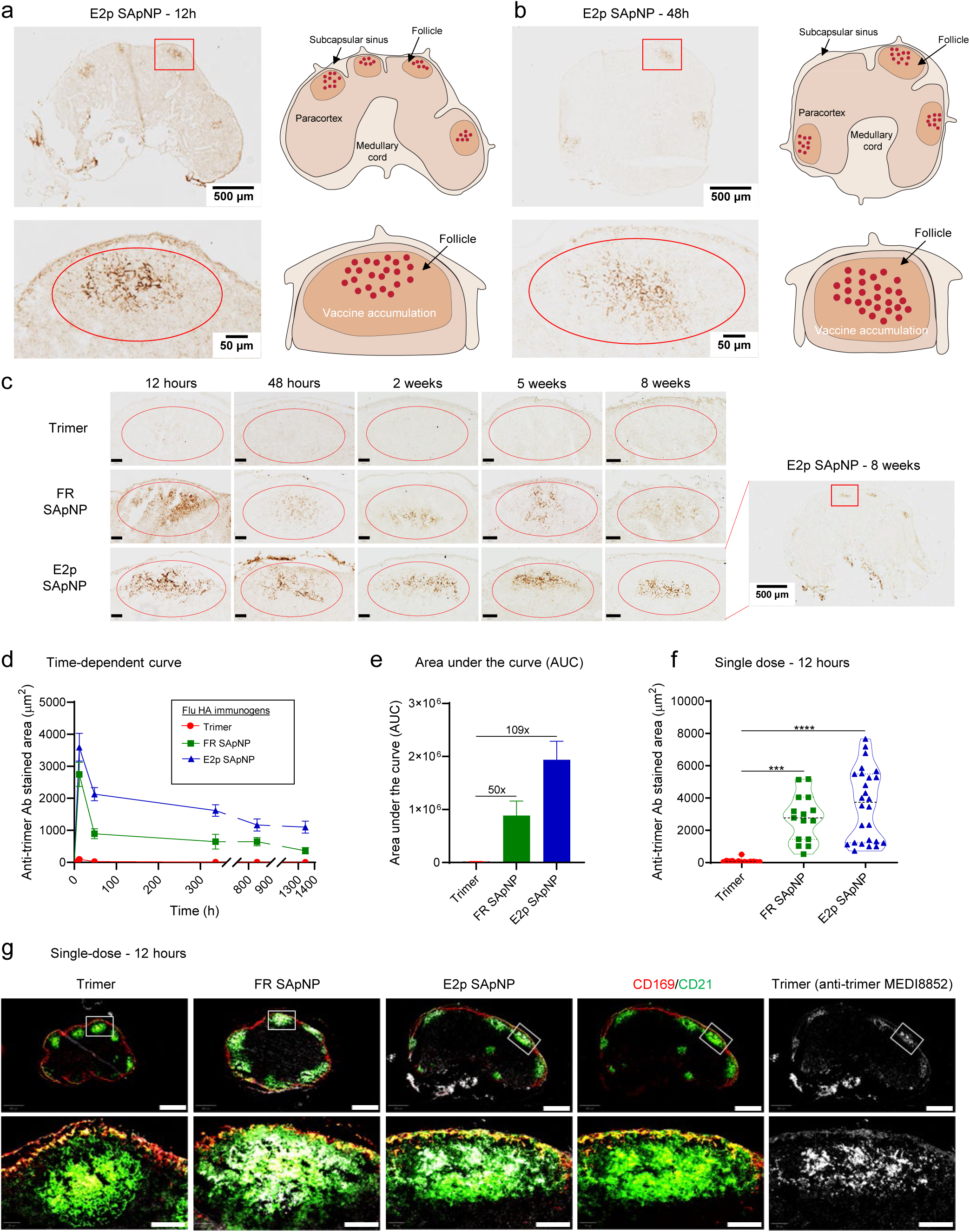
Prolonged retention of influenza HA-presenting SApNPs in lymph node follicles. Distribution of HA trimer-presenting E2p SApNPs in a lymph node at (**a**) 12 h and (**b**) 48 h after a single-dose injection (10 μg/injection, 40 μg/mouse, left). The administrated dose consisted of 80 μl of antigen/AddaVax (AV) adjuvant mixture containing 40 μg of SApNP immunogen and 40 μl of AV adjuvant. The HA stem-specific NAb MEDI8852 was used to stain the sentinel lymph node tissue sections. A schematic illustration of HA-presenting SApNP accumulation in lymph node follicles is shown (right). (**c**) Trafficking and retention of the HA vaccines in lymph node follicles from 12 h to 8 weeks after a single-dose injection. Scale bar = 50 μm for each image. (**d**) Time-dependent curve and (**e**) area under the curve (AUC) of the MEDI8852-stained area in immunohistological images depicting HA vaccine retention in lymph node follicles over 8 weeks (n = 3-5 mice/group). (**f**) Quantification of HA vaccine accumulation in lymph node follicles at 12 h after a single-dose injection. (**g**) Association of HA trimers and trimer-presenting SApNPs with FDC networks in lymph node follicles at 12 h after a single-dose injection. HA trimer and both SApNP vaccines were colocalized with FDC networks. Immunofluorescent images are pseudo-color-coded (CD21^+^, green; CD169^+^, red; MEDI8852, white). Scale bars = 500 and 100 μm for a complete lymph node and the enlarged image of a follicle, respectively. Data points are shown as mean ± SEM for (d) and SD for (e) and (f). Statistical analysis was performed using one-way ANOVA followed by Tukey’s multiple comparison *post hoc* test. ***p < 0.001, ****p < 0.0001.

Next, we investigated the trafficking and retention of HA trimers and SApNPs in lymph node follicles over an 8-week period following a single-dose injection (4 footpads, 10 μg/footpad) (**Fig. 6c**). Immunohistological analyses showed that antigens reached draining lymph nodes and initially accumulated in the subcapsular sinus within 12 h (**Fig. 6c**). Soluble HA trimers and SApNPs, however, exhibited distinct trafficking patterns: trimers entered follicles within 12 h and were completely cleared by 48 h, whereas FR and E2p SApNPs were visualized in follicles at 12 h and remained detectable for at least 8 weeks (**Fig. 6c**). Since the soluble HIV-1 trimer showed markedly higher accumulation at 2 h than 12 h (*78*), the smaller HA trimer was likely cleared more rapidly and therefore exhibited only weak signals at 12 h. Quantification of MEDI8852-stained areas over time revealed that FR and E2p SApNPs were retained ∼112-fold longer than soluble trimers (**Fig. 6c,d**). Area under the curve (AUC) analysis further showed that follicular exposure to SApNPs was 50–109 times greater than that of soluble trimers (**Fig. 6e**). At peak accumulation (12 h), FR and E2p SApNPs—which present 8 and 20 HA trimers, respectively—also exhibited 29–37 times higher accumulation compared with soluble trimers (**Fig. 6f**). These findings are consistent with our previous studies (*78, 82, 84, 100*), which showed that small soluble antigens are rapidly cleared from follicles, whereas large antigen-bearing SApNPs are retained for extended periods. Notably, SApNPs displaying influenza HA or M2e, HIV-1 Env, and SUDV GP remained detectable in follicles for ∼8 weeks, compared with ∼2 weeks for those presenting SARS-CoV-2 spike (*78, 82, 84, 100*). This prolonged retention appears to correlate with antigen thermostability: influenza HA or M2e, HIV-1 Env, and SUDV GP all exhibit T_m_ values >57 °C, while SARS-CoV-2 spike has a lower T_m_ of ∼48 °C, highlighting the importance of antigen stabilization.

FDCs form network structures in the centers of lymph node follicles, where they are essential for antigen retention and presentation to support robust B cell responses (*134–137*). Our previous studies with SARS-CoV-2 spike, HIV-1 Env, influenza M2e, and SUDV GP presented on SApNPs (*78, 82, 84, 100*), as well as earlier work with ovalbumin-conjugated gold NPs (*138, 139*), suggested that FDC networks may serve as the principal follicular component responsible for retaining HA SApNPs. To test this hypothesis, we intradermally injected the vaccine antigens and collected sentinel lymph nodes at the peak of SApNP accumulation (12 h) and additional time points from 48 h to 8 weeks (**Fig. 6g; Fig. S6b-e**). For immunohistological analysis, lymph node sections were stained with the bNAb MEDI8852 (*118*) for HA antigens, anti-CD21 (green) for FDCs, and anti-CD169 (red) for subcapsular sinus macrophages. At 12 h, immunofluorescent signals from both FR and E2p SApNPs showed strong colocalization with CD21 FDC networks (**Fig. 6g**), confirming that HA-presenting SApNPs associate with FDCs within follicles.

### Interaction of HA SApNPs with FDCs, B cells, and phagocytic cells in lymph nodes

FDC networks act as reservoirs that capture and preserve native-like antigens, immune complexes, particulate immunogens, viruses, and bacteria on their surfaces and dendrites through complement receptor-mediated mechanisms (*134–136, 140*). Our previous transmission electron microscopy (TEM) analyses showed that FDCs align SApNPs presenting SARS-CoV-2 spike, HIV-1 Env, and SUDV GP (*78, 82, 84*), as well as ovalbumin-conjugated gold NPs (*138, 139*), along their surfaces and dendrites. Here, we visualized the interface between FDC dendrites and B cells to examine how FDC networks process HA-presenting SApNPs formulated with AddaVax (AV), an oil-in-water emulsion adjuvant (∼150 nm), to engage B cells. Mice were intradermally injected with AV-formulated FR or E2p SApNPs (two footpads, 50 μg per footpad), and fresh sentinel lymph nodes were collected at 12 and 48 h post-injection. Tissues were fixed, sectioned, and processed for TEM. At both time points, TEM images revealed FDCs with long dendrites interacting with B cells in lymph node follicles (**Fig. 7a-c**). Intact SApNPs (round granules, yellow arrows) and AV particles (green arrows) were aligned along FDC dendrites and B cell surfaces (**Fig. 7a-c**; **Fig. S7a-d**). By 48 h, HA SApNPs were visible within B cell endolysosomes (**Fig. 7d**; **Fig. S7e, f**), consistent with findings for HIV-1 Env-presenting SApNPs (*78*). These observations highlight the role of FDC networks in retaining and presenting particulate immunogens on extended dendrites to maximize BCR engagement and promote robust B cell activation.

**Fig. 7.**
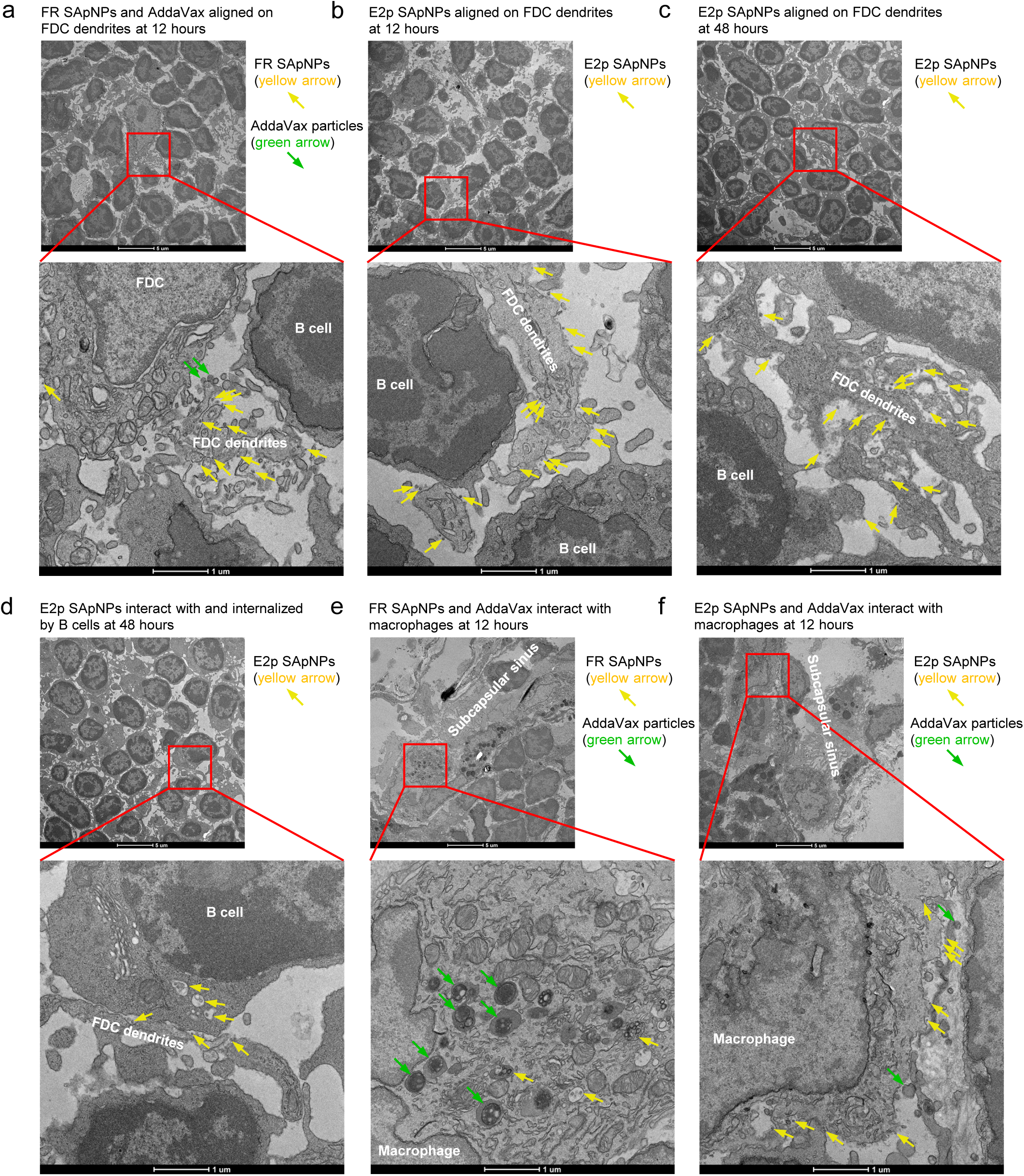
Interaction of HA trimer-presenting SApNPs with FDCs, B cells, and macrophages in lymph nodes. TEM images of HA trimer-presenting (**a**) FR SApNPs (yellow arrows) and (**b**, **c**) E2p SApNPs (yellow arrows), as well as AddaVax (AV) particles (green arrows), aligned along FDC dendrites in lymph node follicles at (**a**, **b**) 12 h and (**c**) 48 h after a single-dose injection (2 footpads, 50 μg/footpad). The administrated dose consisted of 200 μl of antigen/AV adjuvant mix, including 100 μg of SApNP and 100 μl of AV adjuvant, delivered into two hind footpads. (**d**) TEM images showing HA-presenting E2p SApNPs (yellow arrows) inside B cell endolysosomes at 48 h. TEM images of (**e**) FR and (**f**) E2p SApNPs (yellow arrows), along with AV adjuvant particles (green arrows), are shown on the surface or within endolysosomes of subcapsular sinus macrophages at 12 h post-injection.

Antigen-presenting cells, such as macrophages and dendritic cells, not only internalize and process vaccine antigens to support adaptive immunity but also transport intact antigens to migrating B cells and deposit them onto the FDC networks in their native conformation (*135, 136, 141–144*). To assess how adjuvanted SApNPs interact with subcapsular sinus macrophages, we examined the distribution of FR and E2p SApNPs formulated with AV. At 12 h post-injection, both HA-presenting SApNPs and AV adjuvant particles were visible on the surface and within endolysosomal compartments of macrophages (**Fig. 7e, f**), consistent with previous findings for SApNPs presenting SARS-CoV-2 spike, HIV-1 Env, and SUDV GP (*78, 82, 84*). These data indicate that HA-presenting SApNPs adjuvanted with AV are distributed across intercellular and intracellular compartments in lymph nodes. Both FR and E2p SApNPs exhibited similar associations with FDC networks, B cells, and macrophages. Notably, a substantial fraction of adjuvanted SApNPs was processed by macrophages, whereas FDCs retained both SApNPs and AV particles on dendrites to facilitate antigen presentation to B cells.

### Assessment of GC reactions induced by HA trimers and SApNPs in lymph nodes

B cell somatic hypermutation, selection, affinity maturation, and class switching occur during GC reactions, driving the formation of immune memory and NAb development after vaccination (*136, 137, 145, 146*). GC formation and maintenance depend on two critical cellular components: FDC networks, which retain and present antigens, and T follicular helper (T_fh_) cells, which support B cell maturation (*134, 147, 148*). Prolonged retention of HA-presenting SApNPs by FDC networks is expected to enhance the magnitude and durability of GC reactions compared with soluble HA trimers. To test this, we assessed GC responses after a single-dose immunization (four footpads, 10 μg per footpad) with adjuvanted HA-presenting SApNPs. Using established protocols (*78, 82, 84, 100*), we performed immunohistological analyses to identify GC B cells (GL7) and T_fh_ cells (CD4 Bcl6) in lymph node sections. At week 2 post-immunization, robust GCs (GL7^+^, red) were observed in close association with FDC networks (CD21^+^, green), with well-defined light zone (LZ) and dark zone (DZ) structures within B cell follicles (B220^+^, blue) (**Fig. 8a**, left). T_fh_ cells (CD4^+^Bcl6^+^, co-labeled cyan and red) were enriched in the LZ, consistent with their role in B cell maturation and GC maintenance (**Fig. 8a, right**). We next examined GC responses induced by HA trimers and SApNPs at weeks 2, 5, and 8 post-vaccination (**Fig. 8b; Fig. S8a-c**). GC size, defined as the occupied area within follicles, was used to quantify GC responses from immunohistological images (*78, 82, 84, 100*). All three adjuvanted HA immunogens induced robust GC formation, with the E2p SApNP group producing the largest GCs at week 2 after a single-dose immunization (**Fig. 8b; Fig. S8a**). GC size gradually declined across all groups, but both SApNPs induced GCs that were 2.0–2.8-fold larger than those elicited by HA trimers (**Fig. 8b, c**).

**Fig. 8.**
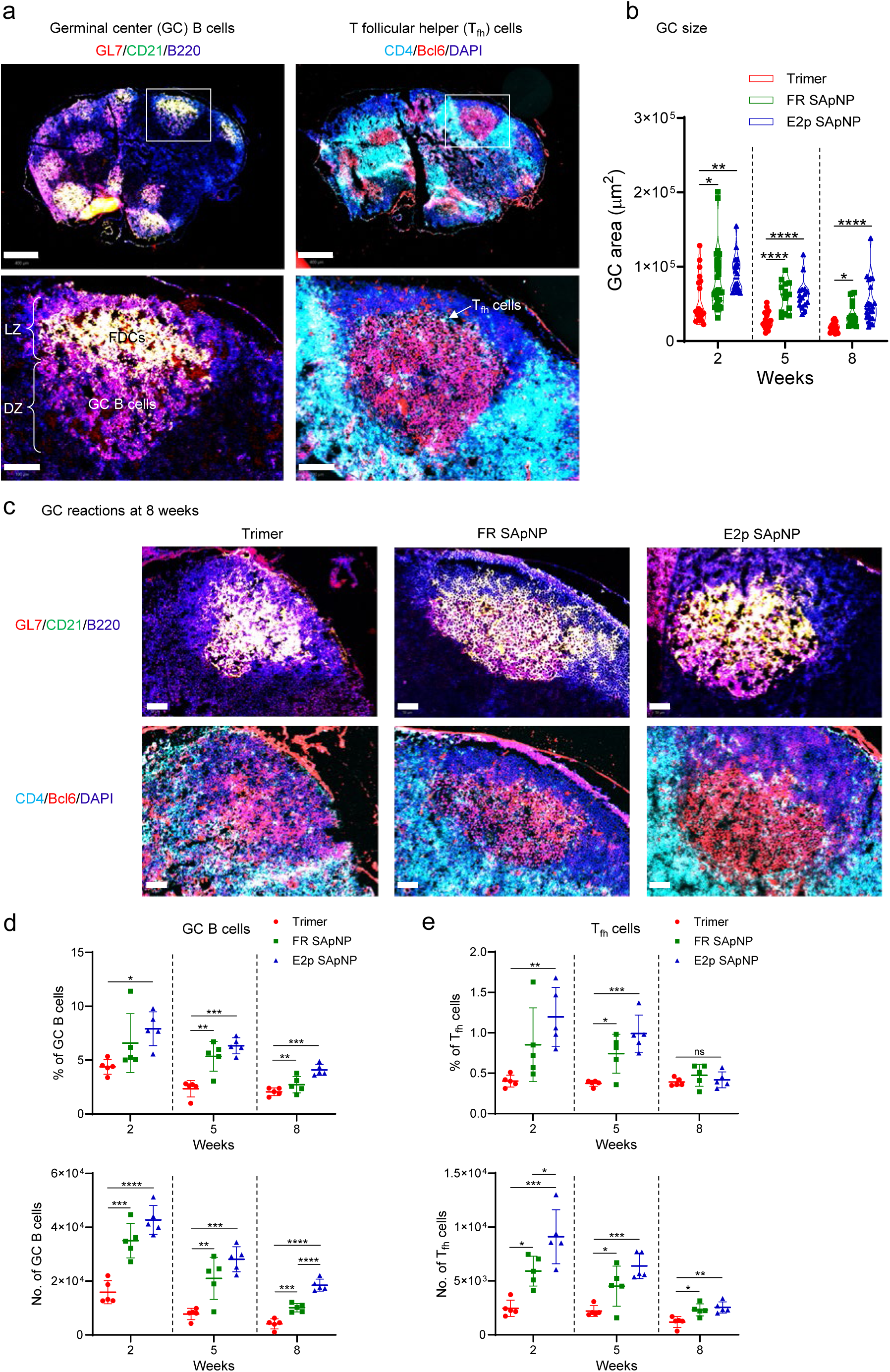
Induction of robust and long-lived germinal centers by HA trimer-presenting SApNPs in lymph nodes. (**a**) Top: Representative immunofluorescent images showing germinal centers (GCs) induced by HA trimer-presenting SApNP vaccines at 2 weeks after a single-dose injection (10 μg/injection, 40 μg/mouse). The administrated dose consisted of 80 μl of antigen/AddaVax (AV) adjuvant mix containing 40 μg of SApNP and 40 μl of AV adjuvant. Bottom: FDC networks (CD21^+^, green) associated with well-organized light zone (LZ) and dark zone (DZ) compartments in lymph node B cell follicles (B220^+^, blue). GC B cells (GL7^+^, red) and T_fh_ cells are primarily located in the DZ and LZ, respectively. Scale bars = 500 μm (whole lymph node) and 100 μm (enlarged follicle). (**b**) Assessment of GC size from immunofluorescent images at 2, 5, and 8 weeks after a single-dose injection (n = 5 mice/group). (**c**) Representative GC immunofluorescent images from HA vaccines at 8 weeks post-immunization. Scale bar = 50 μm (enlarged lymph node follicle). (**d**) Quantification of GC B cells and (**e**) T_fh_ cells by flow cytometry following a single-dose injection (n = 5 mice/group). Data are presented as mean ± SD. Statistical analysis was performed using one-way ANOVA followed by Tukey’s multiple comparison *post hoc* test for each time point. ns = not significant; *p < 0.05, **p < 0.01, ***p < 0.001, ****p < 0.0001.

We further assessed GC responses by flow cytometry. Following established protocols (*78, 82, 84, 100*), mice were immunized with three adjuvanted HA immunogens. Sentinel lymph nodes were collected at 2, 5, and 8 weeks after a single-dose injection (four footpads, 10 μg per injection) and processed into single-cell suspensions. Samples were stained with an antibody cocktail to identify GC B cells (B220^+^GL7^+^CD95^+^) and T_fh_ cells (CD3^+^CD4^+^CXCR5^+^PD1^+^) (**Fig. S9**). Frequencies and absolute numbers of GC B and T_fh_ cells were quantified. Flow cytometry analysis revealed that the E2p SApNP group elicited the highest frequencies and absolute numbers of GC B cells at week 2 (**Fig. 8d**), consistent with our immunohistological findings. Although GC B and T_fh_ cell populations declined over time in all groups, the SApNPs consistently outperformed the soluble trimer at every time point. At week 8, FR and E2p SApNPs induced 2.5–4.5-fold more GC B cells and 1.9–2.1-fold more T_fh_ cells compared to the HA trimer (**Fig. 8d,e**). Together, these results demonstrate that large, multivalent SApNPs elicit more robust and durable GC responses than soluble HA trimers, thereby promoting potent and long-lived humoral immunity.

### Antibody responses and protective efficacy of designed CA09 HA trimers and SApNPs

The immunogenicity and protective efficacy of CA09 HA-N95L trimers and SApNPs—carrying wild-type and modified glycans—were evaluated in BALB/c mice. Animals were immunized intradermally at weeks 0, 3, and 6 using a prime-boost regimen, with HA immunogens formulated in AddaVax (AV), and sera were collected at weeks 2, 5, and 8 (**Fig. 9a**). Serum binding antibody (bAb) responses were measured by ELISA using a CA09 HA-N95L(1TD0) probe incorporating an alternative trimerization motif (PDB 1TD0) to avoid foldon-specific reactivity. All groups mounted vaccine-matched bAb responses as early as 2 weeks after the first dose (**Fig. 9b; Figs. S10a,b**). EC_50_ titers increased by 15- to 556-fold following the second immunization and were further boosted or sustained after the third dose. All CA09 H1 HA-immune sera exhibited cross-activity to an HK68 H3 HA-N95L(1TD0) probe (**Fig. 9c; Fig. S10c**). Among these, unmodified and Kif-treated trimer groups elicited significantly higher EC_50_ titers than all SApNP groups, showing a distinct pattern of heterologous bAb induction. For vaccine-matched hemagglutination inhibition (HAI), SApNP groups demonstrated early onset of activity against CA09 H1N1, with detectable titers after a single immunization, whereas trimer groups showed no measurable HAI at the same time point (**Fig. 9d; Figs. S10d,e**). After the second dose, HAI titers increased across all groups, with SApNPs outperforming their trimer counterparts: 1,040-1,680 for SApNPs vs. 620 for trimers (unmodified), and 1,120-1,600 for SApNPs vs. 640 for trimers (Kif-treated). After the third dose, HAI titers in both unmodified and Kif-treated SApNP groups were further boosted or remained elevated. A similar trend was observed in trimer groups, although their titers remained lower than those elicited by the corresponding SApNPs. Against non-matched post-2009 H1N1 strains MI45 and WI22, sera from all groups showed cross-HAI activity, with SApNPs eliciting higher responder frequencies and greater titers than trimers (**Fig. 9d; Fig. S10f**). However, none of the groups generated detectable cross-HAI activity against pre-2009 H1N1 strains PR8 and NC99, indicating a lineage-restricted response. The Kif/Endo H-treated trimer elicited the lowest vaccine-matched bAb responses and the weakest HAI activity against the matched CA09 H1N1 across all time points, indicating that glycan trimming reduced HA-specific antibody responses.

**Fig. 9.**
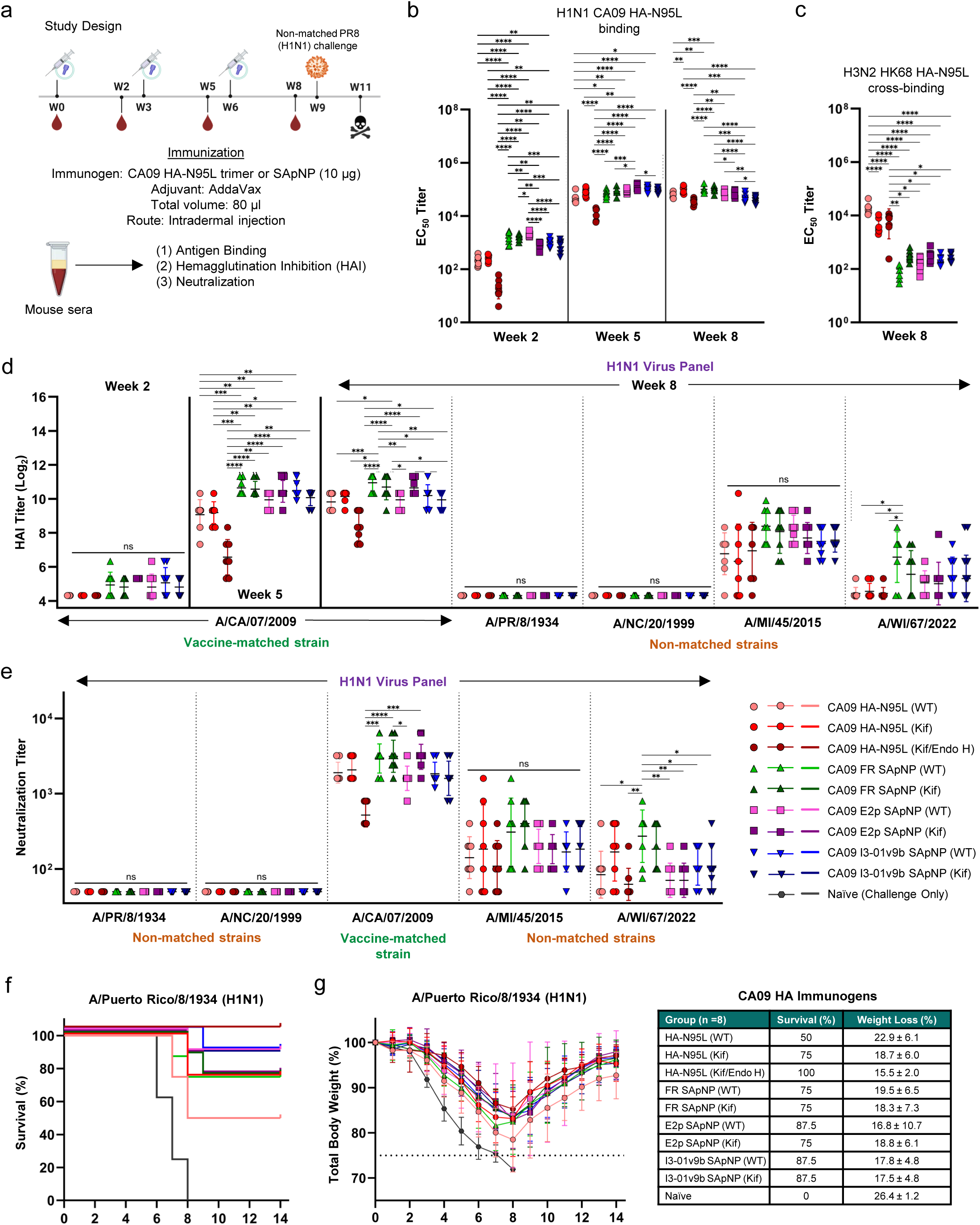
Antibody responses and protective efficacy of designed CA09 HA trimers and SApNPs. (**a**) Schematic of the mouse immunization and viral challenge schedule for CA09 H1 HA vaccines (n = 8 mice/group). Mice were administered 80 μl of antigen/AV adjuvant mix containing 10 μg of immunogen and 40 μl of AV. Mice were immunized at weeks 0, 3, and 6, and challenged at week 9. (**b-c**) CA09 H1 HA vaccine-induced serum binding antibody (bAb) responses against (**b**) vaccine-matched CA09 H1 HA N95L 1TD0 trimer and (**c**) non-matched HK68 H3 HA N95L 1TD0 trimer. EC_50_ values were derived from dose-response curves (OD_450_) generated against the coating antigens, with the calculated geometric means displayed on the plots. (**d**) CA09 H1 HA-induced sera hemagglutination inhibition against vaccine matched CA09, and non-matched PR8, NC99, MI15, and WI22 H1N1 viruses. HAI titers were defined as the highest serum dilutions at which complete hemagglutination inhibition was observed. (**e**) CA09 H1 HA-induced neutralizing antibody (NAb) responses against PR8, NC99, CA09, MI15, and WI22 H1N1 viruses. Neutralization titers were defined as the highest serum dilutions at which no viral infection was observed. (**f**) Survival and weight loss of CA09 H1 HA-immunized mice challenged with non-vaccine-matched, mouse-adapted LD_50_ × 10 of PR8 (H1N1). Mice were monitored for survival, weight loss, and morbidities for 14 days post-challenge. All data points are shown as mean ± SD. Data were analyzed using one-way ANOVA, with Tukey’s multiple comparison test used for *post hoc* analysis. Statistical significance is shown as the following: ns (not significant), \**p* < 0.05, \*\**p* < 0.01, \*\*\**p* < 0.001, and \*\*\*\**p* < 0.0001. The illustration of the mouse immunization study design was created using BioRender.com.

NAb responses were assessed using virus neutralization assays. All groups—except the Kif/Endo H-treated trimer—achieved average vaccine-matched neutralization titers exceeding 1,000 after three immunizations (**Fig. 9e; Figs. S10g,h**). Consistent with HAI results, all CA09 HA-N95L trimer and SApNP groups elicited cross-neutralizing activity against post-2009 H1N1 strains MI15 and WI22, with ∼98% and ∼71% of SApNP-immunized mice showing detectable neutralization of MI15 and WI22, respectively. In contrast, the Kif/Endo H-treated trimer group exhibited reduced cross-neutralization compared with the unmodified and Kif-treated trimers. No neutralizing activity against pre-2009 H1N1 strains PR8 and NC99 was observed in any group. Three weeks after the final vaccine dose, mice were challenged intranasally with 10× LD_50_ of PR8-ma (non-vaccine-matched H1N1) and monitored for survival and body weight loss over 14 days. The Kif/Endo H-treated trimer group achieved 100% survival (**Fig. 9f**), which might be mediated by T cell immunity and Fc-dependent, non-neutralizing but protective antibody functions. Groups immunized with unmodified E2p and I3-01v9 SApNPs, as well as Kif-treated I3-01v9 SApNPs, showed 87.5% survival, followed by the Kif-treated trimer, FR SApNPs (regardless of glycan treatment), and Kif-treated E2p SApNP at 75%. The unmodified trimer group exhibited the lowest survival (50%), while all naïve mice succumbed by day 8. Compared with their timer counterparts, SApNP groups demonstrated 25–37.5% higher survival for unmodified constructs and 0–12.5% higher survival for Kif-treated constructs. In terms of body weight loss, unmodified and Kif-treated SApNP groups showed slightly less (up to 6%) weight loss than their trimer equivalents (**Fig. 9g**). The least weight loss was observed in the Kif/Endo H-treated trimer group (∼16%).

### Antibody responses and protective efficacy of designed HK68 HA trimers and SApNPs

To evaluate the immunogenicity and protective efficacy of H3-based vaccine candidates, HK68 HA-N95L trimers and corresponding SApNPs—with or without glycan modification—were tested in BALB/c mice using the same immunization schedule as in the CA09 study (**Fig. 10a**). Serum bAb titers were measured by ELISA using a matched HK68 HA-N95L(1TD0) probe (**Fig. 10b; Figs. S11a,b**). All vaccine groups elicited detectable antibody responses by week 2 post-prime. These titers increased substantially (13- to 145-fold) after the second dose and either plateaued or rose modestly after the third dose. Heterologous binding was evaluated with a CA09 HA-N95L(1TD0) probe (**Fig. 10c; Fig. S11c**). As observed previously, unmodified and Kif-treated trimers induced significantly stronger cross-reactive bAb responses than their SApNP-displayed counterparts, suggesting that soluble HA trimers may offer a potential advantage through greater antibody accessibility to the stem region. HAI analysis of the HK68 vaccine groups recapitulated the trends observed for the CA09 series (**Fig. 10d; Fig. S11d**). Following the prime, most SApNP groups—except unmodified FR and E2p SApNPs—exhibited early HAI activity, while all trimer groups failed to elicit detectable titers. After the second dose, both trimer and SApNP groups showed marked increases in HAI titers; unmodified SApNP groups reached average titers of 240-440, compared with 130 for their trimer counterparts. After the third dose, unmodified SApNPs reached average titers of 520–720, again outperforming the ∼303 average observed for trimers. Interestingly, the Kif-treated trimer group achieved titers comparable to Kif-treated SApNP groups after the second dose (330 vs. 270-340) and outperformed all other trimer and SApNP groups after the third (920 vs. 580-720). In contrast, the glycan-trimmed (Kif/Endo H) trimer consistently produced the lowest HAI titers among the trimer groups. No cross-HAI activity was detected against the antigenically distant H3N2 strains BR07 and HK14, underscoring the subtype and lineage specificity of the vaccine-induced antibody responses (**Fig. 10d; Fig. S11e**).

**Fig. 10.**
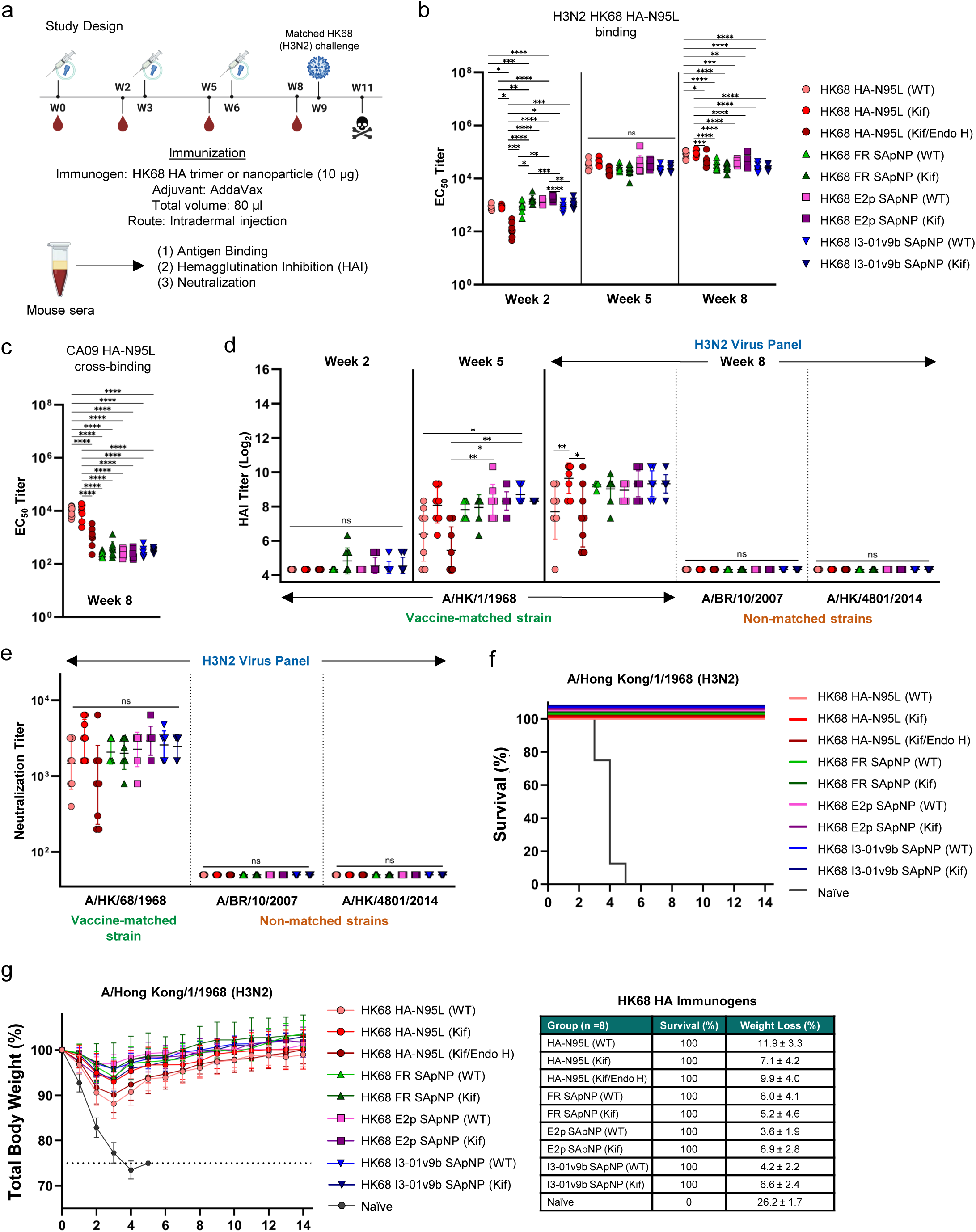
Antibody responses and protective efficacy of designed HK68 HA trimers and SApNPs. (**a**) Schematic of the mouse immunization and viral challenge schedule for HK68 H3 HA vaccines (n = 8 mice/group). Mice were administered 80 μl of antigen/AV adjuvant mix containing 10 μg of immunogen and 40 μl of AV. Mice were immunized at weeks 0, 3, and 6, and challenged at week 9. (**b-c**) HK68 H3 HA vaccine-induced binding antibody (bAb) responses against (**b**) vaccine-matched HK68 H3 HA N95L 1TD0 trimer and (**c**) non-matched CA09 H1 HA N95L 1TD0 trimer. EC_50_ values were derived from dose-response curves (OD_450_) generated against the coating antigens, with the calculated geometric means displayed on the plots. (**d**) HK68 H3 HA-induced sera hemagglutination inhibition against vaccine-matched HK68, and non-matched BR07 and HK14 H3N2 viruses. HAI titers were defined as the highest serum dilutions at which complete hemagglutination inhibition was observed. (**e**) HK68 H3 HA-induced neutralizing antibody (NAb) responses against HK68, BR07, and HK14 viruses. Neutralization titers were defined as the highest serum dilutions at which no viral infection was observed. (**f**) Survival and weight loss of mice challenged with vaccine-matched, mouse-adapted LD_50_ × 10 of HK68 (H3N2). Mice were monitored for survival, weight loss, and morbidities for 14 days post-challenge. All data points are shown as mean ± SD. Data were analyzed using one-way ANOVA, with Tukey’s multiple comparison test used for *post hoc* analysis. Statistical significance is shown as the following: ns (not significant), \**p* < 0.05, \*\**p* < 0.01, \*\*\**p* < 0.001, and \*\*\*\**p* < 0.0001. The illustration of the mouse immunization study design was created using BioRender.com.

In virus neutralization assays, all HK68 vaccine groups—except the Kif/Endo H-treated trimer—achieved average vaccine-matched titers exceeding 1,000 after three immunizations (**Fig. 10e; Fig. S11f**). While the unmodified trimer was less effective at eliciting autologous NAb responses, Kif treatment enhanced neutralization titers in the trimer group compared with the SApNPs, although the differences were not statistically significant. Consistent with HAI results, no cross-neutralization was detected against the heterologous H3N2 strains BR07 and HK14. To assess protective efficacy, mice were challenged with 10× LD_50_ of HK68-ma (vaccine-matched) and monitored for survival and weight loss over 14 days. All vaccinated groups achieved 100% survival (**Fig. 10f**), with an average weight loss of ∼7% across groups (**Fig. 10g**). Notably, mice immunized with unmodified and Kif-treated SApNPs showed 5.9–8.3% and 0.2–1.9% less weight loss, respectively, than their corresponding trimer groups. In contrast, naïve mice experienced ∼26% weight loss on average, with all succumbing to infection by day 5.

### In vivo assessment of stabilized HA trimers from other IAV subtypes and IBV

In addition to evaluating stabilized IAV H1 (CA09) and H3 (HK68) HA immunogens bearing the conserved N95L mutation, we also assessed the NAb responses elicited by stabilized HA trimers from IAV subtypes H5 and H7 and the IBV Victoria lineage. BALB/c mice were immunized with VN04 HA-N95L, GD17 HA-N95L, or BB08 HA-Q95L trimers using a similar prime-boost regimen: intradermal injections at weeks 0, 3, and 6 with AV-adjuvanted HA trimers, followed by serum collection at weeks 2, 5, and 8. The serum NAb response in the VN04 trimer group was measured against the vaccine-matched virus using a pseudovirus neutralization assay. Potent neutralizing activity was observed after the second dose and remained elevated following the third (**Fig. S12a**). A similar response pattern was observed for the GD17 trimer group (**Fig. S12b**). Sera from the BB08 trimer group exhibited robust HAI titers averaging 256 against the vaccine-matched strain (**Fig. S12c**). As expected, BB08 HA-immunized mice exhibited no cross-HAI activity against the non-matched FL06 strain from the Yamagata lineage and achieved only 40% survival after lethal FL06 challenge (**Fig. S12d**). These results confirm that the N95L and Q95L mutations can be extended beyond IAV H1 and H3 to generate stabilized HA immunogens for other IAV subtypes (H5 and H7) and IBV strains, eliciting strong immune responses upon vaccination.

## DISCUSSION

Vaccines are essential for influenza prevention, but their efficacy is often undermined by antigenic drift in circulating seasonal strains and the emergence of pandemic strains through antigenic shift (*131*). Inactivated vaccines remain the standard for seasonal influenza, and live attenuated vaccines are approved for use in specific populations. Despite continued efforts to improve these platforms (*30*), vaccine development is shifting from egg-based production to recombinant DNA, RNA, and protein formats that offer greater flexibility, efficiency, and scalability. Recent studies have identified conserved sites of vulnerability in viral antigens and incorporated potent adjuvants to enhance the breadth, potency, and durability of immune responses (*149*). The overarching goal is to develop universal vaccines that elicit balanced bNAb and T cell responses against diverse IAVs and IBVs—an advance that could transform global influenza prevention (*46, 150, 151*).

HA, the most abundant glycoprotein on the influenza virus surface, mediates viral entry by binding to host cell sialic acids to initiate fusion. As the primary target of NAb responses, HA is a major focus of influenza vaccine design, with renewed interest in NA as a complementary antigen (*152–154*). Owing to the high sequence variability of the HA globular head, most universal influenza vaccine efforts have focused on stem-based antigens. However, the isolation of diverse head-directed bNAbs that neutralize viruses across groups, subtypes, and types indicates that HA remains a viable target for broadly protective vaccine design. Despite this potential, relatively few studies over the past decade have addressed HA stabilization. One early approach introduced inter-protomer disulfide bonds to covalently lock HA in a rigid, closed trimer conformation (*72*), but this strategy may also promote postfusion six-helix bundle formation, as observed for PR8 H1N1 HA (*72*). More recently, a “repair and stabilize” approach was proposed (*73*), in which rare strain-specific HA residues are reverted to the subtype consensus, followed by the introduction of pHS1/2 mutations to enhance HA expression and acid resistance. This strategy increased HA expression by up to 2.5-fold in two H3N2 strains but required additional strain-specific mutations in others, limiting its applicability to a universal vaccine framework.

In this study, we hypothesized that a universal HA stabilization framework is essential for developing a broadly protective HA-based influenza vaccine. Informed by our previous filovirus and RSV studies (*82, 83, 86*), we analyzed central cavity-lining residues implicated in the open trimer conformation of CA09 HA (*88–90*). This analysis revealed a conserved motif across all IAV and IBV subtypes: a buried hydrophilic residue—N95 or Q95—flanked by hydrophobic stretches, suggesting a role in HA metastability. To test this, we designed and characterized HA constructs carrying the N95L (IAV H1, H3, H5, H7) or Q95L (IBV Victoria) substitution, either alone or in combination with pHS1/2 and other stabilizing mutations. Using biochemical, biophysical, and structural approaches, we evaluated the impact of these mutations on HA stability under neutral and acidic conditions. The N95L/Q95L substitution alone prevented trimer dissociation (CA09 HA), enhanced thermostability across multiple subtypes (IAV H3, H5, H7, and both IBV lineages), and, in some cases, increased acid resistance (H1 and H5). The addition of pHS1/2 mutations further improved these properties. Notably, while pHS2 was originally developed for IAV HAs (*73*), a pHS2-inspired NS mutation markedly improved expression and thermostability in IBV HAs. To assess multivalent display, CA09 and HK68 HA-N95L trimers were presented on ferritin as well as on multilayered E2p and I3-01v9b/c SApNPs, using a simple and robust IEX workflow for SApNP purification. Antigenic profiling revealed similar strain-, subtype-, and group-level constraints in antibody recognition for both trimers and SApNPs, suggesting an intrinsic immune barrier that is unlikely to be overcome by HA stabilization or multivalent display.

In vivo evaluation yielded two major findings. First, lymph node analyses revealed distinct behaviors of SApNPs versus soluble trimers. Multivalent display of stabilized HA trimers on FR and E2p SApNPs promoted strong interactions with FDC networks, B cells, and phagocytic cells. The large size and high thermostability of SApNPs enabled prolonged retention and presentation in lymph node follicles (over eight weeks), driving more robust and durable GC responses than soluble trimers. Although both HA trimers and SApNPs elicited strong antibody responses in mice, SApNPs did not show a significant advantage, likely due to reduced accessibility of certain HA epitopes on SApNPs and the short three-week interval used in the present immunization regimen. Second, glycan modification exerted complex effects on vaccine-induced immune responses. In line with our recent filovirus and HCV studies (*82, 121*), Kif treatment preserved antigenicity and modestly improved NAb responses compared with unmodified antigens. In contrast, our HIV-1 vaccine work showed that glycan trimming with Endo H enhanced responder rates and NAb induction (*78*), with similar benefits reported for glycan-trimmed influenza HA and SARS-CoV-2 spike vaccines (*155, 156*). Here, mice immunized with glycan-trimmed CA09 HA showed the highest survival following heterosubtypic H1N1 challenge, suggesting a protective advantage despite suboptimal HAI and NAb titers. Together, these findings underscore the multifaceted role of glycan modifications in shaping vaccine immunity and warrant further investigation.

Our future efforts will focus on several fronts. First, HA trimers and SApNPs incorporating both N95L/Q95L and pHS1/2 mutations will be evaluated in vivo and compared with constructs containing either mutation alone. Given their enhanced thermostability and acid resistance, the combined mutations are expected to induce more robust immune responses than individual ones. Second, this HA-based strategy could be integrated with M2e- and/or NA-based approaches to broaden protection (*99, 100*). We recently developed an SApNP vaccine displaying tandemly linked IAV and IBV M2e peptides, which conferred 100% and 70% protection against IAV and IBV challenge, respectively (*99*). By eliciting both potent NAb and T cell responses, an HA+M2e SApNP vaccine may offer broad protection against diverse influenza viruses, including pandemic strains. Third, the impact of glycan modifications warrants further investigation. A key distinction between this study and that of Chen et al. (*156*), which reported enhanced HAI and NAb titers and improved protection, lies in the expression system. Our HA constructs were expressed exclusively in ExpiCHO cells, while Chen et al. (*156*) produced HA trimers in HEK293 cells prior to glycan trimming. Comparative analyses across these and other expression hosts, such as insect cells used to produce the recombinant FluBlok vaccine (*157*) and the fungal C1 system (*158*), could provide critical insights into how glycosylation shapes vaccine-induced immunity.

## METHODS

### Construct design, expression, and purification of influenza HA immunogens

HA genes from group 1 IAVs—A/California/04/2009 (H1N1; GenBank: FJ966082) and A/Viet Nam/1203/2004 (H5N1; GenBank: AY818135)—and group 2 IAVs—A/Hong Kong/1/1968 (H3N2; GenBank: CY044261) and A/Guangdong/HP001/2017 (H7N9; GenBank: KY643843)— were used to assess the effects of stabilizing mutations N95L, R106M, E103L, H26W (pHS1 (*73*)), and K51I/E103I (pHS2 (*73*)), either introduced individually or in combination. Two IBV strains—B/Brisbane/60/2008 (GenBank: FJ766840, Victoria lineage) and B/Florida/4/2006 (GenBank: EU515992, Yamagata lineage)—were used to evaluate the effects of Q95L and pHS2-equivalent N398L/S450L (NS) mutations. For all soluble HA constructs, the ectodomains were truncated at residue 175 in HA2 (IAV) or residue 523 (IBV) and fused at the C-terminus to a foldon motif followed by a His_6_ tag for IMAC purification using a nickel column. For HA-presenting SApNPs, truncated HA was genetically fused to the NP-forming subunit—without a linker for the 24-meric FR and via a single glycine (“G”) linker for the 60-meric E2p and I3-01v9b platforms.

All HA trimers and HA-presenting SApNPs were produced in ExpiCHO cells (Thermo Fisher). Briefly, ExpiCHO cells were thawed and incubated in ExpiCHO^TM^ Expression Medium (Thermo Fisher) in a shaker incubator at 37 °C, at 135 rpm, with 8% CO_2_. When the cells reached a density of 10 × 10^6^ ml^−1^, ExpiCHO^TM^ Expression Medium was added to adjust the cell density to 6 × 10^6^ ml^−1^ for transfection. ExpiFectamine^TM^ CHO/plasmid DNA complexes were prepared following the manufacturer’s instructions. For glycan-modified constructs, kifunensine (10 mg/l, Tocris Bioscience) was added at the time of transfection to inhibit α-mannosidase I and generate oligomannose-type glycans. For a 100-ml transfection volume, 100 μg of HA antigen plasmid and 320 μl of ExpiFectamine^TM^ CHO reagent were mixed in 7.7 ml of cold OptiPRO™ medium (Thermo Fisher). After the first feed on day 1, ExpiCHO cells were cultured at 33 °C, at 115 rpm, with 8% CO_2_ according to the Max Titer protocol, with a second feed on day 5 (Thermo Fisher). Supernatants were harvested 13–14 days post-transfection, clarified by centrifugation at 3,724 × g for 25 min, and filtered through a 0.45 μm membrane (Thermo Fisher). All soluble HA trimers were purified using Ni Sepharose excel resin (Cytiva); bound proteins were eluted twice with 15 ml of 0.5 M imidazole and buffer-exchanged into Tris-buffered saline (TBS; pH 7.2). Several HA-specific antibodies were used to prepare affinity columns for IAC purification. HA trimers and SApNPs were eluted twice with 25 ml of 3 M MgCl_2_ (pH 8.0) into 25 ml of TN75 buffer (20 mM Tris-HCl, 75 mM NaCl), followed by buffer exchange into TN75. Nickel- or IAC-purified HA trimers and IAC-purified SApNPs were further polished by SEC using a Superdex 200 Increase 10/300 GL column and a Superose 6 Increase 10/300 GL column (Cytiva), respectively. Selected SEC fractions were flash-frozen in liquid nitrogen and stored at −80 °C until further use.

A two-resin IEX process was used to purify HA-presenting SApNPs. ExpiCHO culture supernatants were first clarified by centrifugation at 3,724 × g for 25 minutes and filtered through a 0.22 μm membrane (Millipore). For ÄKTA-based purification, the clarified supernatant was diluted four with 20 mM Tris-HCl buffer (pH 7.5) and applied to a 5 mL HiTrap Capto Q column (Cytiva) on an ÄKTA Pure 25 M system, with UV absorbance at 280 nm (UV_280_) monitored using a single-wavelength detector. The column was washed with 10 column volumes (CV) of 20 mM Tris-HCl, 20 mM NaCl, pH 7.5, and bound proteins were eluted with 4 CV of 20 mM Tris-HCl, 200 mM NaCl, pH 7.5, collected in 2 mL fractions. The fraction with the highest UV_280_ reading was loaded onto a 5 mL Capto Core 700 column (self-packed, Cytiva) and run in flow-through mode on the same system. SApNPs were recovered in the first CV of the flow-through. For gravity-based purification, 20 mL of clarified supernatant was diluted fourfold in 20 mM Tris-HCl (pH 7.5) and applied to a gravity flow column containing 3 mL of Capto Q resin (Cytiva). The column was washed twice with 10 CV of 20 mM Tris-HCl, 20 mM NaCl, pH 7.5, and eluted with 2 CV of 20 mM Tris-HCl, 200 mM NaCl, pH 7.5. The entire elution fraction was then applied to a second gravity column containing 3 mL of Capto Core 700 resin (Cytiva). The final flow-through was concentrated and buffer-exchanged into PBS using 100 kDa Amicon filters (Millipore). Purified SApNPs were flash-frozen in liquid nitrogen and stored at −80 °C until further use.

### Expression and purification of influenza antibodies

All antibodies were transiently expressed in ExpiCHO cells (Thermo Fisher). At 12-14 days post-transfection, the cultures were centrifuged at 3,724 × g for 25 minutes, and the supernatants filtered through a 0.45-μm membrane (Millipore). IgGs were purified using protein A affinity resin (Cytiva) and eluted with 0.3 M citric acid (pH 3.0). The pH of the eluate was immediately adjusted to 7.0 by adding 2 M Tris-Base (pH 9.0). Eluted antibodies were then concentrated and buffer-exchanged into phosphate buffered saline (PBS) using 10 kDa Amicon filters (Millipore). IgG concentrations were quantified by UV_280_ using theoretical extinction coefficients.

### Glycan trimming by endoglycosidase H (endo H) and enzyme removal

The protocol for endo H treatment and enzyme removal was adapted from our previous study (*78*). HA trimers and SApNPs were expressed in the presence of kifunensine (Kif) to enable glycan trimming by endo H. Briefly, 1 mg of HA antigen was mixed with 50 μl of 10× GlycoBuffer 3, 250 μl of endo-Hf (a fusion of endo H and MBP; NEB, catalog no. P0703L), and nuclease-free H_2_O (if necessary) to a final volume of 500 μl. The reaction volume was scaled proportionally as needed. The mixture was incubated at room temperature (25°C) for 5 h to allow enzymatic processing of HA glycans. Following incubation, the reaction mixture was subjected to SEC using a Superdex 200 Increase 10/300 GL column to remove the MBP-tagged endo H. SEC-purified fractions were flash-frozen in liquid nitrogen and stored at −80 °C until further use.

### SDS-PAGE and BN-PAGE

HA trimers and HA-presenting SApNPs were analyzed by sodium dodecyl sulfate-polyacrylamide gel electrophoresis (SDS-PAGE) and blue native-polyacrylamide gel electrophoresis (BN-PAGE). The proteins were mixed with loading dye and loaded on a 10% Tris-Glycine Gel (Bio-Rad) or a 4-12% Bis-Tris NativePAGE^TM^ gel (Life Technologies). Two conditions were used in SDS-PAGE. Under non-reducing conditions, the proteins were mixed with 6× Laemmli SDS sample buffer (Thermo Scientific, catalog no. J60660) and heated to 80 °C for 5 min. Under reducing conditions, the proteins were mixed with a different Laemmli buffer (Thermo Scientific, catalog no. J61337.AC) and boiled at 100 °C for 5 min. After loading, SDS-PAGE gels were run for 25 min at 250 V using SDS running buffer (Bio-Rad). SDS-PAGE gels were stained with InstantBlue (Abcam). For BN-PAGE, the proteins were mixed with 4× native dye and loaded on a BN-PAGE gel. The gel was run for 2–2.5 h at 150 V with NativePAGE^TM^ running buffer (Life Technologies) according to the manufacturer’s instructions. BN-PAGE gels were stained with Coomassie Brilliant Blue R-250 (Bio-Rad) and destained using a solution of 6% ethanol and 3% glacial acetic acid. Gel images were acquired using a ChemiDoc^TM^ XRS+ system and processed with Image Lab version 6.1.0 build 7 Standard Edition (Bio-Rad).

### Differential scanning calorimetry (DSC)

Melting temperature (T_m_) and other thermal parameters of HA trimers and SApNPs—expressed in ExpiCHO cells and purified by different methods—were determined using a MicroCal PEAQ-DSC Man instrument (Malvern). Nickel/SEC- and IAC/SEC-purified HAs and IEX-purified SApNPs were diluted in PBS to a final concentration of 0.5–5 μM. Thermal denaturation was performed at a scan rate of 60 °C/h from 20 °C to 100 °C. Data processing, including buffer correction, normalization, and baseline subtraction, was performed using MicroCal PEAQ-DSC software. Gaussian fitting was conducted using GraphPad Prism version 10.3.1.

### Baculovirus expression and purification of HA for crystal structure determination

The ectodomains of influenza HK68 H3 HA from A/Hong Kong/1/1968 (H3N2) and SH13 H7 HA from A/Shanghai/2/2013 (H7N9) with N95L mutation in HA2 domain were expressed and purified essentially as previously described (*159*). The cDNAs corresponding to the ectodomains (HA1 residues 11-329 and HA2 residues 1-176) of HK68 H3 HA and SH13 H7 HA with HA2 N95L mutation were incorporated into a baculovirus transfer vector, pFastbacHT-A (Invitrogen), with an N-terminal gp67 signal peptide, a C-terminal foldon trimerization domain and His_6_-tag, with a trypsin cleavage site incorporated to enable cleavage between the HA ectodomain and the foldon trimerization domain/His_6_-tag. The constructed plasmids were used to transform DH10bac competent bacterial cells to form a recombinant bacmid and the purified recombinant bacmids were used to transfect Sf9 insect cells for overexpression. HA protein was produced in suspension cultures of High Five cells with recombinant baculovirus at a multiplicity of infection (MOI) of 5-10 and incubated at 28 °C shaking at 110 RPM. After 72 h, High Five cells were removed by centrifugation and supernatants containing secreted, soluble HA protein were concentrated and buffer-exchanged into 20 mM Tris pH 8.0, 150 mM NaCl, and then further purified by metal affinity chromatography using Ni-nitrilotriacetic acid (NTA) resin (Qiagen). For crystal structure determination, the foldon trimerization domain and His6-tag were cleaved from the HA ectodomain with trypsin for HK68 H3 HA and thrombin for SH13 H7 HA and purified further by size exclusion chromatography on a Hiload 16/90 Superdex 200 column (GE healthcare) in 20 mM Tris pH 8.0, 150 mM NaCl and 0.02% NaN3. The purified HAs were quantified by optical absorbance at 280 nm, and purity and integrity were analyzed by reducing and nonreducing sodium dodecyl sulphate–polyacrylamide gel electrophoresis (SDS-PAGE).

### Crystal structure determination

Crystallization experiments were set up using the sitting drop vapor diffusion method using automated crystal screening on our robotic CrystalMation system (Rigaku). Diffraction quality crystals for HK68 H3 HA ectodomain trimer with HA2 N95L mutation were obtained by mixing 0.1 µL HA protein at 6.8 mg/ml in 20 mM Tris pH 8.0, 150 mM NaCl and 0.02% (v/v) NaN_3_ with 0.1 µL of the well solution in 0.2 M sodium formate, 10% ethylene glycol and 20% (w/v) polyethylene glycol 3,350 at 20°C. Diffraction quality crystals for SH13 H7 HA ectodomain trimer with HA2 N95L mutation were obtained by mixing 0.1 µL HA protein at 12.8 mg/ml in 20 mM Tris pH 8.0, 150 mM NaCl and 0.02% (v/v) NaN_3_ with 0.1 µL of the well solution in 0.08 M sodium cacodylate, pH 6.5, 0.16 M magnesium acetate, 20% (v/v) glycerol and 16% (w/v), and 16% (w/v) polyethylene glycol 8,000 at 20°C. The crystals were flashed-cooled at 100 K without additional cryo-protectant. Diffraction data were collected at synchrotron beamlines (Table S1). Data for all crystals were integrated and scaled with HKL2000 (*160*). Data collection statistics are summarized in **Table S1**. The crystal structures of HA2 N95L mutants of HK68 H3 HA and SH13 H7 HA were determined by molecular replacement (MR) using the program Phaser (*161*) using the wild-type HK68 H3 HA structure (PDB 4FNK) and wild-type SH13 H7 HA (PDB 4N5J) as input MR models, respectively. Initial rigid body refinement was performed in REFMAC5 (*162*), and subsequently simulated annealing and restrained refinement (including TLS refinement) were carried out in Phenix (*163*). Between rounds of refinements, model building was examined and modified with the program Coot (*164*). Final statistics for these structures are summarized in **Table S1**. The JCSG validation suite (qc-check.usc.edu/QC/qc_check.pl) and MolProbity (*165*) were used to validate the structural quality. All figures were generated with PyMol (www.pymol.org).

### Enzyme-linked immunosorbent assay (ELISA)

Costar^TM^ 96-well, high-binding, flat-bottom, half-area plates (Corning) were coated with 50 µl of PBS containing 0.1 μg of the appropriate HA trimer or SApNP antigens. Plates were incubated overnight at 4 °C and washed five times with PBST (PBS + 0.05% [v/v] Tween 20). Wells were then blocked for 1 h at room temperature with 150 µl of blocking buffer consisting of 4% (w/v) blotting-grade blocker (Bio-Rad) in PBS, followed by five additional PBST washes. To assess antibody binding, antibodies were diluted in blocking buffer to a starting concentration of 10 µg/ml and subjected to a 10-fold serial dilution. A 50 μl volume of each antibody dilution was added to the appropriate wells. For serum binding analysis, mouse serum was diluted 40-fold in blocking buffer and also subjected to a 10-fold dilution series, with 50 μl of each dilution added per well. All plates were incubated for 1 h at room temperature and then washed five times with PBST. For detection, a 1:5000 dilution of goat anti-human IgG secondary antibody was used for antibody binding, and a 1:3000 dilution of horseradish peroxidase (HRP)-conjugated goat anti-mouse IgG secondary antibody (Jackson ImmunoResearch Laboratories) was used for serum binding. In both cases, 50 μl of the appropriate secondary antibody diluted in PBST was added to each well. After 1 h incubation at room temperature, plates were washed six times with PBST. Finally, wells were developed with 50 µl of 3,3’,5,5’-tetramethylbenzidine (TMB; Life Sciences) for 3–5 minutes, and the reaction was stopped by adding 50 µl of 2 N sulfuric acid. Absorbance was immediately read at 450 nm using a BioTek Synergy plate reader. EC_50_ values were calculated by fitting dose-response curves derived from ELISA binding data using GraphPad Prism version 10.3.1. In antibody binding analyses, if the optical density at 450 nm (OD_450_) did not reach 0.5 at the starting antibody concentration (10 µg/ml), binding was considered undetectable, and an EC_50_ value of 100 µg/ml was assigned to facilitate plotting and comparison.

### Negative-stain electron microscopy (nsEM)

The nsEM analysis was performed by the Core Microscopy Facility at The Scripps Research Institute. Briefly, HA trimer and SApNP samples were prepared at concentrations of 0.008 and 0.01 mg/ml, respectively. Carbon-coated copper grids (400 mesh) were glow-discharged prior to sample application. A volume of 8 µl of each sample was adsorbed onto the grids for 2 min, followed by removal of excess liquid. Grids were negatively stained with 2% (w/v) uranyl formate for 2 min. Excess stain was wicked away, and the grids were allowed to dry. For HA trimers, samples were imaged using a Talos L120C transmission electron microscope (TEM; Thermo Fisher) operated at 120 kV. Approximately 30 images were collected at 73,000× magnification using a CETA 16M CMOS camera, with a resolution of 1.93 Å/pixel and a defocus range of 0.5-2 µm. For SApNPs, fewer than 10 images were acquired at 52,000× magnification, with a pixel size of 2.05 Å and a defocus set to −1.50 µm. Image processing of HA trimers was conducted using the high-performance computing core facility at The Scripps Research Institute. Raw images were converted to MRC format using EMAN2 (*166*), and further processed in cryoSPARC v4.3.0 (*167*). Contrast transfer function (CTF) estimation was performed using patch-based CTF correction. Particles were picked using blob and template-based methods and extracted with a box size of 150 pixels. Minimum and maximum particle sizes were set to 70 and 260 Å, respectively, during blob picking. Ab initio 3D reconstruction was followed by heterogeneous and homogeneous refinement. All EM and model-fitting images were rendered using UCSF Chimera (*168*) and Chimera X (*169*).

### Dynamic Light Scattering (DLS)

Hydrodynamic radius and particle size distributions of HA-presenting SApNPs were measured using a Zetasizer Ultra instrument (Malvern). Briefly, SApNP samples were diluted to 0.2 mg/ml in 1× PBS buffer, and 30 μl of each sample was loaded into a quartz batch cuvette (Malvern, catalog no. ZEN2112). Measurements were conducted at 25 °C in backscattering mode. Data analysis was performed using the Zetasizer Ultra software, and particle size distributions were plotted using GraphPad Prism version 10.3.1.

### Propagation of influenza viruses

The following reagents were obtained for hemagglutination inhibition (HAI), virus neutralization, and viral challenge experiments from BEI Resources, National Institute of Allergy and Infectious Diseases (NIAID), National Institutes of Health (NIH) (NR catalog codes) or International Reagent Resource, World Health Organization Collaborating Center for Surveillance, Epidemiology and Control of Influenza, Centers for Disease Control and Prevention (CDC) (FR catalog codes): A/California/04/2009 (CA09, H1N1; NR-136583), A/Puerto Rico/8/1934 (PR8, H1N1; NR-348), A/New Caledonia/20/1999 (NC99, H1N1; FR-395), A/Michigan/45/2015 (MI15, H1N1; FR-1483), A/Wisconsin/67/2022 (WI22, H1N1; FR-1857), A/Hong Kong/1/1968 (HK68, H3N2; NR-28620), A/Brisbane/10/2007 (BR07, H3N2; NR-12283), A/Hong Kong/4801/2014 (HK14; H3N2; FR-1453), B/Florida/4/2006 (FL06, Yamagata lineage; NR-41795), B/Brisbane/60/2008 (BB08, Victoria lineage; NR-42005), A/Puerto Rico/8/1934 Mouse-Adapted (PR8-ma, H1N1; NR-28652), and A/Hong Kong/1/1968 Mouse-Adapted (HK68-ma, H3N2; NR-28621). Mouse-adapted B/Florida/4/2006 (FL06-ma, Yamagata lineage) was kindly provided by Dr. Eric Weaver. Viruses were propagated as described previously with modifications (*100*). Briefly, 4.4 × 10^6^ Madin-Darby canine kidney (MDCK) cells (CCL-34; ATCC) were plated overnight in 100 mm cell culture dishes. The next day, cells were washed with PBS and infected with virus at a multiplicity of infection (MOI) between 0.001 to 1 in serum-free media for 1 h. Next, cells were washed, and 12 ml of serum-free DMEM containing 0.2% w/v bovine serum albumin (BSA; VWR International) and 1 μg/mL N-tosyl-L-phenylalanyl chloromethyl ketone (TPCK)-treated trypsin (Sigma-Aldrich) was added to the dishes. Cells were incubated for 72–96 h, after which the supernatant was collected, centrifuged at 4000 rpm for 10 min, aliquoted, and frozen at −80 °C until use.

### Immunostaining and quantification of influenza viruses

The 50 Tissue Culture Infectious Dose (TCID_50_) of viruses was determined through a viral titration assay, with immunostaining. First, propagated viruses were subjected to a 10-fold dilution series in serum-free media containing 0.2% w/v BSA and 1 μg/ml TPCK-treated trypsin. Next, 96-well plates seeded the previous day with MDCK cells (2.5 × 10^4^ cells/well) were washed, and incubated with 100 μl/well of the virus dilutions for 24 h at 37 °C. The next day, the supernatants were removed, cells were fixed with 3.7% w/v formaldehyde (Sigma-Aldrich) for 1 h, and then permeabilized with ice-cold methanol (Thermo Fisher) for 20 min. Plates were then washed with deionized water and incubated with 50 μl/well of influenza HA stem antibody MEDI8852 (0.5 μg/ml) for 1 h. Plates were then washed four times and incubated with 50 μl/well of a 1:2000 dilution of HRP-goat anti-human secondary IgG (Jackson ImmunoResearch Laboratories, Inc.). Next, plates were washed, and virus-infected cells were visualized through incubation with TrueBlue peroxidase substrate (25 μl/well) for 20 – 30 min. Reactions were stopped by washing the plates two times with deionized water. Plates were then imaged using a Bioreader 7000 (BIOSYS Scientific). TCID_50_ values were calculated using the Reed–Muench and Spearman–Karber methods (*170*).

### Mouse immunization, sample collection, and challenge studies

We utilized a mouse immunization protocol similar to our previous studies (*78, 82, 86, 99, 100*). All animal experiments were conducted following the guidelines of the Association for the Assessment and Accreditation of Laboratory Animal Care (AAALAC). Animal protocols were approved by the Institutional Animal Care and Use Committee (IACUC) at The Scripps Research Institute (TSRI). Six-to-eight-week-old female BALB/c mice were purchased from The Jackson Laboratory and housed in ventilated cages in environmentally controlled rooms in the immunology building of TSRI. For mechanistic studies of HA immunogen trafficking, retention, presentation, and vaccine-induced GC reactions in lymph nodes, adjuvanted vaccine antigens were administered intradermally into mouse footpads using a 29-gauge insulin needle under 3% isoflurane anesthesia with oxygen. Each mouse received 80 μl of antigen/AV adjuvant (InvivoGen) mix, comprising 40 μg of HA antigen (10 μg per foot) and 40 μl of AV adjuvant. For immunogenicity and challenge studies, mice were immunized intradermally via footpad injections at weeks 0, 3, and 6. Each mouse received 80 μl of antigen/AV adjuvant mix, containing 10 μg of HA antigen (2.5 μg per foot) and 40 μl of AV. Blood samples were collected from the maxillary/facial vein at weeks 2, 5, and 8. Serum was isolated from blood by centrifugation at 14,000 rpm for 10 min, heat-inactivated at 56 °C for 30 min, and centrifuged again at 8,000 rpm for 10 min. Heat-inactivated serum was stored at −4 °C until further use in binding and neutralization assays. At week 9, three weeks after the final immunization, mice were lightly anesthetized with isoflurane and challenged with live virus (25 μl/nostril). CA09 H1 HA-immunized mice were challenged with LD_50_ × 10 of mouse-adapted PR8 (H1N1, non-vaccine-matched). HK68 H3 HA-immunized mice were challenged with LD_50_ × 10 of mouse-adapted HK68 (H3N2, vaccine-matched). BB08 (Victoria lineage) HA-immunized mice were challenged with LD_50_ × 5 of mouse-adapted FL06 (Yamagata lineage; non-vaccine-matched). LD_50_ doses for challenge strains were determined in BALB/c mice using the Reed–Muench and Spearman–Karber methods (*170*). Survival, weight loss, and morbidity were monitored for 14 days post-challenge. Mice that exhibited >25% weight loss or showed visible signs of distress were euthanized.

### Histology, immunostaining, and imaging for lymph node tissues

Vaccine distribution, trafficking, retention, cellular interaction, and GC reactions in sentinel lymph nodes were studied in mice. CA09 HA-N95L trimer and its FR and E2p SApNPs, formulated with AV adjuvant, were administered via intradermal injection into four footpads. Protocols for mouse injection, tissue collection, and data analysis followed those described in previous studies (*78, 82, 84, 100*). Each mouse received 80 μl of antigen/adjuvant mixture containing 40 μg of antigen (10 μg per footpad). Sentinel lymph nodes were collected at time points ranging from 12 h to 8 weeks after a single-dose injection for immunohistological analysis. Lymph nodes were embedded in frozen section compound (VWR International, catalog no. 95057-838) in plastic cryomolds (Tissue-Tek at VWR, catalog no. 4565), rapidly frozen in liquid nitrogen, and stored at −80 °C prior to transfer to the Core Facility of TSRI for tissue processing, immunostaining, and imaging. Lymph node tissues were sectioned using a cryostat and placed on charged slides. Tissues were sectioned using a cryostat and mounted on charged slides. Sections were fixed in 10% neutral-buffered formalin and permeabilized in PBS containing 0.5% Triton X-100. Slides were blocked with Protein Block (Agilent) to minimize nonspecific antibody binding prior to immunostaining. Primary antibodies were applied to lymph node sections and incubated overnight at 4 °C. After washing with TBS containing 0.1% Tween-20 (TBST), biotinylated or fluorophore-conjugated secondary antibodies were applied, and samples were incubated for 1 h at 25 °C. Sections were stained with human antibodies FluA-20 (*119*), MEDI8852 (*118*), and CR9114 (*126*) (1:50), followed by biotinylated goat anti-human secondary antibody (Abcam, catalog no. ab7152, 1:300). Detection was performed using streptavidin-HRP reagent (Vectastain Elite ABC-HRP Kit, Vector, catalog no. PK-6100) and diaminobenzidine (ImmPACT DAB, Vector, catalog no. SK-4105).

To study interactions between HA immunogens and cellular components in lymph nodes, various cell types were identified by immunostaining. FDCs were labeled using an anti-CD21 primary antibody (Abcam, cat. no. ab75985, 1:1800), followed by an anti-rabbit secondary antibody conjugated to Alexa Fluor 555 (Thermo Fisher, cat. no. A21428, 1:200). Subcapsular sinus macrophages were labeled with an anti-sialoadhesin (CD169) antibody (Abcam, cat. no. ab53443, 1:600), followed by an anti-rat secondary antibody conjugated to Alexa Fluor 488 (Abcam, cat. no. ab150165, 1:200). B cells were detected using an anti-B220 antibody (eBioscience, cat. no. 14-0452-82, 1:100), followed by an anti-rat secondary antibody conjugated to Alexa Fluor 647 (Thermo Fisher, cat. no. A21247, 1:200). HA trimer- and SApNP-induced GC reactions were also assessed. GC B cells were stained using a FITC-conjugated rat anti-GL7 antibody (BioLegend, cat. no. 144604, 1:250). T_fh_ cells were identified using an anti-CD4 antibody (BioLegend, cat. no. 100402, 1:100), followed by an anti-rat secondary antibody conjugated to Alexa Fluor 488 (Abcam, cat. no. ab150165, 1:1000). GC-forming cells were stained with an anti-Bcl6 antibody (Abcam, cat. no. ab220092, 1:300), followed by an anti-rabbit secondary antibody conjugated to Alexa Fluor 555 (Thermo Fisher, cat. no. A21428, 1:1000). Cell nuclei were counterstained with 4′,6-diamidino-2-phenylindole (DAPI; Sigma-Aldrich, cat. no. D9542, 100 ng/ml). Stained lymph node sections were scanned using a Leica AT2 whole-slide scanner. Vaccine immunogen trafficking and GC formation in lymph nodes were quantified from bright-field and fluorescence images using ImageJ software (*171*).

### TEM analysis of HA-presenting SApNPs in lymph node tissues

To assess interactions between HA immunogens and cellular components in lymph nodes at both intracellular and intercellular levels, TEM analysis was conducted by the Core Microscopy Facility at TSRI. Lymph node isolation, processing, and analysis followed procedures established in our previous studies (*78, 82, 84*). To visualize AV-formulated HA SApNPs in association with FDCs and phagocytic cells, mice were administered 200 μl of an antigen/adjuvant mixture containing 100 μg of HA immunogen and 100 μl of AP, delivered via intradermal injection into the two hind footpads (50 μg per footpad). Fresh sentinel lymph nodes were collected at 12 and 48 h after a single-dose injection. Tissues were bisected and fixed overnight at 4 °C in oxygenated fixative comprising 2.5% glutaraldehyde and 4% paraformaldehyde in 0.1 M sodium cacodylate buffer (pH 7.4). Samples were then washed with 0.1 M sodium cacodylate buffer and post-fixed in 1% osmium tetroxide and 1.5% potassium ferrocyanide in the same buffer at 4 °C for 1 h. After rinsing with Corning Cell Culture Grade Water, tissues were stained with 0.5% uranyl acetate overnight at 4 °C. Samples were subsequently rinsed with double-distilled H_2_O, dehydrated through a graded ethanol series followed by acetone, and infiltrated with LX-112 epoxy resin (Ladd), which was polymerized at 60 °C. Ultrathin sections (70 nm) were cut and mounted on copper grids for TEM imaging. Sections were imaged at 80 kV using a Talos L120C TEM (Thermo Fisher), and images were acquired with a CETA 16M CMOS camera for further analysis.

### Lymph node disaggregation, cell staining, and flow cytometry

To access vaccine-induced humoral immunity, the frequencies and numbers of GC B cells (B220^+^GL7^+^CD95^+^) and T_fh_ cells (CD3^+^CD4^+^CXCR5^+^PD-1^+^) were quantified by flow cytometry. Lymph node tissue collection, processing, and analysis followed protocols established in previous studies (*78, 82, 84, 100*). Each mouse received 80 μl of an antigen/adjuvant mixture—containing 40 μg of HA antigen (10 μg per footpad) formulated with AV—via intradermal injection into four footpads. Sentinel lymph nodes were isolated at 2, 5, and 8 weeks after a single-dose injection, and mechanically disaggregated. Tissues were transferred to enzyme digestion solution in 1.5 ml Eppendorf tubes containing 958 μl of Hanks’ Balanced Salt Solution (HBSS; Thermo Fisher Scientific, catalog no. 14185052), 40 μl of 10 mg/ml collagenase IV (Sigma-Aldrich, catalog no. C5138), and 2 μl of 10 mg/ml DNase I (Roche, catalog no. 10104159001). Samples were incubated at 37 °C for 30 min on a rotator, followed by filtration through a 70 μm cell strainer to obtain single-cell suspensions. Cells were centrifuged at 400 × g for 10 min and resuspended in HBSS blocking buffer containing 0.5% (w/v) bovine serum albumin and 2 mM EDTA. To block Fc receptor–mediated nonspecific binding, cells were incubated on ice for 30 min with anti-CD16/32 antibody (BioLegend, catalog no. 101302, 1:50). Cells were then transferred to 96-well V-bottom microplates and stained with a pre-mixed antibody cocktail on ice for 30 min. The staining cocktail included: Zombie NIR live/dead stain (BioLegend, catalog no. 423106, 1:100), Brilliant Violet 510 anti-mouse/human CD45R/B220 antibody (BioLegend, catalog no. 103247, 1:300), FITC anti-mouse CD3 antibody (BioLegend, catalog no. 100204, 1:300), Alexa Fluor 700 anti-mouse CD4 antibody (BioLegend, catalog no. 100536, 1:300), PE anti-mouse/human GL7 antibody (BioLegend, catalog no. 144608, 1:500), Brilliant Violet 605 anti-mouse CD95 (Fas) antibody (BioLegend, catalog no. 152612, 1:500), Brilliant Violet 421 anti-mouse CD185 (CXCR5) antibody (BioLegend, catalog no. 145511, 1:500), and PE/Cyanine7 anti-mouse CD279 (PD-1) antibody (BioLegend, catalog no. 135216, 1:500). Following staining, cells were centrifuged at 400 × *g* for 10 min, washed with HBSS blocking solution, and fixed with 1.6% paraformaldehyde (Thermo Fisher Scientific, catalog no. 28906) in HBSS on ice for 30 min. Finally, cells were spun down again at 400 × *g* for 10 min and stored in HBSS blocking solution at 4 °C until acquisition. Flow cytometry was performed on a five-laser ZE5 cell analyzer (Yeti model; Bio-Rad) equipped with Everest software at the TSRI Core Facility. Data analysis was conducted using FlowJo version 10 software, and the results were plotted using GraphPad Prism version 10.3.1.

### Hemagglutination inhibition (HAI) assays

Hemagglutination inhibition (HAI) assays, which determine the serum quantity required to inhibit the agglutinating activity of influenza viruses, were performed according to WHO guidelines (*172, 173*). To determine the hemagglutinating units (HA units) for each virus strain, 50 μl of 2-fold serially diluted virus was incubated with 50 μl of 0.75% (w/v) turkey red blood cells (RBCs; 5% packed, Innovative Research) in V-bottom 96-well plates for 30 min at room temperature (RT). A single HA unit was defined as the last virus dilution that caused complete agglutination of RBCs. To assess HAI activity, mouse sera were pretreated with receptor-destroying enzyme (RDE; Fisher Scientific, catalog no. NC9553641) by diluting one part serum with three parts RDE. The sera were then incubated at 37 °C for 18–20 h and inactivated at 56 °C for 30 min, after which six parts PBS were added to achieve a final 1:10 dilution. The serum was then further diluted 4-fold in PBS and subjected to a 2-fold dilution series (25 μl/well) in V-bottom plates. The selected virus was adjusted to 8 HA units in 25 μl and added to each well. Following a 30-min incubation at RT, 50 μl of 0.75% RBCs were added to each well and incubated for an additional 30 min. The HAI titer was defined as the reciprocal of the highest serum dilution that fully inhibited hemagglutination. Samples with no detectable HAI activity were assigned a titer of 20 (i.e., half the starting dilution of 1:40). For HAI assays (and neutralization assays mentioned below), H1 HA-immune sera were tested against a panel of vaccine-matched (CA09) and non-matched H1N1 strains, whereas H3 HA-immune sera were tested against vaccine-matched (HK68) and non-matched H3N2 strains.

### Live virus microneutralization assays

Live virus neutralization assays were performed to assess the neutralizing activity of vaccine-induced mouse sera. Mouse sera were pretreated with RDE by diluting one part serum with three parts RDE. The sera were then incubated at 37 °C for 18–20 h and inactivated at 56 °C for 30 min, after which six parts PBS were added to achieve a final 1:10 dilution. Next, sera were diluted another 10-fold to produce 100-fold dilutions and then subjected to a 2-fold dilution series incubated with TCID_50_ × 100 of virus diluent (final concentration of 0.2% w/v BSA and 1 μg/ml TPCK-treated trypsin in DMEM) for 1 h at 37 °C. Next, 96-well plates seeded the previous day with MDCK cells (2.5 × 10^4^ cells/well) were washed, and incubated with 100 μl/well of the serum-virus dilutions for 24 h at 37 °C. The next day, supernatants were removed, and cells were fixed and subjected to the immunostaining protocol described above for virus quantification. Neutralization titers were quantified as the reciprocal of the highest serum dilutions that contained no observable virus. Samples with no detectable neutralization were assigned a titer of 50 (1/2 the starting dilution of 1:100).

### Pseudovirus production and neutralization assays

Influenza HA pseudotyped particles (pps) were generated by the co-transfection of HEK293T cells with envelope-deficient HIV-1 pNL4–3.lucR−E− plasmid (NIH AIDS reagent program: https://www.aidsreagent.org/) and an expression plasmid encoding the HA gene of avian H5 A/Vietnam/1203/2004 (VN04) (GenBank: AY818135) or H7 A/Guangdong/Th005/2017 (GD17) (GenBank: KY643843) strain at a 4:1 ratio using a Lipofectamine 3000 transfection protocol (Thermo Fisher). After 72 h, pseudoviruses were collected from the supernatant by centrifugation at 3724 × g for 10 min, aliquoted, and stored at −80°C until use. For pseudovirus neutralization assays, mouse sera starting at a dilution of 100× were serially diluted 3-fold and incubated with pseudoviruses at 37 °C for 1 h in white solid-bottom 96-half-well plates (Corning). Next, 8.0 × 10^3^ HEK293T cells were added to each well and plates were then incubated at 37°C for 60–72 h. After incubation, the supernatant was removed, and the cells were lysed. Luciferase reporter gene expression was quantified through the addition of Bright-Glo Luciferase substrate (Promega) according to the manufacturer’s instructions. Luciferase activity in relative light units (RLU) was measured on a BioTek microplate reader with Gen 5 software. Values from experimental wells were compared against virus-containing wells, with background luminescence from a series of uninfected wells subtracted from both.

### Statistical analysis

Data were collected from 8 mice per group in immunization and challenge studies and 3-5 mice per group in the mechanistic study. All ELISA, hemagglutination inhibition and neutralization assays were performed in duplicate. Different vaccine constructs, adjuvant-formulated HA immunogens, were compared using one-way analysis of variation (ANOVA), followed by Tukey’s multiple comparison *post hoc* test. Statistical significance is indicated as the following in the figures: ns (not significant), \**p* < 0.05, \*\**p* < 0.01, \*\*\**p* < 0.001, and \*\*\*\**p* < 0.0001. The graphs were generated, and statistical analyses were performed using GraphPad Prism 10.3.1 software.

## ACKNOWLEDGEMENTS

We acknowledge K. Vanderpool, T. Fassel, and S. Henderson of the Microscopy Core Facility at TSRI for their expert assistance with negative-stain EM analysis. We thank Henry Tien for automated robotic crystal screening. We thank M. Livneh, K. Spencer, and S. Henderson of the Histology and Microscopy Core Facility at TSRI for their expertise and technical support in immunohistology. We acknowledge A. Saluk, B. Seegers, and B. Monteverde of the Flow Cytometry Core Facility at TSRI for their technical support. The mouse-adapted influenza B virus B/Florida/4/2006 (Yamagata lineage) was kindly provided by Dr. Eric Weaver and Matthew Pekarek from the Nebraska Center for Virology at University of Nebraska-Lincoln as a gift.

## Author contributions

Project design by Y.-N.Z, K.B.G., and J.Z.; HA trimer and SApNP design by L.H., and J.Z; antibody expression and purification by G.W., C.D., and L.H.; HA trimer and SApNP expression by G.W., C.D., Y.-Z.L., and L.H.; SDS and BN PAGE by G.W., C.D., Y.-Z.L., and L.H.; DSC and DSL by Y.-Z.L., and L.H.; ELISA by Y.-Z.L. and Y.-N.Z.; X-ray crystallographic analysis by X.Z., and I.A.W.; negative-stain EM by Y.-Z.L. and J.Z.; virus preparation by Y.-N.Z., and K.B.G.; mouse immunization and viral challenge by Y.-N.Z., and K.B.G.; HAI and neutralization assays by K.B.G.; lymph node analysis by Y.-N.Z.; manuscript written by Y.-N.Z., K.B.G., Y.-Z.L., X.Z., M.N., I.A.W., and J.Z. All authors were asked to comment on the manuscript.

## Data and material availability

.All data needed to evaluate the conclusions of this research are available in the main text and Supplementary Information. The atomic coordinates and structure factors have been deposited in the Protein Data Bank (PDB) under accession codes AAA and BBB for the HK68 and SH13 HA-N95L constructs, respectively. The authors declare that the data supporting the findings of this study are available within the article and its Supplementary Information files. Source data are provided with this paper.

