## Supplemental materials for "Single-component self-assembling protein nanoparticles displaying stabilized prefusion-closed hemagglutinin trimers for influenza vaccine development"

**Fig. S1**

a

**>H1N1-CA09-HA-[N95L H26W/K51I/E103I]-foldon-His$_6$**

MKAILVVLLYTFATANADTLCIGYHANNSTDTVDTVLEKNVTVTHSVNLLEDKHNGKLCKLRGVAPLHLGKCNIAGWILGNPECESLSTA
SSWSYIVETPSSDNGTCYPGDFIDYEELREQLSSVSSFERFEIFPKTSSWPNHDSNKGVTAACPHAGAKSFYKNLIWLVKKGNSYPKLSK
SYINDKGKEVLVLWGIHHPSTSADQQSLYQNADTYVFVGSSRYSKKFKPEIAIRPKVRDQEGRMNYYWTLVEPGDKITFEATGNLVVPRY
AFAMERNAGSGIIISDTPVHDCNTTCQTPKGAINTSLPFQNIHPITIGKCPKYVKSTKLRLATGLRNIPSIQSRGLFGAIAGFIEGGWTG
**H26W** **K51I** **N95L** **E103I**
MVDGWYGYHHQNEQGSGYAADLKSTQNAIDEITNKVNSVIEKMNTQFTAVGKEFNHLEKRIENLNKKVDDGFLDIWTYNAELLVLLENER
TLDYHDSNVKNLYEKVRSQLKNNAKEIGNGCFEFYHKCDNTCMESVKNGTYDYPKYSEEAKLNREEIDG**AS**GYIPEAPRDGQAYVRKDGE
WVLLSTFLGSHHHHHH

**>H5N1-VN04-HA-[N95L H26W/K51I/E103I]-foldon-His$_6$**

MEKIVLLFAIVSLVKSDQICIGYHANNSTEQVDTIMEKNVTVTHAQDILEKKHNGKLCDLDGVKPLILRDCSVAGWLLGNPMCDEFINVP
EWSYIVEKANPVNDLCYPGDFNDYEELKHLLSRINHFEKIQIIPKSSWSSHEASLGVSSACPYQGKSSFFRNVVWLIKKNSTYPTIKRSY
NNTNQEDLLVLWGIHHPNDAAEQTKLYQNPTTYISVGTSTLNQRLVPRIATRSKVNGQSGRMEFFWTILKPNDAINFESNGNFIAPEYAY
KIVKKGDSTIMKSELEYGNCNTKCQTPMGAINSSMPFHNIHPLTIGECPKYVKSNRLVLATGLRNSPQRERRRKKRGLFGAIAGFIEGGW
**H26W** **K51I** **N95L** **E103I**
QGMVDGWYGYHHSNEQGSGYAADKESTQKAIDGVTNKVNSIIDKMNTQFEAVGREFNNLERRIENLNKKMEDGFLDVWTYLAELLVLMEN
ERTLDFHDSNVKNLYDKVRLQLRDNAKELGNGCFEFYHKCDNECMESVRNGTYDYPQYSEEARLKREEIS**AS**GYIPEAPRDGQAYVRKDG
EWVLLSTFLGSHHHHHH

: Leader sequence
: N95L mutation
: H26W, K51I, and E103I mutation
: Foldon trimerization motif
GS: Linker; **AS**: Enzymatic site; HHHHHH: His$_6$-tag

b

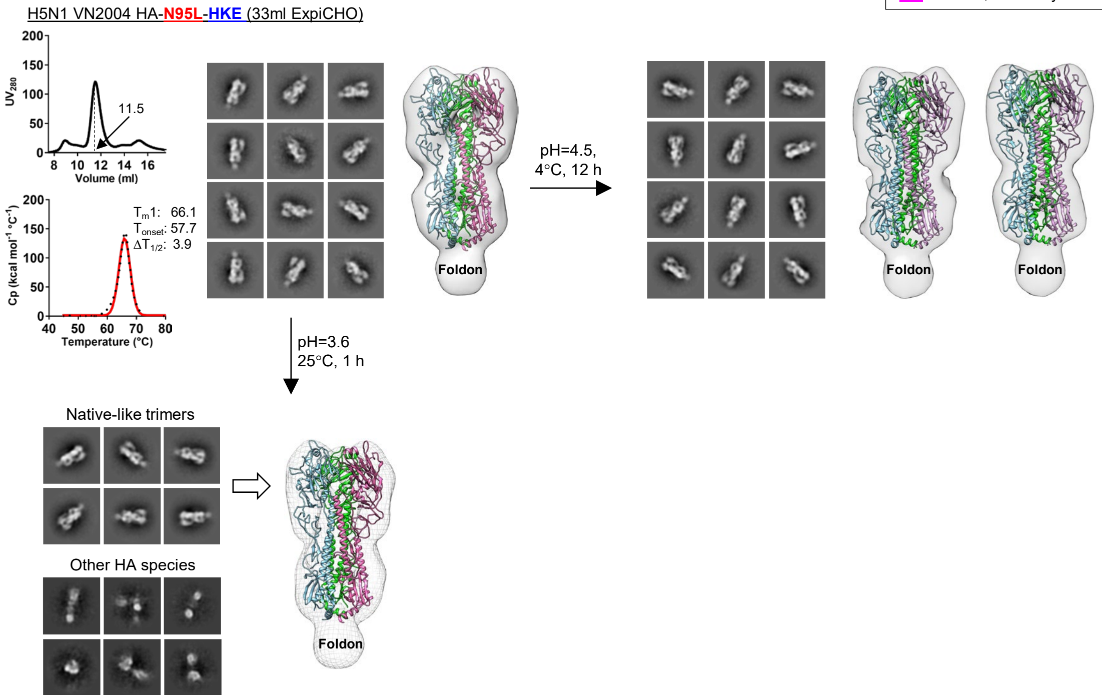

c

**>H1N1-CA09-HA-[R106M and R106M/E103L w/o N95L]-foldon-His$_6$**

MKAILVVLLYTFATANADTLCIGYHANNSTDTVDTVLEKNVTVTHSVNLLEDKHNGKLCKLRGVAPLHLGKCNIAGWILGNPECESLSTA
SSWSYIVETPSSDNGTCYPGDFIDYEELREQLSSVSSFERFEIFPKTSSWPNHDSNKGVTAACPHAGAKSFYKNLIWLVKKGNSYPKLSK
SYINDKGKEVLVLWGIHHPSTSADQQSLYQNADTYVFVGSSRYSKKFKPEIAIRPKVRDQEGRMNYYWTLVEPGDKITFEATGNLVVPRY
AFAMERNAGSGIIISDTPVHDCNTTCQTPKGAINTSLPFQNIHPITIGKCPKYVKSTKLRLATGLRNIPSIQSRGLFGAIAGFIEGGWTG
**N95L** **E103I**
MVDGWYGYHHQNEQGSGYAADLKSTQNAIDEITNKVNSVIEKMNTQFTAVGKEFNHLEKRIENLNKKVDDGFLDIWTYNAELLVLLENER
**R106M**
TLDYHDSNVKNLYEKVRSQLKNNAKEIGNGCFEFYHKCDNTCMESVKNGTYDYPKYSEEAKLNREEIDGASGYIPEAPRDGQAYVRKDGE
WVLLSTFLGSHHHHHH

: Leader sequence
xxxx: N95L mutation
: R106M mutation and/or E103L mutation
: Foldon trimerization motif
GS: Linker; **AS**: Enzymatic site; HHHHHH: His$_6$-tag

Fig. S1

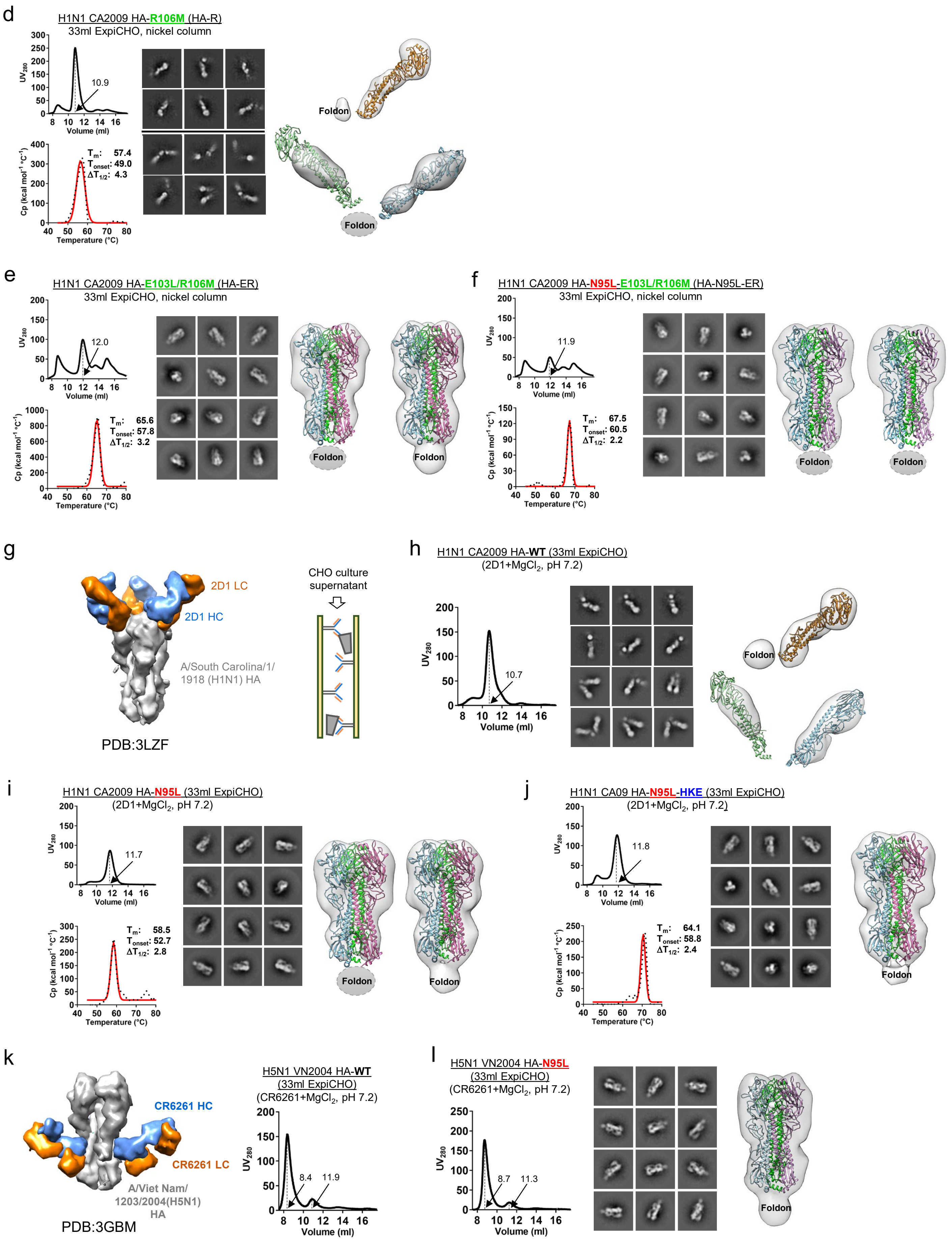

d
H1N1 CA2009 HA-R106M (HA-R)
33ml ExpiCHO, nickel column
UV280
Volume (ml)
10.9
Cp (kcal mol-1 °C-1)
Temperature (°C)
Tm: 57.4
Tonset: 49.0
ΔT1/2: 4.3
Foldon
Foldon
e
H1N1 CA2009 HA-E103L/R106M (HA-ER)
33ml ExpiCHO, nickel column
12.0
Tm: 65.6
Tonset: 57.8
ΔT1/2: 3.2
Foldon
Foldon
f
H1N1 CA2009 HA-N95L-E103L/R106M (HA-N95L-ER)
33ml ExpiCHO, nickel column
11.9
Tm: 67.5
Tonset: 60.5
ΔT1/2: 2.2
Foldon
Foldon
g
2D1 LC
2D1 HC
A/South Carolina/1/1918 (H1N1) HA
PDB:3LZF
CHO culture supernatant
h
H1N1 CA2009 HA-WT (33ml ExpiCHO)
(2D1+MgCl2, pH 7.2)
10.7
Foldon
i
H1N1 CA2009 HA-N95L (33ml ExpiCHO)
(2D1+MgCl2, pH 7.2)
11.7
Tm: 58.5
Tonset: 52.7
ΔT1/2: 2.8
Foldon
Foldon
j
H1N1 CA09 HA-N95L-HKE (33ml ExpiCHO)
(2D1+MgCl2, pH 7.2)
11.8
Tm: 64.1
Tonset: 58.8
ΔT1/2: 2.4
Foldon
k
CR6261 HC
CR6261 LC
A/Viet Nam/1203/2004(H5N1) HA
PDB:3GBM
H5N1 VN2004 HA-WT
(33ml ExpiCHO)
(CR6261+MgCl2, pH 7.2)
8.4
11.9
l
H5N1 VN2004 HA-N95L
(33ml ExpiCHO)
(CR6261+MgCl2, pH 7.2)
8.7
11.3
Foldon

m

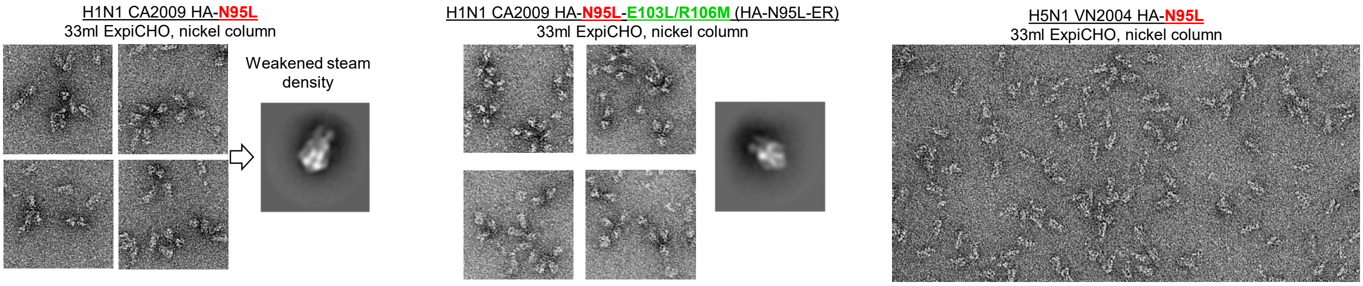

**Fig. S1. Design and in vitro characterization of group-1 IAV HA constructs.** (**a**) Amino acid sequences of H1N1 CA2009 and H5N1 VN2004 HA constructs containing N95L and the triple pHS1/2 mutations H26W/K51I/E103I (HKE). The signal peptide, HKE mutations, N95L mutation, restriction site (AS), foldon, linker (GS), and $His_6$ tag are highlighted in yellow, green, red, blue font, gray, magenta, and cyan, respectively. The foldon is a C-terminal trimerization motif, and the $His_6$ tag facilitates purification by nickel affinity chromatography. Since both elements are present in all HA constructs, they are not included in the construct names beyond this panel. (**b**) In vitro characterization of VN04 HA-N95L-HKE under various conditions. Top left: Characterization at neutral pH, including SEC profile following ExpiCHO expression and nickel purification (top left), DSC thermogram with labeled thermal parameters (bottom left), representative 2D class averages from nsEM (middle), and 3D reconstruction (right). Top right: nsEM characterization following mild acidic challenge (pH 4.5, 4 °C, 12 h), including representative 2D class averages (left) and 3D reconstruction (right). Bottom left: nsEM characterization after brief exposure to lower pH at 25 °C, including 2D class averages (left) and 3D reconstruction (right). (**c**) Amino acid sequences of CA09 HA constructs containing R106M, E103L/R106M (ER), and N95L+E103L/R106M mutations. The signal peptide, N95L mutation, E103L/R106M mutations, restriction site (AS), foldon, linker (GS), and $His_6$ tag are highlighted in yellow, red, green, blue font, gray, magenta, and cyan, respectively. (**d**)-(**f**) In vitro characterization of three CA09 HA constructs following ExpiCHO expression and nickel purification: (**d**) R106M, (**e**) E103L/R106M (ER), and (**f**) N95L+E103L/R106M. (**g**) Structural model of an HA trimer in complex with bNAb 2D1 (PDB 3LZF), which targets an epitope near the receptor-binding site (RBS) on the head domain (left), and schematic of the 2D1 IAC column. (**h**)-(**j**) In vitro characterization of three CA09 HA constructs following ExpiCHO expression and 2D1 purification: (**h**) WT HA, (**i**) HA-N95L, and (**j**) HA-N95L-HKE. Panel layout follows the format of the neutral pH panel in (**b**). (**k**) Structural model of an H5 HA trimer in complex with bNAb CR6261 (PDB 3GBM), which binds a conserved epitope on the stem region (left), and SEC profile of VN04 WT HA following ExpiCHO expression and CR6261 purification. (**l**) In vitro characterization of H5 HA-N95L following ExpiCHO expression and CR6261 purification, similar to (**h**)-(**j**) but without DSC analysis. (**m**) Micrographs from nsEM analysis to assess anisotropy. Left: CA09 HA-N95L; middle: CA09 HA-N95L-HKE; right: VN04 HA-N95L.

**a**

**>H3N2-HK68-HA-[N95L H26W/K51I/E103I]-foldon-His$_6$**

```
MKTIIALSYIFCLALGQDLPGNDNSTATLCLGHHAVPNGTLVKTITDDQIEVTNATELVQSSSTGKICNNPHRILDGIDCTLIDALLGDP
HCDVFQNETWDLFVERSKAFSNCYPYDVPDYASLRSLVASSGTLEFITEGFTWTGVTQNGGSNACKRGPGSGFFSRLNWLTKSGSTYPVL
NVTMPNNDNFDKLYIWGVHHPSTNQEQTSLYVQASGRVTVSTRRSQQTIIPNIGSRPWVRGLSSRISIYWTIVKPGDVLVINSNGNLIAP
RGYFKMRTGKSSIMRSDAPIDTCISECITPNGSIPNDKPFQNVNKITYGACPKYVKQNTLKLATGMRNVPEKQTRGLFGAIAGFIENGWE
         H26W                     K51I                                        N95L       E103I
GMIDGWYGFRHQNSEGTGQAADLKSTQAAIDQINGKLNRVIEKTNEKFHQIEKEFSEVEGRIQDLEKYVEDTKIDLWSYLAELLVALENQ
HTIDLTDSEMNKLFEKTRRQLRENAEDMGNGCFKIYHKCDNACIESIRNGTYDHDVYRDEALNNRFQIKASGYIPEAPRDGQAYVRKDGE
WVLLSTFLGSHHHHHH
```

**>H7N9-GD17-HA-[N95L H26W/K51I/E103I]-foldon-His$_6$**

```
MNTQILVFALIAIIPTNADKICLGHHAVSNGTKVNTLTERGVEVVNATETVERTNTPRICSKGKRTVDLGQCGLLGTITGPPQCDQFLEF
SADLIIERREGSDVCYPGKFVNEEALRQILRESGGIDKEPMGFTYNGIRTNGVTSACRRSGSSFYAEMKWLLSNTDNAAFPQMTKSYKNT
KESPAIIVWGIHHSVSTAEQTKLYGSGNKLVTVGSSNYQQSFVPSPGARPQVNGQSGRIDFHWLILNPNDTVTFSFNGAFIAPDRASFLR
GKSMGIQSRVQVDANCEGDCYHSGGTIISNLPFQNIDSRAVGKCPRYVKQRSLLLATGMKNVPEVPKRKRTARGLFGAIAGFIENGWEGL
       H26W                     K51I                                        N95L       E103I
IDGWYGFRHQNAQGEGTAADYKSTQSAIDQITGKLNRLIAKTNQQFKLIDNEFNEVEKQIGNVINWTRDSITEVWSYLAELLVAMENQHT
IDLADSEMDKLYERVKRQLRENAEEDGTGCFEIFHKCDDDCMASIRNNTYDHRKYREEAMQNRIQIDASGYIPEAPRDGQAYVRKDGEWV
LLSTFLGSHHHHHH
```

(yellow): Leader sequence
(red): N95L mutation
(green): H26W, K51I, and E103I mutation
(grey): Foldon trimerization motif
GS: Linker; **AS**: Enzymatic site; HHHHHH: His$_6$-tag

**b**

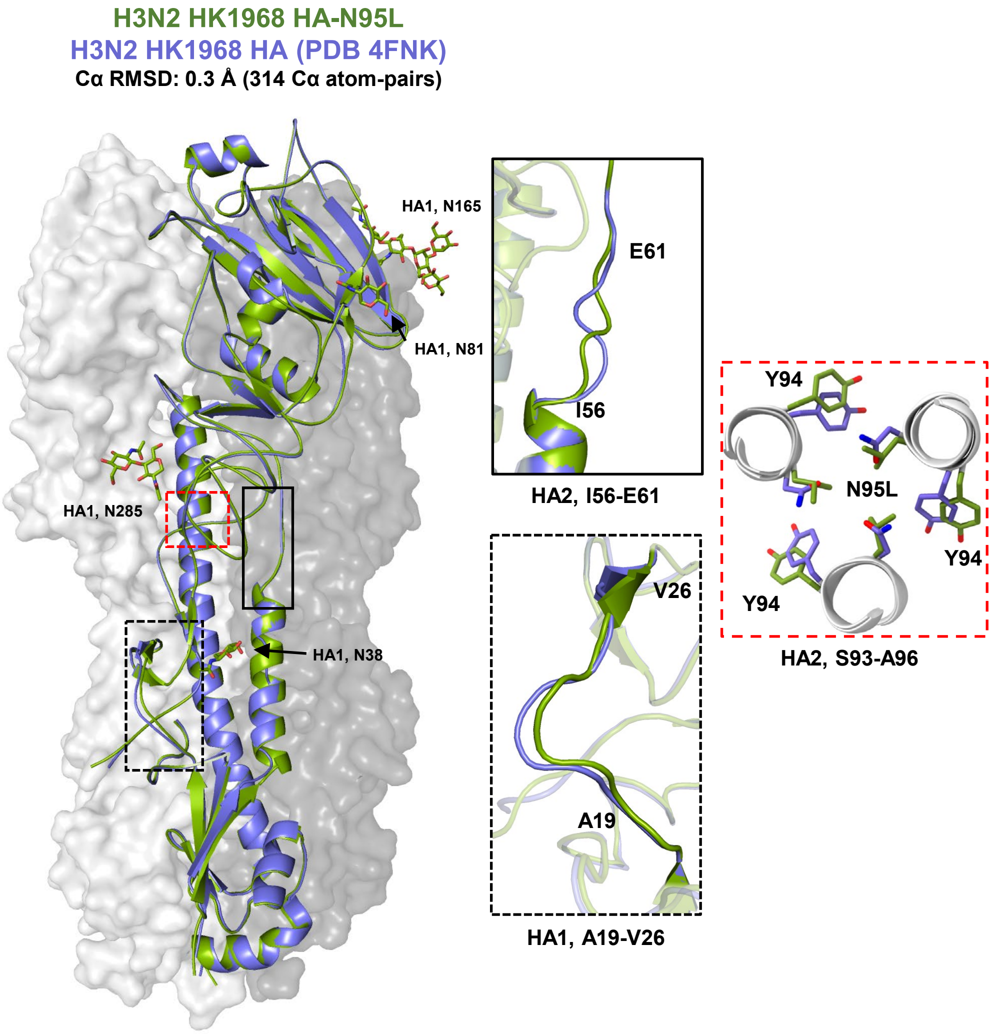

**Fig. S2**

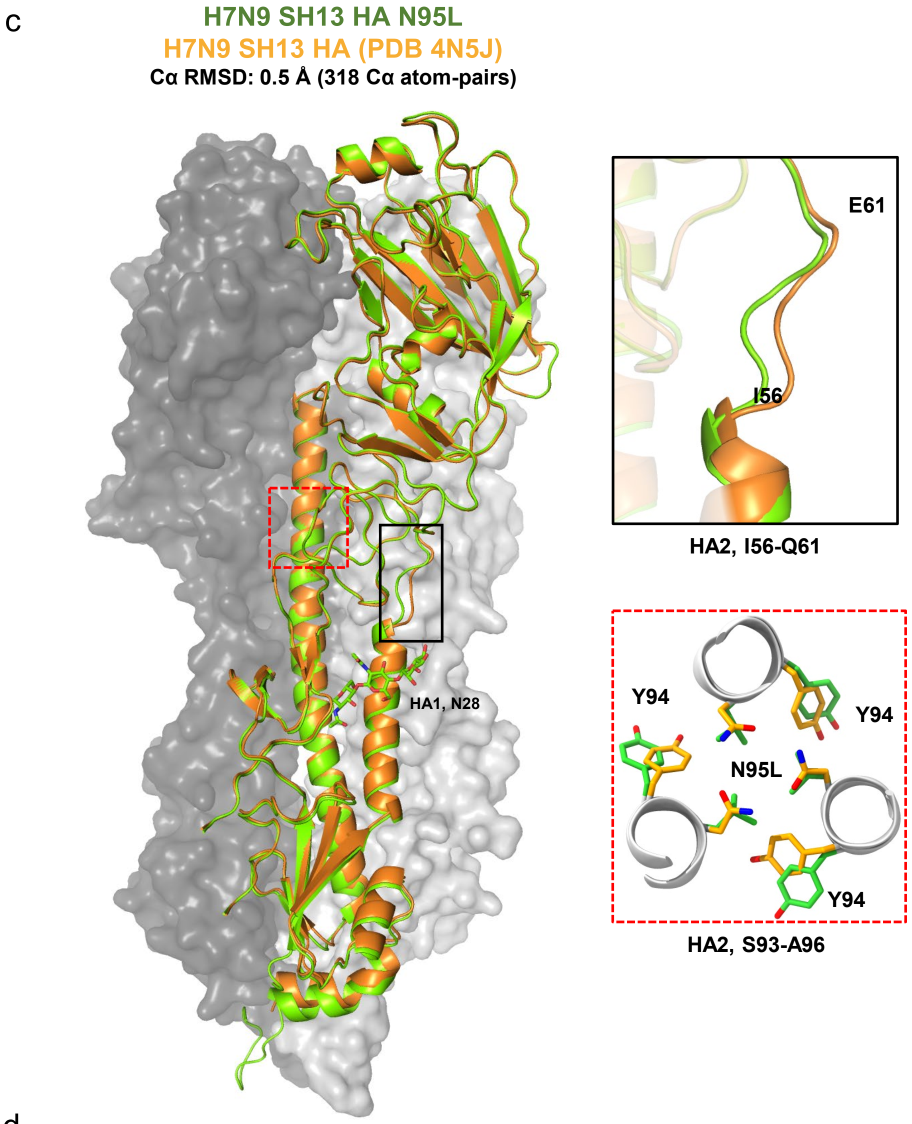
c
H7N9 SH13 HA N95L
H7N9 SH13 HA (PDB 4N5J)
Cα RMSD: 0.5 Å (318 Cα atom-pairs)
HA1, N28
E61
I56
HA2, I56-Q61
Y94
Y94
N95L
Y94
HA2, S93-A96

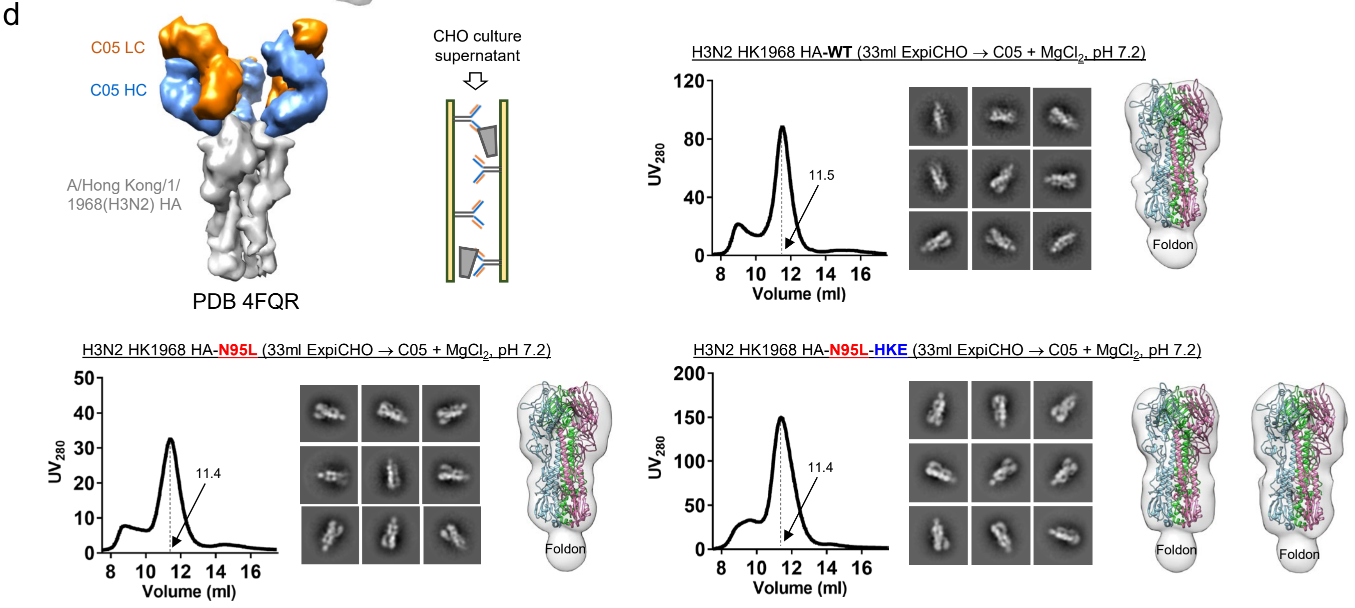
d
C05 LC
C05 HC
A/Hong Kong/1/
1968(H3N2) HA
PDB 4FQR
CHO culture
supernatant
H3N2 HK1968 HA-WT (33ml ExpiCHO → C05 + MgCl2, pH 7.2)
UV280
Volume (ml)
11.5
Foldon
H3N2 HK1968 HA-N95L (33ml ExpiCHO → C05 + MgCl2, pH 7.2)
11.4
Foldon
H3N2 HK1968 HA-N95L-HKE (33ml ExpiCHO → C05 + MgCl2, pH 7.2)
11.4
Foldon
Foldon

Fig. S2

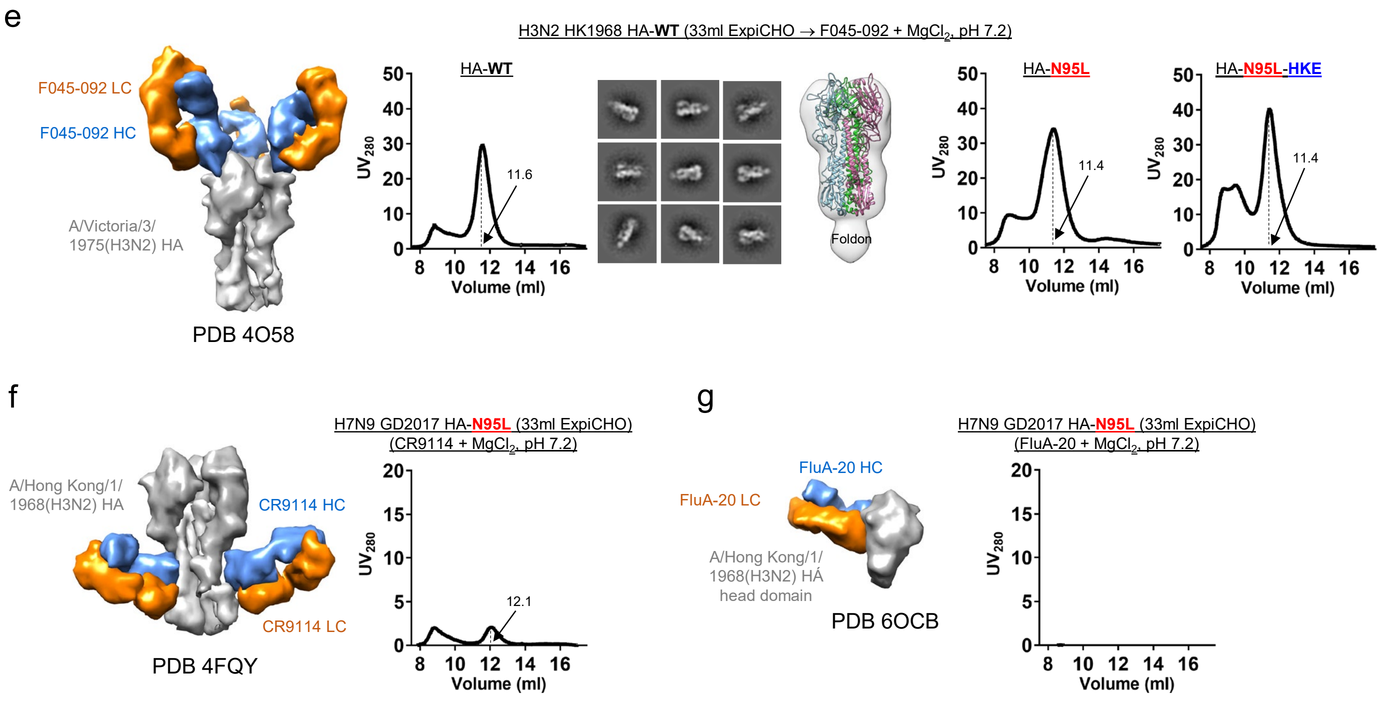

**Fig. S2. In vitro and structural characterization of group-2 IAV HA constructs.** (**a**) Amino acid sequences of HK68 and GD17 HA constructs containing the N95L mutation and the pHS1/2 mutations H26W/K51I/E103I (HKE). The signal peptide, HKE mutations, N95L mutation, restriction site (AS), foldon, linker (GS), and $His_6$ tag are highlighted in yellow, green, red, blue font, gray, magenta, and cyan, respectively. Since the foldon and $His_6$ tag are present in all constructs, they are not included in the construct names beyond this panel. (**b**) Superposition of the crystal structures of H3N2 HK1968 WT HA (cauliflower; PDB 4FNK) and HA-N95L (green), with one HA protomer shown as a ribbon model within the molecular surface of the trimer. Three HA2 regions that differ between the structures, A19-V26, I56-E61 and S93-A96, are circled and shown enlarged. (**c**) Superposition of the crystal structures of H7N9 SH2013 WT HA (cauliflower; PDB 4N5J) and HA-N95L (green), with one HA protomer shown as a ribbon model within the trimer surface. Two HA2 regions (I56-E61 and S93-A96) are circled and enlarged. (**d**) Purification of H3N2 HK1968 HA constructs by IAC using a C05 column. Top left: structural model of an H3 HA trimer in complex with bNAb C05 (PDB 4FQR), which targets the RBS via an unusually long HCDR3 loop, and schematic of the C05 IAC column. SEC profiles, 2D class averages, and 3D reconstructs are shown for H3N2 HK1968 WT HA (top right), HA-N95L (bottom left), and HA-N95L-HKE (bottom right). The trimer peak is labeled on each SEC profile. (**e**) Purification of H3N2 HK1968 HA constructs by IAC using a F045-092 column. Left: structural model of an H3 HA trimer in complex with the H3-specific bNAb F045-092, which inserts a 23-residue HCDR3 loop into the RBS, mimicking sialic acid binding. SEC profiles are shown for WT HA, HA-N95L, and HA-95L-HKE, with 2D class averages and 3D reconstructions shown for WT HA (center panels). (**f**) Purification of H7N9 GD2017 HA-N95L by IAC using a CR9114 column. Left: structural model of an H3 HA trimer in complex with bNAb CR9114, which targets the conserved stem. Right: SEC profile following IAC purification; the trimer peak is labeled. (**g**) Purification of H7N9 GD2017 HA-N95L by IAC using a FluA-20 column. Left: structural model of an H3 HA head domain in complex with bNAb FluA-20, which targets the trimer interface of the head domain. Right: SEC profile after IAC purification.

a

**>FluB-BB08-HA-[Q95L N51L/S103L]-foldon-His$_6$ (IAV H3 numbering)**

MKAIIVLLMVVTSNADRICTGITSSNSPHVVKTATQGEVNVTGVIPLTTTPTKSHFANLKGTETRGKLCPKCLNCTDLDVALGRPKCTGK
IPSARVSILHEVRPVTSGCFPIMHDRTKIRQLPNLLRGYEHIRLSTHNVINAENAPGGPYKIGTSGSCPNITNGNGFFATMAWAVPKNDK
NKTATNPLTIEVPYICTEGEDQITVWGFHSDNETQMAKLYGDSKPQKFTSSANGVTTHYVSQIGGFPNQTEDGGLPQSGRIVVDYMVQKS
GKTGTITYQRGILLPQKVWCASGRSKVIKGSLPLIGEADCLHEKYGGLNKSKPYYTGEHAKAIGNCPIWVKTPLKLANGTKYRPPAKLLK
**N51L**
ERGFFGAIAGFLEGGWEGMIAGWHGYTSHGAHGVAVAADLKSTQEAINKITKNLNSLSELEVKNLQRLSGAMDELHNEILELDEKVDDLR
**Q95L** **S103L**
ADTISSQIELAVLLSNEGIINSEDEHLLALERKLKKMLGPSAVEIGNGCFETKHKCNQTCLDRIAAGTFDAGEFSLPTFDSLNITAAS**AS**
GYIPEAPRDGQAYVRKDGEWVLLSTFLGSHHHHHH

**>FluB-FL06-HA-[Q95L N51L/S103L]-foldon-His$_6$ (IAV H3 numbering)**

MKAIIVLLMVVTSNADRICTGITSSNSPHVVKTATQGEVNVTGVIPLTTTPTKSYFANLKGTRTRGKLCPDCLNCTDLDVALGRPMCVGT
TPSAKASILHEVKPVTSGCFPIMHDRTKIRQLPNLLRGYENIRLSTQNVIDAEKAPGGPYRLGTSGSCPNATSKSGFFATMAWAVPKDNN
KNATNPLTVEVPYICTEGEDQITVWGFHSDDKTQMKNLYGDSNPQKFTSSANGVTTHYVSQIGSFPDQTEDGGLPQSGRIVVDYMMQKPG
KTGTIVYQRGVLLPQKVWCASGRSKVIKGSLPLIGEADCLHEKYGGLNKSKPYYTGEHAKAIGNCPIWVKTPLKLANGTKYRPPAKLLKE
**N51L**
RGFFGAIAGFLEGGWEGMIAGWHGYTSHGAHGVAVAADLKSTQEAINKITKNLNSLSELEVKNLQRLSGAMDELHNEILELDEKVDDLRA
**Q95L** **S103L**
DTISSQIELAVLLSNEGIINSEDEHLLALERKLKKMLGPSAVEIGNGCFETKHKCNQTCLDRIAAGTFNAGEFSLPTFDSLNITAAS**AS**G
YIPEAPRDGQAYVRKDGEWVLLSTFLGSHHHHHH

: Leader sequence
: Q95L mutation
: N51L and S103L mutation
: Foldon trimerization motif
GS: Linker; **AS**: Enzymatic site; HHHHHH: His$_6$-tag

b

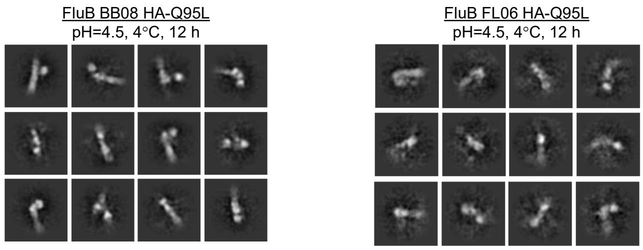

**Fig. S3. Design and in vitro characterization of IBV HA constructs.** (**a**) Amino acid sequences of FluB BB08 and FL06 HA constructs Q95L and the pHS2-inspired mutations (N51L/S103L, or NS). The signal peptide, NS mutation, Q95L mutation, restriction site (AS), foldon, linker (GS), and His$_6$ tag are highlighted in yellow, green, red, blue font, gray, magenta, and cyan, respectively. The foldon is a C-terminal trimerization motif, and the His$_6$ tag facilitates purification by nickel affinity chromatography. Since both elements are present in all HA constructs, they are not included in the construct names beyond this panel. (**b**) representative 2D class averages from nsEM analysis of BB08 (left) and FL06 (right) HA-Q95L proteins upon mild acidic challenge (pH 4.5, 4 °C, 12 h). The HA-Q95L proteins were transiently expressed in ExpiCHO cells and purified using a nickel column.

**Fig. S4**

**a**

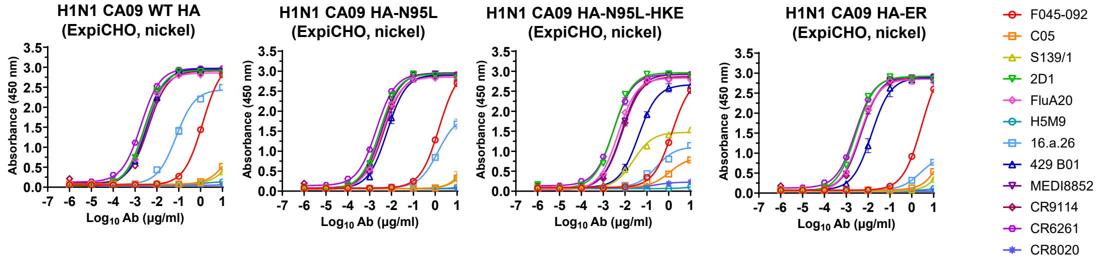

##### H1N1 CA09 WT HA (ExpiCHO, nickel)

| F045-092 | | C05 | | S139/1 | | 2D1 | | FluA20 | | H5M9 | |
|---|---|---|---|---|---|---|---|---|---|---|---|
| EC50 | STD | EC50 | STD | EC50 | STD | EC50 | STD | EC50 | STD | EC50 | STD |
| 1.25 | 0.0297 | 100 | 0 | 100 | 0 | 0.00265 | 1.90919E-05 | 0.0028 | 0.000113844 | 100 | 0 |

| 16.a.26 | | 429 B01 | | MEDI8852 | | CR9114 | | CR6261 | | CR8020 | |
|---|---|---|---|---|---|---|---|---|---|---|---|
| EC50 | STD | EC50 | STD | EC50 | STD | EC50 | STD | EC50 | STD | EC50 | STD |
| 0.0757 | 0.0103 | 0.00347 | 0.000209 | 0.00304 | 8.98026E-05 | 0.00411 | 0.000153 | 0.00187 | 0.000124 | 100 | 0 |

##### H1N1 CA09 HA-N95L (ExpiCHO, nickel)

| F045-092 | | C05 | | S139/1 | | 2D1 | | FluA20 | | H5M9 | |
|---|---|---|---|---|---|---|---|---|---|---|---|
| EC50 | STD | EC50 | STD | EC50 | STD | EC50 | STD | EC50 | STD | EC50 | STD |
| 1.51 | 0.0728 | 100 | 0 | 100 | 0 | 0.00287 | 0.000204 | 0.00367 | 0.000148 | 100 | 0 |

| 16.a.26 | | 429 B01 | | MEDI8852 | | CR9114 | | CR6261 | | CR8020 | |
|---|---|---|---|---|---|---|---|---|---|---|---|
| EC50 | STD | EC50 | STD | EC50 | STD | EC50 | STD | EC50 | STD | EC50 | STD |
| 1.34 | 0.0304 | 0.00632 | 0.00102 | 0.00347 | 0.000157 | 0.00438 | 0.000239 | 0.00202 | 2.12132E-05 | 100 | 0 |

##### H1N1 CA09 HA-N95L-HKE (ExpiCHO, nickel)

| F045-092 | | C05 | | S139/1 | | 2D1 | | FluA20 | | H5M9 | |
|---|---|---|---|---|---|---|---|---|---|---|---|
| EC50 | STD | EC50 | STD | EC50 | STD | EC50 | STD | EC50 | STD | EC50 | STD |
| 1.41 | 0.0127 | 100 | 0 | 0.0243 | 0.00264 | 0.00293 | 0.000228 | 0.00479 | 4.6E-05 | 100 | 0 |

| 16.a.26 | | 429 B01 | | MEDI8852 | | CR9114 | | CR6261 | | CR8020 | |
|---|---|---|---|---|---|---|---|---|---|---|---|
| EC50 | STD | EC50 | STD | EC50 | STD | EC50 | STD | EC50 | STD | EC50 | STD |
| 0.264 | 0.0156 | 0.041 | 7.07E-05 | 0.00705 | 0.00048 | 0.00732 | 0.000486 | 0.003 | 0.000262 | 100 | 0 |

##### H1N1 CA09 HA-ER (E103L/R106M) (ExpiCHO, nickel)

| F045-092 | | C05 | | S139/1 | | 2D1 | | FluA20 | | H5M9 | |
|---|---|---|---|---|---|---|---|---|---|---|---|
| EC50 | STD | EC50 | STD | EC50 | STD | EC50 | STD | EC50 | STD | EC50 | STD |
| 2.85 | 0.0255 | 100 | 0 | 100 | 0 | 0.00276 | 0.00017 | 0.00405 | 9.69E-05 | 100 | 0 |

| 16.a.26 | | 429 B01 | | MEDI8852 | | CR9114 | | CR6261 | | CR8020 | |
|---|---|---|---|---|---|---|---|---|---|---|---|
| EC50 | STD | EC50 | STD | EC50 | STD | EC50 | STD | EC50 | STD | EC50 | STD |
| 2.51 | 0.688 | 0.0146 | 0.00389 | 0.00475 | 0.00036 | 0.00461 | 0.000277 | 0.00259 | 0.00012 | 100 | 0 |

**b**

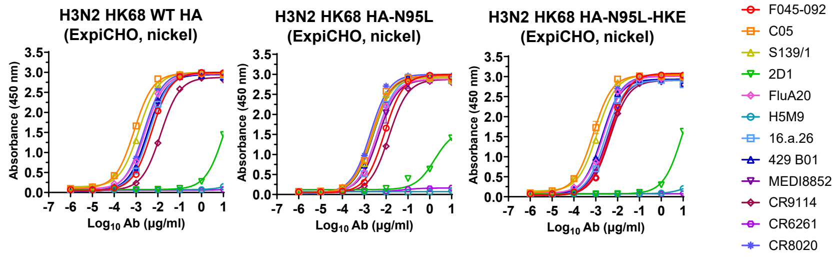

##### H3N2 HK68 WT HA (ExpiCHO, nickel)

| F045-092 | | C05 | | S139/1 | | 2D1 | | FluA20 | | H5M9 | |
|---|---|---|---|---|---|---|---|---|---|---|---|
| EC50 | STD | EC50 | STD | EC50 | STD | EC50 | STD | EC50 | STD | EC50 | STD |
| 0.00524 | 0.000262 | 0.000807 | 4.38E-05 | 0.00139 | 0.000112 | 11.5 | 1.03 | 0.00267 | 0.000217 | 100 | 0 |

| 16.a.26 | | 429 B01 | | MEDI8852 | | CR9114 | | CR6261 | | CR8020 | |
|---|---|---|---|---|---|---|---|---|---|---|---|
| EC50 | STD | EC50 | STD | EC50 | STD | EC50 | STD | EC50 | STD | EC50 | STD |
| 0.00403 | 0.000235 | 0.0033 | 3.68E-05 | 0.00376 | 0.000296 | 0.0135 | 0.000891 | 100 | 0 | 0.00228 | 1.63E-05 |

##### H3N2 HK68 HA-N95L (ExpiCHO, nickel)

| F045-092 | | C05 | | S139/1 | | 2D1 | | FluA20 | | H5M9 | |
|---|---|---|---|---|---|---|---|---|---|---|---|
| EC50 | STD | EC50 | STD | EC50 | STD | EC50 | STD | EC50 | STD | EC50 | STD |
| 0.00834 | 0.000436 | 0.00222 | 0.000104 | 0.00244 | 4.95E-05 | 1.66 | 0.0219 | 0.00369 | 4.38E-05 | 100 | 0 |

| 16.a.26 | | 429 B01 | | MEDI8852 | | CR9114 | | CR6261 | | CR8020 | |
|---|---|---|---|---|---|---|---|---|---|---|---|
| EC50 | STD | EC50 | STD | EC50 | STD | EC50 | STD | EC50 | STD | EC50 | STD |
| 0.00374 | 0.000207 | 0.00261 | 0.000208 | 0.00453 | 0.00137 | 0.0136 | 7.07E-05 | 100 | 0 | 0.00191 | 5.73E-05 |

##### H3N2 HK68 HA-N95L-HKE (ExpiCHO, nickel)

| F045-092 | | C05 | | S139/1 | | 2D1 | | FluA20 | | H5M9 | |
|---|---|---|---|---|---|---|---|---|---|---|---|
| EC50 | STD | EC50 | STD | EC50 | STD | EC50 | STD | EC50 | STD | EC50 | STD |
| 0.00444 | 0.000184 | 0.000773 | 0.000136 | 0.00117 | 2.76E-05 | 10.6 | 2.75 | 0.00227 | 2.33E-05 | 100 | 0 |

| 16.a.26 | | 429 B01 | | MEDI8852 | | CR9114 | | CR6261 | | CR8020 | |
|---|---|---|---|---|---|---|---|---|---|---|---|
| EC50 | STD | EC50 | STD | EC50 | STD | EC50 | STD | EC50 | STD | EC50 | STD |
| 0.00274 | 0.00012 | 0.00195 | 7.35E-05 | 0.00350 | 0.00025 | 0.00442 | 0.000135 | 100 | 0 | 0.00195 | 5.66E-05 |

Fig. S4

c

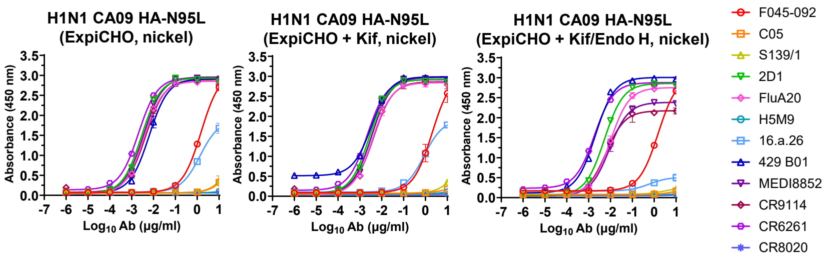

##### H1N1 CA09 HA-N95L (ExpiCHO, nickel)

| F045-092 | | C05 | | S139/1 | | 2D1 | | FluA20 | | H5M9 | |
|---|---|---|---|---|---|---|---|---|---|---|---|
| EC50 | STD | EC50 | STD | EC50 | STD | EC50 | STD | EC50 | STD | EC50 | STD |
| 1.51 | 0.0728 | 100 | 0 | 100 | 0 | 0.00287 | 0.000204 | 0.00367 | 0.000148 | 100 | 0 |

| 16.a.26 | | 429 B01 | | MEDI8852 | | CR9114 | | CR6261 | | CR8020 | |
|---|---|---|---|---|---|---|---|---|---|---|---|
| EC50 | STD | EC50 | STD | EC50 | STD | EC50 | STD | EC50 | STD | EC50 | STD |
| 1.34 | 0.0304 | 0.00632 | 0.00102 | 0.00347 | 0.000157 | 0.00438 | 0.000239 | 0.00202 | 2.12132E-05 | 100 | 0 |

##### H1N1 CA09 HA-N95L (ExpiCHO + KIF, nickel)

| F045-092 | | C05 | | S139/1 | | 2D1 | | FluA20 | | H5M9 | |
|---|---|---|---|---|---|---|---|---|---|---|---|
| EC50 | STD | EC50 | STD | EC50 | STD | EC50 | STD | EC50 | STD | EC50 | STD |
| 2.05 | 0.525 | 100 | 0 | 100 | 0 | 0.00275 | 8.34E-05 | 0.00386 | 0.000422 | 100 | 0 |

| 16.a.26 | | 429 B01 | | MEDI8852 | | CR9114 | | CR6261 | | CR8020 | |
|---|---|---|---|---|---|---|---|---|---|---|---|
| EC50 | STD | EC50 | STD | EC50 | STD | EC50 | STD | EC50 | STD | EC50 | STD |
| 0.869 | 0.140 | 0.00341 | 0.000315 | 0.00348 | 0.000221 | 0.00424 | 4.95E-05 | 0.00218 | 0.000244 | 100 | 0 |

##### H1N1 CA09 HA-N95L (ExpiCHO + Kif/Endo H, nickel)

| F045-092 | | C05 | | S139/1 | | 2D1 | | FluA20 | | H5M9 | |
|---|---|---|---|---|---|---|---|---|---|---|---|
| EC50 | STD | EC50 | STD | EC50 | STD | EC50 | STD | EC50 | STD | EC50 | STD |
| 1.52 | 0.239 | 100 | 0 | 100 | 0 | 0.00529 | 0.000237 | 0.00979 | 0.00128 | 100 | 0 |

| 16.a.26 | | 429 B01 | | MEDI8852 | | CR9114 | | CR6261 | | CR8020 | |
|---|---|---|---|---|---|---|---|---|---|---|---|
| EC50 | STD | EC50 | STD | EC50 | STD | EC50 | STD | EC50 | STD | EC50 | STD |
| 0.587 | 0.138 | 0.00204 | 5.44E-05 | 0.00899 | 0.0015 | 0.00616 | 0.000508 | 0.00184 | 9.05E-05 | 100 | 0 |

d

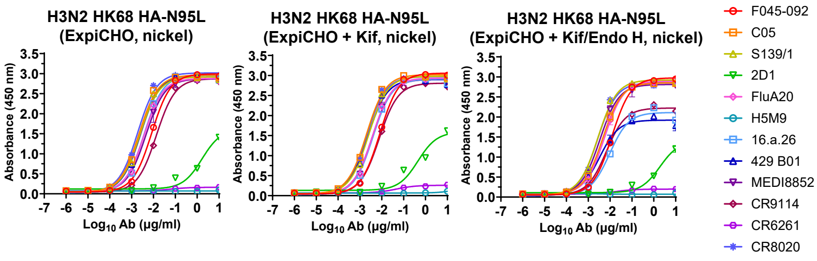

##### H3N2 HK68 HA-N95L (ExpiCHO, nickel)

| F045-092 | | C05 | | S139/1 | | 2D1 | | FluA20 | | H5M9 | |
|---|---|---|---|---|---|---|---|---|---|---|---|
| EC50 | STD | EC50 | STD | EC50 | STD | EC50 | STD | EC50 | STD | EC50 | STD |
| 0.008336 | 0.000436 | 0.00222 | 0.000104 | 0.002441 | 4.95E-05 | 1.655427 | 0.02192 | 0.003694 | 4.38E-05 | 100 | 0 |

| 16.a.26 | | 429 B01 | | MEDI8852 | | CR9114 | | CR6261 | | CR8020 | |
|---|---|---|---|---|---|---|---|---|---|---|---|
| EC50 | STD | EC50 | STD | EC50 | STD | EC50 | STD | EC50 | STD | EC50 | STD |
| 0.003742 | 0.000207 | 0.002605 | 0.000208 | 0.004531 | 0.001365 | 0.01355 | 7.07E-05 | 100 | 0 | 0.001905 | 5.73E-05 |

##### H3N2 HK68 HA-N95L (ExpiCHO + Kif, nickel)

| F045-092 | | C05 | | S139/1 | | 2D1 | | FluA20 | | H5M9 | |
|---|---|---|---|---|---|---|---|---|---|---|---|
| EC50 | STD | EC50 | STD | EC50 | STD | EC50 | STD | EC50 | STD | EC50 | STD |
| 0.008143 | 0.000356 | 0.001999 | 5.16E-05 | 0.002482 | 8.27E-05 | 0.452619 | 0.057629 | 0.003812 | 0.000174 | 100 | 0 |

| 16.a.26 | | 429 B01 | | MEDI8852 | | CR9114 | | CR6261 | | CR8020 | |
|---|---|---|---|---|---|---|---|---|---|---|---|
| EC50 | STD | EC50 | STD | EC50 | STD | EC50 | STD | EC50 | STD | EC50 | STD |
| 0.003971 | 0.000353 | 0.002285 | 0.000214 | 0.003714 | 0.000477 | 0.007597 | 0.000372 | 100 | 0 | 0.00204 | 0.000133 |

##### H3N2 HK68 HA-N95L (ExpiCHO + Kif/Endo H, nickel)

| F045-092 | | C05 | | S139/1 | | 2D1 | | FluA20 | | H5M9 | |
|---|---|---|---|---|---|---|---|---|---|---|---|
| EC50 | STD | EC50 | STD | EC50 | STD | EC50 | STD | EC50 | STD | EC50 | STD |
| 0.012087 | 0.000396 | 0.004569 | 0.000513 | 0.002804 | 0.000134 | 1.916498 | 0.003536 | 0.005528 | 0.00081 | 100 | 0 |

| 16.a.26 | | 429 B01 | | MEDI8852 | | CR9114 | | CR6261 | | CR8020 | |
|---|---|---|---|---|---|---|---|---|---|---|---|
| EC50 | STD | EC50 | STD | EC50 | STD | EC50 | STD | EC50 | STD | EC50 | STD |
| 0.009516 | 0.000181 | 0.003325 | 0.000171 | 0.003414 | 0.000332 | 0.006722 | 0.00024 | 100 | 0 | 0.002662 | 4.6E-05 |

**Fig. S4. Antigenic analyses of H1N2 CA09 and H3N2 HK68 HA constructs with and without glycan modifications.** (**a**) ELISA analysis of CA09 WT HA, HA-ER, HA-N95L, and HA-N95L-HKE trimers binding to 12 antibodies in the IgG form, showing both binding curves (top) and summary of $EC_{50}$ values and standard deviations (bottom). (**b**) ELISA analysis of HK68 WT HA, HA-N95L, and HA-N95L-HKE trimers binding to 12 antibodies in the IgG form, showing both binding curves (top) and summary of $EC_{50}$ values and standard deviations (bottom). (**c**) ELISA analysis of unmodified, Kif-treated, and Kif/Endo H-treated CA09 HA-N95L trimers binding to 12 antibodies in the IgG form, showing both binding curves (top) and summary of $EC_{50}$ values and standard deviations (bottom). (**d**) ELISA analysis of unmodified, Kif-treated, and Kif/endo H-treated HK68 HA-N95L trimers binding to 12 antibodies in the IgG form, showing both binding curves (top) and summary of $EC_{50}$ values and standard deviations (bottom). In ELISA, each well was coated with 0.1 μg of the appropriate antigen, and IgG was diluted in a 10-fold dilution series from a starting concentration of 10 μg/ml for all tested antibodies. Error bars represent the difference between duplicate values at each concentration tested for each sample.

a

**>H1N1-CA09-HA-N95L-FR**

MKAILVVLLYTFATANADTLCIGYHANNSTDTVDTVLEKNVTVTHSVNLLEDKHNGKLCKLRGVAPLHLGKCNIAGWILGNPECESLSTA
SSWSYIVETPSSDNGTCYPGDFIDYEELREQLSSVSSFERFEIFPKTSSWPNHDSNKGVTAACPHAGAKSFYKNLIWLVKKGNSYPKLSK
SYINDKGKEVLVLWGIHHPSTSADQQSLYQNADTYVFVGSSRYSKKFKPEIAIRPKVRDQEGRMNYYWTLVEPGDKITFEATGNLVVPRY
AFAMERNAGSGIIISDTPVHDCNTTCQTPKGAINTSLPFQNIHPITIGKCPKYVKSTKLRLATGLRNIPSIQSRGLFGAIAGFIEGGWTG
MVDGWYGYHHQNEQGSGYAADLKSTQNAIDEITNKVNSVIEKMNTQFTAVGKEFNHLEKRIENLNKKVDDGFLDIWTYNAELLVLLENER
TLDYHDSNVKNLYEKVRSQLKNNAKEIGNGCFEFYHKCDNTCMESVKNGTYDYPKYSEEAKLNREEIDGASDIIKLLNEQVNKEMQSSNL
YMSMSSWCYTHSLDGAGLFLFDHAAEEYEHAKKLIIFLNENNVPVQLTSISAPEHKFEGLTQIFQKAYEHEQHISESINNIVDHAIKSKD
HATFNFLQWYVAEQHEEEVLFKDILDKIELIGNENHGLYLADQYVKGIAKSRKS

**>H1N1-CA09-HA-N95L-E2p-LD4-PADRE**

MKAILVVLLYTFATANADTLCIGYHANNSTDTVDTVLEKNVTVTHSVNLLEDKHNGKLCKLRGVAPLHLGKCNIAGWILGNPECESLSTA
SSWSYIVETPSSDNGTCYPGDFIDYEELREQLSSVSSFERFEIFPKTSSWPNHDSNKGVTAACPHAGAKSFYKNLIWLVKKGNSYPKLSK
SYINDKGKEVLVLWGIHHPSTSADQQSLYQNADTYVFVGSSRYSKKFKPEIAIRPKVRDQEGRMNYYWTLVEPGDKITFEATGNLVVPRY
AFAMERNAGSGIIISDTPVHDCNTTCQTPKGAINTSLPFQNIHPITIGKCPKYVKSTKLRLATGLRNIPSIQSRGLFGAIAGFIEGGWTG
MVDGWYGYHHQNEQGSGYAADLKSTQNAIDEITNKVNSVIEKMNTQFTAVGKEFNHLEKRIENLNKKVDDGFLDIWTYNAELLVLLENER
TLDYHDSNVKNLYEKVRSQLKNNAKEIGNGCFEFYHKCDNTCMESVKNGTYDYPKYSEEAKLNREEIDGASGAAAKPATTEGEFPETREK
MSGIRRAIAKAMVHSKHTAPHVTLMDEADVTKLVAHRKKFKAIAAEKGIKLTFLPYVVKALVSALREYPVLNTAIDDETEEIIQKHYYNI
GIAADTDRGLLVPVIKHADRKPIFALAQEINELAEKARDGKLTPGEMKGASCTITNIGSAGGQWFTPVINHPEVAILGIGRIAEKPIVRD
GEIVAAPMLALSLSFDHRMIDGATAQKALNHIKRLLSDPELLLMGGGGSFSEEQKKALDLAFYFDRRLTPEWRRYLSQRLGLNEEQIERW
FRRKEQQIGWSHPQFEKGSAKFVAAWTLKAAA

**>H1N1-CA09-HA-N95L-I3-01v9b-LD7-PADRE**

MKAILVVLLYTFATANADTLCIGYHANNSTDTVDTVLEKNVTVTHSVNLLEDKHNGKLCKLRGVAPLHLGKCNIAGWILGNPECESLSTA
SSWSYIVETPSSDNGTCYPGDFIDYEELREQLSSVSSFERFEIFPKTSSWPNHDSNKGVTAACPHAGAKSFYKNLIWLVKKGNSYPKLSK
SYINDKGKEVLVLWGIHHPSTSADQQSLYQNADTYVFVGSSRYSKKFKPEIAIRPKVRDQEGRMNYYWTLVEPGDKITFEATGNLVVPRY
AFAMERNAGSGIIISDTPVHDCNTTCQTPKGAINTSLPFQNIHPITIGKCPKYVKSTKLRLATGLRNIPSIQSRGLFGAIAGFIEGGWTG
MVDGWYGYHHQNEQGSGYAADLKSTQNAIDEITNKVNSVIEKMNTQFTAVGKEFNHLEKRIENLNKKVDDGFLDIWTYNAELLVLLENER
TLDYHDSNVKNLYEKVRSQLKNNAKEIGNGCFEFYHKCDNTCMESVKNGTYDYPKYSEEAKLNREEIDGASGAEKMIKEIGSGSEELQKK
MEELFKKHKIVAVLRANSVEEAKMKALAVFVGGVHLIEITFTVPDADTVIKELSFLKELGAIIGAGTVTSVEQCRKAVESGAEFIVSPHL
DAEITVFCLEKGVFYMPGVMTPTELVKAMKLGHNILKLFPGEVVGPQFVKAMKGPFPNVKFVPTGGVNLDNVCEWFKAGVLAVGVGSALV
KGTIAEVAAKAAAFVEKIRGCTEGGGGSSPAVDIGDRLDELEKALEALSAEDGHDDVGQRLESLLRRWNSRRADGSAKFVAAWTLKAAA

: Leader sequence
: N95L mutation
G: Linker; **AS**: Enzymatic site
: FR, E2p-LD4-PADRE (E2p-L4P), I3-01v9b-LD7-PADRE (I3-01v9b-L7P)

b

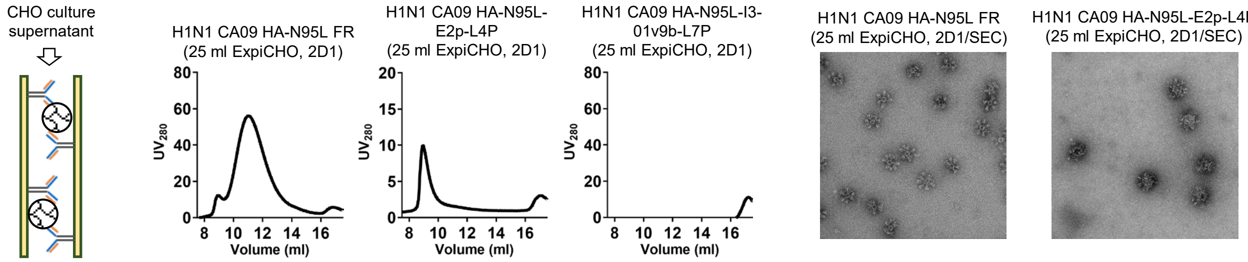

c

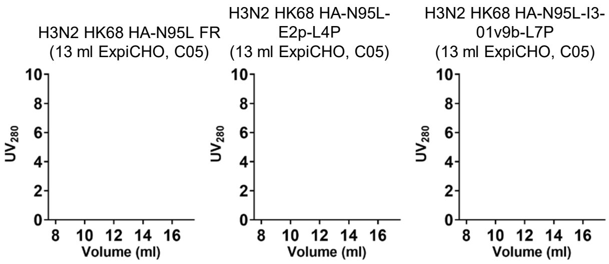

### Fig. S5

d

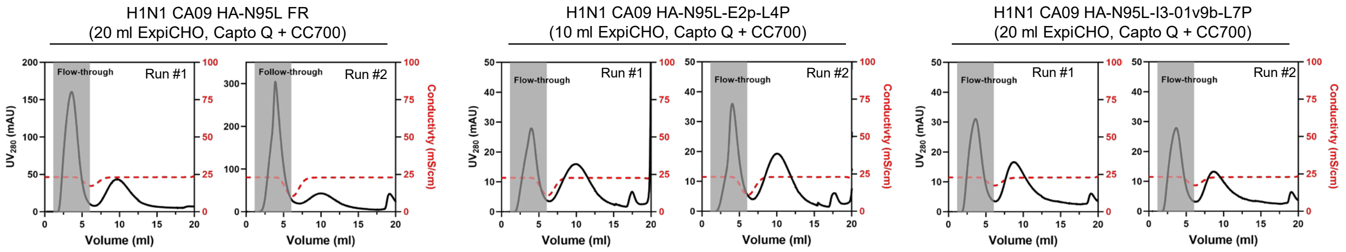
H1N1 CA09 HA-N95L FR
(20 ml ExpiCHO, Capto Q + CC700)
H1N1 CA09 HA-N95L-E2p-L4P
(10 ml ExpiCHO, Capto Q + CC700)
H1N1 CA09 HA-N95L-I3-01v9b-L7P
(20 ml ExpiCHO, Capto Q + CC700)
Flow-through
Follow-through
Run #1
Run #2
UV280 (mAU)
Conductivity (mS/cm)
Volume (ml)

e

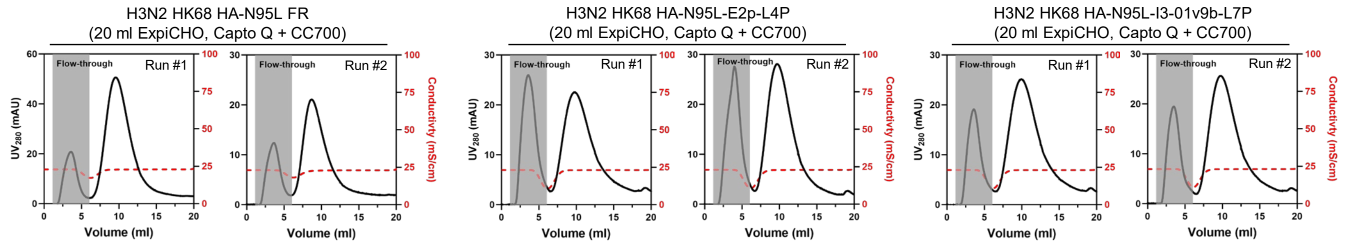
H3N2 HK68 HA-N95L FR
(20 ml ExpiCHO, Capto Q + CC700)
H3N2 HK68 HA-N95L-E2p-L4P
(20 ml ExpiCHO, Capto Q + CC700)
H3N2 HK68 HA-N95L-I3-01v9b-L7P
(20 ml ExpiCHO, Capto Q + CC700)
Flow-through
Run #1
Run #2
UV280 (mAU)
Conductivity (mS/cm)
Volume (ml)

f

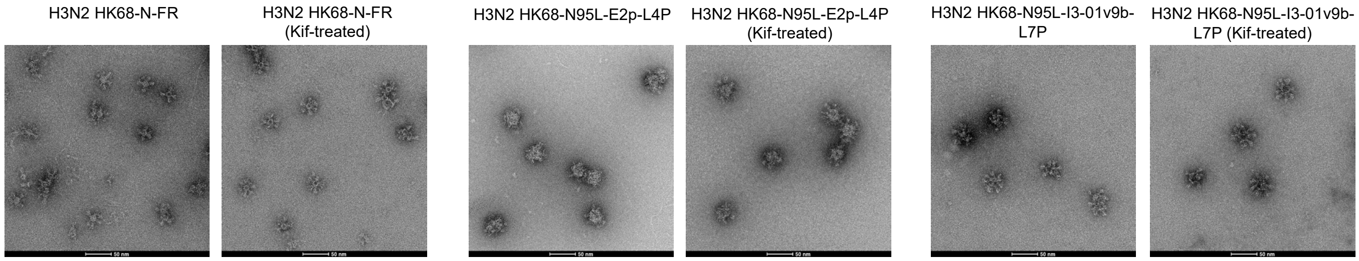
H3N2 HK68-N-FR
H3N2 HK68-N-FR (Kif-treated)
H3N2 HK68-N95L-E2p-L4P
H3N2 HK68-N95L-E2p-L4P (Kif-treated)
H3N2 HK68-N95L-I3-01v9b-L7P
H3N2 HK68-N95L-I3-01v9b-L7P (Kif-treated)

g

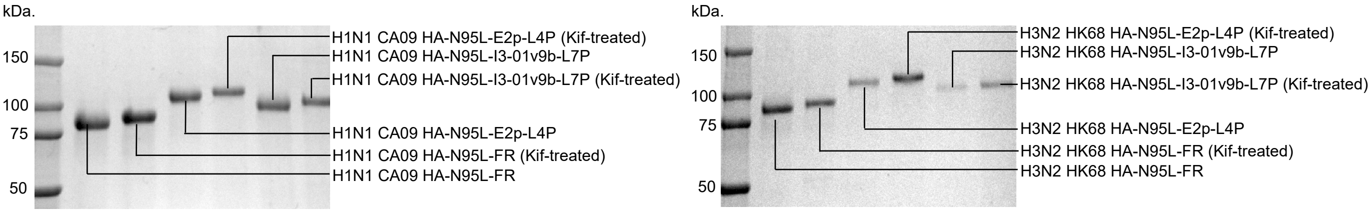
kDa.
150
100
75
50
H1N1 CA09 HA-N95L-E2p-L4P (Kif-treated)
H1N1 CA09 HA-N95L-I3-01v9b-L7P
H1N1 CA09 HA-N95L-I3-01v9b-L7P (Kif-treated)
H1N1 CA09 HA-N95L-E2p-L4P
H1N1 CA09 HA-N95L-FR (Kif-treated)
H1N1 CA09 HA-N95L-FR
H3N2 HK68 HA-N95L-E2p-L4P (Kif-treated)
H3N2 HK68 HA-N95L-I3-01v9b-L7P
H3N2 HK68 HA-N95L-I3-01v9b-L7P (Kif-treated)
H3N2 HK68 HA-N95L-E2p-L4P
H3N2 HK68 HA-N95L-FR (Kif-treated)
H3N2 HK68 HA-N95L-FR

h

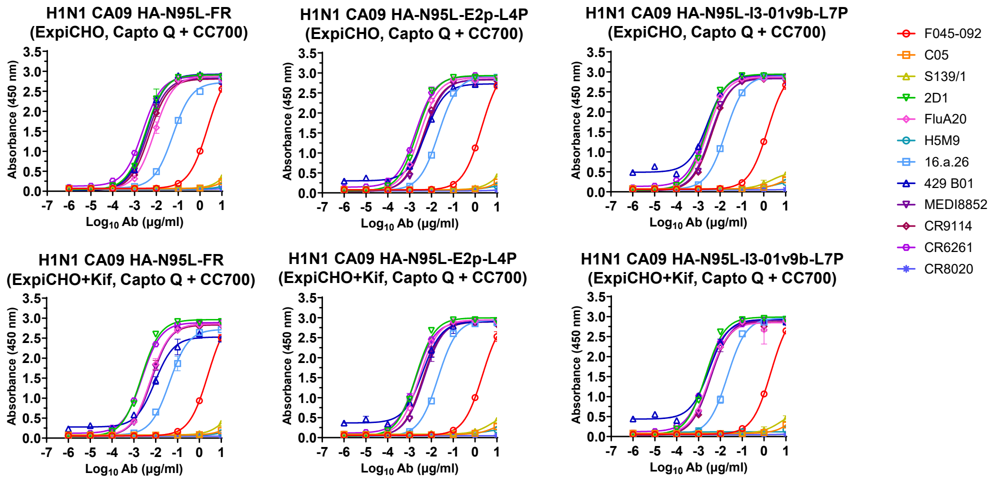
H1N1 CA09 HA-N95L-FR (ExpiCHO, Capto Q + CC700)
H1N1 CA09 HA-N95L-E2p-L4P (ExpiCHO, Capto Q + CC700)
H1N1 CA09 HA-N95L-I3-01v9b-L7P (ExpiCHO, Capto Q + CC700)
H1N1 CA09 HA-N95L-FR (ExpiCHO+Kif, Capto Q + CC700)
H1N1 CA09 HA-N95L-E2p-L4P (ExpiCHO+Kif, Capto Q + CC700)
H1N1 CA09 HA-N95L-I3-01v9b-L7P (ExpiCHO+Kif, Capto Q + CC700)
Absorbance (450 nm)
Log10 Ab (μg/ml)
F045-092
C05
S139/1
2D1
FluA20
H5M9
16.a.26
429 B01
MEDI8852
CR9114
CR6261
CR8020

### Fig. S5

##### H1N1 CA09 HA-N95L-FR (ExpiCHO, Capto Q + CC700)

| F045-092 | | C05 | | S139/1 | | 2D1 | | FluA20 | | H5M9 | |
|---|---|---|---|---|---|---|---|---|---|---|---|
| EC50 | STD | EC50 | STD | EC50 | STD | EC50 | STD | EC50 | STD | EC50 | STD |
| 2.26 | 0.188 | 100 | 0 | 100 | 0 | 0.00323 | 0.000630 | 0.00818 | 0.001161069 | 100 | 0 |

| 16.a.26 | | 429 B01 | | MEDI8852 | | CR9114 | | CR6261 | | CR8020 | |
|---|---|---|---|---|---|---|---|---|---|---|---|
| EC50 | STD | EC50 | STD | EC50 | STD | EC50 | STD | EC50 | STD | EC50 | STD |
| 0.0568 | 0.00443 | 0.00348 | 0.000339 | 0.00394 | 0.000331 | 0.00481 | 0.000759 | 0.00237 | 8.20244E-05 | 100 | 0 |

##### H1N1 CA09 HA-N95L-FR (ExpiCHO + Kif, Capto Q + CC700)

| F045-092 | | C05 | | S139/1 | | 2D1 | | FluA20 | | H5M9 | |
|---|---|---|---|---|---|---|---|---|---|---|---|
| EC50 | STD | EC50 | STD | EC50 | STD | EC50 | STD | EC50 | STD | EC50 | STD |
| 2.49 | 0.0898 | 100 | 0 | 100 | 0 | 0.00217 | 0.000214 | 0.00588 | 0.001 | 100 | 0 |

| 16.a.26 | | 429 B01 | | MEDI8852 | | CR9114 | | CR6261 | | CR8020 | |
|---|---|---|---|---|---|---|---|---|---|---|---|
| EC50 | STD | EC50 | STD | EC50 | STD | EC50 | STD | EC50 | STD | EC50 | STD |
| 0.0436 | 0.00931 | 0.00926 | 0.00168 | 0.00646 | 0.000819 | 0.00576 | 0.00081 | 0.00237 | 7.92E-05 | 100 | 0 |

##### H1N1 CA09 HA-N95L-E2p-L4P (ExpiCHO, Capto Q + CC700)

| F045-092 | | C05 | | S139/1 | | 2D1 | | FluA20 | | H5M9 | |
|---|---|---|---|---|---|---|---|---|---|---|---|
| EC50 | STD | EC50 | STD | EC50 | STD | EC50 | STD | EC50 | STD | EC50 | STD |
| 1.97 | 0.161 | 100 | 0 | 100 | 0 | 0.00216 | 6.01E-05 | 0.00293 | 0.000238 | 100 | 0 |

| 16.a.26 | | 429 B01 | | MEDI8852 | | CR9114 | | CR6261 | | CR8020 | |
|---|---|---|---|---|---|---|---|---|---|---|---|
| EC50 | STD | EC50 | STD | EC50 | STD | EC50 | STD | EC50 | STD | EC50 | STD |
| 0.0211 | 0.000445 | 0.00570 | 0.000313 | 0.00483 | 0.000129 | 0.00418 | 7.42E-05 | 0.00192 | 0.000167 | 100 | 0 |

##### H1N1 CA09 HA-N95L-E2p-L4P (ExpiCHO + Kif, Capto Q + CC700)

| F045-092 | | C05 | | S139/1 | | 2D1 | | FluA20 | | H5M9 | |
|---|---|---|---|---|---|---|---|---|---|---|---|
| EC50 | STD | EC50 | STD | EC50 | STD | EC50 | STD | EC50 | STD | EC50 | STD |
| 2.19 | 0.353 | 100 | 0 | 100 | 0 | 0.00193 | 4.45477E-05 | 0.00278 | 0.000122 | 100 | 0 |

| 16.a.26 | | 429 B01 | | MEDI8852 | | CR9114 | | CR6261 | | CR8020 | |
|---|---|---|---|---|---|---|---|---|---|---|---|
| EC50 | STD | EC50 | STD | EC50 | STD | EC50 | STD | EC50 | STD | EC50 | STD |
| 0.0201 | 0.00128 | 0.00401 | 0.000684 | 0.00458 | 0.000834 | 0.00401 | 0.000561 | 0.002 | 0.000204 | 100 | 0 |

##### H1N1 CA09 HA-N95L-I3-01v9b-L7P (ExpiCHO, Capto Q + CC700)

| F045-092 | | C05 | | S139/1 | | 2D1 | | FluA20 | | H5M9 | |
|---|---|---|---|---|---|---|---|---|---|---|---|
| EC50 | STD | EC50 | STD | EC50 | STD | EC50 | STD | EC50 | STD | EC50 | STD |
| 1.53 | 0.188 | 100 | 0 | 100 | 0 | 0.00225 | 2.83E-05 | 0.00256 | 0.000231 | 100 | 0 |

| 16.a.26 | | 429 B01 | | MEDI8852 | | CR9114 | | CR6261 | | CR8020 | |
|---|---|---|---|---|---|---|---|---|---|---|---|
| EC50 | STD | EC50 | STD | EC50 | STD | EC50 | STD | EC50 | STD | EC50 | STD |
| 0.0179 | 0.00139 | 0.00265 | 9.26E-05 | 0.003672 | 4.6E-05 | 0.00411 | 0.000239 | 0.00216 | 4.1E-05 | 100 | 0 |

##### H1N1 CA09 HA-N95L-I3-01v9b-L7P (ExpiCHO + Kif, Capto Q + CC700)

| F045-092 | | C05 | | S139/1 | | 2D1 | | FluA20 | | H5M9 | |
|---|---|---|---|---|---|---|---|---|---|---|---|
| EC50 | STD | EC50 | STD | EC50 | STD | EC50 | STD | EC50 | STD | EC50 | STD |
| 2.10 | 0.0438 | 100 | 0 | 100 | 0 | 0.00210 | 8.49E-06 | 0.00314 | 0.000672 | 100 | 0 |

| 16.a.26 | | 429 B01 | | MEDI8852 | | CR9114 | | CR6261 | | CR8020 | |
|---|---|---|---|---|---|---|---|---|---|---|---|
| EC50 | STD | EC50 | STD | EC50 | STD | EC50 | STD | EC50 | STD | EC50 | STD |
| 0.0211 | 0.00241 | 0.00314 | 0.0003 | 0.00364 | 0.000351 | 0.00344 | 6.15E-05 | 0.00211 | 0.000205 | 100 | 0 |

i

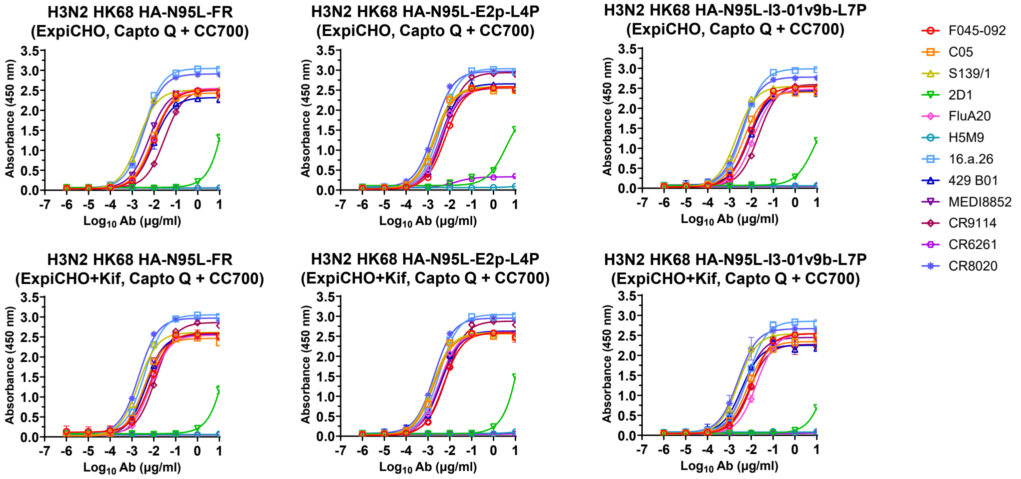

**Fig. S5**

**H3N2 HK68 HA-N95L-FR (ExpiCHO, Capto Q + CC700)**

| F045-092 | | C05 | | S139/1 | | 2D1 | | FluA20 | | H5M9 | |
|---|---|---|---|---|---|---|---|---|---|---|---|
| EC50 | STD | EC50 | STD | EC50 | STD | EC50 | STD | EC50 | STD | EC50 | STD |
| 0.00999 | 0.00118 | 0.00790 | 0.000706 | 0.00194 | 5.44472E-05 | 50.7 | 4.12 | 0.00979 | 0.000127 | 100 | 0 |

| 16.a.26 | | 429 B01 | | MEDI8852 | | CR9114 | | CR6261 | | CR8020 | |
|---|---|---|---|---|---|---|---|---|---|---|---|
| EC50 | STD | EC50 | STD | EC50 | STD | EC50 | STD | EC50 | STD | EC50 | STD |
| 0.00339 | 0.000177 | 0.00970 | 0.00202 | 0.00571 | 9.54594E-05 | 0.0304 | 0.00119 | 100 | 0 | 0.00345 | 0.000102 |

**H3N2 HK68 HA-N95L-FR (ExpiCHO + Kif, Capto Q + CC700)**

| F045-092 | | C05 | | S139/1 | | 2D1 | | FluA20 | | H5M9 | |
|---|---|---|---|---|---|---|---|---|---|---|---|
| EC50 | STD | EC50 | STD | EC50 | STD | EC50 | STD | EC50 | STD | EC50 | STD |
| 0.00701 | 0.000207 | 0.00359 | 0.000443 | 0.002 | 4.95E-05 | 39.4 | 17.3 | 0.00710 | 0.000202 | 100 | 0 |

| 16.a.26 | | 429 B01 | | MEDI8852 | | CR9114 | | CR6261 | | CR8020 | |
|---|---|---|---|---|---|---|---|---|---|---|---|
| EC50 | STD | EC50 | STD | EC50 | STD | EC50 | STD | EC50 | STD | EC50 | STD |
| 0.00375 | 7.5E-05 | 0.00471 | 0.000483 | 0.00391 | 0.000175 | 0.0121 | 0.000516 | 100 | 0 | 0.00204 | 4.81E-05 |

**H3N2 HK68 HA-N95L-E2p-L4P (ExpiCHO, Capto Q + CC700)**

| F045-092 | | C05 | | S139/1 | | 2D1 | | FluA20 | | H5M9 | |
|---|---|---|---|---|---|---|---|---|---|---|---|
| EC50 | STD | EC50 | STD | EC50 | STD | EC50 | STD | EC50 | STD | EC50 | STD |
| 0.00622 | 0.000107 | 0.00217 | 1.13E-05 | 0.00190 | 2.9E-05 | 3.78 | 0.290621 | 0.00381 | 2.83E-06 | 100 | 0 |

| 16.a.26 | | 429 B01 | | MEDI8852 | | CR9114 | | CR6261 | | CR8020 | |
|---|---|---|---|---|---|---|---|---|---|---|---|
| EC50 | STD | EC50 | STD | EC50 | STD | EC50 | STD | EC50 | STD | EC50 | STD |
| 0.00370 | 0.000207 | 0.00388 | 0.000169 | 0.00421 | 9.9E-06 | 0.00610 | 0.000311 | 100 | 0 | 0.00192 | 1.27E-05 |

**H3N2 HK68 HA-N95L-E2p-L4P (ExpiCHO + Kif, Capto Q + CC700)**

| F045-092 | | C05 | | S139/1 | | 2D1 | | FluA20 | | H5M9 | |
|---|---|---|---|---|---|---|---|---|---|---|---|
| EC50 | STD | EC50 | STD | EC50 | STD | EC50 | STD | EC50 | STD | EC50 | STD |
| 0.00529 | 0.000323 | 0.00188 | 8.2E-05 | 0.00165 | 0.000105 | 35.0 | 0.552 | 0.00314 | 0.000115 | 100 | 0 |

| 16.a.26 | | 429 B01 | | MEDI8852 | | CR9114 | | CR6261 | | CR8020 | |
|---|---|---|---|---|---|---|---|---|---|---|---|
| EC50 | STD | EC50 | STD | EC50 | STD | EC50 | STD | EC50 | STD | EC50 | STD |
| 0.00326 | 0.00031 | 0.00356 | 0.000346 | 0.00382 | 0.000453 | 0.00685 | 0.000388 | 100 | 0 | 0.0019 | 1.77E-05 |

**H3N2 HK68 HA-N95L-I3-01v9b-L7P (ExpiCHO, Capto Q + CC700)**

| F045-092 | | C05 | | S139/1 | | 2D1 | | FluA20 | | H5M9 | |
|---|---|---|---|---|---|---|---|---|---|---|---|
| EC50 | STD | EC50 | STD | EC50 | STD | EC50 | STD | EC50 | STD | EC50 | STD |
| 0.00780 | 0.000535 | 0.00433 | 0.00012 | 0.00211 | 5.59E-05 | 8.22 | 0.7064 | 0.0122 | 4.95E-05 | 100 | 0 |

| 16.a.26 | | 429 B01 | | MEDI8852 | | CR9114 | | CR6261 | | CR8020 | |
|---|---|---|---|---|---|---|---|---|---|---|---|
| EC50 | STD | EC50 | STD | EC50 | STD | EC50 | STD | EC50 | STD | EC50 | STD |
| 0.00457 | 0.000322 | 0.00815 | 0.00049 | 0.00703 | 0.000122 | 0.0211 | 0.000686 | 100 | 0 | 0.00345 | 0.000101 |

**H3N2 HK68 HA-N95L-I3-01v9b-L7P (ExpiCHO + Kif, Capto Q + CC700)**

| F045-092 | | C05 | | S139/1 | | 2D1 | | FluA20 | | H5M9 | |
|---|---|---|---|---|---|---|---|---|---|---|---|
| EC50 | STD | EC50 | STD | EC50 | STD | EC50 | STD | EC50 | STD | EC50 | STD |
| 0.0111 | 0.0007 | 0.00683 | 0.000561 | 0.00251 | 3.68E-05 | 56.9 | 24.9 | 0.0176 | 0.00238 | 100 | 0 |

| 16.a.26 | | 429 B01 | | MEDI8852 | | CR9114 | | CR6261 | | CR8020 | |
|---|---|---|---|---|---|---|---|---|---|---|---|
| EC50 | STD | EC50 | STD | EC50 | STD | EC50 | STD | EC50 | STD | EC50 | STD |
| 0.00711 | 0.000414 | 0.00362 | 0.000178 | 0.00509 | 0.000262 | 0.00841 | 0.000447 | 100 | 0 | 0.00253 | 0.00102 |

**Fig. S5. Design and in vitro characterization of HA-presenting 1c-SApNPs.** (**a**) Amino acid sequences of CA09 HA-N95L-presenting FR, E2p-LD4-PADRE (or E2p-L4P), and I3-01v9b-LD7-PADRE (or I3-01v9b-L7P). FR forms a 24-mer, while E2p-L4P and I3-01v9b-L7P assemble into multilayered 60-mers. The signal peptide, N95L mutation, restriction site (AS), linker (GS), and SApNP backbone are highlighted in yellow, red, blue font, magenta, and cyan, respectively. (**b**) 2D1-based IAC purification of CA09 1c-SApNPs. Left: Schematic of a 2D1-based IAC column for 1c-SApNP purification. Middle: SEC profiles of 2D1-purified CA09 HA-N95L FR, E2p-L4P, and I3-01v9b-L7P materials obtained from a Superose 6 10/300 GL column. Right: nsEM micrographs of 2D1/SEC-purified CA09 HA-N95L FR and E2p-L4P 1c-SApNPs. (**c**) SEC profiles of C05-purified HK68 HA-N95L FR, E2p-L4P, and I3-01v9b-L7P materials obtained from a Superose 6 10/300 GL column. (**d,e**) Post-CC700 flow-through chromatographic profiles for CA09 and HK68 HA-N95L-presenting FR, E2p-L4P, and I3-01v9b-L7P samples obtained from an ÄKTA pure 25 M instrument. Of note, two runs are shown for each sample. (**f**) nsEM micrographs of unmodified and Kif-treated IEX-purified HK68 HA-N95L FR, E2p-L4P, and I3-01v9b-L7P 1c-SApNPs. (**g**) SDS-PAGE gels of unmodified and Kif-treated HA-N95L FR, E2p-L4P, and I3-01v9b-L7P 1c-SApNPs derived from CA09 (left) and HK68 (right). Bands are labeled on the gels. (**h**) ELISA analysis of unmodified and Kif-treated IEX-purified CA09 HA-N95L-presenting FR, E2p-L4P, and I3-01v9b-L7P 1c-SApNPs binding to 12 antibodies in the IgG form, showing both binding curves (top) and summary of $EC_{50}$ values and standard deviations (bottom). (**i**) ELISA analysis of unmodified and Kif-treated IEX-purified HK68 HA-N95L-presenting FR, E2p-L4P, and I3-01v9b-L7P 1c-SApNPs binding to 12 antibodies in the IgG form, showing both binding curves (top) and summary of $EC_{50}$ values and standard deviations (bottom). The ELISA data shown in (**h, i**) were obtained as follows. Briefly, each well was coated with 0.1 μg of the appropriate antigen, and IgG was diluted in a 10-fold dilution series from a starting concentration of 10 μg/ml for all tested antibodies. Error bars represent the difference between these duplicate values at each concentration tested for each sample.

**Fig. S6**

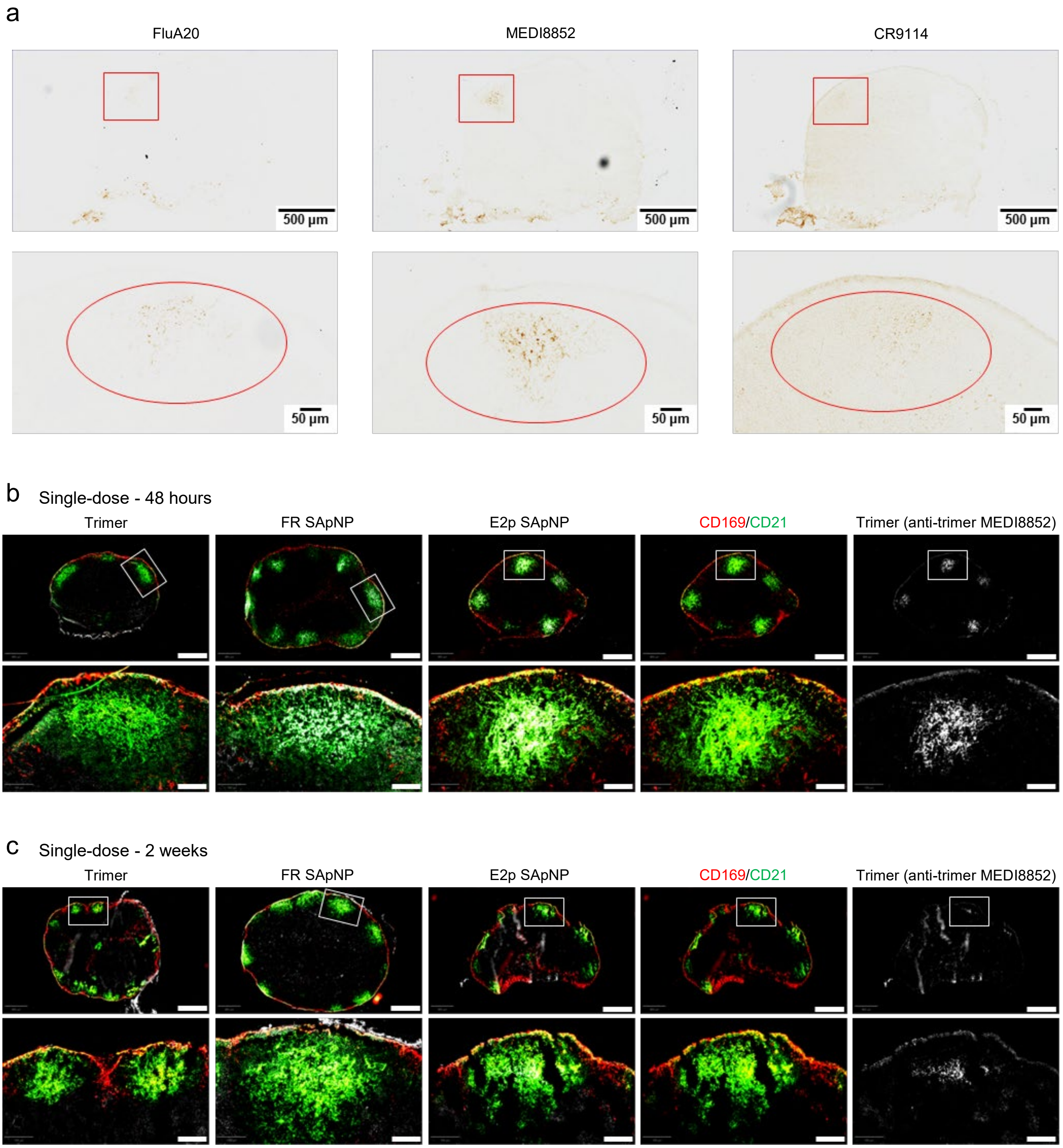
a
FluA20
MEDI8852
CR9114
500 µm
50 µm
b
Single-dose - 48 hours
Trimer
FR SApNP
E2p SApNP
CD169/CD21
Trimer (anti-trimer MEDI8852)
c
Single-dose - 2 weeks
Trimer
FR SApNP
E2p SApNP
CD169/CD21
Trimer (anti-trimer MEDI8852)

**Fig. S6**

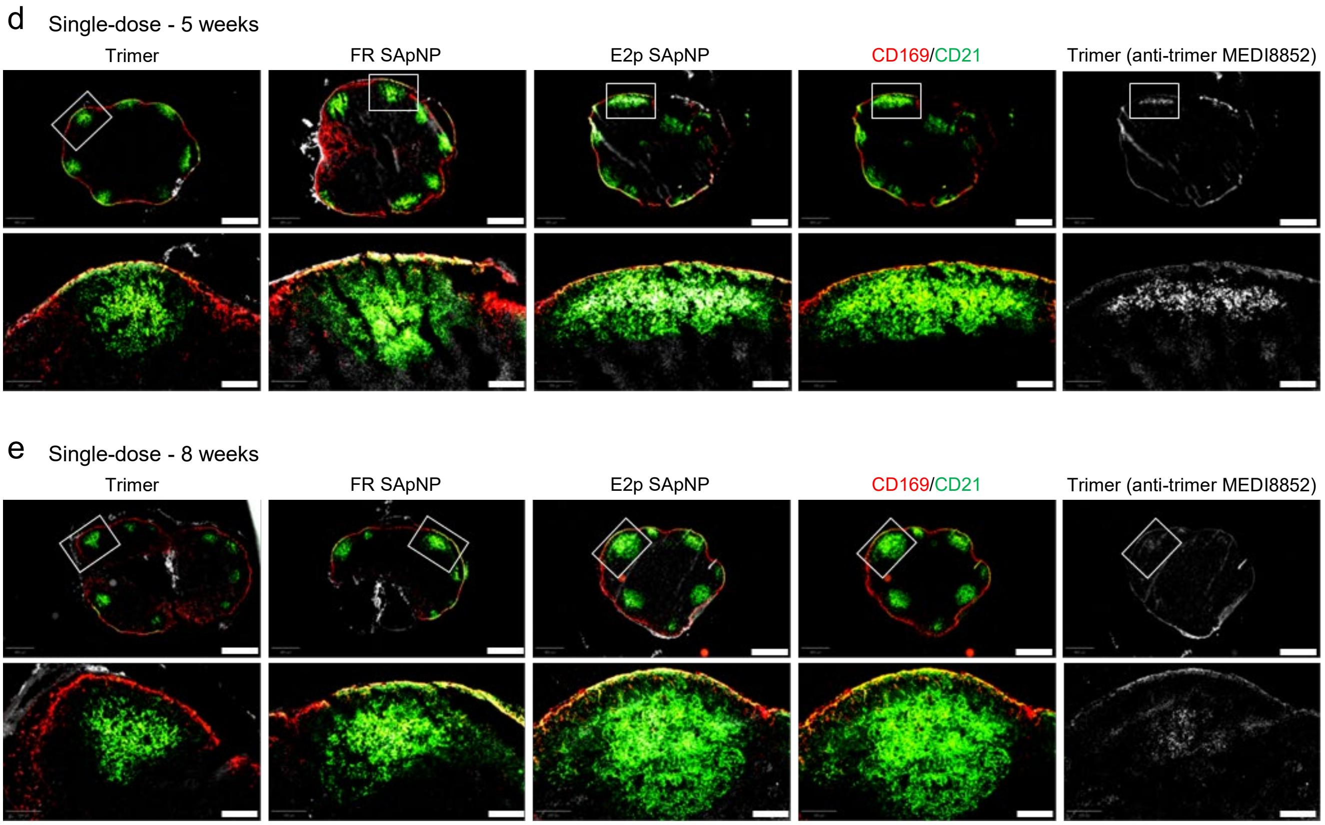

**Fig. S6. Immunohistological images of HA trimer and SApNPs in lymph nodes.** (**a**) Immunostaining images of lymph node tissues from mice injected with HA trimer-presenting E2p SApNPs at 48 hours post-injection, stained using three human anti-trimer antibodies FluA20, MEDI8852, and CR9914. (**b-e**) Colocalization of HA trimer and HA trimer-presenting FR and E2p SApNPs with FDC networks in lymph node follicles at various time points following a single-dose injection (10 µg per injection, 40 µg total per mouse, n = 3-5 mice/group): (**b**) 48 hours, (**c**) 2 weeks, (**d**) 5 weeks, and (**e**) 8 weeks. Immunofluorescent images are pseudo-color-coded as follows: $CD21^+$ (green), $CD169^+$ (red), and anti-trimer MEDI8852 (white). Scale bars: 500 µm (entire lymph node) and 100 µm (enlarged follicle).

**Fig. S7**

a FR SApNPs (yellow arrow) and AddaVax particles (green arrow) aligned on FDC dendrites at 12 hours

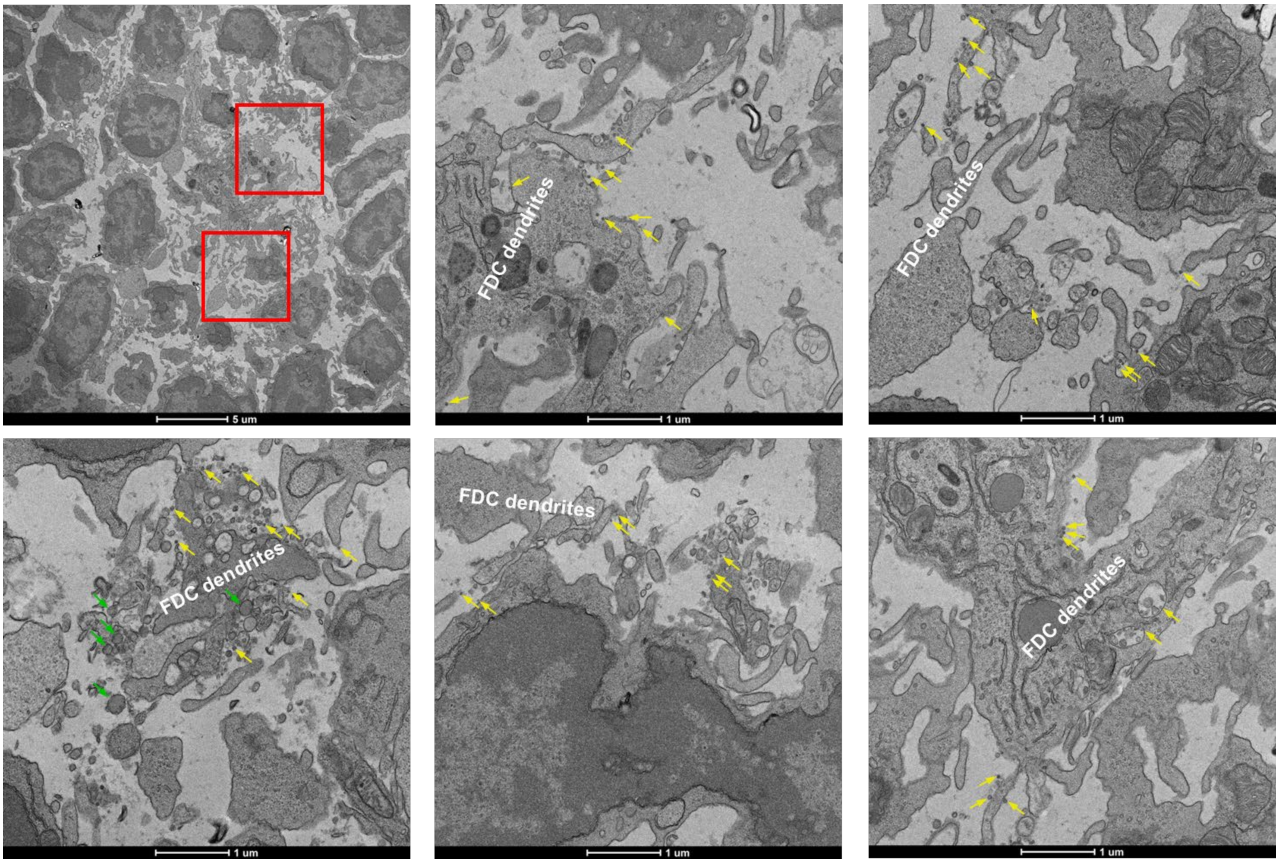

b E2p SApNPs (yellow arrow) aligned on FDC dendrites at 12 hours

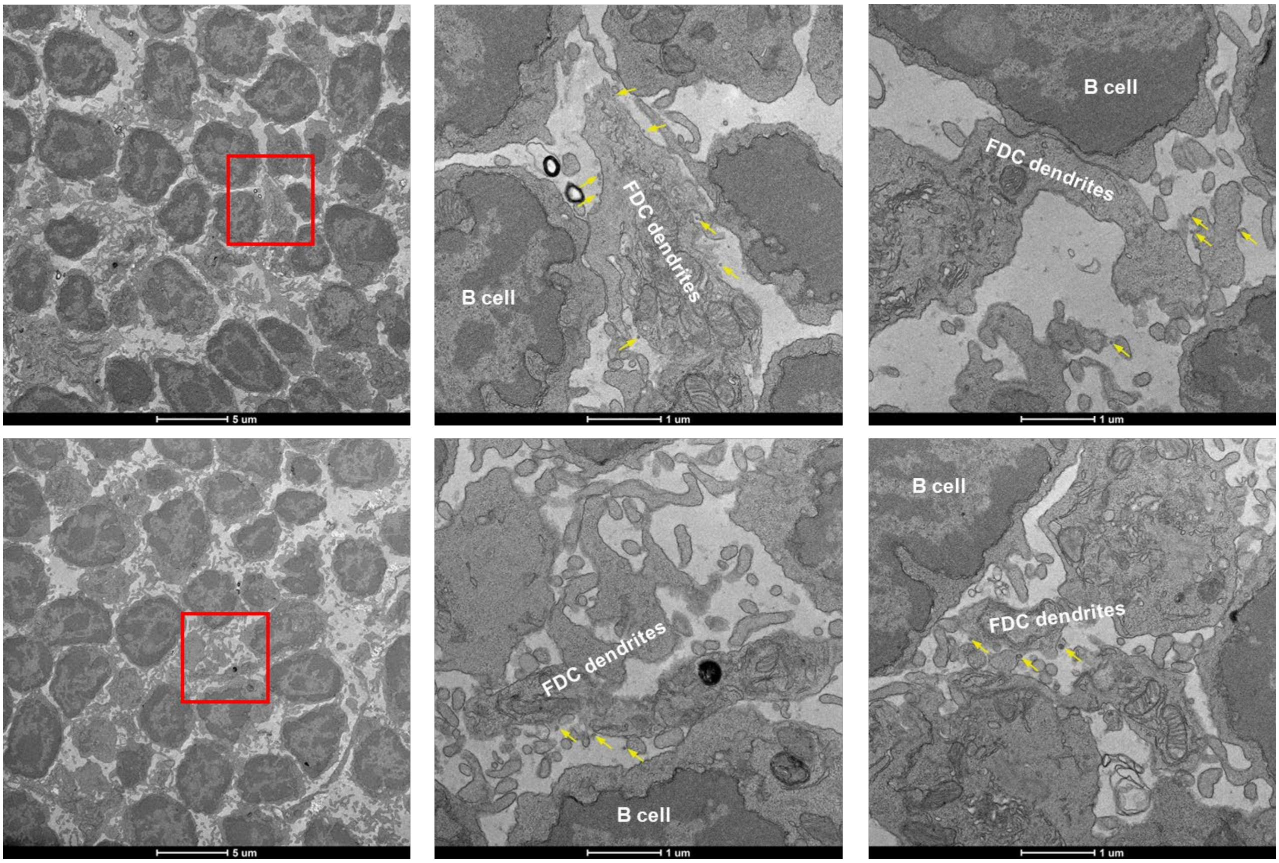

c FR SApNPs (yellow arrow) and AddaVax particles (green arrow) aligned on FDC dendrites at 48 hours

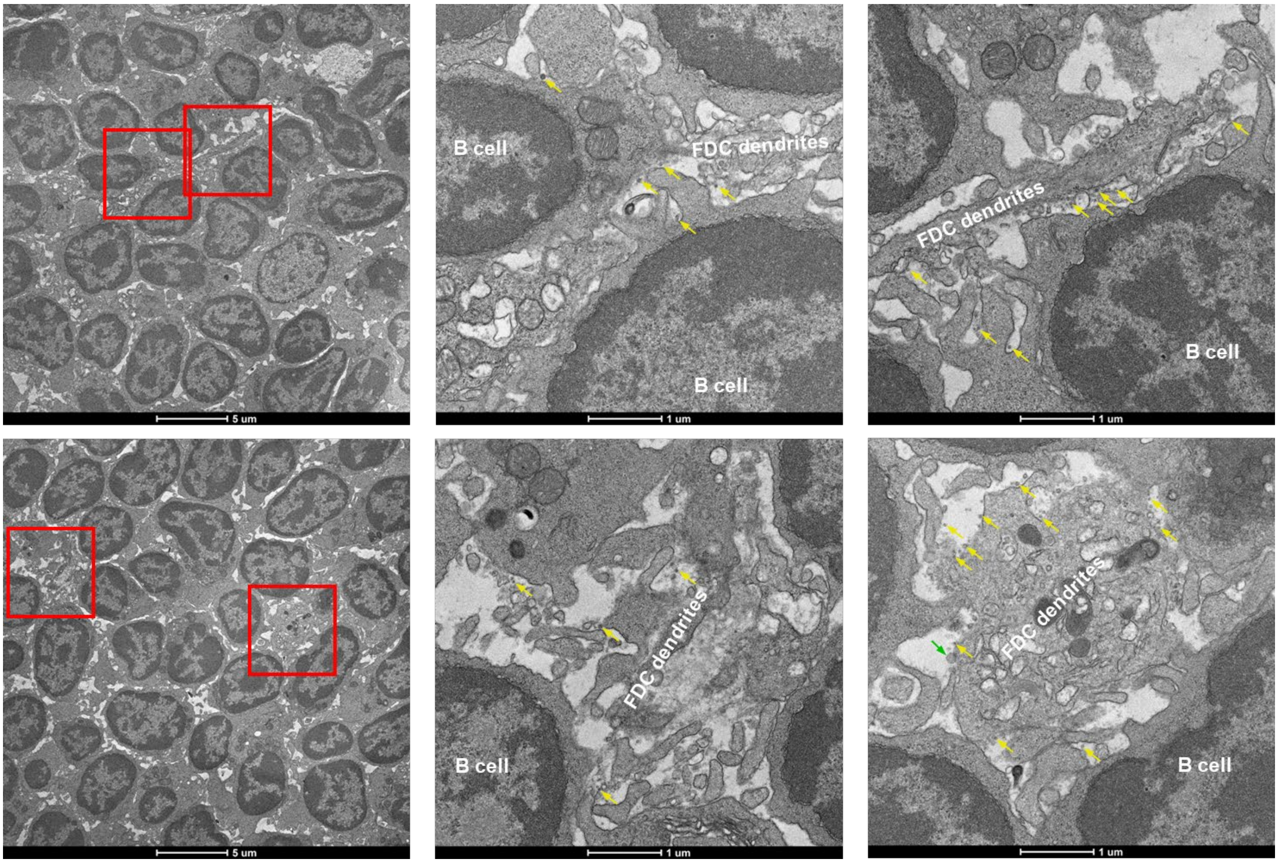

d E2p SApNPs (yellow arrow) aligned on FDC dendrites at 48 hours

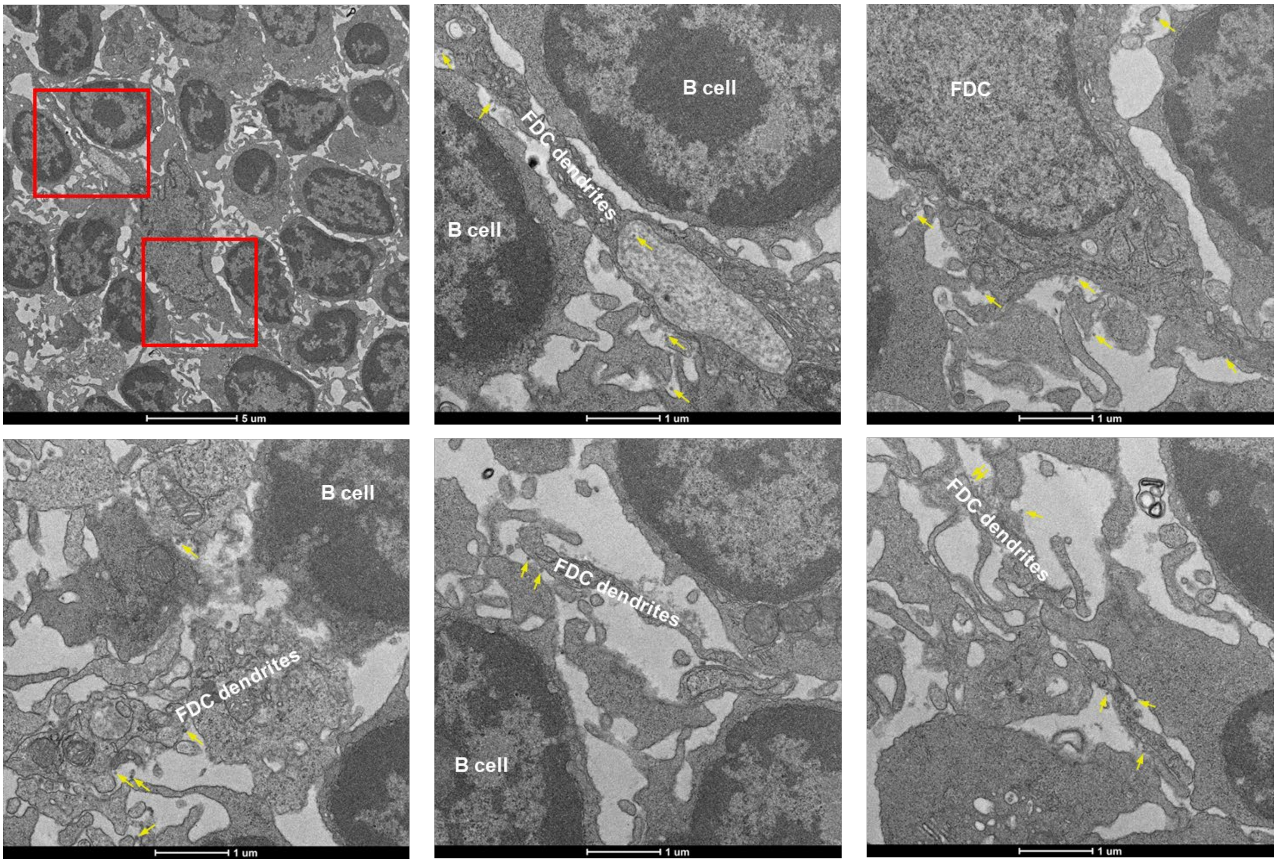

**Fig. S7**

e FR SApNPs (yellow arrow) interact with B cells at 48 hours

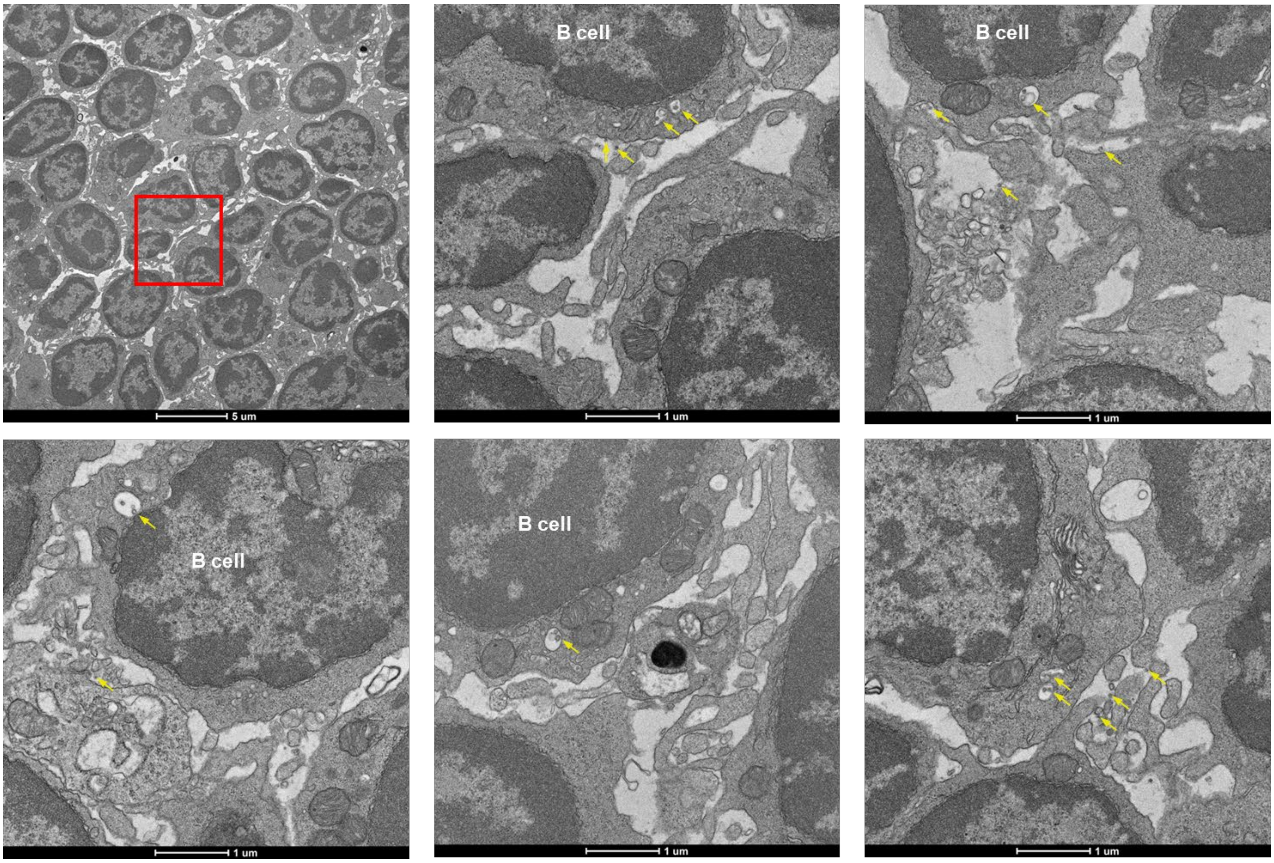

f E2p SApNPs (yellow arrow) interact with B cells at 48 hours

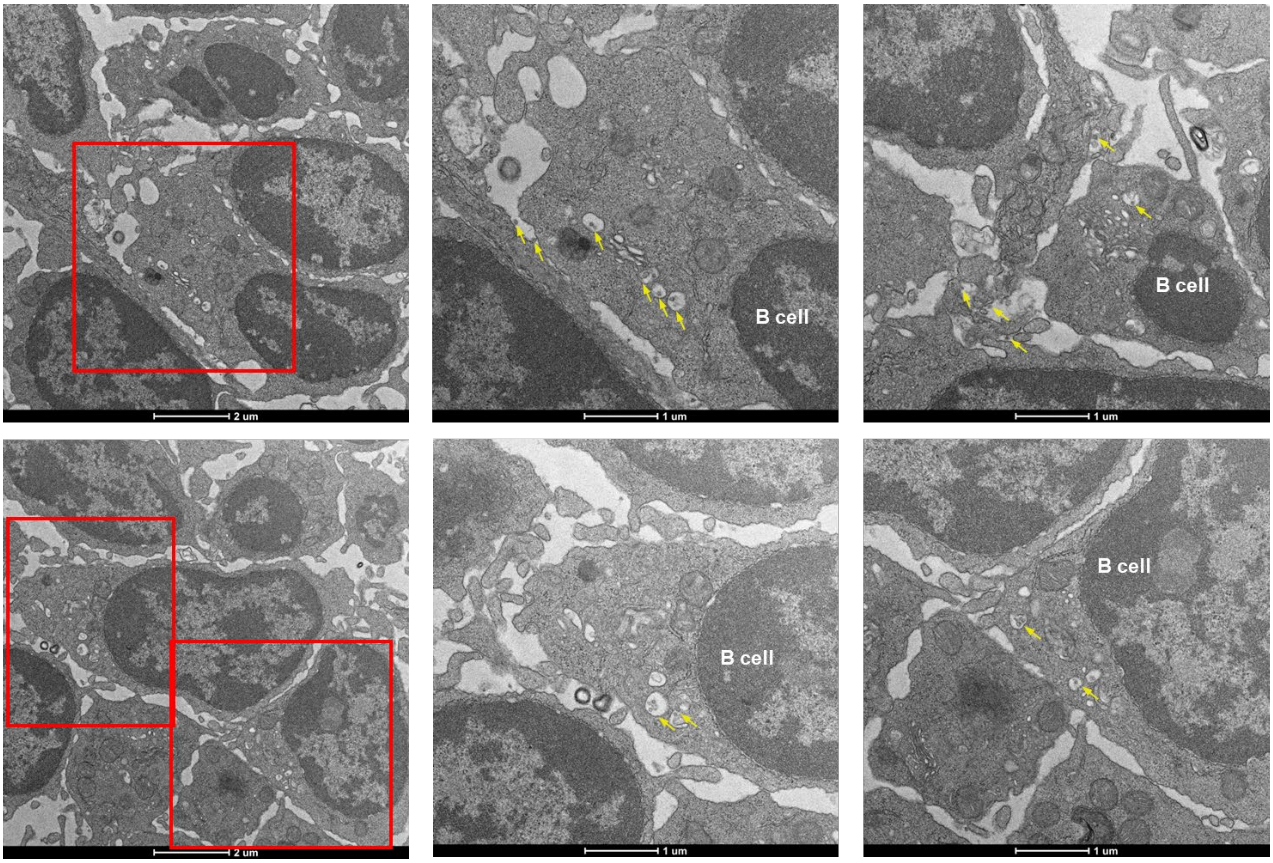

**Fig. S7. TEM images of HA-presenting SApNPs interacting with FDCs and B cells in lymph nodes.** HA-presenting FR or E2p SApNPs and AddaVax (AV) particles aligned on FDC dendrites at (**a**-**b**) 12 hours and (**c**-**d**) 48 hours after a single-dose injection (2 footpads, 50 µg/footpad). (**e**) FR or (**f**) E2p SApNPs associated with B cells at 48 hours post-injection. TEM images were performed on two popliteal sentinel lymph nodes for each SApNP construct. FR or E2p SApNPs are indicated by yellow arrows and AV adjuvants are indicated by green arrows.

Fig. S8

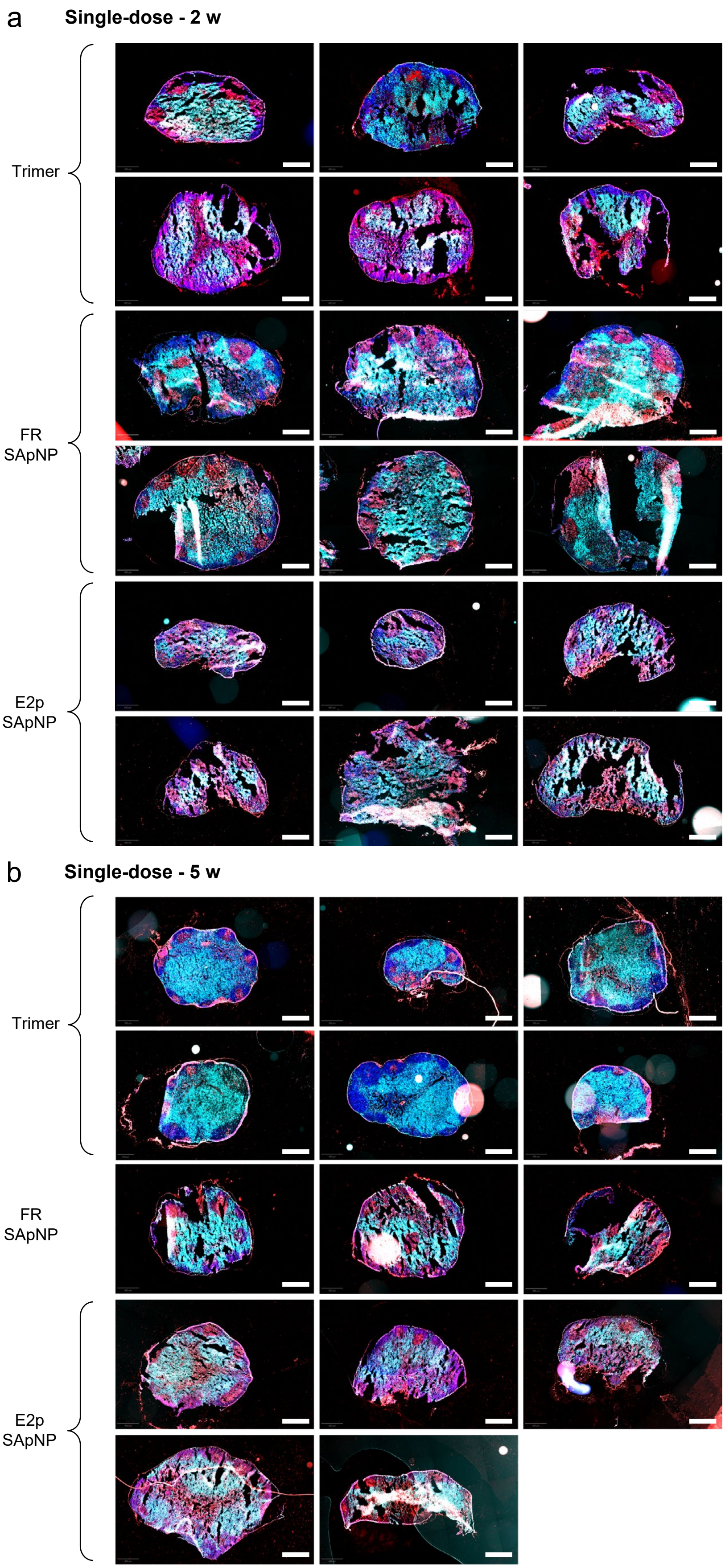
a
Single-dose - 2 w
Trimer
FR
SApNP
E2p
SApNP
b
Single-dose - 5 w
Trimer
FR
SApNP
E2p
SApNP

Fig. S8

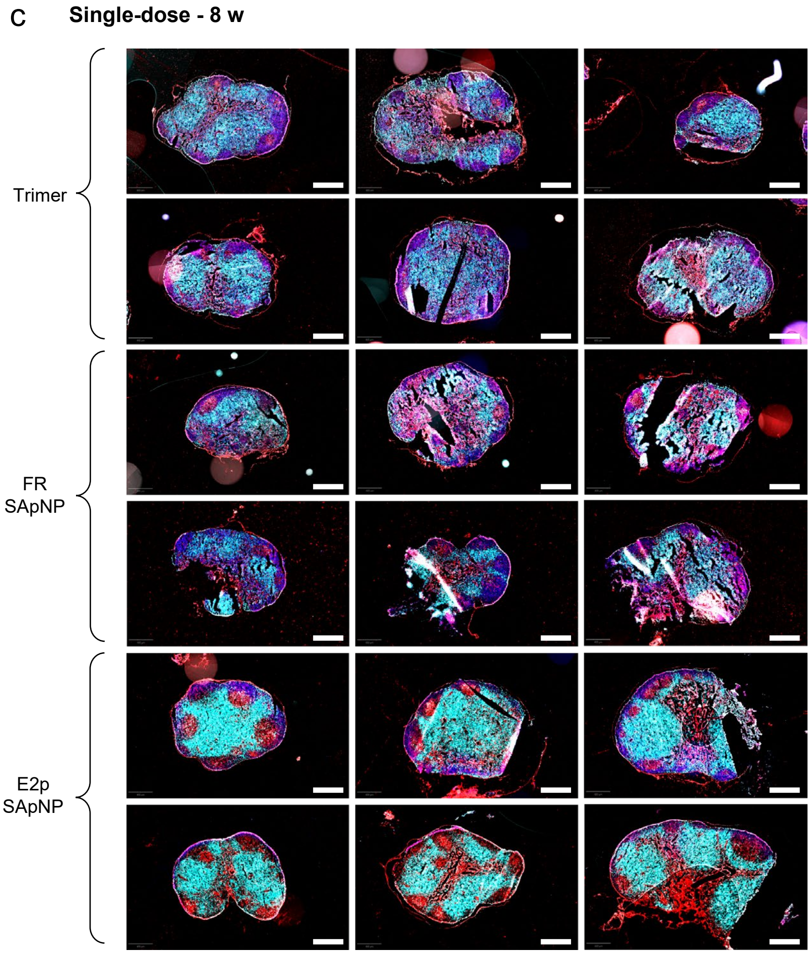

**Fig. S8. Immunohistological analysis of HA trimer and SApNP vaccine-induced germinal centers (GCs).** Immunohistological images of GCs at (**a**) 2, (**b**) 5, and (**c**) 8 weeks following a single-dose injection of HA trimer and HA trimer-presenting FR and E2p SApNPs (10 μg per injection, totaling 40 μg per mouse, n = 5 mice/group). Immunofluorescent images are pseudo-color-coded as follows: BCL6$^+$ (red), CD4$^+$ (cyan), and DAPI (blue). Scale bars: 500 μm (entire lymph node).

Fig. S9

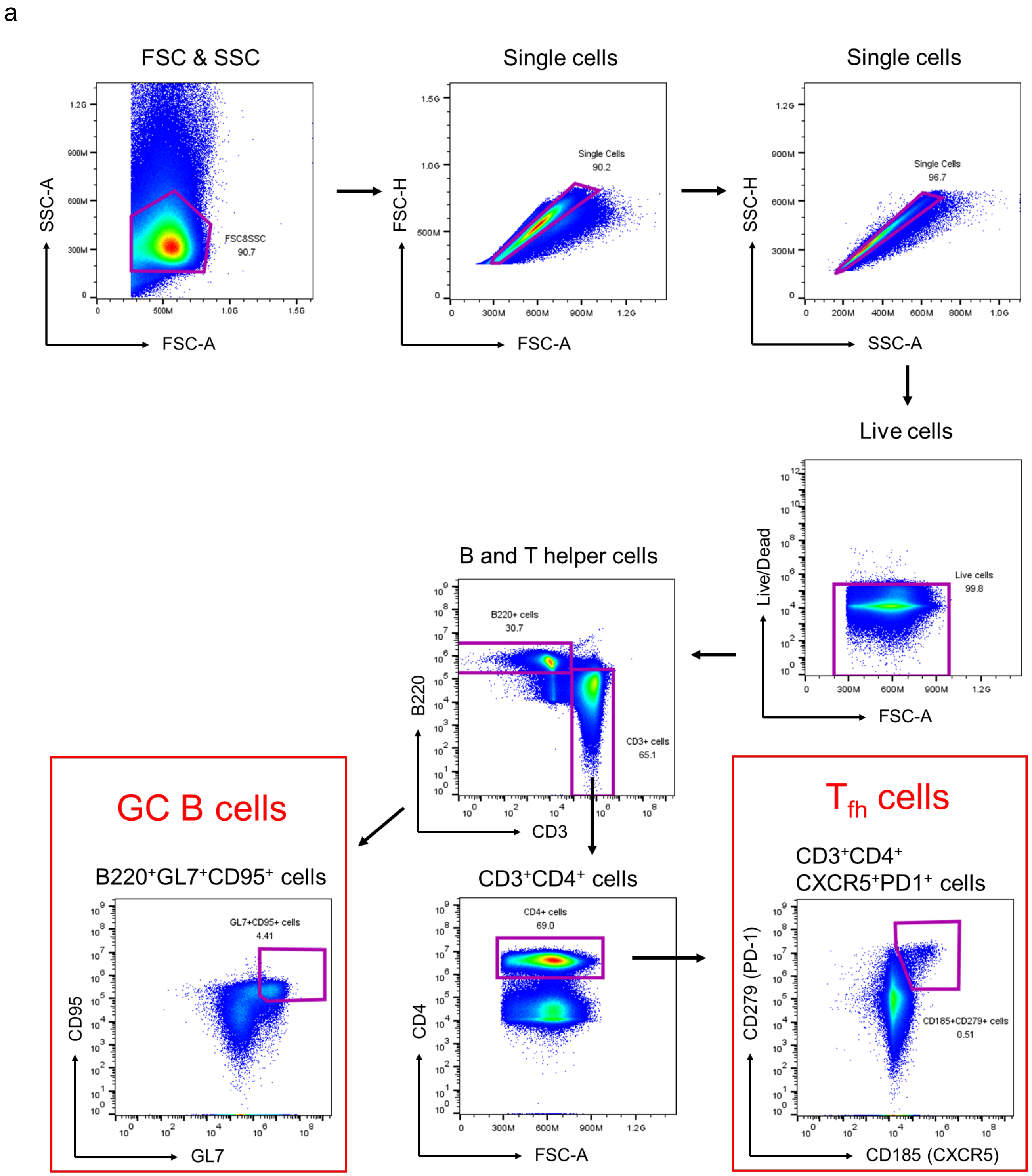

**Fig. S9. Flow cytometry analysis of germinal centers (GCs) induced by HA trimer and SApNPs vaccines.** Gating strategy for analyzing GCs (GC B cells and T follicular helper cells) using flow cytometry (*n* = 5 mice/group).

**Fig. S10**

a **Sera of individual mice immunized with H1N1 CA09 HA vaccines binding to H1N1 CA09 HA-N95L(1TD0) trimer**

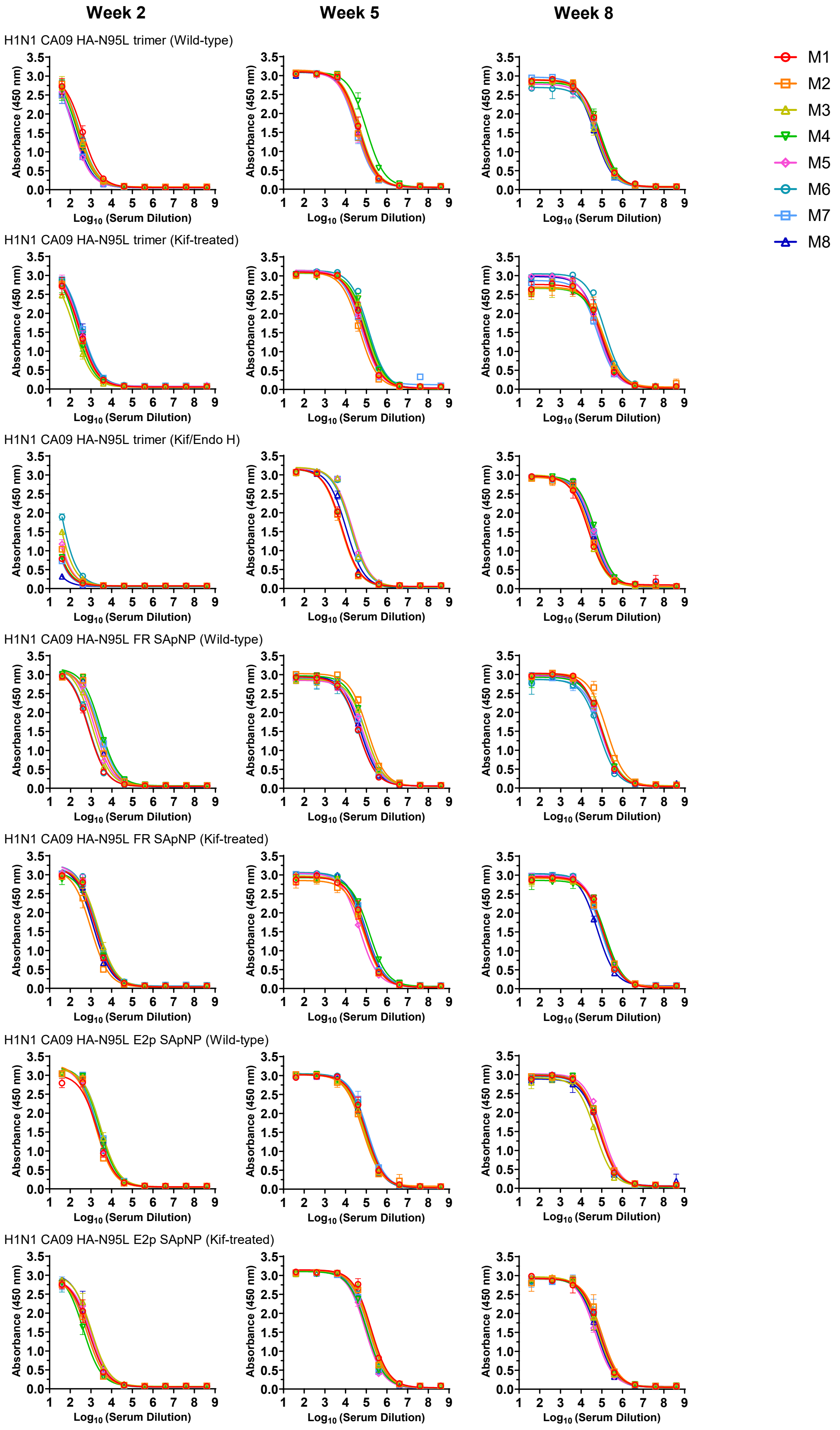

Fig. S10

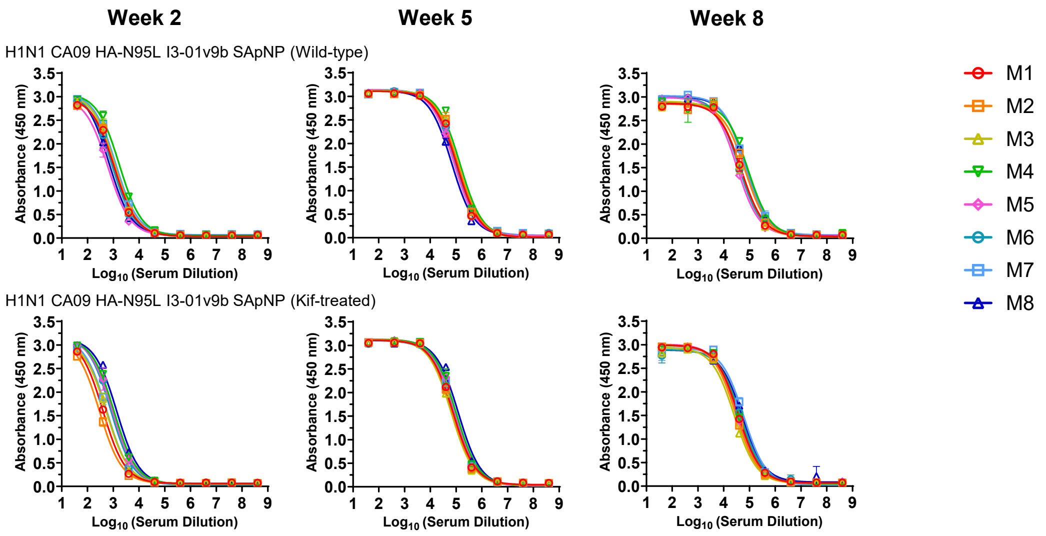

#### b Mouse serum ELISA $EC_{50}$ titers

**Week 2**

| Antigen | $EC_{50}$ titers (week 2) | | | | | | | | Geometric Mean |
|---|---|---|---|---|---|---|---|---|---|
| | M1 | M2 | M3 | M4 | M5 | M6 | M7 | M8 | |
| H1N1 CA09 HA-N95L trimer (Wild-type) | 390 | 208 | 258 | 206 | 120 | 276 | 138 | 162 | 205.4 |
| H1N1 CA09 HA-N95L trimer (Kif-treated) | 287 | 251 | 164 | 201 | 288 | 335 | 367 | 350 | 271.5 |
| H1N1 CA09 HA-N95L trimer (Kif/Endo H) | 13 | 20 | 38 | 14 | 24 | 62 | 12 | 4 | 17.7 |
| H1N1 CA09 HA-N95L FR SApNP (Wild-type) | 726 | 1907 | 1112 | 2733 | 1428 | 768 | 2448 | 1925 | 1469.5 |
| H1N1 CA09 HA-N95L FR SApNP (Kif-treated) | 1643 | 998 | 2228 | 1768 | 1610 | 1828 | 1722 | 1313 | 1600.1 |
| H1N1 CA09 HA-N95L E2p SApNP (Wild-type) | 2103 | 1641 | 2928 | 2548 | 1818 | 2064 | 2939 | 2726 | 2296.6 |
| H1N1 CA09 HA-N95L E2p SApNP (Kif-treated) | 821 | 636 | 1052 | 431 | 996 | 825 | 634 | 1002 | 770.0 |
| H1N1 CA09 HA-N95L I3-01v9b SApNP (Wild-type) | 1073 | 1127 | 1144 | 1699 | 606 | 871 | 1238 | 677 | 1004.9 |
| H1N1 CA09 HA-N95L I3-01v9b SApNP (Kif-treated) | 397 | 292 | 545 | 1068 | 940 | 809 | 560 | 1379 | 667.8 |

**Week 5**

| Antigen | $EC_{50}$ titers (week 5) | | | | | | | | Geometric Mean |
|---|---|---|---|---|---|---|---|---|---|
| | M1 | M2 | M3 | M4 | M5 | M6 | M7 | M8 | |
| H1N1 CA09 HA-N95L trimer (Wild-type) | 45645 | 35970 | 50189 | 103286 | 34990 | 36466 | 31838 | 38555 | 43712 |
| H1N1 CA09 HA-N95L trimer (Kif-treated) | 73661 | 47238 | 93658 | 111711 | 61792 | 126001 | 55362 | 77319 | 76824 |
| H1N1 CA09 HA-N95L trimer (Kif/Endo H) | 6453 | 5930 | 18165 | 6375 | 20312 | 17136 | 17720 | 10463 | 11404 |
| H1N1 CA09 HA-N95L FR SApNP (Wild-type) | 41897 | 112423 | 90740 | 86690 | 71834 | 42244 | 64057 | 52630 | 66426 |
| H1N1 CA09 HA-N95L FR SApNP (Kif-treated) | 84078 | 77322 | 81581 | 136633 | 47734 | 82330 | 89474 | 94645 | 83712 |
| H1N1 CA09 HA-N95L E2p SApNP (Wild-type) | 96412 | 67706 | 80475 | 84699 | 97370 | 101162 | 114953 | 84646 | 89895 |
| H1N1 CA09 HA-N95L E2p SApNP (Kif-treated) | 171683 | 134567 | 151672 | 104855 | 90147 | 133945 | 105077 | 108523 | 122464 |
| H1N1 CA09 HA-N95L I3-01v9b SApNP (Wild-type) | 106202 | 121855 | 115865 | 144854 | 83944 | 94494 | 108025 | 68489 | 103080 |
| H1N1 CA09 HA-N95L I3-01v9b SApNP (Kif-treated) | 76571 | 71428 | 64464 | 99170 | 93902 | 72579 | 88103 | 123175 | 84446 |

**Week 8**

| Antigen | $EC_{50}$ titers (week 8) | | | | | | | | Geometric Mean |
|---|---|---|---|---|---|---|---|---|---|
| | M1 | M2 | M3 | M4 | M5 | M6 | M7 | M8 | |
| H1N1 CA09 HA-N95L trimer (Wild-type) | 70625 | 70623 | 49991 | 86499 | 69245 | 66352 | 49128 | 45388 | 62093 |
| H1N1 CA09 HA-N95L trimer (Kif-treated) | 102368 | 123069 | 134929 | 109653 | 70975 | 142565 | 65092 | 79897 | 99759 |
| H1N1 CA09 HA-N95L trimer (Kif/Endo H) | 22130 | 28369 | 22509 | 49792 | 42211 | 42732 | 39000 | 33632 | 33703 |
| H1N1 CA09 HA-N95L FR SApNP (Wild-type) | 96989 | 167841 | 100539 | 108233 | 90979 | 71819 | 92507 | 93890 | 100066 |
| H1N1 CA09 HA-N95L FR SApNP (Kif-treated) | 117325 | 114201 | 121244 | 143866 | 121910 | 100539 | 88978 | 59516 | 105347 |
| H1N1 CA09 HA-N95L E2p SApNP (Wild-type) | 75377 | 84565 | 48891 | 83403 | 103857 | 77129 | 76227 | 77302 | 76931 |
| H1N1 CA09 HA-N95L E2p SApNP (Kif-treated) | 79581 | 97753 | 65644 | 67552 | 46472 | 77172 | 90870 | 59158 | 71262 |
| H1N1 CA09 HA-N95L I3-01v9b SApNP (Wild-type) | 47249 | 66833 | 46086 | 86836 | 31707 | 37899 | 66004 | 68145 | 53617 |
| H1N1 CA09 HA-N95L I3-01v9b SApNP (Kif-treated) | 36097 | 29493 | 25730 | 38145 | 33831 | 45746 | 59853 | 51928 | 38695 |

#### Statistical analysis

| One-way ANOVA with Tukey's multiple comparisons test (w2) | Statistics | Adjusted P Value |
|---|---|---|
| HA trimer-Wild-type vs. HA trimer-Kif-treated | ns | >0.9999 |
| HA trimer-Wild-type vs. HA trimer-Kif/Endo H | ns | 0.9793 |
| HA trimer-Wild-type vs. HA FR SApNP-Wild-type | **** | <0.0001 |
| HA trimer-Wild-type vs. HA FR SApNP-Kif-treated | **** | <0.0001 |
| HA trimer-Wild-type vs. HA E2p SApNP-Wild-type | **** | <0.0001 |
| HA trimer-Wild-type vs. HA E2p SApNP-Kif-treated | ns | 0.0675 |
| HA trimer-Wild-type vs. HA I3-01v9b SApNP-Wild-type | ** | 0.0011 |
| HA trimer-Wild-type vs. HA I3-01v9b SApNP-Kif-treated | ns | 0.1291 |
| HA trimer-Kif-treated vs. HA trimer-Kif/Endo H | ns | 0.9039 |
| HA trimer-Kif-treated vs. HA FR SApNP-Wild-type | **** | <0.0001 |
| HA trimer-Kif-treated vs. HA FR SApNP-Kif-treated | **** | <0.0001 |
| HA trimer-Kif-treated vs. HA E2p SApNP-Wild-type | **** | <0.0001 |
| HA trimer-Kif-treated vs. HA E2p SApNP-Kif-treated | ns | 0.1451 |
| HA trimer-Kif-treated vs. HA I3-01v9b SApNP-Wild-type | ** | 0.0033 |
| HA trimer-Kif-treated vs. HA I3-01v9b SApNP-Kif-treated | ns | 0.2526 |
| HA trimer-Kif/Endo H vs. HA FR SApNP-Wild-type | **** | <0.0001 |
| HA trimer-Kif/Endo H vs. HA FR SApNP-Kif-treated | **** | <0.0001 |
| HA trimer-Kif/Endo H vs. HA E2p SApNP-Wild-type | **** | <0.0001 |
| HA trimer-Kif/Endo H vs. HA E2p SApNP-Kif-treated | ** | 0.0031 |
| HA trimer-Kif/Endo H vs. HA I3-01v9b SApNP-Wild-type | **** | <0.0001 |
| HA trimer-Kif/Endo H vs. HA I3-01v9b SApNP-Kif-treated | ** | 0.0075 |
| HA FR SApNP-Wild-type vs. HA FR SApNP-Kif-treated | ns | >0.9999 |
| HA FR SApNP-Wild-type vs. HA E2p SApNP-Wild-type | ** | 0.0089 |
| HA FR SApNP-Wild-type vs. HA E2p SApNP-Kif-treated | ** | 0.0012 |
| HA FR SApNP-Wild-type vs. HA I3-01v9b SApNP-Wild-type | ns | 0.0708 |
| HA FR SApNP-Wild-type vs. HA I3-01v9b SApNP-Kif-treated | *** | 0.0005 |
| HA FR SApNP-Kif-treated vs. HA E2p SApNP-Wild-type | * | 0.0101 |
| HA FR SApNP-Kif-treated vs. HA E2p SApNP-Kif-treated | ** | 0.001 |
| HA FR SApNP-Kif-treated vs. HA I3-01v9b SApNP-Wild-type | ns | 0.0637 |
| HA FR SApNP-Kif-treated vs. HA I3-01v9b SApNP-Kif-treated | *** | 0.0004 |
| HA E2p SApNP-Wild-type vs. HA E2p SApNP-Kif-treated | **** | <0.0001 |
| HA E2p SApNP-Wild-type vs. HA I3-01v9b SApNP-Wild-type | **** | <0.0001 |
| HA E2p SApNP-Wild-type vs. HA I3-01v9b SApNP-Kif-treated | **** | <0.0001 |
| HA E2p SApNP-Kif-treated vs. HA I3-01v9b SApNP-Wild-type | ns | 0.9084 |
| HA E2p SApNP-Kif-treated vs. HA I3-01v9b SApNP-Kif-treated | ns | >0.9999 |
| HA I3-01v9b SApNP-Wild-type vs. HA I3-01v9b SApNP-Kif-treated | ns | 0.7838 |

| One-way ANOVA with Tukey's multiple comparisons test (w5) | Statistics | Adjusted P Value |
|---|---|---|
| HA trimer-Wild-type vs. HA trimer-Kif-treated | ns | 0.0811 |
| HA trimer-Wild-type vs. HA trimer-Kif/Endo H | ns | 0.0714 |
| HA trimer-Wild-type vs. HA FR SApNP-Wild-type | ns | 0.497 |
| HA trimer-Wild-type vs. HA FR SApNP-Kif-treated | * | 0.0198 |
| HA trimer-Wild-type vs. HA E2p SApNP-Wild-type | ** | 0.0063 |
| HA trimer-Wild-type vs. HA E2p SApNP-Kif-treated | **** | <0.0001 |
| HA trimer-Wild-type vs. HA I3-01v9b SApNP-Wild-type | **** | <0.0001 |
| HA trimer-Wild-type vs. HA I3-01v9b SApNP-Kif-treated | * | 0.0228 |
| HA trimer-Kif-treated vs. HA trimer-Kif/Endo H | **** | <0.0001 |
| HA trimer-Kif-treated vs. HA FR SApNP-Wild-type | ns | 0.9894 |
| HA trimer-Kif-treated vs. HA FR SApNP-Kif-treated | ns | 0.9998 |
| HA trimer-Kif-treated vs. HA E2p SApNP-Wild-type | ns | 0.992 |
| HA trimer-Kif-treated vs. HA E2p SApNP-Kif-treated | ** | 0.0056 |
| HA trimer-Kif-treated vs. HA I3-01v9b SApNP-Wild-type | ns | 0.4147 |
| HA trimer-Kif-treated vs. HA I3-01v9b SApNP-Kif-treated | ns | >0.9999 |
| HA trimer-Kif/Endo H vs. HA FR SApNP-Wild-type | **** | <0.0001 |
| HA trimer-Kif/Endo H vs. HA FR SApNP-Kif-treated | **** | <0.0001 |
| HA trimer-Kif/Endo H vs. HA E2p SApNP-Wild-type | **** | <0.0001 |
| HA trimer-Kif/Endo H vs. HA E2p SApNP-Kif-treated | **** | <0.0001 |
| HA trimer-Kif/Endo H vs. HA I3-01v9b SApNP-Wild-type | **** | <0.0001 |
| HA trimer-Kif/Endo H vs. HA I3-01v9b SApNP-Kif-treated | **** | <0.0001 |
| HA FR SApNP-Wild-type vs. HA FR SApNP-Kif-treated | ns | 0.8648 |
| HA FR SApNP-Wild-type vs. HA E2p SApNP-Wild-type | ns | 0.6516 |
| HA FR SApNP-Wild-type vs. HA E2p SApNP-Kif-treated | *** | 0.0002 |
| HA FR SApNP-Wild-type vs. HA I3-01v9b SApNP-Wild-type | ns | 0.0589 |
| HA FR SApNP-Wild-type vs. HA I3-01v9b SApNP-Kif-treated | ns | 0.8855 |
| HA FR SApNP-Kif-treated vs. HA E2p SApNP-Wild-type | ns | >0.9999 |
| HA FR SApNP-Kif-treated vs. HA E2p SApNP-Kif-treated | * | 0.0273 |
| HA FR SApNP-Kif-treated vs. HA I3-01v9b SApNP-Wild-type | ns | 0.7567 |
| HA FR SApNP-Kif-treated vs. HA I3-01v9b SApNP-Kif-treated | ns | >0.9999 |
| HA E2p SApNP-Wild-type vs. HA E2p SApNP-Kif-treated | ns | 0.0741 |
| HA E2p SApNP-Wild-type vs. HA I3-01v9b SApNP-Wild-type | ns | 0.9269 |
| HA E2p SApNP-Wild-type vs. HA I3-01v9b SApNP-Kif-treated | ns | >0.9999 |
| HA E2p SApNP-Kif-treated vs. HA I3-01v9b SApNP-Wild-type | ns | 0.7103 |
| HA E2p SApNP-Kif-treated vs. HA I3-01v9b SApNP-Kif-treated | * | 0.0238 |
| HA I3-01v9b SApNP-Wild-type vs. HA I3-01v9b SApNP-Kif-treated | ns | 0.727 |

| One-way ANOVA with Tukey's multiple comparisons test (w8) | Statistics | Adjusted P Value |
|---|---|---|
| HA trimer-Wild-type vs. HA trimer-Kif-treated | ** | 0.0048 |
| HA trimer-Wild-type vs. HA trimer-Kif/Endo H | ns | 0.123 |
| HA trimer-Wild-type vs. HA FR SApNP-Wild-type | ** | 0.0061 |
| HA trimer-Wild-type vs. HA FR SApNP-Kif-treated | *** | 0.001 |
| HA trimer-Wild-type vs. HA E2p SApNP-Wild-type | ns | 0.8574 |
| HA trimer-Wild-type vs. HA E2p SApNP-Kif-treated | ns | 0.9885 |
| HA trimer-Wild-type vs. HA I3-01v9b SApNP-Wild-type | ns | 0.9984 |
| HA trimer-Wild-type vs. HA I3-01v9b SApNP-Kif-treated | ns | 0.3356 |
| HA trimer-Kif-treated vs. HA trimer-Kif/Endo H | **** | <0.0001 |
| HA trimer-Kif-treated vs. HA FR SApNP-Wild-type | ns | >0.9999 |
| HA trimer-Kif-treated vs. HA FR SApNP-Kif-treated | ns | >0.9999 |
| HA trimer-Kif-treated vs. HA E2p SApNP-Wild-type | ns | 0.2409 |
| HA trimer-Kif-treated vs. HA E2p SApNP-Kif-treated | ns | 0.0745 |
| HA trimer-Kif-treated vs. HA I3-01v9b SApNP-Wild-type | *** | 0.0004 |
| HA trimer-Kif-treated vs. HA I3-01v9b SApNP-Kif-treated | **** | <0.0001 |
| HA trimer-Kif/Endo H vs. HA FR SApNP-Wild-type | **** | <0.0001 |
| HA trimer-Kif/Endo H vs. HA FR SApNP-Kif-treated | **** | <0.0001 |
| HA trimer-Kif/Endo H vs. HA E2p SApNP-Wild-type | ** | 0.0017 |
| HA trimer-Kif/Endo H vs. HA E2p SApNP-Kif-treated | ** | 0.0093 |
| HA trimer-Kif/Endo H vs. HA I3-01v9b SApNP-Wild-type | ns | 0.4626 |
| HA trimer-Kif/Endo H vs. HA I3-01v9b SApNP-Kif-treated | ns | 0.9999 |
| HA FR SApNP-Wild-type vs. HA FR SApNP-Kif-treated | ns | 0.9997 |
| HA FR SApNP-Wild-type vs. HA E2p SApNP-Wild-type | ns | 0.2754 |
| HA FR SApNP-Wild-type vs. HA E2p SApNP-Kif-treated | ns | 0.0888 |
| HA FR SApNP-Wild-type vs. HA I3-01v9b SApNP-Wild-type | *** | 0.0006 |
| HA FR SApNP-Wild-type vs. HA I3-01v9b SApNP-Kif-treated | **** | <0.0001 |
| HA FR SApNP-Kif-treated vs. HA E2p SApNP-Wild-type | ns | 0.083 |
| HA FR SApNP-Kif-treated vs. HA E2p SApNP-Kif-treated | * | 0.02 |
| HA FR SApNP-Kif-treated vs. HA I3-01v9b SApNP-Wild-type | **** | <0.0001 |
| HA FR SApNP-Kif-treated vs. HA I3-01v9b SApNP-Kif-treated | **** | <0.0001 |
| HA E2p SApNP-Wild-type vs. HA E2p SApNP-Kif-treated | ns | 0.9998 |
| HA E2p SApNP-Wild-type vs. HA I3-01v9b SApNP-Wild-type | ns | 0.4178 |
| HA E2p SApNP-Wild-type vs. HA I3-01v9b SApNP-Kif-treated | ** | 0.0086 |
| HA E2p SApNP-Kif-treated vs. HA I3-01v9b SApNP-Wild-type | ns | 0.7626 |
| HA E2p SApNP-Kif-treated vs. HA I3-01v9b SApNP-Kif-treated | * | 0.0402 |
| HA I3-01v9b SApNP-Wild-type vs. HA I3-01v9b SApNP-Kif-treated | ns | 0.7876 |

### Fig. S10

**C** **Week-8 sera of individual mice immunized with H1N1 CA09 HA vaccines binding to H3N2 HK68 HA-N95L(1TD0) trimer**

**Week 8**

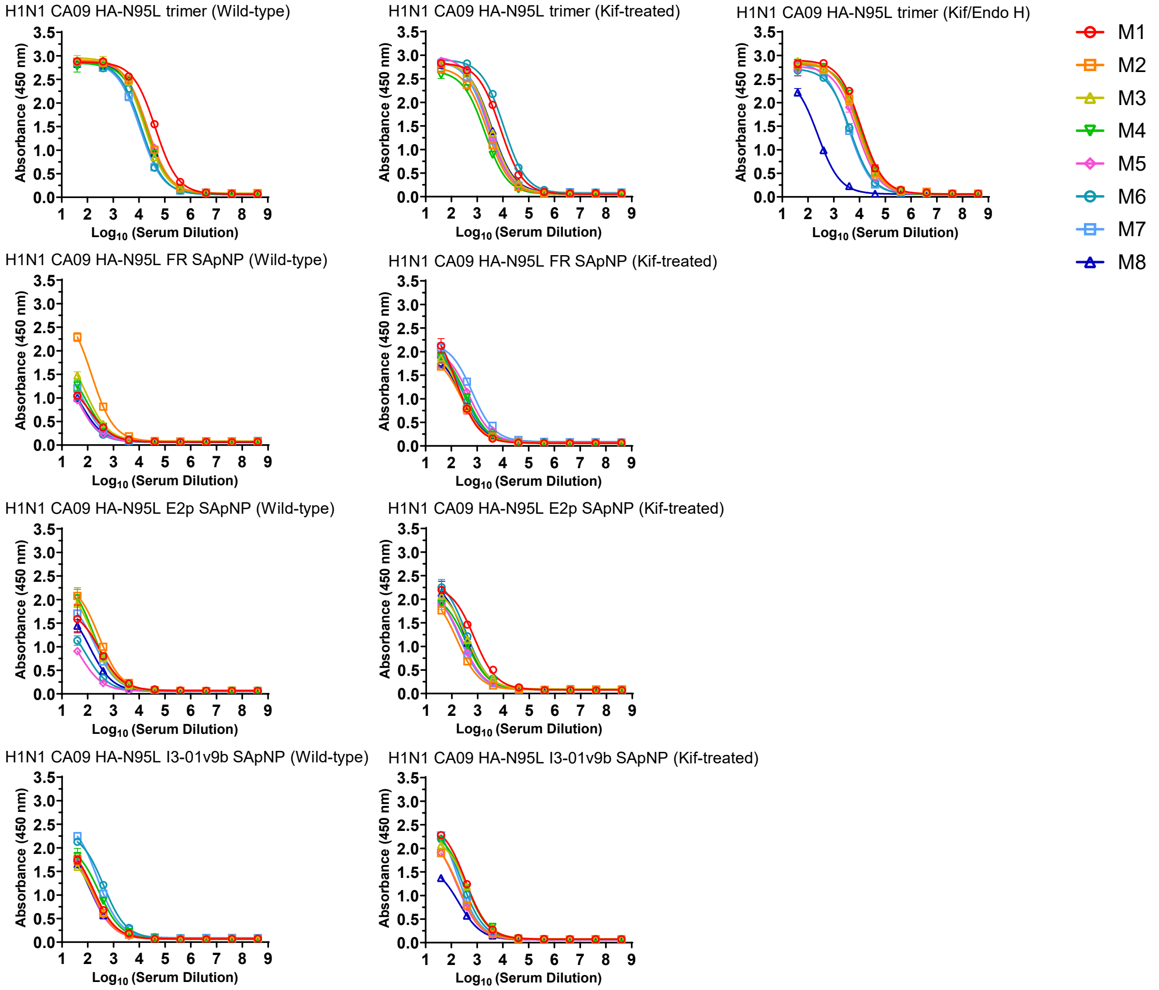

#### Mouse serum ELISA $EC_{50}$ titers

**Week 8**

| Antigen | $EC_{50}$ titers (week 8) | | | | | | | | Geometric Mean |
|---|---|---|---|---|---|---|---|---|---|
| | M1 | M2 | M3 | M4 | M5 | M6 | M7 | M8 | |
| H1N1 CA09 HA-N95L trimer (Wild-type) | 43377 | 20439 | 15951 | 19657 | 21204 | 12549 | 10832 | 19918 | 18853 |
| H1N1 CA09 HA-N95L trimer (Kif-treated) | 7962 | 2486 | 3462 | 1983 | 2762 | 10675 | 2645 | 3646 | 3744 |
| H1N1 CA09 HA-N95L trimer (Kif/Endo H) | 11564 | 9506 | 8612 | 11216 | 7875 | 4482 | 3750 | 239 | 4906 |
| H1N1 CA09 HA-N95L FR SApNP (Wild-type) | 127 | 141 | 89 | 88 | 41 | 28 | 57 | 55 | 69 |
| H1N1 CA09 HA-N95L FR SApNP (Kif-treated) | 157 | 256 | 225 | 362 | 474 | 299 | 647 | 259 | 306 |
| H1N1 CA09 HA-N95L E2p SApNP (Wild-type) | 300 | 275 | 175 | 169 | 51 | 88 | 184 | 125 | 150 |
| H1N1 CA09 HA-N95L E2p SApNP (Kif-treated) | 739 | 162 | 376 | 345 | 242 | 364 | 205 | 318 | 312 |
| H1N1 CA09 HA-N95L I3-01v9b SApNP (Wild-type) | 170 | 135 | 171 | 271 | 126 | 413 | 241 | 124 | 189 |
| H1N1 CA09 HA-N95L I3-01v9b SApNP (Kif-treated) | 364 | 194 | 428 | 370 | 176 | 259 | 173 | 195 | 254 |

#### Statistical analysis

| One-way ANOVA with Tukey's multiple comparisons test (w8) | Statistics | Adjusted P Value |
|---|---|---|
| HA trimer-Wild-type vs. HA trimer-Kif-treated | **** | <0.0001 |
| HA trimer-Wild-type vs. HA trimer-Kif/Endo H | **** | <0.0001 |
| HA trimer-Wild-type vs. HA FR SApNP-Wild-type | **** | <0.0001 |
| HA trimer-Wild-type vs. HA FR SApNP-Kif-treated | **** | <0.0001 |
| HA trimer-Wild-type vs. HA E2p SApNP-Wild-type | **** | <0.0001 |
| HA trimer-Wild-type vs. HA E2p SApNP-Kif-treated | **** | <0.0001 |
| HA trimer-Wild-type vs. HA I3-01v9b SApNP-Wild-type | **** | <0.0001 |
| HA trimer-Wild-type vs. HA I3-01v9b SApNP-Kif-treated | **** | <0.0001 |
| HA trimer-Kif-treated vs. HA trimer-Kif/Endo H | ns | 0.8759 |
| HA trimer-Kif-treated vs. HA FR SApNP-Wild-type | ns | 0.3362 |
| HA trimer-Kif-treated vs. HA FR SApNP-Kif-treated | ns | 0.418 |
| HA trimer-Kif-treated vs. HA E2p SApNP-Wild-type | ns | 0.3647 |
| HA trimer-Kif-treated vs. HA E2p SApNP-Kif-treated | ns | 0.421 |
| HA trimer-Kif-treated vs. HA I3-01v9b SApNP-Wild-type | ns | 0.3759 |
| HA trimer-Kif-treated vs. HA I3-01v9b SApNP-Kif-treated | ns | 0.3964 |
| HA trimer-Kif/Endo H vs. HA FR SApNP-Wild-type | ** | 0.0099 |
| HA trimer-Kif/Endo H vs. HA FR SApNP-Kif-treated | * | 0.0149 |
| HA trimer-Kif/Endo H vs. HA E2p SApNP-Wild-type | * | 0.0115 |
| HA trimer-Kif/Endo H vs. HA E2p SApNP-Kif-treated | * | 0.0151 |
| HA trimer-Kif/Endo H vs. HA I3-01v9b SApNP-Wild-type | * | 0.0121 |
| HA trimer-Kif/Endo H vs. HA I3-01v9b SApNP-Kif-treated | * | 0.0134 |
| HA FR SApNP-Wild-type vs. HA FR SApNP-Kif-treated | ns | >0.9999 |
| HA FR SApNP-Wild-type vs. HA E2p SApNP-Wild-type | ns | >0.9999 |
| HA FR SApNP-Wild-type vs. HA E2p SApNP-Kif-treated | ns | >0.9999 |
| HA FR SApNP-Wild-type vs. HA I3-01v9b SApNP-Wild-type | ns | >0.9999 |
| HA FR SApNP-Wild-type vs. HA I3-01v9b SApNP-Kif-treated | ns | >0.9999 |
| HA FR SApNP-Kif-treated vs. HA E2p SApNP-Wild-type | ns | >0.9999 |
| HA FR SApNP-Kif-treated vs. HA E2p SApNP-Kif-treated | ns | >0.9999 |
| HA FR SApNP-Kif-treated vs. HA I3-01v9b SApNP-Wild-type | ns | >0.9999 |
| HA FR SApNP-Kif-treated vs. HA I3-01v9b SApNP-Kif-treated | ns | >0.9999 |
| HA E2p SApNP-Wild-type vs. HA E2p SApNP-Kif-treated | ns | >0.9999 |
| HA E2p SApNP-Wild-type vs. HA I3-01v9b SApNP-Wild-type | ns | >0.9999 |
| HA E2p SApNP-Wild-type vs. HA I3-01v9b SApNP-Kif-treated | ns | >0.9999 |
| HA E2p SApNP-Kif-treated vs. HA I3-01v9b SApNP-Wild-type | ns | >0.9999 |
| HA E2p SApNP-Kif-treated vs. HA I3-01v9b SApNP-Kif-treated | ns | >0.9999 |
| HA I3-01v9b SApNP-Wild-type vs. HA I3-01v9b SApNP-Kif-treated | ns | >0.9999 |

**Fig. S10**

d

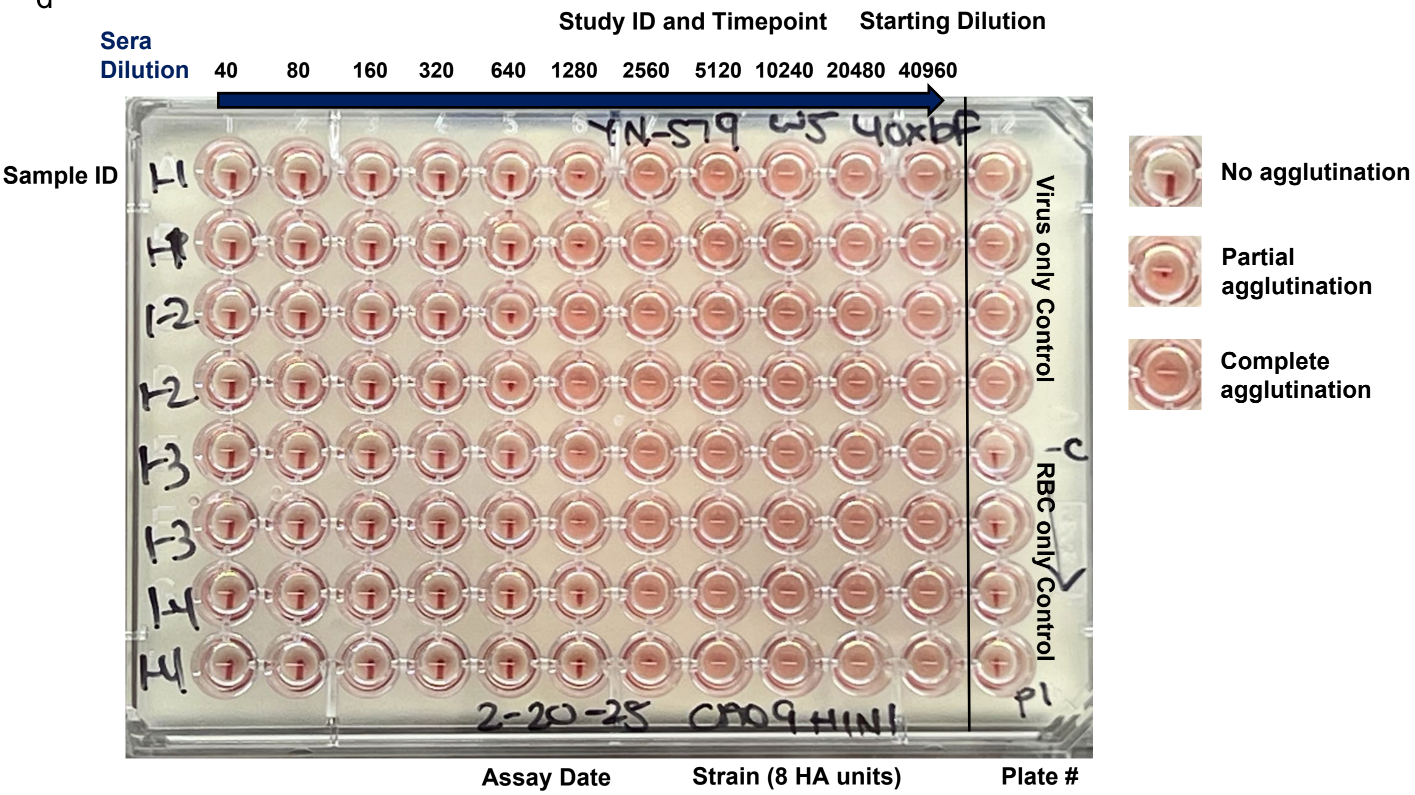

e **H1N1 CA09 HA-induced sera HAI against vaccine matched H1N1 CA09 virus**

**Week 2**

| Antigen | HAI Titer ($Log_2$) (week 2) | | | | | | | |
|---|---|---|---|---|---|---|---|---|
| | M1 | M2 | M3 | M4 | M5 | M6 | M7 | M8 |
| H1N1 CA09 HA-N95L trimer (Wild-type) | 20 | 20 | 20 | 20 | 20 | 20 | 20 | 20 |
| H1N1 CA09 HA-N95L trimer (Wild-type) | 20 | 20 | 20 | 20 | 20 | 20 | 20 | 20 |
| H1N1 CA09 HA-N95L trimer (Kif-treated) | 20 | 20 | 20 | 20 | 20 | 20 | 20 | 20 |
| H1N1 CA09 HA-N95L trimer (Kif-treated) | 20 | 20 | 20 | 20 | 20 | 20 | 20 | 20 |
| H1N1 CA09 HA-N95L trimer (Kif/Endo H) | 20 | 20 | 20 | 20 | 20 | 20 | 20 | 20 |
| H1N1 CA09 HA-N95L trimer (Kif/Endo H) | 20 | 20 | 20 | 20 | 20 | 20 | 20 | 20 |
| H1N1 CA09 HA-N95L FR SApNP (Wild-type) | 20 | 40 | 20 | 80 | 20 | 20 | 40 | 40 |
| H1N1 CA09 HA-N95L FR SApNP (Wild-type) | 20 | 40 | 20 | 80 | 20 | 20 | 40 | 40 |
| H1N1 CA09 HA-N95L FR SApNP (Kif-treated) | 40 | 20 | 40 | 20 | 40 | 20 | 20 | 40 |
| H1N1 CA09 HA-N95L FR SApNP (Kif-treated) | 40 | 20 | 40 | 20 | 40 | 20 | 20 | 40 |
| H1N1 CA09 HA-N95L E2p SApNP (Wild-type) | 20 | 20 | 40 | 80 | 20 | 20 | 40 | 20 |
| H1N1 CA09 HA-N95L E2p SApNP (Wild-type) | 20 | 20 | 40 | 80 | 20 | 20 | 40 | 20 |
| H1N1 CA09 HA-N95L E2p SApNP (Kif-treated) | 40 | 40 | 40 | 40 | 40 | 40 | 40 | 40 |
| H1N1 CA09 HA-N95L E2p SApNP (Kif-treated) | 40 | 40 | 40 | 40 | 40 | 40 | 40 | 40 |
| H1N1 CA09 HA-N95L I3-01v9b SApNP (Wild-type) | 80 | 20 | 20 | 80 | 20 | 40 | 40 | 20 |
| H1N1 CA09 HA-N95L I3-01v9b SApNP (Wild-type) | 80 | 20 | 20 | 80 | 20 | 40 | 40 | 20 |
| H1N1 CA09 HA-N95L I3-01v9b SApNP (Kif-treated) | 20 | 20 | 20 | 40 | 40 | 40 | 20 | 40 |
| H1N1 CA09 HA-N95L I3-01v9b SApNP (Kif-treated) | 20 | 20 | 20 | 40 | 40 | 40 | 20 | 40 |

**Week 5**

| Antigen | HAI Titer ($Log_2$) (week 5) | | | | | | | |
|---|---|---|---|---|---|---|---|---|
| | M1 | M2 | M3 | M4 | M5 | M6 | M7 | M8 |
| H1N1 CA09 HA-N95L trimer (Wild-type) | 640 | 640 | 640 | 1280 | 160 | 320 | 640 | 640 |
| H1N1 CA09 HA-N95L trimer (Wild-type) | 640 | 640 | 640 | 1280 | 160 | 320 | 640 | 640 |
| H1N1 CA09 HA-N95L trimer (Kif-treated) | 640 | 320 | 1280 | 640 | 640 | 640 | 320 | 640 |
| H1N1 CA09 HA-N95L trimer (Kif-treated) | 640 | 320 | 1280 | 640 | 640 | 640 | 320 | 640 |
| H1N1 CA09 HA-N95L trimer (Kif/Endo H) | 80 | 40 | 80 | 40 | 320 | 160 | 80 | 160 |
| H1N1 CA09 HA-N95L trimer (Kif/Endo H) | 80 | 40 | 80 | 40 | 320 | 160 | 80 | 160 |
| H1N1 CA09 HA-N95L FR SApNP (Wild-type) | 1280 | 2560 | 2560 | 1280 | 1280 | 1280 | 1280 | 1280 |
| H1N1 CA09 HA-N95L FR SApNP (Wild-type) | 1280 | 2560 | 2560 | 1280 | 1280 | 1280 | 1280 | 2560 |
| H1N1 CA09 HA-N95L FR SApNP (Kif-treated) | 2560 | 1280 | 1280 | 2560 | 1280 | 1280 | 1280 | 1280 |
| H1N1 CA09 HA-N95L FR SApNP (Kif-treated) | 2560 | 1280 | 1280 | 2560 | 1280 | 1280 | 1280 | 1280 |
| H1N1 CA09 HA-N95L E2p SApNP (Wild-type) | 1280 | 640 | 1280 | 1280 | 640 | 1280 | 1280 | 640 |
| H1N1 CA09 HA-N95L E2p SApNP (Wild-type) | 1280 | 640 | 1280 | 1280 | 640 | 1280 | 1280 | 640 |
| H1N1 CA09 HA-N95L E2p SApNP (Kif-treated) | 2560 | 1280 | 2560 | 1280 | 1280 | 1280 | 1280 | 640 |
| H1N1 CA09 HA-N95L E2p SApNP (Kif-treated) | 2560 | 1280 | 2560 | 1280 | 1280 | 1280 | 1280 | 640 |
| H1N1 CA09 HA-N95L I3-01v9b SApNP (Wild-type) | 2560 | 1280 | 640 | 1280 | 1280 | 1280 | 1280 | 1280 |
| H1N1 CA09 HA-N95L I3-01v9b SApNP (Wild-type) | 2560 | 1280 | 1280 | 2560 | 1280 | 1280 | 1280 | 1280 |
| H1N1 CA09 HA-N95L I3-01v9b SApNP (Kif-treated) | 1280 | 1280 | 640 | 640 | 1280 | 1280 | 1280 | 1280 |
| H1N1 CA09 HA-N95L I3-01v9b SApNP (Kif-treated) | 1280 | 1280 | 640 | 640 | 1280 | 1280 | 1280 | 1280 |

**Week 8**

| Antigen | HAI Titer ($Log_2$) (week 8) | | | | | | | |
|---|---|---|---|---|---|---|---|---|
| | M1 | M2 | M3 | M4 | M5 | M6 | M7 | M8 |
| H1N1 CA09 HA-N95L trimer (Wild-type) | 1280 | 1280 | 640 | 1280 | 640 | 640 | 1280 | 640 |
| H1N1 CA09 HA-N95L trimer (Wild-type) | 1280 | 1280 | 640 | 1280 | 640 | 640 | 1280 | 640 |
| H1N1 CA09 HA-N95L trimer (Kif-treated) | 1280 | 1280 | 1280 | 1280 | 640 | 1280 | 1280 | 640 |
| H1N1 CA09 HA-N95L trimer (Kif-treated) | 1280 | 1280 | 1280 | 1280 | 640 | 1280 | 1280 | 1280 |
| H1N1 CA09 HA-N95L trimer (Kif/Endo H) | 320 | 320 | 160 | 160 | 640 | 640 | 640 | 320 |
| H1N1 CA09 HA-N95L trimer (Kif/Endo H) | 320 | 160 | 160 | 160 | 640 | 640 | 640 | 320 |
| H1N1 CA09 HA-N95L FR SApNP (Wild-type) | 2560 | 2560 | 2560 | 2560 | 1280 | 1280 | 1280 | 2560 |
| H1N1 CA09 HA-N95L FR SApNP (Wild-type) | 2560 | 2560 | 2560 | 2560 | 1280 | 1280 | 1280 | 2560 |
| H1N1 CA09 HA-N95L FR SApNP (Kif-treated) | 2560 | 1280 | 2560 | 2560 | 2560 | 1280 | 1280 | 640 |
| H1N1 CA09 HA-N95L FR SApNP (Kif-treated) | 2560 | 1280 | 2560 | 2560 | 2560 | 1280 | 1280 | 640 |
| H1N1 CA09 HA-N95L E2p SApNP (Wild-type) | 1280 | 640 | 1280 | 1280 | 640 | 1280 | 640 | 1280 |
| H1N1 CA09 HA-N95L E2p SApNP (Wild-type) | 1280 | 640 | 1280 | 1280 | 640 | 1280 | 640 | 1280 |
| H1N1 CA09 HA-N95L E2p SApNP (Kif-treated) | 2560 | 1280 | 2560 | 1280 | 1280 | 1280 | 2560 | 1280 |
| H1N1 CA09 HA-N95L E2p SApNP (Kif-treated) | 1280 | 1280 | 2560 | 1280 | 1280 | 1280 | 2560 | 1280 |
| H1N1 CA09 HA-N95L I3-01v9b SApNP (Wild-type) | 2560 | 1280 | 640 | 1280 | 640 | 1280 | 1280 | 1280 |
| H1N1 CA09 HA-N95L I3-01v9b SApNP (Wild-type) | 2560 | 1280 | 640 | 1280 | 640 | 1280 | 1280 | 1280 |
| H1N1 CA09 HA-N95L I3-01v9b SApNP (Kif-treated) | 1280 | 1280 | 640 | 640 | 640 | 1280 | 1280 | 1280 |
| H1N1 CA09 HA-N95L I3-01v9b SApNP (Kif-treated) | 1280 | 1280 | 640 | 640 | 640 | 1280 | 1280 | 1280 |

### Fig. S10

#### Statistical analysis

| One-way ANOVA with Tukey's multiple comparisons test (w2) | Statistics | Adjusted P Value |
|---|---|---|
| HA trimer-Wild-type vs. HA trimer-Kif-treated | ns | >0.9999 |
| HA trimer-Wild-type vs. HA trimer-Kif/Endo H | ns | >0.9999 |
| HA trimer-Wild-type vs. HA FR SApNP-Wild-type | ns | 0.4659 |
| HA trimer-Wild-type vs. HA FR SApNP-Kif-treated | ns | 0.8871 |
| HA trimer-Wild-type vs. HA E2p SApNP-Wild-type | ns | 0.7003 |
| HA trimer-Wild-type vs. HA E2p SApNP-Kif-treated | ns | 0.126 |
| HA trimer-Wild-type vs. HA I3-01v9b SApNP-Wild-type | ns | 0.126 |
| HA trimer-Wild-type vs. HA I3-01v9b SApNP-Kif-treated | ns | 0.8871 |
| HA trimer-Kif-treated vs. HA trimer-Kif/Endo H | ns | >0.9999 |
| HA trimer-Kif-treated vs. HA FR SApNP-Wild-type | ns | 0.4659 |
| HA trimer-Kif-treated vs. HA FR SApNP-Kif-treated | ns | 0.8871 |
| HA trimer-Kif-treated vs. HA E2p SApNP-Wild-type | ns | 0.7003 |
| HA trimer-Kif-treated vs. HA E2p SApNP-Kif-treated | ns | 0.126 |
| HA trimer-Kif-treated vs. HA I3-01v9b SApNP-Wild-type | ns | 0.126 |
| HA trimer-Kif-treated vs. HA I3-01v9b SApNP-Kif-treated | ns | 0.8871 |
| HA trimer-Kif/Endo H vs. HA FR SApNP-Wild-type | ns | 0.4659 |
| HA trimer-Kif/Endo H vs. HA FR SApNP-Kif-treated | ns | 0.8871 |
| HA trimer-Kif/Endo H vs. HA E2p SApNP-Wild-type | ns | 0.7003 |
| HA trimer-Kif/Endo H vs. HA E2p SApNP-Kif-treated | ns | 0.126 |
| HA trimer-Kif/Endo H vs. HA I3-01v9b SApNP-Wild-type | ns | 0.126 |
| HA trimer-Kif/Endo H vs. HA I3-01v9b SApNP-Kif-treated | ns | 0.8871 |
| HA FR SApNP-Wild-type vs. HA FR SApNP-Kif-treated | ns | 0.9985 |
| HA FR SApNP-Wild-type vs. HA E2p SApNP-Wild-type | ns | >0.9999 |
| HA FR SApNP-Wild-type vs. HA E2p SApNP-Kif-treated | ns | 0.9985 |
| HA FR SApNP-Wild-type vs. HA I3-01v9b SApNP-Wild-type | ns | 0.9985 |
| HA FR SApNP-Wild-type vs. HA I3-01v9b SApNP-Kif-treated | ns | 0.9985 |
| HA FR SApNP-Kif-treated vs. HA E2p SApNP-Wild-type | ns | >0.9999 |
| HA FR SApNP-Kif-treated vs. HA E2p SApNP-Kif-treated | ns | 0.8871 |
| HA FR SApNP-Kif-treated vs. HA I3-01v9b SApNP-Wild-type | ns | 0.8871 |
| HA FR SApNP-Kif-treated vs. HA I3-01v9b SApNP-Kif-treated | ns | >0.9999 |
| HA E2p SApNP-Wild-type vs. HA E2p SApNP-Kif-treated | ns | 0.9774 |
| HA E2p SApNP-Wild-type vs. HA I3-01v9b SApNP-Wild-type | ns | 0.9774 |
| HA E2p SApNP-Wild-type vs. HA I3-01v9b SApNP-Kif-treated | ns | >0.9999 |
| HA E2p SApNP-Kif-treated vs. HA I3-01v9b SApNP-Wild-type | ns | >0.9999 |
| HA E2p SApNP-Kif-treated vs. HA I3-01v9b SApNP-Kif-treated | ns | 0.8871 |
| HA I3-01v9b SApNP-Wild-type vs. HA I3-01v9b SApNP-Kif-treated | ns | 0.8871 |

| One-way ANOVA with Tukey's multiple comparisons test (w5) | Statistics | Adjusted P Value |
|---|---|---|
| HA trimer-Wild-type vs. HA trimer-Kif-treated | ns | >0.9999 |
| HA trimer-Wild-type vs. HA trimer-Kif/Endo H | ns | 0.4024 |
| HA trimer-Wild-type vs. HA FR SApNP-Wild-type | *** | 0.0004 |
| HA trimer-Wild-type vs. HA FR SApNP-Kif-treated | ** | 0.0015 |
| HA trimer-Wild-type vs. HA E2p SApNP-Wild-type | ns | 0.6364 |
| HA trimer-Wild-type vs. HA E2p SApNP-Kif-treated | ** | 0.0049 |
| HA trimer-Wild-type vs. HA I3-01v9b SApNP-Wild-type | ** | 0.0086 |
| HA trimer-Wild-type vs. HA I3-01v9b SApNP-Kif-treated | ns | 0.4024 |
| HA trimer-Kif-treated vs. HA trimer-Kif/Endo H | ns | 0.3493 |
| HA trimer-Kif-treated vs. HA FR SApNP-Wild-type | *** | 0.0006 |
| HA trimer-Kif-treated vs. HA FR SApNP-Kif-treated | ** | 0.002 |
| HA trimer-Kif-treated vs. HA E2p SApNP-Wild-type | ns | 0.6943 |
| HA trimer-Kif-treated vs. HA E2p SApNP-Kif-treated | ** | 0.0065 |
| HA trimer-Kif-treated vs. HA I3-01v9b SApNP-Wild-type | * | 0.0112 |
| HA trimer-Kif-treated vs. HA I3-01v9b SApNP-Kif-treated | ns | 0.4587 |
| HA trimer-Kif/Endo H vs. HA FR SApNP-Wild-type | **** | <0.0001 |
| HA trimer-Kif/Endo H vs. HA FR SApNP-Kif-treated | **** | <0.0001 |
| HA trimer-Kif/Endo H vs. HA E2p SApNP-Wild-type | ** | 0.0037 |
| HA trimer-Kif/Endo H vs. HA E2p SApNP-Kif-treated | **** | <0.0001 |
| HA trimer-Kif/Endo H vs. HA I3-01v9b SApNP-Wild-type | **** | <0.0001 |
| HA trimer-Kif/Endo H vs. HA I3-01v9b SApNP-Kif-treated | ** | 0.0011 |
| HA FR SApNP-Wild-type vs. HA FR SApNP-Kif-treated | ns | >0.9999 |
| HA FR SApNP-Wild-type vs. HA E2p SApNP-Wild-type | ns | 0.1218 |
| HA FR SApNP-Wild-type vs. HA E2p SApNP-Kif-treated | ns | 0.9984 |
| HA FR SApNP-Wild-type vs. HA I3-01v9b SApNP-Wild-type | ns | 0.9927 |
| HA FR SApNP-Wild-type vs. HA I3-01v9b SApNP-Kif-treated | ns | 0.2553 |
| HA FR SApNP-Kif-treated vs. HA E2p SApNP-Wild-type | ns | 0.2553 |
| HA FR SApNP-Kif-treated vs. HA E2p SApNP-Kif-treated | ns | >0.9999 |
| HA FR SApNP-Kif-treated vs. HA I3-01v9b SApNP-Wild-type | ns | 0.9998 |
| HA FR SApNP-Kif-treated vs. HA I3-01v9b SApNP-Kif-treated | ns | 0.4587 |
| HA E2p SApNP-Wild-type vs. HA E2p SApNP-Kif-treated | ns | 0.4587 |
| HA E2p SApNP-Wild-type vs. HA I3-01v9b SApNP-Wild-type | ns | 0.5769 |
| HA E2p SApNP-Wild-type vs. HA I3-01v9b SApNP-Kif-treated | ns | >0.9999 |
| HA E2p SApNP-Kif-treated vs. HA I3-01v9b SApNP-Wild-type | ns | >0.9999 |
| HA E2p SApNP-Kif-treated vs. HA I3-01v9b SApNP-Kif-treated | ns | 0.6943 |
| HA I3-01v9b SApNP-Wild-type vs. HA I3-01v9b SApNP-Kif-treated | ns | 0.7998 |

| One-way ANOVA with Tukey's multiple comparisons test (w8) | Statistics | Adjusted P Value |
|---|---|---|
| HA trimer-Wild-type vs. HA trimer-Kif-treated | ns | 0.9963 |
| HA trimer-Wild-type vs. HA trimer-Kif/Endo H | ns | 0.3574 |
| HA trimer-Wild-type vs. HA FR SApNP-Wild-type | *** | 0.0009 |
| HA trimer-Wild-type vs. HA FR SApNP-Kif-treated | * | 0.0196 |
| HA trimer-Wild-type vs. HA E2p SApNP-Wild-type | ns | >0.9999 |
| HA trimer-Wild-type vs. HA E2p SApNP-Kif-treated | ns | 0.1068 |
| HA trimer-Wild-type vs. HA I3-01v9b SApNP-Wild-type | ns | 0.9302 |
| HA trimer-Wild-type vs. HA I3-01v9b SApNP-Kif-treated | ns | >0.9999 |
| HA trimer-Kif-treated vs. HA trimer-Kif/Endo H | ns | 0.0654 |
| HA trimer-Kif-treated vs. HA FR SApNP-Wild-type | * | 0.0121 |
| HA trimer-Kif-treated vs. HA FR SApNP-Kif-treated | ns | 0.1535 |
| HA trimer-Kif-treated vs. HA E2p SApNP-Wild-type | ns | >0.9999 |
| HA trimer-Kif-treated vs. HA E2p SApNP-Kif-treated | ns | 0.4834 |
| HA trimer-Kif-treated vs. HA I3-01v9b SApNP-Wild-type | ns | >0.9999 |
| HA trimer-Kif-treated vs. HA I3-01v9b SApNP-Kif-treated | ns | >0.9999 |
| HA trimer-Kif/Endo H vs. HA FR SApNP-Wild-type | **** | <0.0001 |
| HA trimer-Kif/Endo H vs. HA FR SApNP-Kif-treated | **** | <0.0001 |
| HA trimer-Kif/Endo H vs. HA E2p SApNP-Wild-type | ns | 0.1978 |
| HA trimer-Kif/Endo H vs. HA E2p SApNP-Kif-treated | **** | <0.0001 |
| HA trimer-Kif/Endo H vs. HA I3-01v9b SApNP-Wild-type | * | 0.0174 |
| HA trimer-Kif/Endo H vs. HA I3-01v9b SApNP-Kif-treated | ns | 0.1978 |
| HA FR SApNP-Wild-type vs. HA FR SApNP-Kif-treated | ns | 0.9874 |
| HA FR SApNP-Wild-type vs. HA E2p SApNP-Wild-type | ** | 0.0026 |
| HA FR SApNP-Wild-type vs. HA E2p SApNP-Kif-treated | ns | 0.7933 |
| HA FR SApNP-Wild-type vs. HA I3-01v9b SApNP-Wild-type | * | 0.0478 |
| HA FR SApNP-Wild-type vs. HA I3-01v9b SApNP-Kif-treated | ** | 0.0026 |
| HA FR SApNP-Kif-treated vs. HA E2p SApNP-Wild-type | * | 0.0478 |
| HA FR SApNP-Kif-treated vs. HA E2p SApNP-Kif-treated | ns | 0.9992 |
| HA FR SApNP-Kif-treated vs. HA I3-01v9b SApNP-Wild-type | ns | 0.3813 |
| HA FR SApNP-Kif-treated vs. HA I3-01v9b SApNP-Kif-treated | * | 0.0478 |
| HA E2p SApNP-Wild-type vs. HA E2p SApNP-Kif-treated | ns | 0.2145 |
| HA E2p SApNP-Wild-type vs. HA I3-01v9b SApNP-Wild-type | ns | 0.9874 |
| HA E2p SApNP-Wild-type vs. HA I3-01v9b SApNP-Kif-treated | ns | >0.9999 |
| HA E2p SApNP-Kif-treated vs. HA I3-01v9b SApNP-Wild-type | ns | 0.7933 |
| HA E2p SApNP-Kif-treated vs. HA I3-01v9b SApNP-Kif-treated | ns | 0.2145 |
| HA I3-01v9b SApNP-Wild-type vs. HA I3-01v9b SApNP-Kif-treated | ns | 0.9874 |

#### f H1N1 CA09 HA-induced sera HAI (at week 8) against non-matched H1N1 PR8, NC99, MI15, and WI22 viruses

### A/PR/8/1934

| Antigen | HAI Titer ($Log_2$) (week 8) | | | | | | | |
|---|---|---|---|---|---|---|---|---|
| | M1 | M2 | M3 | M4 | M5 | M6 | M7 | M8 |
| H1N1 CA09 HA-N95L trimer (Wild-type) | 20 | 20 | 20 | 20 | 20 | 20 | 20 | 20 |
| H1N1 CA09 HA-N95L trimer (Wild-type) | 20 | 20 | 20 | 20 | 20 | 20 | 20 | 20 |
| H1N1 CA09 HA-N95L trimer (Kif-treated) | 20 | 20 | 20 | 20 | 20 | 20 | 20 | 20 |
| H1N1 CA09 HA-N95L trimer (Kif-treated) | 20 | 20 | 20 | 20 | 20 | 20 | 20 | 20 |
| H1N1 CA09 HA-N95L trimer (Kif/Endo H) | 20 | 20 | 20 | 20 | 20 | 20 | 20 | 20 |
| H1N1 CA09 HA-N95L trimer (Kif/Endo H) | 20 | 20 | 20 | 20 | 20 | 20 | 20 | 20 |
| H1N1 CA09 HA-N95L FR SApNP (Wild-type) | 20 | 20 | 20 | 20 | 20 | 20 | 20 | 20 |
| H1N1 CA09 HA-N95L FR SApNP (Wild-type) | 20 | 20 | 20 | 20 | 20 | 20 | 20 | 20 |
| H1N1 CA09 HA-N95L FR SApNP (Kif-treated) | 20 | 20 | 20 | 20 | 20 | 20 | 20 | 20 |
| H1N1 CA09 HA-N95L FR SApNP (Kif-treated) | 20 | 20 | 20 | 20 | 20 | 20 | 20 | 20 |
| H1N1 CA09 HA-N95L E2p SApNP (Wild-type) | 20 | 20 | 20 | 20 | 20 | 20 | 20 | 20 |
| H1N1 CA09 HA-N95L E2p SApNP (Wild-type) | 20 | 20 | 20 | 20 | 20 | 20 | 20 | 20 |
| H1N1 CA09 HA-N95L E2p SApNP (Kif-treated) | 20 | 20 | 20 | 20 | 20 | 20 | 20 | 20 |
| H1N1 CA09 HA-N95L E2p SApNP (Kif-treated) | 20 | 20 | 20 | 20 | 20 | 20 | 20 | 20 |
| H1N1 CA09 HA-N95L I3-01v9b SApNP (Wild-type) | 20 | 20 | 20 | 20 | 20 | 20 | 20 | 20 |
| H1N1 CA09 HA-N95L I3-01v9b SApNP (Wild-type) | 20 | 20 | 20 | 20 | 20 | 20 | 20 | 20 |
| H1N1 CA09 HA-N95L I3-01v9b SApNP (Kif-treated) | 20 | 20 | 20 | 20 | 20 | 20 | 20 | 20 |
| H1N1 CA09 HA-N95L I3-01v9b SApNP (Kif-treated) | 20 | 20 | 20 | 20 | 20 | 20 | 20 | 20 |

### A/NC/20/1999

| Antigen | HAI Titer ($Log_2$) (week 8) | | | | | | | |
|---|---|---|---|---|---|---|---|---|
| | M1 | M2 | M3 | M4 | M5 | M6 | M7 | M8 |
| H1N1 CA09 HA-N95L trimer (Wild-type) | 20 | 20 | 20 | 20 | 20 | 20 | 20 | 20 |
| H1N1 CA09 HA-N95L trimer (Wild-type) | 20 | 20 | 20 | 20 | 20 | 20 | 20 | 20 |
| H1N1 CA09 HA-N95L trimer (Kif-treated) | 20 | 20 | 20 | 20 | 20 | 20 | 20 | 20 |
| H1N1 CA09 HA-N95L trimer (Kif-treated) | 20 | 20 | 20 | 20 | 20 | 20 | 20 | 20 |
| H1N1 CA09 HA-N95L trimer (Kif/Endo H) | 20 | 20 | 20 | 20 | 20 | 20 | 20 | 20 |
| H1N1 CA09 HA-N95L trimer (Kif/Endo H) | 20 | 20 | 20 | 20 | 20 | 20 | 20 | 20 |
| H1N1 CA09 HA-N95L FR SApNP (Wild-type) | 20 | 20 | 20 | 20 | 20 | 20 | 20 | 20 |
| H1N1 CA09 HA-N95L FR SApNP (Wild-type) | 20 | 20 | 20 | 20 | 20 | 20 | 20 | 20 |
| H1N1 CA09 HA-N95L FR SApNP (Kif-treated) | 20 | 20 | 20 | 20 | 20 | 20 | 20 | 20 |
| H1N1 CA09 HA-N95L FR SApNP (Kif-treated) | 20 | 20 | 20 | 20 | 20 | 20 | 20 | 20 |
| H1N1 CA09 HA-N95L E2p SApNP (Wild-type) | 20 | 20 | 20 | 20 | 20 | 20 | 20 | 20 |
| H1N1 CA09 HA-N95L E2p SApNP (Wild-type) | 20 | 20 | 20 | 20 | 20 | 20 | 20 | 20 |
| H1N1 CA09 HA-N95L E2p SApNP (Kif-treated) | 20 | 20 | 20 | 20 | 20 | 20 | 20 | 20 |
| H1N1 CA09 HA-N95L E2p SApNP (Kif-treated) | 20 | 20 | 20 | 20 | 20 | 20 | 20 | 20 |
| H1N1 CA09 HA-N95L I3-01v9b SApNP (Wild-type) | 20 | 20 | 20 | 20 | 20 | 20 | 20 | 20 |
| H1N1 CA09 HA-N95L I3-01v9b SApNP (Wild-type) | 20 | 20 | 20 | 20 | 20 | 20 | 20 | 20 |
| H1N1 CA09 HA-N95L I3-01v9b SApNP (Kif-treated) | 20 | 20 | 20 | 20 | 20 | 20 | 20 | 20 |
| H1N1 CA09 HA-N95L I3-01v9b SApNP (Kif-treated) | 20 | 20 | 20 | 20 | 20 | 20 | 20 | 20 |

### A/MI/45/2015

| Antigen | HAI Titer ($Log_2$) (week 8) | | | | | | | |
|---|---|---|---|---|---|---|---|---|
| | M1 | M2 | M3 | M4 | M5 | M6 | M7 | M8 |
| H1N1 CA09 HA-N95L trimer (Wild-type) | 160 | 160 | 160 | 320 | 160 | 20 | 80 | 80 |
| H1N1 CA09 HA-N95L trimer (Wild-type) | 160 | 160 | 160 | 320 | 160 | 20 | 40 | 80 |
| H1N1 CA09 HA-N95L trimer (Kif-treated) | 40 | 160 | 1280 | 80 | 20 | 320 | 20 | 20 |
| H1N1 CA09 HA-N95L trimer (Kif-treated) | 40 | 160 | 1280 | 80 | 20 | 320 | 20 | 20 |
| H1N1 CA09 HA-N95L trimer (Kif/Endo H) | 40 | 320 | 40 | 320 | 320 | 160 | 20 | 320 |
| H1N1 CA09 HA-N95L trimer (Kif/Endo H) | 40 | 320 | 40 | 320 | 320 | 160 | 20 | 320 |
| H1N1 CA09 HA-N95L FR SApNP (Wild-type) | 160 | 640 | 160 | 320 | 640 | 160 | 320 | 640 |
| H1N1 CA09 HA-N95L FR SApNP (Wild-type) | 160 | 1280 | 160 | 320 | 640 | 160 | 320 | 640 |
| H1N1 CA09 HA-N95L FR SApNP (Kif-treated) | 640 | 80 | 320 | 640 | 640 | 320 | 160 | 80 |
| H1N1 CA09 HA-N95L FR SApNP (Kif-treated) | 640 | 80 | 320 | 640 | 640 | 320 | 160 | 80 |
| H1N1 CA09 HA-N95L E2p SApNP (Wild-type) | 320 | 160 | 320 | 320 | 640 | 640 | 160 | 320 |
| H1N1 CA09 HA-N95L E2p SApNP (Wild-type) | 160 | 160 | 320 | 320 | 640 | 640 | 160 | 320 |
| H1N1 CA09 HA-N95L E2p SApNP (Kif-treated) | 640 | 160 | 160 | 320 | 160 | 160 | 320 | 80 |
| H1N1 CA09 HA-N95L E2p SApNP (Kif-treated) | 640 | 160 | 160 | 320 | 160 | 160 | 320 | 80 |
| H1N1 CA09 HA-N95L I3-01v9b SApNP (Wild-type) | 160 | 320 | 160 | 320 | 80 | 80 | 160 | 320 |
| H1N1 CA09 HA-N95L I3-01v9b SApNP (Wild-type) | 60 | 320 | 160 | 320 | 80 | 80 | 160 | 320 |
| H1N1 CA09 HA-N95L I3-01v9b SApNP (Kif-treated) | 320 | 320 | 80 | 160 | 160 | 160 | 320 | 160 |
| H1N1 CA09 HA-N95L I3-01v9b SApNP (Kif-treated) | 320 | 320 | 80 | 160 | 160 | 160 | 320 | 160 |

#### Statistical analysis

A/MI/45/2015

| One-way ANOVA with Tukey's multiple comparisons test (w8) | Statistics | Adjusted P Value |
|---|---|---|
| HA trimer-Wild-type vs. HA trimer-Kif-treated | ns | 0.9914 |
| HA trimer-Wild-type vs. HA trimer-Kif/Endo H | ns | >0.9999 |
| HA trimer-Wild-type vs. HA FR SApNP-Wild-type | ns | 0.2548 |
| HA trimer-Wild-type vs. HA FR SApNP-Kif-treated | ns | 0.5765 |
| HA trimer-Wild-type vs. HA E2p SApNP-Wild-type | ns | 0.636 |
| HA trimer-Wild-type vs. HA E2p SApNP-Kif-treated | ns | 0.9864 |
| HA trimer-Wild-type vs. HA I3-01v9b SApNP-Wild-type | ns | >0.9999 |
| HA trimer-Wild-type vs. HA I3-01v9b SApNP-Kif-treated | ns | 0.9994 |
| HA trimer-Kif-treated vs. HA trimer-Kif/Endo H | ns | >0.9999 |
| HA trimer-Kif-treated vs. HA FR SApNP-Wild-type | ns | 0.8114 |
| HA trimer-Kif-treated vs. HA FR SApNP-Kif-treated | ns | 0.9795 |
| HA trimer-Kif-treated vs. HA E2p SApNP-Wild-type | ns | 0.9883 |
| HA trimer-Kif-treated vs. HA E2p SApNP-Kif-treated | ns | >0.9999 |
| HA trimer-Kif-treated vs. HA I3-01v9b SApNP-Wild-type | ns | >0.9999 |
| HA trimer-Kif-treated vs. HA I3-01v9b SApNP-Kif-treated | ns | >0.9999 |
| HA trimer-Kif/Endo H vs. HA FR SApNP-Wild-type | ns | 0.5316 |
| HA trimer-Kif/Endo H vs. HA FR SApNP-Kif-treated | ns | 0.8551 |
| HA trimer-Kif/Endo H vs. HA E2p SApNP-Wild-type | ns | 0.8926 |
| HA trimer-Kif/Endo H vs. HA E2p SApNP-Kif-treated | ns | 0.9999 |
| HA trimer-Kif/Endo H vs. HA I3-01v9b SApNP-Wild-type | ns | >0.9999 |
| HA trimer-Kif/Endo H vs. HA I3-01v9b SApNP-Kif-treated | ns | >0.9999 |
| HA FR SApNP-Wild-type vs. HA FR SApNP-Kif-treated | ns | 0.9998 |
| HA FR SApNP-Wild-type vs. HA E2p SApNP-Wild-type | ns | 0.9994 |
| HA FR SApNP-Wild-type vs. HA E2p SApNP-Kif-treated | ns | 0.8448 |
| HA FR SApNP-Wild-type vs. HA I3-01v9b SApNP-Wild-type | ns | 0.5391 |
| HA FR SApNP-Wild-type vs. HA I3-01v9b SApNP-Kif-treated | ns | 0.636 |
| HA FR SApNP-Kif-treated vs. HA E2p SApNP-Wild-type | ns | >0.9999 |
| HA FR SApNP-Kif-treated vs. HA E2p SApNP-Kif-treated | ns | 0.9864 |
| HA FR SApNP-Kif-treated vs. HA I3-01v9b SApNP-Wild-type | ns | 0.8602 |
| HA FR SApNP-Kif-treated vs. HA I3-01v9b SApNP-Kif-treated | ns | 0.9164 |
| HA E2p SApNP-Wild-type vs. HA E2p SApNP-Kif-treated | ns | 0.9927 |
| HA E2p SApNP-Wild-type vs. HA I3-01v9b SApNP-Wild-type | ns | 0.8968 |
| HA E2p SApNP-Wild-type vs. HA I3-01v9b SApNP-Kif-treated | ns | 0.9424 |
| HA E2p SApNP-Kif-treated vs. HA I3-01v9b SApNP-Wild-type | ns | 0.9999 |
| HA E2p SApNP-Kif-treated vs. HA I3-01v9b SApNP-Kif-treated | ns | >0.9999 |
| HA I3-01v9b SApNP-Wild-type vs. HA I3-01v9b SApNP-Kif-treated | ns | >0.9999 |

### A/WI/67/2022

| Antigen | HAI Titer ($Log_2$) (week 8) | | | | | | | |
|---|---|---|---|---|---|---|---|---|
| | M1 | M2 | M3 | M4 | M5 | M6 | M7 | M8 |
| H1N1 CA09 HA-N95L trimer (Wild-type) | 20 | 20 | 20 | 20 | 20 | 40 | 20 | 20 |
| H1N1 CA09 HA-N95L trimer (Wild-type) | 20 | 20 | 20 | 20 | 20 | 40 | 20 | 20 |
| H1N1 CA09 HA-N95L trimer (Kif-treated) | 20 | 40 | 40 | 20 | 20 | 20 | 20 | 20 |
| H1N1 CA09 HA-N95L trimer (Kif-treated) | 20 | 40 | 40 | 20 | 20 | 20 | 20 | 20 |
| H1N1 CA09 HA-N95L trimer (Kif/Endo H) | 20 | 40 | 20 | 20 | 20 | 20 | 20 | 20 |
| H1N1 CA09 HA-N95L trimer (Kif/Endo H) | 20 | 40 | 20 | 20 | 20 | 20 | 20 | 20 |
| H1N1 CA09 HA-N95L FR SApNP (Wild-type) | 20 | 320 | 160 | 160 | 160 | 20 | 160 | 80 |
| H1N1 CA09 HA-N95L FR SApNP (Wild-type) | 20 | 320 | 160 | 160 | 160 | 20 | 160 | 80 |
| H1N1 CA09 HA-N95L FR SApNP (Kif-treated) | 160 | 20 | 160 | 20 | 80 | 80 | 20 | 20 |
| H1N1 CA09 HA-N95L FR SApNP (Kif-treated) | 160 | 20 | 160 | 20 | 80 | 80 | 20 | 20 |
| H1N1 CA09 HA-N95L E2p SApNP (Wild-type) | 40 | 20 | 40 | 20 | 80 | 20 | 40 | 40 |
| H1N1 CA09 HA-N95L E2p SApNP (Wild-type) | 40 | 20 | 40 | 20 | 80 | 20 | 40 | 40 |
| H1N1 CA09 HA-N95L E2p SApNP (Kif-treated) | 160 | 20 | 20 | 20 | 20 | 40 | 20 | 40 |
| H1N1 CA09 HA-N95L E2p SApNP (Kif-treated) | 320 | 20 | 20 | 20 | 20 | 40 | 20 | 40 |
| H1N1 CA09 HA-N95L I3-01v9b SApNP (Wild-type) | 20 | 80 | 80 | 40 | 20 | 20 | 160 | 40 |
| H1N1 CA09 HA-N95L I3-01v9b SApNP (Wild-type) | 20 | 80 | 80 | 40 | 20 | 20 | 160 | 40 |
| H1N1 CA09 HA-N95L I3-01v9b SApNP (Kif-treated) | 80 | 40 | 20 | 40 | 20 | 320 | 20 | 20 |
| H1N1 CA09 HA-N95L I3-01v9b SApNP (Kif-treated) | 80 | 40 | 20 | 40 | 20 | 320 | 20 | 20 |

A/WI/67/2022

| One-way ANOVA with Tukey's multiple comparisons test (w8) | Statistics | Adjusted P Value |
|---|---|---|
| HA trimer-Wild-type vs. HA trimer-Kif-treated | ns | >0.9999 |
| HA trimer-Wild-type vs. HA trimer-Kif/Endo H | ns | >0.9999 |
| HA trimer-Wild-type vs. HA FR SApNP-Wild-type | * | 0.0252 |
| HA trimer-Wild-type vs. HA FR SApNP-Kif-treated | ns | 0.8718 |
| HA trimer-Wild-type vs. HA E2p SApNP-Wild-type | ns | >0.9999 |
| HA trimer-Wild-type vs. HA E2p SApNP-Kif-treated | ns | 0.9911 |
| HA trimer-Wild-type vs. HA I3-01v9b SApNP-Wild-type | ns | 0.9764 |
| HA trimer-Wild-type vs. HA I3-01v9b SApNP-Kif-treated | ns | 0.7561 |
| HA trimer-Kif-treated vs. HA trimer-Kif/Endo H | ns | >0.9999 |
| HA trimer-Kif-treated vs. HA FR SApNP-Wild-type | * | 0.0313 |
| HA trimer-Kif-treated vs. HA FR SApNP-Kif-treated | ns | 0.9021 |
| HA trimer-Kif-treated vs. HA E2p SApNP-Wild-type | ns | >0.9999 |
| HA trimer-Kif-treated vs. HA E2p SApNP-Kif-treated | ns | 0.9951 |
| HA trimer-Kif-treated vs. HA I3-01v9b SApNP-Wild-type | ns | 0.9851 |
| HA trimer-Kif-treated vs. HA I3-01v9b SApNP-Kif-treated | ns | 0.8059 |
| HA trimer-Kif/Endo H vs. HA FR SApNP-Wild-type | * | 0.0252 |
| HA trimer-Kif/Endo H vs. HA FR SApNP-Kif-treated | ns | 0.8718 |
| HA trimer-Kif/Endo H vs. HA E2p SApNP-Wild-type | ns | >0.9999 |
| HA trimer-Kif/Endo H vs. HA E2p SApNP-Kif-treated | ns | 0.9911 |
| HA trimer-Kif/Endo H vs. HA I3-01v9b SApNP-Wild-type | ns | 0.9764 |
| HA trimer-Kif/Endo H vs. HA I3-01v9b SApNP-Kif-treated | ns | 0.7561 |
| HA FR SApNP-Wild-type vs. HA FR SApNP-Kif-treated | ns | 0.5543 |
| HA FR SApNP-Wild-type vs. HA E2p SApNP-Wild-type | ns | 0.086 |
| HA FR SApNP-Wild-type vs. HA E2p SApNP-Kif-treated | ns | 0.2368 |
| HA FR SApNP-Wild-type vs. HA I3-01v9b SApNP-Wild-type | ns | 0.314 |
| HA FR SApNP-Wild-type vs. HA I3-01v9b SApNP-Kif-treated | ns | 0.3564 |
| HA FR SApNP-Kif-treated vs. HA E2p SApNP-Wild-type | ns | 0.9851 |
| HA FR SApNP-Kif-treated vs. HA E2p SApNP-Kif-treated | ns | 0.9998 |
| HA FR SApNP-Kif-treated vs. HA I3-01v9b SApNP-Wild-type | ns | >0.9999 |
| HA FR SApNP-Kif-treated vs. HA I3-01v9b SApNP-Kif-treated | ns | >0.9999 |
| HA E2p SApNP-Wild-type vs. HA E2p SApNP-Kif-treated | ns | >0.9999 |
| HA E2p SApNP-Wild-type vs. HA I3-01v9b SApNP-Wild-type | ns | 0.9995 |
| HA E2p SApNP-Wild-type vs. HA I3-01v9b SApNP-Kif-treated | ns | 0.9641 |
| HA E2p SApNP-Kif-treated vs. HA I3-01v9b SApNP-Wild-type | ns | >0.9999 |
| HA E2p SApNP-Kif-treated vs. HA I3-01v9b SApNP-Kif-treated | ns | 0.9994 |
| HA I3-01v9b SApNP-Wild-type vs. HA I3-01v9b SApNP-Kif-treated | ns | >0.9999 |

g

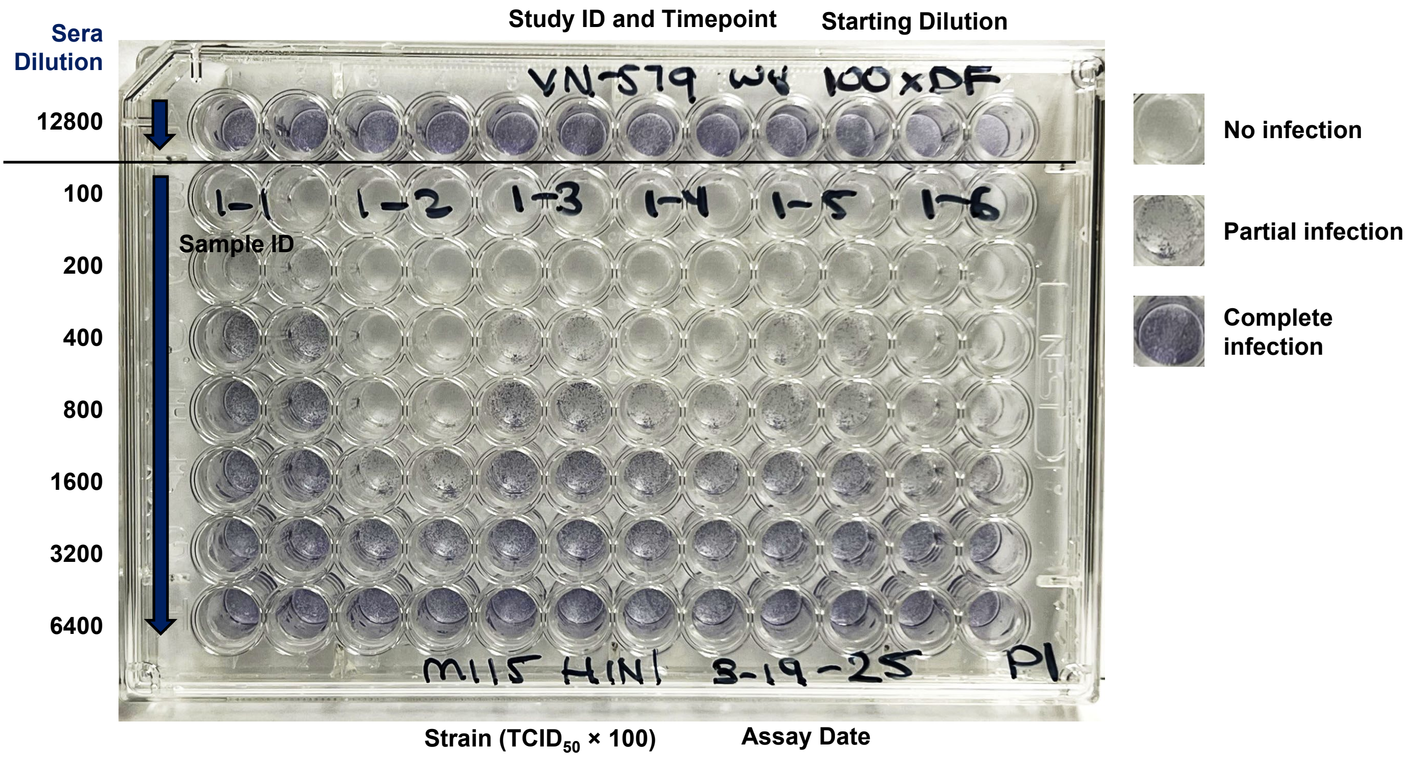

h **H1N1 CA09 HA-induced NAb responses against H1N1 PR8, NC99, CA09, MI15, and WI22 viruses**

**A/PR/8/1934**

| Antigen | Neutralization Titer (week 8) | | | | | | | |
|---|---|---|---|---|---|---|---|---|
| | M1 | M2 | M3 | M4 | M5 | M6 | M7 | M8 |
| H1N1 CA09 HA-N95L trimer (Wild-type) | 50 | 50 | 50 | 50 | 50 | 50 | 50 | 50 |
| H1N1 CA09 HA-N95L trimer (Wild-type) | 50 | 50 | 50 | 50 | 50 | 50 | 50 | 50 |
| H1N1 CA09 HA-N95L trimer (Kif-treated) | 50 | 50 | 50 | 50 | 50 | 50 | 50 | 50 |
| H1N1 CA09 HA-N95L trimer (Kif-treated) | 50 | 50 | 50 | 50 | 50 | 50 | 50 | 50 |
| H1N1 CA09 HA-N95L trimer (Kif/Endo H) | 50 | 50 | 50 | 50 | 50 | 50 | 50 | 50 |
| H1N1 CA09 HA-N95L trimer (Kif/Endo H) | 50 | 50 | 50 | 50 | 50 | 50 | 50 | 50 |
| H1N1 CA09 HA-N95L FR SApNP (Wild-type) | 50 | 50 | 50 | 50 | 50 | 50 | 50 | 50 |
| H1N1 CA09 HA-N95L FR SApNP (Wild-type) | 50 | 50 | 50 | 50 | 50 | 50 | 50 | 50 |
| H1N1 CA09 HA-N95L FR SApNP (Kif-treated) | 50 | 50 | 50 | 50 | 50 | 50 | 50 | 50 |
| H1N1 CA09 HA-N95L FR SApNP (Kif-treated) | 50 | 50 | 50 | 50 | 50 | 50 | 50 | 50 |
| H1N1 CA09 HA-N95L E2p SApNP (Wild-type) | 50 | 50 | 50 | 50 | 50 | 50 | 50 | 50 |
| H1N1 CA09 HA-N95L E2p SApNP (Wild-type) | 50 | 50 | 50 | 50 | 50 | 50 | 50 | 50 |
| H1N1 CA09 HA-N95L E2p SApNP (Kif-treated) | 50 | 50 | 50 | 50 | 50 | 50 | 50 | 50 |
| H1N1 CA09 HA-N95L E2p SApNP (Kif-treated) | 50 | 50 | 50 | 50 | 50 | 50 | 50 | 50 |
| H1N1 CA09 HA-N95L I3-01v9b SApNP (Wild-type) | 50 | 50 | 50 | 50 | 50 | 50 | 50 | 50 |
| H1N1 CA09 HA-N95L I3-01v9b SApNP (Wild-type) | 50 | 50 | 50 | 50 | 50 | 50 | 50 | 50 |
| H1N1 CA09 HA-N95L I3-01v9b SApNP (Kif-treated) | 50 | 50 | 50 | 50 | 50 | 50 | 50 | 50 |
| H1N1 CA09 HA-N95L I3-01v9b SApNP (Kif-treated) | 50 | 50 | 50 | 50 | 50 | 50 | 50 | 50 |

**A/NC/20/1999**

| Antigen | Neutralization Titer (week 8) | | | | | | | |
|---|---|---|---|---|---|---|---|---|
| | M1 | M2 | M3 | M4 | M5 | M6 | M7 | M8 |
| H1N1 CA09 HA-N95L trimer (Wild-type) | 50 | 50 | 50 | 50 | 50 | 50 | 50 | 50 |
| H1N1 CA09 HA-N95L trimer (Wild-type) | 50 | 50 | 50 | 50 | 50 | 50 | 50 | 50 |
| H1N1 CA09 HA-N95L trimer (Kif-treated) | 50 | 50 | 50 | 50 | 50 | 50 | 50 | 50 |
| H1N1 CA09 HA-N95L trimer (Kif-treated) | 50 | 50 | 50 | 50 | 50 | 50 | 50 | 50 |
| H1N1 CA09 HA-N95L trimer (Kif/Endo H) | 50 | 50 | 50 | 50 | 50 | 50 | 50 | 50 |
| H1N1 CA09 HA-N95L trimer (Kif/Endo H) | 50 | 50 | 50 | 50 | 50 | 50 | 50 | 50 |
| H1N1 CA09 HA-N95L FR SApNP (Wild-type) | 50 | 50 | 50 | 50 | 50 | 50 | 50 | 50 |
| H1N1 CA09 HA-N95L FR SApNP (Wild-type) | 50 | 50 | 50 | 50 | 50 | 50 | 50 | 50 |
| H1N1 CA09 HA-N95L FR SApNP (Kif-treated) | 50 | 50 | 50 | 50 | 50 | 50 | 50 | 50 |
| H1N1 CA09 HA-N95L FR SApNP (Kif-treated) | 50 | 50 | 50 | 50 | 50 | 50 | 50 | 50 |
| H1N1 CA09 HA-N95L E2p SApNP (Wild-type) | 50 | 50 | 50 | 50 | 50 | 50 | 50 | 50 |
| H1N1 CA09 HA-N95L E2p SApNP (Wild-type) | 50 | 50 | 50 | 50 | 50 | 50 | 50 | 50 |
| H1N1 CA09 HA-N95L E2p SApNP (Kif-treated) | 50 | 50 | 50 | 50 | 50 | 50 | 50 | 50 |
| H1N1 CA09 HA-N95L E2p SApNP (Kif-treated) | 50 | 50 | 50 | 50 | 50 | 50 | 50 | 50 |
| H1N1 CA09 HA-N95L I3-01v9b SApNP (Wild-type) | 50 | 50 | 50 | 50 | 50 | 50 | 50 | 50 |
| H1N1 CA09 HA-N95L I3-01v9b SApNP (Wild-type) | 50 | 50 | 50 | 50 | 50 | 50 | 50 | 50 |
| H1N1 CA09 HA-N95L I3-01v9b SApNP (Kif-treated) | 50 | 50 | 50 | 50 | 50 | 50 | 50 | 50 |
| H1N1 CA09 HA-N95L I3-01v9b SApNP (Kif-treated) | 50 | 50 | 50 | 50 | 50 | 50 | 50 | 50 |

**A/CA/07/2009**

| Antigen | Neutralization Titer (week 8) | | | | | | | |
|---|---|---|---|---|---|---|---|---|
| | M1 | M2 | M3 | M4 | M5 | M6 | M7 | M8 |
| H1N1 CA09 HA-N95L trimer (Wild-type) | 1600 | 3200 | 1600 | 1600 | 1600 | 1600 | 3200 | 1600 |
| H1N1 CA09 HA-N95L trimer (Wild-type) | 1600 | 3200 | 1600 | 1600 | 1600 | 1600 | 3200 | 1600 |
| H1N1 CA09 HA-N95L trimer (Kif-treated) | 1600 | 3200 | 3200 | 3200 | 1600 | 1600 | 1600 | 1600 |
| H1N1 CA09 HA-N95L trimer (Kif-treated) | 1600 | 3200 | 3200 | 3200 | 1600 | 1600 | 1600 | 1600 |
| H1N1 CA09 HA-N95L trimer (Kif/Endo H) | 400 | 400 | 400 | 400 | 800 | 800 | 800 | 400 |
| H1N1 CA09 HA-N95L trimer (Kif/Endo H) | 400 | 400 | 400 | 400 | 800 | 800 | 800 | 400 |
| H1N1 CA09 HA-N95L FR SApNP (Wild-type) | 3200 | 6400 | 3200 | 3200 | 3200 | 1600 | 1600 | 3200 |
| H1N1 CA09 HA-N95L FR SApNP (Wild-type) | 3200 | 6400 | 3200 | 3200 | 3200 | 1600 | 1600 | 3200 |
| H1N1 CA09 HA-N95L FR SApNP (Kif-treated) | 3200 | 3200 | 1600 | 6400 | 6400 | 3200 | 3200 | 1600 |
| H1N1 CA09 HA-N95L FR SApNP (Kif-treated) | 3200 | 1600 | 3200 | 6400 | 6400 | 3200 | 1600 | 1600 |
| H1N1 CA09 HA-N95L E2p SApNP (Wild-type) | 1600 | 1600 | 1600 | 1600 | 800 | 3200 | 1600 | 1600 |
| H1N1 CA09 HA-N95L E2p SApNP (Wild-type) | 1600 | 1600 | 1600 | 1600 | 800 | 3200 | 1600 | 1600 |
| H1N1 CA09 HA-N95L E2p SApNP (Kif-treated) | 3200 | 1600 | 3200 | 3200 | 3200 | 3200 | 6400 | 1600 |
| H1N1 CA09 HA-N95L E2p SApNP (Kif-treated) | 3200 | 3200 | 3200 | 3200 | 3200 | 3200 | 6400 | 1600 |
| H1N1 CA09 HA-N95L I3-01v9b SApNP (Wild-type) | 3200 | 1600 | 1600 | 3200 | 800 | 1600 | 1600 | 1600 |
| H1N1 CA09 HA-N95L I3-01v9b SApNP (Wild-type) | 3200 | 1600 | 1600 | 3200 | 1600 | 1600 | 1600 | 1600 |
| H1N1 CA09 HA-N95L I3-01v9b SApNP (Kif-treated) | 1600 | 1600 | 1600 | 800 | 800 | 3200 | 3200 | 1600 |
| H1N1 CA09 HA-N95L I3-01v9b SApNP (Kif-treated) | 1600 | 1600 | 1600 | 800 | 800 | 3200 | 3200 | 1600 |

**Statistical analysis**

**A/CA/07/2009**

| One-way ANOVA with Tukey's multiple comparisons test (w8) | Statistics | Adjusted P Value |
|---|---|---|
| HA trimer-Wild-type vs. HA trimer-Kif-treated | ns | >0.9999 |
| HA trimer-Wild-type vs. HA trimer-Kif/Endo H | ns | 0.1871 |
| HA trimer-Wild-type vs. HA FR SApNP-Wild-type | ns | 0.4223 |
| HA trimer-Wild-type vs. HA FR SApNP-Kif-treated | ns | 0.1543 |
| HA trimer-Wild-type vs. HA E2p SApNP-Wild-type | ns | 0.9998 |
| HA trimer-Wild-type vs. HA E2p SApNP-Kif-treated | ns | 0.3147 |
| HA trimer-Wild-type vs. HA I3-01v9b SApNP-Wild-type | ns | >0.9999 |
| HA trimer-Wild-type vs. HA I3-01v9b SApNP-Kif-treated | ns | >0.9999 |
| HA trimer-Kif-treated vs. HA trimer-Kif/Endo H | ns | 0.082 |
| HA trimer-Kif-treated vs. HA FR SApNP-Wild-type | ns | 0.6637 |
| HA trimer-Kif-treated vs. HA FR SApNP-Kif-treated | ns | 0.3147 |
| HA trimer-Kif-treated vs. HA E2p SApNP-Wild-type | ns | 0.9914 |
| HA trimer-Kif-treated vs. HA E2p SApNP-Kif-treated | ns | 0.5418 |
| HA trimer-Kif-treated vs. HA I3-01v9b SApNP-Wild-type | ns | >0.9999 |
| HA trimer-Kif-treated vs. HA I3-01v9b SApNP-Kif-treated | ns | 0.9981 |
| HA trimer-Kif/Endo H vs. HA FR SApNP-Wild-type | *** | 0.0003 |
| HA trimer-Kif/Endo H vs. HA FR SApNP-Kif-treated | **** | <0.0001 |
| HA trimer-Kif/Endo H vs. HA E2p SApNP-Wild-type | ns | 0.4811 |
| HA trimer-Kif/Endo H vs. HA E2p SApNP-Kif-treated | *** | 0.0001 |
| HA trimer-Kif/Endo H vs. HA I3-01v9b SApNP-Wild-type | ns | 0.2248 |
| HA trimer-Kif/Endo H vs. HA I3-01v9b SApNP-Kif-treated | ns | 0.3665 |
| HA FR SApNP-Wild-type vs. HA FR SApNP-Kif-treated | ns | 0.9998 |
| HA FR SApNP-Wild-type vs. HA E2p SApNP-Wild-type | ns | 0.1543 |
| HA FR SApNP-Wild-type vs. HA E2p SApNP-Kif-treated | ns | >0.9999 |
| HA FR SApNP-Wild-type vs. HA I3-01v9b SApNP-Wild-type | ns | 0.3665 |
| HA FR SApNP-Wild-type vs. HA I3-01v9b SApNP-Kif-treated | ns | 0.2248 |
| HA FR SApNP-Kif-treated vs. HA E2p SApNP-Wild-type | * | 0.0406 |
| HA FR SApNP-Kif-treated vs. HA E2p SApNP-Kif-treated | ns | >0.9999 |
| HA FR SApNP-Kif-treated vs. HA I3-01v9b SApNP-Wild-type | ns | 0.1261 |
| HA FR SApNP-Kif-treated vs. HA I3-01v9b SApNP-Kif-treated | ns | 0.0654 |
| HA E2p SApNP-Wild-type vs. HA E2p SApNP-Kif-treated | ns | 0.1021 |
| HA E2p SApNP-Wild-type vs. HA I3-01v9b SApNP-Wild-type | ns | >0.9999 |
| HA E2p SApNP-Wild-type vs. HA I3-01v9b SApNP-Kif-treated | ns | >0.9999 |
| HA E2p SApNP-Kif-treated vs. HA I3-01v9b SApNP-Wild-type | ns | 0.2673 |
| HA E2p SApNP-Kif-treated vs. HA I3-01v9b SApNP-Kif-treated | ns | 0.1543 |
| HA I3-01v9b SApNP-Wild-type vs. HA I3-01v9b SApNP-Kif-treated | ns | >0.9999 |

### Fig. S10

A/MI/45/2015

| Antigen | Neutralization Titer (week 8) | | | | | | | |
|---|---|---|---|---|---|---|---|---|
| | M1 | M2 | M3 | M4 | M5 | M6 | M7 | M8 |
| H1N1 CA09 HA-N95L trimer (Wild-type) | 100 | 400 | 100 | 200 | 100 | 200 | 200 | 50 |
| H1N1 CA09 HA-N95L trimer (Wild-type) | 100 | 400 | 100 | 200 | 100 | 200 | 200 | 50 |
| H1N1 CA09 HA-N95L trimer (Kif-treated) | 100 | 800 | 1600 | 200 | 50 | 400 | 50 | 50 |
| H1N1 CA09 HA-N95L trimer (Kif-treated) | 100 | 800 | 1600 | 200 | 50 | 400 | 50 | 50 |
| H1N1 CA09 HA-N95L trimer (Kif/Endo H) | 50 | 400 | 50 | 200 | 100 | 50 | 100 | 200 |
| H1N1 CA09 HA-N95L trimer (Kif/Endo H) | 50 | 400 | 50 | 200 | 100 | 50 | 100 | 200 |
| H1N1 CA09 HA-N95L FR SApNP (Wild-type) | 100 | 1600 | 100 | 400 | 800 | 100 | 400 | 400 |
| H1N1 CA09 HA-N95L FR SApNP (Wild-type) | 100 | 1600 | 100 | 400 | 800 | 100 | 400 | 400 |
| H1N1 CA09 HA-N95L FR SApNP (Kif-treated) | 400 | 200 | 800 | 800 | 800 | 400 | 400 | 100 |
| H1N1 CA09 HA-N95L FR SApNP (Kif-treated) | 400 | 200 | 800 | 800 | 800 | 400 | 400 | 100 |
| H1N1 CA09 HA-N95L E2p SApNP (Wild-type) | 200 | 200 | 200 | 200 | 400 | 400 | 100 | 100 |
| H1N1 CA09 HA-N95L E2p SApNP (Wild-type) | 200 | 200 | 200 | 200 | 400 | 400 | 100 | 100 |
| H1N1 CA09 HA-N95L E2p SApNP (Kif-treated) | 400 | 100 | 100 | 200 | 200 | 200 | 200 | 200 |
| H1N1 CA09 HA-N95L E2p SApNP (Kif-treated) | 400 | 100 | 100 | 200 | 200 | 200 | 200 | 200 |
| H1N1 CA09 HA-N95L I3-01v9b SApNP (Wild-type) | 100 | 400 | 200 | 200 | 200 | 50 | 200 | 200 |
| H1N1 CA09 HA-N95L I3-01v9b SApNP (Wild-type) | 100 | 400 | 200 | 200 | 200 | 50 | 200 | 200 |
| H1N1 CA09 HA-N95L I3-01v9b SApNP (Kif-treated) | 400 | 200 | 100 | 100 | 100 | 400 | 400 | 100 |
| H1N1 CA09 HA-N95L I3-01v9b SApNP (Kif-treated) | 400 | 200 | 100 | 100 | 100 | 400 | 400 | 100 |

#### Statistical analysis

A/MI/45/2015

| One-way ANOVA with Tukey's multiple comparisons test (w8) | Statistics | Adjusted P Value |
|---|---|---|
| HA trimer-Wild-type vs. HA trimer-Kif-treated | ns | 0.7548 |
| HA trimer-Wild-type vs. HA trimer-Kif/Endo H | ns | >0.9999 |
| HA trimer-Wild-type vs. HA FR SApNP-Wild-type | ns | 0.383 |
| HA trimer-Wild-type vs. HA FR SApNP-Kif-treated | ns | 0.383 |
| HA trimer-Wild-type vs. HA E2p SApNP-Wild-type | ns | >0.9999 |
| HA trimer-Wild-type vs. HA E2p SApNP-Kif-treated | ns | >0.9999 |
| HA trimer-Wild-type vs. HA I3-01v9b SApNP-Wild-type | ns | >0.9999 |
| HA trimer-Wild-type vs. HA I3-01v9b SApNP-Kif-treated | ns | >0.9999 |
| HA trimer-Kif-treated vs. HA trimer-Kif/Endo H | ns | 0.6434 |
| HA trimer-Kif-treated vs. HA FR SApNP-Wild-type | ns | 0.9997 |
| HA trimer-Kif-treated vs. HA FR SApNP-Kif-treated | ns | 0.9997 |
| HA trimer-Kif-treated vs. HA E2p SApNP-Wild-type | ns | 0.9324 |
| HA trimer-Kif-treated vs. HA E2p SApNP-Kif-treated | ns | 0.8689 |
| HA trimer-Kif-treated vs. HA I3-01v9b SApNP-Wild-type | ns | 0.849 |
| HA trimer-Kif-treated vs. HA I3-01v9b SApNP-Kif-treated | ns | 0.9324 |
| HA trimer-Kif/Endo H vs. HA FR SApNP-Wild-type | ns | 0.2841 |
| HA trimer-Kif/Endo H vs. HA FR SApNP-Kif-treated | ns | 0.2841 |
| HA trimer-Kif/Endo H vs. HA E2p SApNP-Wild-type | ns | 0.9997 |
| HA trimer-Kif/Endo H vs. HA E2p SApNP-Kif-treated | ns | >0.9999 |
| HA trimer-Kif/Endo H vs. HA I3-01v9b SApNP-Wild-type | ns | >0.9999 |
| HA trimer-Kif/Endo H vs. HA I3-01v9b SApNP-Kif-treated | ns | 0.9997 |
| HA FR SApNP-Wild-type vs. HA FR SApNP-Kif-treated | ns | >0.9999 |
| HA FR SApNP-Wild-type vs. HA E2p SApNP-Wild-type | ns | 0.6434 |
| HA FR SApNP-Wild-type vs. HA E2p SApNP-Kif-treated | ns | 0.5249 |
| HA FR SApNP-Wild-type vs. HA I3-01v9b SApNP-Wild-type | ns | 0.4955 |
| HA FR SApNP-Wild-type vs. HA I3-01v9b SApNP-Kif-treated | ns | 0.6434 |
| HA FR SApNP-Kif-treated vs. HA E2p SApNP-Wild-type | ns | 0.6434 |
| HA FR SApNP-Kif-treated vs. HA E2p SApNP-Kif-treated | ns | 0.5249 |
| HA FR SApNP-Kif-treated vs. HA I3-01v9b SApNP-Wild-type | ns | 0.4955 |
| HA FR SApNP-Kif-treated vs. HA I3-01v9b SApNP-Kif-treated | ns | 0.6434 |
| HA E2p SApNP-Wild-type vs. HA E2p SApNP-Kif-treated | ns | >0.9999 |
| HA E2p SApNP-Wild-type vs. HA I3-01v9b SApNP-Wild-type | ns | >0.9999 |
| HA E2p SApNP-Wild-type vs. HA I3-01v9b SApNP-Kif-treated | ns | >0.9999 |
| HA E2p SApNP-Kif-treated vs. HA I3-01v9b SApNP-Wild-type | ns | >0.9999 |
| HA E2p SApNP-Kif-treated vs. HA I3-01v9b SApNP-Kif-treated | ns | >0.9999 |
| HA I3-01v9b SApNP-Wild-type vs. HA I3-01v9b SApNP-Kif-treated | ns | >0.9999 |

A/WI/67/2022

| Antigen | Neutralization Titer (week 8) | | | | | | | |
|---|---|---|---|---|---|---|---|---|
| | M1 | M2 | M3 | M4 | M5 | M6 | M7 | M8 |
| H1N1 CA09 HA-N95L trimer (Wild-type) | 100 | 100 | 50 | 100 | 50 | 400 | 50 | 50 |
| H1N1 CA09 HA-N95L trimer (Wild-type) | 100 | 100 | 50 | 100 | 50 | 400 | 50 | 50 |
| H1N1 CA09 HA-N95L trimer (Kif-treated) | 200 | 400 | 400 | 400 | 50 | 200 | 50 | 100 |
| H1N1 CA09 HA-N95L trimer (Kif-treated) | 200 | 400 | 400 | 400 | 50 | 200 | 50 | 100 |
| H1N1 CA09 HA-N95L trimer (Kif/Endo H) | 50 | 50 | 50 | 50 | 50 | 50 | 200 | 50 |
| H1N1 CA09 HA-N95L trimer (Kif/Endo H) | 50 | 50 | 50 | 50 | 50 | 100 | 200 | 50 |
| H1N1 CA09 HA-N95L FR SApNP (Wild-type) | 100 | 800 | 400 | 400 | 400 | 100 | 400 | 200 |
| H1N1 CA09 HA-N95L FR SApNP (Wild-type) | 100 | 800 | 400 | 400 | 400 | 50 | 400 | 200 |
| H1N1 CA09 HA-N95L FR SApNP (Kif-treated) | 400 | 100 | 400 | 100 | 200 | 400 | 100 | 100 |
| H1N1 CA09 HA-N95L FR SApNP (Kif-treated) | 400 | 100 | 400 | 100 | 200 | 400 | 100 | 100 |
| H1N1 CA09 HA-N95L E2p SApNP (Wild-type) | 200 | 50 | 100 | 50 | 100 | 50 | 50 | 50 |
| H1N1 CA09 HA-N95L E2p SApNP (Wild-type) | 200 | 50 | 100 | 50 | 100 | 50 | 50 | 50 |
| H1N1 CA09 HA-N95L E2p SApNP (Kif-treated) | 200 | 50 | 50 | 50 | 100 | 50 | 50 | 100 |
| H1N1 CA09 HA-N95L E2p SApNP (Kif-treated) | 200 | 50 | 50 | 50 | 100 | 50 | 50 | 100 |
| H1N1 CA09 HA-N95L I3-01v9b SApNP (Wild-type) | 50 | 200 | 200 | 100 | 100 | 50 | 200 | 100 |
| H1N1 CA09 HA-N95L I3-01v9b SApNP (Wild-type) | 50 | 200 | 200 | 100 | 100 | 50 | 200 | 100 |
| H1N1 CA09 HA-N95L I3-01v9b SApNP (Kif-treated) | 200 | 100 | 50 | 100 | 100 | 400 | 100 | 50 |
| H1N1 CA09 HA-N95L I3-01v9b SApNP (Kif-treated) | 200 | 50 | 50 | 100 | 100 | 400 | 100 | 50 |

A/WI/67/2022

| One-way ANOVA with Tukey's multiple comparisons test (w8) | Statistics | Adjusted P Value |
|---|---|---|
| HA trimer-Wild-type vs. HA trimer-Kif-treated | ns | 0.6803 |
| HA trimer-Wild-type vs. HA trimer-Kif/Endo H | ns | 0.9992 |
| HA trimer-Wild-type vs. HA FR SApNP-Wild-type | * | 0.0107 |
| HA trimer-Wild-type vs. HA FR SApNP-Kif-treated | ns | 0.6803 |
| HA trimer-Wild-type vs. HA E2p SApNP-Wild-type | ns | 0.9999 |
| HA trimer-Wild-type vs. HA E2p SApNP-Kif-treated | ns | 0.9999 |
| HA trimer-Wild-type vs. HA I3-01v9b SApNP-Wild-type | ns | >0.9999 |
| HA trimer-Wild-type vs. HA I3-01v9b SApNP-Kif-treated | ns | >0.9999 |
| HA trimer-Kif-treated vs. HA trimer-Kif/Endo H | ns | 0.2746 |
| HA trimer-Kif-treated vs. HA FR SApNP-Wild-type | ns | 0.581 |
| HA trimer-Kif-treated vs. HA FR SApNP-Kif-treated | ns | >0.9999 |
| HA trimer-Kif-treated vs. HA E2p SApNP-Wild-type | ns | 0.356 |
| HA trimer-Kif-treated vs. HA E2p SApNP-Kif-treated | ns | 0.356 |
| HA trimer-Kif-treated vs. HA I3-01v9b SApNP-Wild-type | ns | 0.8 |
| HA trimer-Kif-treated vs. HA I3-01v9b SApNP-Kif-treated | ns | 0.8727 |
| HA trimer-Kif/Endo H vs. HA FR SApNP-Wild-type | ** | 0.0013 |
| HA trimer-Kif/Endo H vs. HA FR SApNP-Kif-treated | ns | 0.2746 |
| HA trimer-Kif/Endo H vs. HA E2p SApNP-Wild-type | ns | >0.9999 |
| HA trimer-Kif/Endo H vs. HA E2p SApNP-Kif-treated | ns | >0.9999 |
| HA trimer-Kif/Endo H vs. HA I3-01v9b SApNP-Wild-type | ns | 0.9946 |
| HA trimer-Kif/Endo H vs. HA I3-01v9b SApNP-Kif-treated | ns | 0.9844 |
| HA FR SApNP-Wild-type vs. HA FR SApNP-Kif-treated | ns | 0.581 |
| HA FR SApNP-Wild-type vs. HA E2p SApNP-Wild-type | ** | 0.0022 |
| HA FR SApNP-Wild-type vs. HA E2p SApNP-Kif-treated | ** | 0.0022 |
| HA FR SApNP-Wild-type vs. HA I3-01v9b SApNP-Wild-type | * | 0.0194 |
| HA FR SApNP-Wild-type vs. HA I3-01v9b SApNP-Kif-treated | * | 0.0297 |
| HA FR SApNP-Kif-treated vs. HA E2p SApNP-Wild-type | ns | 0.356 |
| HA FR SApNP-Kif-treated vs. HA E2p SApNP-Kif-treated | ns | 0.356 |
| HA FR SApNP-Kif-treated vs. HA I3-01v9b SApNP-Wild-type | ns | 0.8 |
| HA FR SApNP-Kif-treated vs. HA I3-01v9b SApNP-Kif-treated | ns | 0.8727 |
| HA E2p SApNP-Wild-type vs. HA E2p SApNP-Kif-treated | ns | >0.9999 |
| HA E2p SApNP-Wild-type vs. HA I3-01v9b SApNP-Wild-type | ns | 0.9986 |
| HA E2p SApNP-Wild-type vs. HA I3-01v9b SApNP-Kif-treated | ns | 0.9946 |
| HA E2p SApNP-Kif-treated vs. HA I3-01v9b SApNP-Wild-type | ns | 0.9986 |
| HA E2p SApNP-Kif-treated vs. HA I3-01v9b SApNP-Kif-treated | ns | 0.9946 |
| HA I3-01v9b SApNP-Wild-type vs. HA I3-01v9b SApNP-Kif-treated | ns | >0.9999 |

**Fig. S10. Immunogenicity of H1N1 CA09 HA vaccines in mice.** (**a**) ELISA curves of mouse sera from H1N1 CA09 HA-N95L trimer and SApNP vaccine groups (n = 8 mice/group) binding to the coating antigen to H1N1 CA09 HA-N95L(1TD0) trimer (**b**) (Top) Summary of geometric mean $EC_{50}$ titers measured for H1N1 CA09 HA-N95L vaccine groups against H1N1 CA09 HA-N95L(1TD0) trimer. Color coding indicates $EC_{50}$ levels (green to red: low to high binding). (Bottom) Summary of statistical analysis performed for each timepoint. (**c**) ELISA curves of mouse sera from H1N1 CA09 HA-N95L vaccine groups at week 8 after three immunizations binding to H3N2 HK68 HA-N95L(1TD0) trimer. Summary of geometric mean $EC_{50}$ titers and statistical analysis. (**d**) Hemagglutination inhibition (HAI) titers elicited by HA trimer and SApNPs vaccines assay plate template. Each receptor-destroy enzyme (RDE)-treated serum sample was diluted to produce a 40-fold dilution, which then underwent a 2-fold dilution series. Next, 8 hemagglutination units (HA units) of virus were incubated with serum samples for 30 min at room temperature (RT), after which 0.75% w/v turkey red blood cells (RBCs) were added to each well and incubated for 30 min at RT. No agglutination (minimal virus binding to RBCs) was visualized as a "teardrop" formation in the well. Partial agglutination (partial virus binding to RBCs) was visualized as a partial spot formation. Complete agglutination (complete virus binding to RBCs) was visualized by a clear appearance in the well (no red spot formation). HAI titers were thereby determined by the highest sera dilution at which complete hemagglutination inhibition (no agglutination) was observed. Each sample was run in duplicate. Virus only control wells included virus + RBCs only and RBC only control wells included RBCs only. (**e**) H1N1 CA09 HA-N95L-induced sera HAI against vaccine matched H1N1 CA09 virus. Color coding indicates HAI titers (green to red: low to high binding). Summary of statistical analysis performed for each timepoint. (**f**) H1N1 CA09 HA-N95L-induced sera HAI against non-matched H1N1 PR8, NC99, MI15, and WI22 viruses at week 8 after three immunizations. Summary of HAI titers and statistical analysis. (**g**) Neutralization titers elicited by HA trimer and SApNPs vaccines assay plate template. Each receptor-destroy enzyme (RDE)-treated serum sample was diluted to produce at least a 100-fold dilution, was further subjected to a 2-fold dilution series with $TCID_{50} \times 100$ of virus, and was then incubated for 1 h at 37 °C. Next, dilutions were transferred to washed, pre-plated MDCK cells and incubated for 24 h at 37 °C. Supernatants were then removed, and cells were fixed, permeabilized, and stained with anti-HA antibody MEDI8852 (primary antibody), followed by goat anti-mouse IgG (secondary antibody). Infected wells were visualized using True Blue substrate. No virus infection was visualized as no blue color in the well. Partial infection was visualized by interspersed blue color in the well. Complete infection was visualized by blue color in the full well. Neutralization titers were thereby determined by the highest sera dilution at which no blue color (no infection) was observed. Each sample was run in duplicate. Antibody MEDI8852 was used as a positive control and antibody FLD 194 was used as a negative control. (**h**) H1N1 CA09 HA-N95L-induced NAb responses against H1N1 PR8, NC99, CA09, MI15, and WI22 viruses at week 8 after three immunizations. Summary of neutralization titers and statistical analysis. Error bars represent the difference between duplicate measurements at each concentration for each sample. $EC_{50}$ values were calculated using GraphPad Prism version 10.3.1. Data were analyzed using one-way ANOVA, followed by Tukey's multiple comparison post hoc test for each timepoint. For significance, ns (not significant), $*p < 0.05$, $**p < 0.01$, $***p < 0.001$, and $****p < 0.0001$.

Fig. S11

a Sera of individual mice immunized with H3N2 HK68 HA vaccines binding to H3N2 HK68 HA-N95L(1TD0) trimer

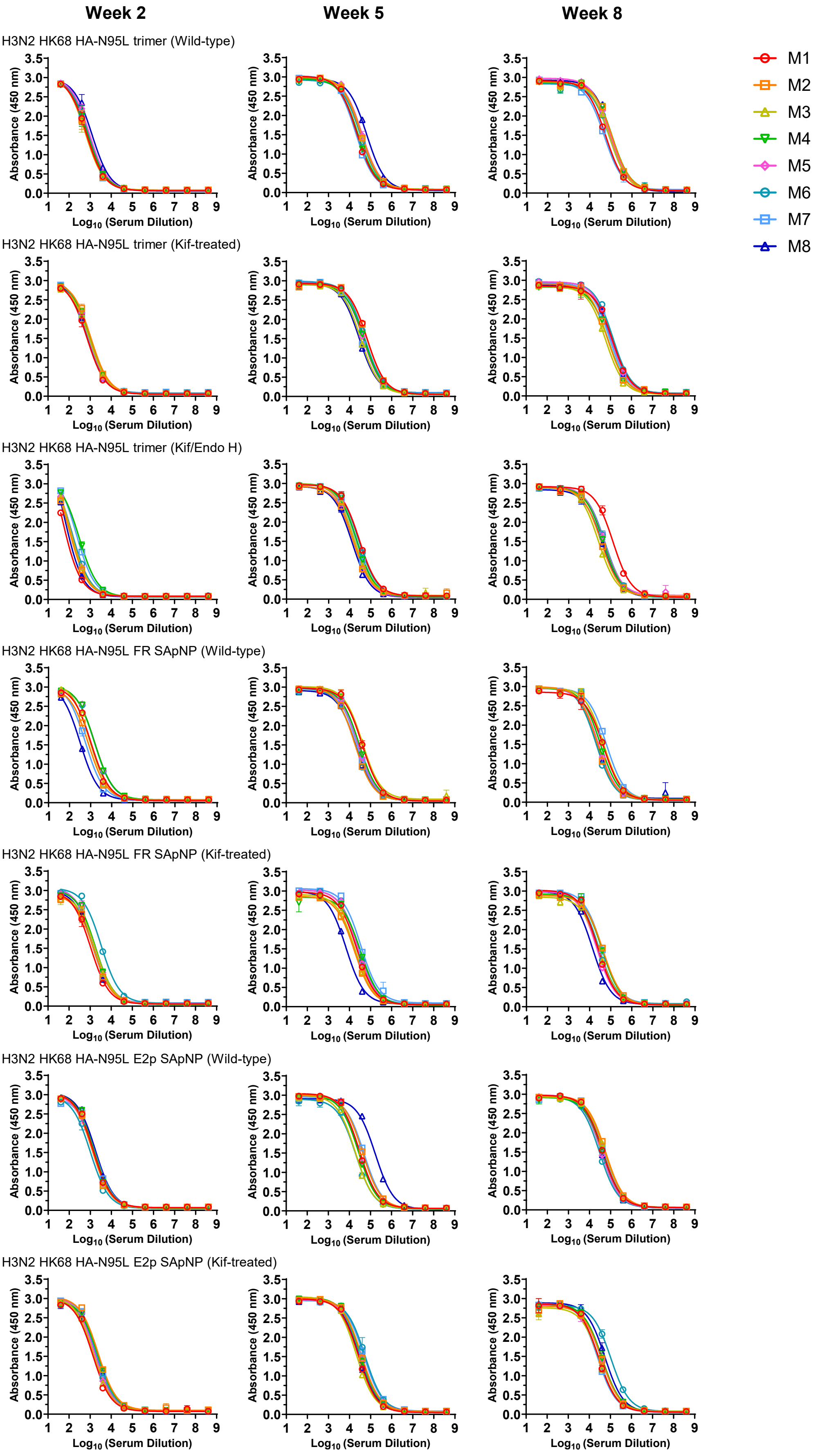

Fig. S11

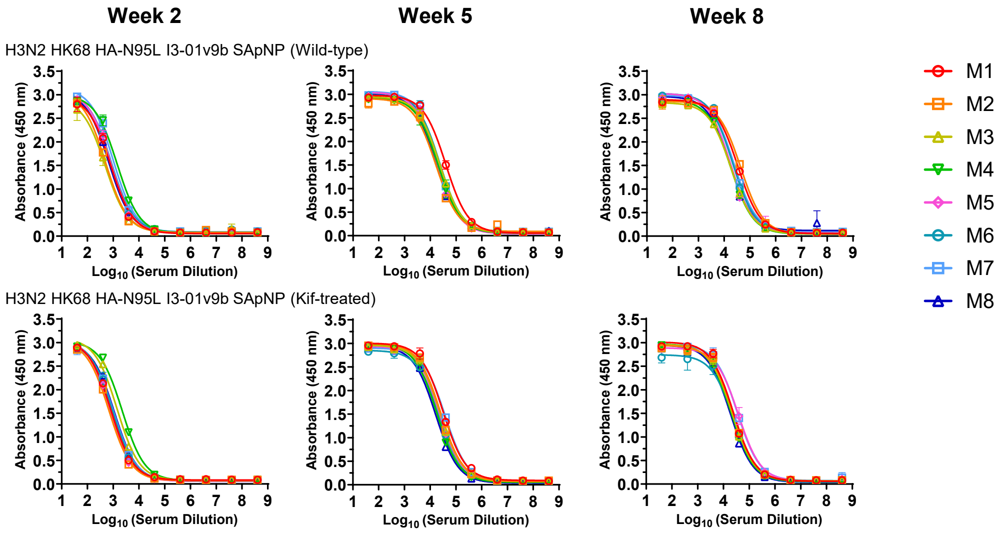

#### b Mouse serum ELISA $EC_{50}$ titers

**Week 2**

| Antigen | $EC_{50}$ titers (week 2) | | | | | | | | Geometric Mean |
|---|---|---|---|---|---|---|---|---|---|
| | M1 | M2 | M3 | M4 | M5 | M6 | M7 | M8 | |
| H3N2 HK68 HA-N95L trimer (Wild-type) | 674 | 625 | 588 | 765 | 924 | 830 | 656 | 1227 | 764.7 |
| H3N2 HK68 HA-N95L trimer (Kif-treated) | 707 | 1086 | 942 | 968 | 864 | 846 | 776 | 776 | 863.3 |
| H3N2 HK68 HA-N95L trimer (Kif/Endo H) | 47 | 98 | 109 | 301 | 110 | 138 | 222 | 62 | 115.3 |
| H3N2 HK68 HA-N95L FR SApNP (Wild-type) | 1068 | 789 | 920 | 1547 | 1611 | 1560 | 565 | 316 | 925.2 |
| H3N2 HK68 HA-N95L FR SApNP (Kif-treated) | 1062 | 1476 | 1779 | 1769 | 1689 | 3352 | 1407 | 1347 | 1642.3 |
| H3N2 HK68 HA-N95L E2p SApNP (Wild-type) | 1352 | 1232 | 1238 | 1489 | 1275 | 1006 | 1709 | 1754 | 1361.5 |
| H3N2 HK68 HA-N95L E2p SApNP (Kif-treated) | 1326 | 2419 | 1438 | 2361 | 1668 | 1872 | 2056 | 2191 | 1875.6 |
| H3N2 HK68 HA-N95L I3-01v9b SApNP (Wild-type) | 793 | 504 | 492 | 1433 | 788 | 828 | 1007 | 669 | 770.3 |
| H3N2 HK68 HA-N95L I3-01v9b SApNP (Kif-treated) | 814 | 691 | 1433 | 2253 | 756 | 943 | 1048 | 1099 | 1051.2 |

**Week 5**

| Antigen | $EC_{50}$ titers (week 5) | | | | | | | | Geometric Mean |
|---|---|---|---|---|---|---|---|---|---|
| | M1 | M2 | M3 | M4 | M5 | M6 | M7 | M8 | |
| H3N2 HK68 HA-N95L trimer (Wild-type) | 22274 | 34531 | 35286 | 25173 | 38852 | 33139 | 19558 | 65237 | 32082 |
| H3N2 HK68 HA-N95L trimer (Kif-treated) | 68285 | 68326 | 34048 | 48509 | 43378 | 51284 | 31217 | 28817 | 44516 |
| H3N2 HK68 HA-N95L trimer (Kif/Endo H) | 28865 | 14516 | 16430 | 21739 | 15227 | 27106 | 20247 | 12720 | 18845 |
| H3N2 HK68 HA-N95L FR SApNP (Wild-type) | 40636 | 18110 | 41592 | 28906 | 25830 | 18858 | 23428 | 20312 | 25941 |
| H3N2 HK68 HA-N95L FR SApNP (Kif-treated) | 22007 | 16950 | 18435 | 33563 | 24721 | 26179 | 34783 | 6845 | 20839 |
| H3N2 HK68 HA-N95L E2p SApNP (Wild-type) | 30288 | 40993 | 18979 | 27157 | 29327 | 19913 | 48236 | 170510 | 36388 |
| H3N2 HK68 HA-N95L E2p SApNP (Kif-treated) | 26543 | 36075 | 20821 | 32363 | 37312 | 54593 | 48026 | 27766 | 33920 |
| H3N2 HK68 HA-N95L I3-01v9b SApNP (Wild-type) | 39771 | 16483 | 23655 | 20384 | 18540 | 23034 | 21857 | 17103 | 21771 |
| H3N2 HK68 HA-N95L I3-01v9b SApNP (Kif-treated) | 31993 | 25479 | 24746 | 18285 | 23493 | 23087 | 37570 | 16372 | 24337 |

**Week 8**

| Antigen | $EC_{50}$ titers (week 8) | | | | | | | | Geometric Mean |
|---|---|---|---|---|---|---|---|---|---|
| | M1 | M2 | M3 | M4 | M5 | M6 | M7 | M8 | |
| H3N2 HK68 HA-N95L trimer (Wild-type) | 56490 | 112348 | 85351 | 115989 | 91333 | 86085 | 48648 | 110928 | 84767 |
| H3N2 HK68 HA-N95L trimer (Kif-treated) | 125660 | 73554 | 60624 | 80118 | 110390 | 131766 | 90716 | 112526 | 95056 |
| H3N2 HK68 HA-N95L trimer (Kif/Endo H) | 126902 | 34647 | 25925 | 44630 | 45542 | 51947 | 52165 | 39163 | 47189 |
| H3N2 HK68 HA-N95L FR SApNP (Wild-type) | 45112 | 23594 | 45620 | 32327 | 26430 | 20450 | 62749 | 21941 | 32243 |
| H3N2 HK68 HA-N95L FR SApNP (Kif-treated) | 24404 | 42901 | 28387 | 39255 | 21815 | 27218 | 37958 | 13595 | 27807 |
| H3N2 HK68 HA-N95L E2p SApNP (Wild-type) | 41988 | 57029 | 53673 | 41870 | 35579 | 29789 | 47331 | 36564 | 42101 |
| H3N2 HK68 HA-N95L E2p SApNP (Kif-treated) | 28123 | 38507 | 29874 | 39655 | 36866 | 105427 | 24741 | 55806 | 40247 |
| H3N2 HK68 HA-N95L I3-01v9b SApNP (Wild-type) | 34358 | 44026 | 18513 | 17794 | 17427 | 21541 | 24006 | 16445 | 22832 |
| H3N2 HK68 HA-N95L I3-01v9b SApNP (Kif-treated) | 23675 | 24379 | 21434 | 19922 | 37936 | 24485 | 34679 | 18679 | 24911 |

#### Statistical analysis

| One-way ANOVA with Tukey's multiple comparisons test (w2) | Statistics | Adjusted P Value |
|---|---|---|
| HA trimer-Wild-type vs. HA trimer-Kif-treated | ns | >0.9999 |
| HA trimer-Wild-type vs. HA trimer-Kif/Endo H | * | 0.0373 |
| HA trimer-Wild-type vs. HA FR SApNP-Wild-type | ns | 0.9177 |
| HA trimer-Wild-type vs. HA FR SApNP-Kif-treated | *** | 0.0003 |
| HA trimer-Wild-type vs. HA E2p SApNP-Wild-type | ns | 0.0768 |
| HA trimer-Wild-type vs. HA E2p SApNP-Kif-treated | **** | <0.0001 |
| HA trimer-Wild-type vs. HA I3-01v9b SApNP-Wild-type | ns | >0.9999 |
| HA trimer-Wild-type vs. HA I3-01v9b SApNP-Kif-treated | ns | 0.7107 |
| HA trimer-Kif-treated vs. HA trimer-Kif/Endo H | * | 0.0108 |
| HA trimer-Kif-treated vs. HA FR SApNP-Wild-type | ns | 0.9922 |
| HA trimer-Kif-treated vs. HA FR SApNP-Kif-treated | ** | 0.0013 |
| HA trimer-Kif-treated vs. HA E2p SApNP-Wild-type | ns | 0.2029 |
| HA trimer-Kif-treated vs. HA E2p SApNP-Kif-treated | **** | <0.0001 |
| HA trimer-Kif-treated vs. HA I3-01v9b SApNP-Wild-type | ns | >0.9999 |
| HA trimer-Kif-treated vs. HA I3-01v9b SApNP-Kif-treated | ns | 0.9207 |
| HA trimer-Kif/Endo H vs. HA FR SApNP-Wild-type | *** | 0.0006 |
| HA trimer-Kif/Endo H vs. HA FR SApNP-Kif-treated | **** | <0.0001 |
| HA trimer-Kif/Endo H vs. HA E2p SApNP-Wild-type | **** | <0.0001 |
| HA trimer-Kif/Endo H vs. HA E2p SApNP-Kif-treated | **** | <0.0001 |
| HA trimer-Kif/Endo H vs. HA I3-01v9b SApNP-Wild-type | * | 0.0251 |
| HA trimer-Kif/Endo H vs. HA I3-01v9b SApNP-Kif-treated | *** | 0.0001 |
| HA FR SApNP-Wild-type vs. HA FR SApNP-Kif-treated | * | 0.0217 |
| HA FR SApNP-Wild-type vs. HA E2p SApNP-Wild-type | ns | 0.7375 |
| HA FR SApNP-Wild-type vs. HA E2p SApNP-Kif-treated | ** | 0.0012 |
| HA FR SApNP-Wild-type vs. HA I3-01v9b SApNP-Wild-type | ns | 0.9562 |
| HA FR SApNP-Wild-type vs. HA I3-01v9b SApNP-Kif-treated | ns | >0.9999 |
| HA FR SApNP-Kif-treated vs. HA E2p SApNP-Wild-type | ns | 0.6791 |
| HA FR SApNP-Kif-treated vs. HA E2p SApNP-Kif-treated | ns | 0.9906 |
| HA FR SApNP-Kif-treated vs. HA I3-01v9b SApNP-Wild-type | *** | 0.0005 |
| HA FR SApNP-Kif-treated vs. HA I3-01v9b SApNP-Kif-treated | ns | 0.0679 |
| HA E2p SApNP-Wild-type vs. HA E2p SApNP-Kif-treated | ns | 0.1584 |
| HA E2p SApNP-Wild-type vs. HA I3-01v9b SApNP-Wild-type | ns | 0.1085 |
| HA E2p SApNP-Wild-type vs. HA I3-01v9b SApNP-Kif-treated | ns | 0.931 |
| HA E2p SApNP-Kif-treated vs. HA I3-01v9b SApNP-Wild-type | **** | <0.0001 |
| HA E2p SApNP-Kif-treated vs. HA I3-01v9b SApNP-Kif-treated | ** | 0.0047 |
| HA I3-01v9b SApNP-Wild-type vs. HA I3-01v9b SApNP-Kif-treated | ns | 0.7954 |

| One-way ANOVA with Tukey's multiple comparisons test (w5) | Statistics | Adjusted P Value |
|---|---|---|
| HA trimer-Wild-type vs. HA trimer-Kif-treated | ns | 0.9335 |
| HA trimer-Wild-type vs. HA trimer-Kif/Endo H | ns | 0.85 |
| HA trimer-Wild-type vs. HA FR SApNP-Wild-type | ns | 0.9983 |
| HA trimer-Wild-type vs. HA FR SApNP-Kif-treated | ns | 0.9616 |
| HA trimer-Wild-type vs. HA E2p SApNP-Wild-type | ns | 0.8827 |
| HA trimer-Wild-type vs. HA E2p SApNP-Kif-treated | ns | >0.9999 |
| HA trimer-Wild-type vs. HA I3-01v9b SApNP-Wild-type | ns | 0.9546 |
| HA trimer-Wild-type vs. HA I3-01v9b SApNP-Kif-treated | ns | 0.9899 |
| HA trimer-Kif-treated vs. HA trimer-Kif/Endo H | ns | 0.1413 |
| HA trimer-Kif-treated vs. HA FR SApNP-Wild-type | ns | 0.5473 |
| HA trimer-Kif-treated vs. HA FR SApNP-Kif-treated | ns | 0.2807 |
| HA trimer-Kif-treated vs. HA E2p SApNP-Wild-type | ns | >0.9999 |
| HA trimer-Kif-treated vs. HA E2p SApNP-Kif-treated | ns | 0.9621 |
| HA trimer-Kif-treated vs. HA I3-01v9b SApNP-Wild-type | ns | 0.2638 |
| HA trimer-Kif-treated vs. HA I3-01v9b SApNP-Kif-treated | ns | 0.4082 |
| HA trimer-Kif/Endo H vs. HA FR SApNP-Wild-type | ns | 0.9971 |
| HA trimer-Kif/Endo H vs. HA FR SApNP-Kif-treated | ns | >0.9999 |
| HA trimer-Kif/Endo H vs. HA E2p SApNP-Wild-type | ns | 0.1008 |
| HA trimer-Kif/Endo H vs. HA E2p SApNP-Kif-treated | ns | 0.7881 |
| HA trimer-Kif/Endo H vs. HA I3-01v9b SApNP-Wild-type | ns | >0.9999 |
| HA trimer-Kif/Endo H vs. HA I3-01v9b SApNP-Kif-treated | ns | 0.9997 |
| HA FR SApNP-Wild-type vs. HA FR SApNP-Kif-treated | ns | >0.9999 |
| HA FR SApNP-Wild-type vs. HA E2p SApNP-Wild-type | ns | 0.4496 |
| HA FR SApNP-Wild-type vs. HA E2p SApNP-Kif-treated | ns | 0.9949 |
| HA FR SApNP-Wild-type vs. HA I3-01v9b SApNP-Wild-type | ns | >0.9999 |
| HA FR SApNP-Wild-type vs. HA I3-01v9b SApNP-Kif-treated | ns | >0.9999 |
| HA FR SApNP-Kif-treated vs. HA E2p SApNP-Wild-type | ns | 0.212 |
| HA FR SApNP-Kif-treated vs. HA E2p SApNP-Kif-treated | ns | 0.9328 |
| HA FR SApNP-Kif-treated vs. HA I3-01v9b SApNP-Wild-type | ns | >0.9999 |
| HA FR SApNP-Kif-treated vs. HA I3-01v9b SApNP-Kif-treated | ns | >0.9999 |
| HA E2p SApNP-Wild-type vs. HA E2p SApNP-Kif-treated | ns | 0.9257 |
| HA E2p SApNP-Wild-type vs. HA I3-01v9b SApNP-Wild-type | ns | 0.198 |
| HA E2p SApNP-Wild-type vs. HA I3-01v9b SApNP-Kif-treated | ns | 0.3214 |
| HA E2p SApNP-Kif-treated vs. HA I3-01v9b SApNP-Wild-type | ns | 0.9226 |
| HA E2p SApNP-Kif-treated vs. HA I3-01v9b SApNP-Kif-treated | ns | 0.9781 |
| HA I3-01v9b SApNP-Wild-type vs. HA I3-01v9b SApNP-Kif-treated | ns | >0.9999 |

| One-way ANOVA with Tukey's multiple comparisons test (w8) | Statistics | Adjusted P Value |
|---|---|---|
| HA trimer-Wild-type vs. HA trimer-Kif-treated | ns | 0.9857 |
| HA trimer-Wild-type vs. HA trimer-Kif/Endo H | * | 0.0162 |
| HA trimer-Wild-type vs. HA FR SApNP-Wild-type | **** | <0.0001 |
| HA trimer-Wild-type vs. HA FR SApNP-Kif-treated | **** | <0.0001 |
| HA trimer-Wild-type vs. HA E2p SApNP-Wild-type | *** | 0.0007 |
| HA trimer-Wild-type vs. HA E2p SApNP-Kif-treated | ** | 0.0013 |
| HA trimer-Wild-type vs. HA I3-01v9b SApNP-Wild-type | **** | <0.0001 |
| HA trimer-Wild-type vs. HA I3-01v9b SApNP-Kif-treated | **** | <0.0001 |
| HA trimer-Kif-treated vs. HA trimer-Kif/Endo H | *** | 0.0007 |
| HA trimer-Kif-treated vs. HA FR SApNP-Wild-type | **** | <0.0001 |
| HA trimer-Kif-treated vs. HA FR SApNP-Kif-treated | **** | <0.0001 |
| HA trimer-Kif-treated vs. HA E2p SApNP-Wild-type | **** | <0.0001 |
| HA trimer-Kif-treated vs. HA E2p SApNP-Kif-treated | **** | <0.0001 |
| HA trimer-Kif-treated vs. HA I3-01v9b SApNP-Wild-type | **** | <0.0001 |
| HA trimer-Kif-treated vs. HA I3-01v9b SApNP-Kif-treated | **** | <0.0001 |
| HA trimer-Kif/Endo H vs. HA FR SApNP-Wild-type | ns | 0.6802 |
| HA trimer-Kif/Endo H vs. HA FR SApNP-Kif-treated | ns | 0.3338 |
| HA trimer-Kif/Endo H vs. HA E2p SApNP-Wild-type | ns | 0.9869 |
| HA trimer-Kif/Endo H vs. HA E2p SApNP-Kif-treated | ns | 0.997 |
| HA trimer-Kif/Endo H vs. HA I3-01v9b SApNP-Wild-type | ns | 0.1173 |
| HA trimer-Kif/Endo H vs. HA I3-01v9b SApNP-Kif-treated | ns | 0.16 |
| HA FR SApNP-Wild-type vs. HA FR SApNP-Kif-treated | ns | 0.9998 |
| HA FR SApNP-Wild-type vs. HA E2p SApNP-Wild-type | ns | 0.9955 |
| HA FR SApNP-Wild-type vs. HA E2p SApNP-Kif-treated | ns | 0.9825 |
| HA FR SApNP-Wild-type vs. HA I3-01v9b SApNP-Wild-type | ns | 0.9774 |
| HA FR SApNP-Wild-type vs. HA I3-01v9b SApNP-Kif-treated | ns | 0.9908 |
| HA FR SApNP-Kif-treated vs. HA E2p SApNP-Wild-type | ns | 0.9054 |
| HA FR SApNP-Kif-treated vs. HA E2p SApNP-Kif-treated | ns | 0.8223 |
| HA FR SApNP-Kif-treated vs. HA I3-01v9b SApNP-Wild-type | ns | 0.9998 |
| HA FR SApNP-Kif-treated vs. HA I3-01v9b SApNP-Kif-treated | ns | >0.9999 |
| HA E2p SApNP-Wild-type vs. HA E2p SApNP-Kif-treated | ns | >0.9999 |
| HA E2p SApNP-Wild-type vs. HA I3-01v9b SApNP-Wild-type | ns | 0.622 |
| HA E2p SApNP-Wild-type vs. HA I3-01v9b SApNP-Kif-treated | ns | 0.7128 |
| HA E2p SApNP-Kif-treated vs. HA I3-01v9b SApNP-Wild-type | ns | 0.4938 |
| HA E2p SApNP-Kif-treated vs. HA I3-01v9b SApNP-Kif-treated | ns | 0.5874 |
| HA I3-01v9b SApNP-Wild-type vs. HA I3-01v9b SApNP-Kif-treated | ns | >0.9999 |

Fig. S11

**C** **Week-8 sera of individual mice immunized with H3N2 HK68 HA vaccines binding to H1N1 CA09 HA-N95L(1TD0) trimer**

**Week 8**

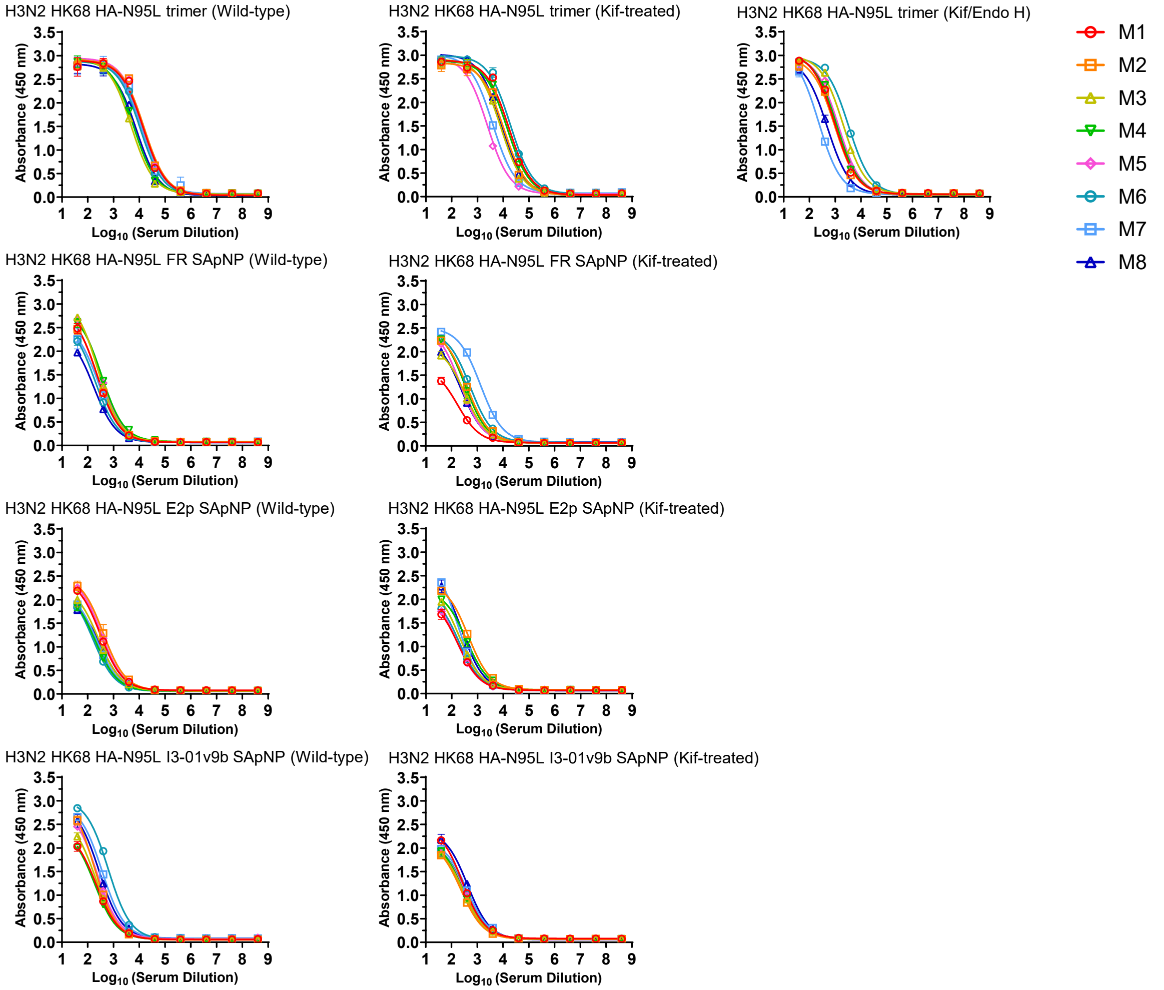

#### Mouse serum ELISA $EC_{50}$ titers

**Week 8**

| Antigen | $EC_{50}$ titers (week 8) | | | | | | | | Geometric Mean |
|---|---|---|---|---|---|---|---|---|---|
| | M1 | M2 | M3 | M4 | M5 | M6 | M7 | M8 | |
| H3N2 HK68 HA-N95L trimer (Wild-type) | 14232 | 15244 | 4872 | 6115 | 14531 | 10760 | 12122 | 7671 | 9923 |
| H3N2 HK68 HA-N95L trimer (Kif-treated) | 16492 | 10654 | 8010 | 14168 | 2428 | 18890 | 3919 | 8300 | 8594 |
| H3N2 HK68 HA-N95L trimer (Kif/Endo H) | 959 | 962 | 2042 | 1131 | 1319 | 3313 | 228 | 492 | 1006 |
| H3N2 HK68 HA-N95L FR SApNP (Wild-type) | 233 | 239 | 247 | 344 | 295 | 194 | 258 | 171 | 242 |
| H3N2 HK68 HA-N95L FR SApNP (Kif-treated) | 172 | 404 | 338 | 336 | 221 | 532 | 1338 | 241 | 363 |
| H3N2 HK68 HA-N95L E2p SApNP (Wild-type) | 306 | 401 | 257 | 195 | 337 | 158 | 223 | 321 | 264 |
| H3N2 HK68 HA-N95L E2p SApNP (Kif-treated) | 174 | 439 | 201 | 374 | 150 | 183 | 161 | 245 | 223 |
| H3N2 HK68 HA-N95L I3-01v9b SApNP (Wild-type) | 212 | 179 | 198 | 186 | 227 | 627 | 361 | 279 | 258 |
| H3N2 HK68 HA-N95L I3-01v9b SApNP (Kif-treated) | 271 | 236 | 282 | 272 | 326 | 351 | 402 | 422 | 314 |

#### Statistical analysis

| One-way ANOVA with Tukey's multiple comparisons test (w8) | Statistics | Adjusted P Value |
|---|---|---|
| HA trimer-Wild-type vs. HA trimer-Kif-treated | ns | >0.9999 |
| HA trimer-Wild-type vs. HA trimer-Kif/Endo H | **** | <0.0001 |
| HA trimer-Wild-type vs. HA FR SApNP-Wild-type | **** | <0.0001 |
| HA trimer-Wild-type vs. HA FR SApNP-Kif-treated | **** | <0.0001 |
| HA trimer-Wild-type vs. HA E2p SApNP-Wild-type | **** | <0.0001 |
| HA trimer-Wild-type vs. HA E2p SApNP-Kif-treated | **** | <0.0001 |
| HA trimer-Wild-type vs. HA I3-01v9b SApNP-Wild-type | **** | <0.0001 |
| HA trimer-Wild-type vs. HA I3-01v9b SApNP-Kif-treated | **** | <0.0001 |
| HA trimer-Kif-treated vs. HA trimer-Kif/Endo H | **** | <0.0001 |
| HA trimer-Kif-treated vs. HA FR SApNP-Wild-type | **** | <0.0001 |
| HA trimer-Kif-treated vs. HA FR SApNP-Kif-treated | **** | <0.0001 |
| HA trimer-Kif-treated vs. HA E2p SApNP-Wild-type | **** | <0.0001 |
| HA trimer-Kif-treated vs. HA E2p SApNP-Kif-treated | **** | <0.0001 |
| HA trimer-Kif-treated vs. HA I3-01v9b SApNP-Wild-type | **** | <0.0001 |
| HA trimer-Kif-treated vs. HA I3-01v9b SApNP-Kif-treated | **** | <0.0001 |
| HA trimer-Kif/Endo H vs. HA FR SApNP-Wild-type | ns | 0.9931 |
| HA trimer-Kif/Endo H vs. HA FR SApNP-Kif-treated | ns | 0.9984 |
| HA trimer-Kif/Endo H vs. HA E2p SApNP-Wild-type | ns | 0.9942 |
| HA trimer-Kif/Endo H vs. HA E2p SApNP-Kif-treated | ns | 0.9928 |
| HA trimer-Kif/Endo H vs. HA I3-01v9b SApNP-Wild-type | ns | 0.9945 |
| HA trimer-Kif/Endo H vs. HA I3-01v9b SApNP-Kif-treated | ns | 0.9957 |
| HA FR SApNP-Wild-type vs. HA FR SApNP-Kif-treated | ns | >0.9999 |
| HA FR SApNP-Wild-type vs. HA E2p SApNP-Wild-type | ns | >0.9999 |
| HA FR SApNP-Wild-type vs. HA E2p SApNP-Kif-treated | ns | >0.9999 |
| HA FR SApNP-Wild-type vs. HA I3-01v9b SApNP-Wild-type | ns | >0.9999 |
| HA FR SApNP-Wild-type vs. HA I3-01v9b SApNP-Kif-treated | ns | >0.9999 |
| HA FR SApNP-Kif-treated vs. HA E2p SApNP-Wild-type | ns | >0.9999 |
| HA FR SApNP-Kif-treated vs. HA E2p SApNP-Kif-treated | ns | >0.9999 |
| HA FR SApNP-Kif-treated vs. HA I3-01v9b SApNP-Wild-type | ns | >0.9999 |
| HA FR SApNP-Kif-treated vs. HA I3-01v9b SApNP-Kif-treated | ns | >0.9999 |
| HA E2p SApNP-Wild-type vs. HA E2p SApNP-Kif-treated | ns | >0.9999 |
| HA E2p SApNP-Wild-type vs. HA I3-01v9b SApNP-Wild-type | ns | >0.9999 |
| HA E2p SApNP-Wild-type vs. HA I3-01v9b SApNP-Kif-treated | ns | >0.9999 |
| HA E2p SApNP-Kif-treated vs. HA I3-01v9b SApNP-Wild-type | ns | >0.9999 |
| HA E2p SApNP-Kif-treated vs. HA I3-01v9b SApNP-Kif-treated | ns | >0.9999 |
| HA I3-01v9b SApNP-Wild-type vs. HA I3-01v9b SApNP-Kif-treated | ns | >0.9999 |

### Fig. S11

d

#### H3N2 HK68 HA-induced sera HAI against vaccine matched H3N2 HK68 virus

##### Week 2

| Antigen | HAI Titer $(Log_2)$ (week 2) | | | | | | | |
|---|---|---|---|---|---|---|---|---|
| | M1 | M2 | M3 | M4 | M5 | M6 | M7 | M8 |
| H3N2 HK68 HA-N95L trimer (Wild-type) | 20 | 20 | 20 | 20 | 20 | 20 | 20 | 20 |
| H3N2 HK68 HA-N95L trimer (Wild-type) | 20 | 20 | 20 | 20 | 20 | 20 | 20 | 20 |
| H3N2 HK68 HA-N95L trimer (Kif-treated) | 20 | 20 | 20 | 20 | 20 | 20 | 20 | 20 |
| H3N2 HK68 HA-N95L trimer (Kif-treated) | 20 | 20 | 20 | 20 | 20 | 20 | 20 | 20 |
| H3N2 HK68 HA-N95L trimer (Kif/Endo H) | 20 | 20 | 20 | 20 | 20 | 20 | 20 | 20 |
| H3N2 HK68 HA-N95L trimer (Kif/Endo H) | 20 | 20 | 20 | 20 | 20 | 20 | 20 | 20 |
| H3N2 HK68 HA-N95L FR SApNP (Wild-type) | 20 | 20 | 20 | 20 | 20 | 20 | 20 | 20 |
| H3N2 HK68 HA-N95L FR SApNP (Wild-type) | 20 | 20 | 20 | 20 | 20 | 20 | 20 | 20 |
| H3N2 HK68 HA-N95L FR SApNP (Kif-treated) | 20 | 40 | 20 | 20 | 20 | 80 | 20 | 40 |
| H3N2 HK68 HA-N95L FR SApNP (Kif-treated) | 20 | 40 | 20 | 20 | 20 | 80 | 20 | 40 |
| H3N2 HK68 HA-N95L E2p SApNP (Wild-type) | 20 | 20 | 20 | 20 | 20 | 20 | 20 | 20 |
| H3N2 HK68 HA-N95L E2p SApNP (Wild-type) | 20 | 20 | 20 | 20 | 20 | 20 | 20 | 20 |
| H3N2 HK68 HA-N95L E2p SApNP (Kif-treated) | 20 | 20 | 20 | 40 | 20 | 20 | 40 | 20 |
| H3N2 HK68 HA-N95L E2p SApNP (Kif-treated) | 20 | 20 | 20 | 40 | 20 | 20 | 40 | 20 |
| H3N2 HK68 HA-N95L I3-01v9b SApNP (Wild-type) | 20 | 20 | 20 | 40 | 20 | 20 | 20 | 20 |
| H3N2 HK68 HA-N95L I3-01v9b SApNP (Wild-type) | 20 | 20 | 20 | 40 | 20 | 20 | 20 | 20 |
| H3N2 HK68 HA-N95L I3-01v9b SApNP (Kif-treated) | 20 | 20 | 20 | 40 | 20 | 20 | 20 | 40 |
| H3N2 HK68 HA-N95L I3-01v9b SApNP (Kif-treated) | 20 | 20 | 20 | 40 | 20 | 20 | 20 | 40 |

##### Week 5

| Antigen | HAI Titer $(Log_2)$ (week 5) | | | | | | | |
|---|---|---|---|---|---|---|---|---|
| | M1 | M2 | M3 | M4 | M5 | M6 | M7 | M8 |
| H3N2 HK68 HA-N95L trimer (Wild-type) | 320 | 320 | 20 | 20 | 80 | 160 | 40 | 160 |
| H3N2 HK68 HA-N95L trimer (Wild-type) | 320 | 160 | 20 | 20 | 80 | 160 | 40 | 160 |
| H3N2 HK68 HA-N95L trimer (Kif-treated) | 640 | 640 | 80 | 320 | 320 | 320 | 160 | 160 |
| H3N2 HK68 HA-N95L trimer (Kif-treated) | 640 | 640 | 80 | 320 | 320 | 320 | 160 | 160 |
| H3N2 HK68 HA-N95L trimer (Kif/Endo H) | 160 | 20 | 20 | 40 | 20 | 80 | 160 | 20 |
| H3N2 HK68 HA-N95L trimer (Kif/Endo H) | 160 | 20 | 20 | 40 | 20 | 80 | 160 | 20 |
| H3N2 HK68 HA-N95L FR SApNP (Wild-type) | 320 | 160 | 320 | 320 | 320 | 160 | 160 | 160 |
| H3N2 HK68 HA-N95L FR SApNP (Wild-type) | 320 | 160 | 320 | 320 | 320 | 160 | 160 | 160 |
| H3N2 HK68 HA-N95L FR SApNP (Kif-treated) | 320 | 320 | 320 | 320 | 320 | 320 | 160 | 80 |
| H3N2 HK68 HA-N95L FR SApNP (Kif-treated) | 320 | 320 | 320 | 320 | 320 | 320 | 160 | 80 |
| H3N2 HK68 HA-N95L E2p SApNP (Wild-type) | 320 | 320 | 160 | 160 | 320 | 160 | 640 | 1280 |
| H3N2 HK68 HA-N95L E2p SApNP (Wild-type) | 320 | 320 | 160 | 160 | 320 | 160 | 320 | 1280 |
| H3N2 HK68 HA-N95L E2p SApNP (Kif-treated) | 320 | 320 | 160 | 640 | 320 | 320 | 320 | 320 |
| H3N2 HK68 HA-N95L E2p SApNP (Kif-treated) | 320 | 320 | 160 | 640 | 320 | 320 | 320 | 320 |
| H3N2 HK68 HA-N95L I3-01v9b SApNP (Wild-type) | 640 | 320 | 640 | 640 | 320 | 320 | 320 | 320 |
| H3N2 HK68 HA-N95L I3-01v9b SApNP (Wild-type) | 640 | 320 | 640 | 640 | 320 | 320 | 320 | 320 |
| H3N2 HK68 HA-N95L I3-01v9b SApNP (Kif-treated) | 320 | 320 | 320 | 320 | 320 | 320 | 320 | 320 |
| H3N2 HK68 HA-N95L I3-01v9b SApNP (Kif-treated) | 320 | 320 | 320 | 320 | 320 | 320 | 320 | 320 |

##### Week 8

| Antigen | HAI Titer $(Log_2)$ (week 8) | | | | | | | |
|---|---|---|---|---|---|---|---|---|
| | M1 | M2 | M3 | M4 | M5 | M6 | M7 | M8 |
| H3N2 HK68 HA-N95L trimer (Wild-type) | 160 | 640 | 160 | 20 | 320 | 320 | 160 | 640 |
| H3N2 HK68 HA-N95L trimer (Wild-type) | 160 | 640 | 160 | 20 | 320 | 320 | 160 | 640 |
| H3N2 HK68 HA-N95L trimer (Kif-treated) | 1280 | 1280 | 320 | 640 | 1280 | 1280 | 320 | 1280 |
| H3N2 HK68 HA-N95L trimer (Kif-treated) | 1280 | 1280 | 320 | 640 | 640 | 1280 | 320 | 1280 |
| H3N2 HK68 HA-N95L trimer (Kif/Endo H) | 1280 | 40 | 80 | 160 | 160 | 640 | 320 | 40 |
| H3N2 HK68 HA-N95L trimer (Kif/Endo H) | 1280 | 40 | 80 | 160 | 160 | 640 | 320 | 40 |
| H3N2 HK68 HA-N95L FR SApNP (Wild-type) | 640 | 640 | 640 | 640 | 640 | 640 | 640 | 640 |
| H3N2 HK68 HA-N95L FR SApNP (Wild-type) | 640 | 640 | 640 | 640 | 320 | 640 | 640 | 640 |
| H3N2 HK68 HA-N95L FR SApNP (Kif-treated) | 640 | 640 | 640 | 640 | 320 | 640 | 640 | 160 |
| H3N2 HK68 HA-N95L FR SApNP (Kif-treated) | 640 | 1280 | 640 | 640 | 320 | 640 | 640 | 160 |
| H3N2 HK68 HA-N95L E2p SApNP (Wild-type) | 640 | 640 | 320 | 320 | 640 | 320 | 640 | 640 |
| H3N2 HK68 HA-N95L E2p SApNP (Wild-type) | 640 | 640 | 320 | 320 | 640 | 320 | 640 | 640 |
| H3N2 HK68 HA-N95L E2p SApNP (Kif-treated) | 320 | 640 | 640 | 1280 | 640 | 640 | 320 | 1280 |
| H3N2 HK68 HA-N95L E2p SApNP (Kif-treated) | 320 | 640 | 640 | 1280 | 640 | 640 | 320 | 1280 |
| H3N2 HK68 HA-N95L I3-01v9b SApNP (Wild-type) | 640 | 1280 | 1280 | 640 | 320 | 640 | 640 | 320 |
| H3N2 HK68 HA-N95L I3-01v9b SApNP (Wild-type) | 640 | 1280 | 1280 | 640 | 320 | 640 | 640 | 320 |
| H3N2 HK68 HA-N95L I3-01v9b SApNP (Kif-treated) | 320 | 640 | 640 | 640 | 640 | 640 | 1280 | 640 |
| H3N2 HK68 HA-N95L I3-01v9b SApNP (Kif-treated) | 320 | 640 | 640 | 640 | 640 | 640 | 1280 | 640 |

#### Statistical analysis

| One-way ANOVA with Tukey's multiple comparisons test (w2) | Statistics | Adjusted P Value |
|---|---|---|
| HA trimer-Wild-type vs. HA trimer-Kif-treated | ns | >0.9999 |
| HA trimer-Wild-type vs. HA trimer-Kif/Endo H | ns | >0.9999 |
| HA trimer-Wild-type vs. HA FR SApNP-Wild-type | ns | >0.9999 |
| HA trimer-Wild-type vs. HA FR SApNP-Kif-treated | ns | 0.1097 |
| HA trimer-Wild-type vs. HA E2p SApNP-Wild-type | ns | >0.9999 |
| HA trimer-Wild-type vs. HA E2p SApNP-Kif-treated | ns | 0.9623 |
| HA trimer-Wild-type vs. HA I3-01v9b SApNP-Wild-type | ns | 0.9997 |
| HA trimer-Wild-type vs. HA I3-01v9b SApNP-Kif-treated | ns | 0.9623 |
| HA trimer-Kif-treated vs. HA trimer-Kif/Endo H | ns | >0.9999 |
| HA trimer-Kif-treated vs. HA FR SApNP-Wild-type | ns | >0.9999 |
| HA trimer-Kif-treated vs. HA FR SApNP-Kif-treated | ns | 0.1097 |
| HA trimer-Kif-treated vs. HA E2p SApNP-Wild-type | ns | >0.9999 |
| HA trimer-Kif-treated vs. HA E2p SApNP-Kif-treated | ns | 0.9623 |
| HA trimer-Kif-treated vs. HA I3-01v9b SApNP-Wild-type | ns | 0.9997 |
| HA trimer-Kif-treated vs. HA I3-01v9b SApNP-Kif-treated | ns | 0.9623 |
| HA trimer-Kif/Endo H vs. HA FR SApNP-Wild-type | ns | >0.9999 |
| HA trimer-Kif/Endo H vs. HA FR SApNP-Kif-treated | ns | 0.1097 |
| HA trimer-Kif/Endo H vs. HA E2p SApNP-Wild-type | ns | >0.9999 |
| HA trimer-Kif/Endo H vs. HA E2p SApNP-Kif-treated | ns | 0.9623 |
| HA trimer-Kif/Endo H vs. HA I3-01v9b SApNP-Wild-type | ns | 0.9997 |
| HA trimer-Kif/Endo H vs. HA I3-01v9b SApNP-Kif-treated | ns | 0.9623 |
| HA FR SApNP-Wild-type vs. HA FR SApNP-Kif-treated | ns | 0.1097 |
| HA FR SApNP-Wild-type vs. HA E2p SApNP-Wild-type | ns | >0.9999 |
| HA FR SApNP-Wild-type vs. HA E2p SApNP-Kif-treated | ns | 0.9623 |
| HA FR SApNP-Wild-type vs. HA I3-01v9b SApNP-Wild-type | ns | 0.9997 |
| HA FR SApNP-Wild-type vs. HA I3-01v9b SApNP-Kif-treated | ns | 0.9623 |
| HA FR SApNP-Kif-treated vs. HA E2p SApNP-Wild-type | ns | 0.1097 |
| HA FR SApNP-Kif-treated vs. HA E2p SApNP-Kif-treated | ns | 0.7219 |
| HA FR SApNP-Kif-treated vs. HA I3-01v9b SApNP-Wild-type | ns | 0.3489 |
| HA FR SApNP-Kif-treated vs. HA I3-01v9b SApNP-Kif-treated | ns | 0.7219 |
| HA E2p SApNP-Wild-type vs. HA E2p SApNP-Kif-treated | ns | 0.9623 |
| HA E2p SApNP-Wild-type vs. HA I3-01v9b SApNP-Wild-type | ns | 0.9997 |
| HA E2p SApNP-Wild-type vs. HA I3-01v9b SApNP-Kif-treated | ns | 0.9623 |
| HA E2p SApNP-Kif-treated vs. HA I3-01v9b SApNP-Wild-type | ns | 0.9997 |
| HA E2p SApNP-Kif-treated vs. HA I3-01v9b SApNP-Kif-treated | ns | >0.9999 |
| HA I3-01v9b SApNP-Wild-type vs. HA I3-01v9b SApNP-Kif-treated | ns | 0.9997 |

| One-way ANOVA with Tukey's multiple comparisons test (w5) | Statistics | Adjusted P Value |
|---|---|---|
| HA trimer-Wild-type vs. HA trimer-Kif-treated | ns | 0.3305 |
| HA trimer-Wild-type vs. HA trimer-Kif/Endo H | ns | 0.9975 |
| HA trimer-Wild-type vs. HA FR SApNP-Wild-type | ns | 0.9302 |
| HA trimer-Wild-type vs. HA FR SApNP-Kif-treated | ns | 0.7772 |
| HA trimer-Wild-type vs. HA E2p SApNP-Wild-type | ns | 0.0558 |
| HA trimer-Wild-type vs. HA E2p SApNP-Kif-treated | ns | 0.2687 |
| HA trimer-Wild-type vs. HA I3-01v9b SApNP-Wild-type | * | 0.0152 |
| HA trimer-Wild-type vs. HA I3-01v9b SApNP-Kif-treated | ns | 0.3995 |
| HA trimer-Kif-treated vs. HA trimer-Kif/Endo H | ns | 0.0649 |
| HA trimer-Kif-treated vs. HA FR SApNP-Wild-type | ns | 0.9782 |
| HA trimer-Kif-treated vs. HA FR SApNP-Kif-treated | ns | 0.9986 |
| HA trimer-Kif-treated vs. HA E2p SApNP-Wild-type | ns | 0.9958 |
| HA trimer-Kif-treated vs. HA E2p SApNP-Kif-treated | ns | >0.9999 |
| HA trimer-Kif-treated vs. HA I3-01v9b SApNP-Wild-type | ns | 0.9302 |
| HA trimer-Kif-treated vs. HA I3-01v9b SApNP-Kif-treated | ns | >0.9999 |
| HA trimer-Kif/Endo H vs. HA FR SApNP-Wild-type | ns | 0.5128 |
| HA trimer-Kif/Endo H vs. HA FR SApNP-Kif-treated | ns | 0.2987 |
| HA trimer-Kif/Endo H vs. HA E2p SApNP-Wild-type | ** | 0.0062 |
| HA trimer-Kif/Endo H vs. HA E2p SApNP-Kif-treated | * | 0.0479 |
| HA trimer-Kif/Endo H vs. HA I3-01v9b SApNP-Wild-type | ** | 0.0013 |
| HA trimer-Kif/Endo H vs. HA I3-01v9b SApNP-Kif-treated | ns | 0.0868 |
| HA FR SApNP-Wild-type vs. HA FR SApNP-Kif-treated | ns | >0.9999 |
| HA FR SApNP-Wild-type vs. HA E2p SApNP-Wild-type | ns | 0.6307 |
| HA FR SApNP-Wild-type vs. HA E2p SApNP-Kif-treated | ns | 0.9591 |
| HA FR SApNP-Wild-type vs. HA I3-01v9b SApNP-Wild-type | ns | 0.3305 |
| HA FR SApNP-Wild-type vs. HA I3-01v9b SApNP-Kif-treated | ns | 0.9897 |
| HA FR SApNP-Kif-treated vs. HA E2p SApNP-Wild-type | ns | 0.839 |
| HA FR SApNP-Kif-treated vs. HA E2p SApNP-Kif-treated | ns | 0.9958 |
| HA FR SApNP-Kif-treated vs. HA I3-01v9b SApNP-Wild-type | ns | 0.5521 |
| HA FR SApNP-Kif-treated vs. HA I3-01v9b SApNP-Kif-treated | ns | 0.9996 |
| HA E2p SApNP-Wild-type vs. HA E2p SApNP-Kif-treated | ns | 0.9986 |
| HA E2p SApNP-Wild-type vs. HA I3-01v9b SApNP-Wild-type | ns | >0.9999 |
| HA E2p SApNP-Wild-type vs. HA I3-01v9b SApNP-Kif-treated | ns | 0.9897 |
| HA E2p SApNP-Kif-treated vs. HA I3-01v9b SApNP-Wild-type | ns | 0.9591 |
| HA E2p SApNP-Kif-treated vs. HA I3-01v9b SApNP-Kif-treated | ns | >0.9999 |
| HA I3-01v9b SApNP-Wild-type vs. HA I3-01v9b SApNP-Kif-treated | ns | 0.8903 |

| One-way ANOVA with Tukey's multiple comparisons test (w8) | Statistics | Adjusted P Value |
|---|---|---|
| HA trimer-Wild-type vs. HA trimer-Kif-treated | ** | 0.0052 |
| HA trimer-Wild-type vs. HA trimer-Kif/Endo H | ns | >0.9999 |
| HA trimer-Wild-type vs. HA FR SApNP-Wild-type | ns | 0.5151 |
| HA trimer-Wild-type vs. HA FR SApNP-Kif-treated | ns | 0.6868 |
| HA trimer-Wild-type vs. HA E2p SApNP-Wild-type | ns | 0.8918 |
| HA trimer-Wild-type vs. HA E2p SApNP-Kif-treated | ns | 0.17 |
| HA trimer-Wild-type vs. HA I3-01v9b SApNP-Wild-type | ns | 0.17 |
| HA trimer-Wild-type vs. HA I3-01v9b SApNP-Kif-treated | ns | 0.2821 |
| HA trimer-Kif-treated vs. HA trimer-Kif/Endo H | * | 0.011 |
| HA trimer-Kif-treated vs. HA FR SApNP-Wild-type | ns | 0.5909 |
| HA trimer-Kif-treated vs. HA FR SApNP-Kif-treated | ns | 0.4207 |
| HA trimer-Kif-treated vs. HA E2p SApNP-Wild-type | ns | 0.2143 |
| HA trimer-Kif-treated vs. HA E2p SApNP-Kif-treated | ns | 0.9301 |
| HA trimer-Kif-treated vs. HA I3-01v9b SApNP-Wild-type | ns | 0.9301 |
| HA trimer-Kif-treated vs. HA I3-01v9b SApNP-Kif-treated | ns | 0.8271 |
| HA trimer-Kif/Endo H vs. HA FR SApNP-Wild-type | ns | 0.6764 |
| HA trimer-Kif/Endo H vs. HA FR SApNP-Kif-treated | ns | 0.8271 |
| HA trimer-Kif/Endo H vs. HA E2p SApNP-Wild-type | ns | 0.9613 |
| HA trimer-Kif/Endo H vs. HA E2p SApNP-Kif-treated | ns | 0.274 |
| HA trimer-Kif/Endo H vs. HA I3-01v9b SApNP-Wild-type | ns | 0.274 |
| HA trimer-Kif/Endo H vs. HA I3-01v9b SApNP-Kif-treated | ns | 0.4207 |
| HA FR SApNP-Wild-type vs. HA FR SApNP-Kif-treated | ns | >0.9999 |
| HA FR SApNP-Wild-type vs. HA E2p SApNP-Wild-type | ns | 0.9992 |
| HA FR SApNP-Wild-type vs. HA E2p SApNP-Kif-treated | ns | 0.9992 |
| HA FR SApNP-Wild-type vs. HA I3-01v9b SApNP-Wild-type | ns | 0.9992 |
| HA FR SApNP-Wild-type vs. HA I3-01v9b SApNP-Kif-treated | ns | >0.9999 |
| HA FR SApNP-Kif-treated vs. HA E2p SApNP-Wild-type | ns | >0.9999 |
| HA FR SApNP-Kif-treated vs. HA E2p SApNP-Kif-treated | ns | 0.992 |
| HA FR SApNP-Kif-treated vs. HA I3-01v9b SApNP-Wild-type | ns | 0.992 |
| HA FR SApNP-Kif-treated vs. HA I3-01v9b SApNP-Kif-treated | ns | 0.9992 |
| HA E2p SApNP-Wild-type vs. HA E2p SApNP-Kif-treated | ns | 0.9301 |
| HA E2p SApNP-Wild-type vs. HA I3-01v9b SApNP-Wild-type | ns | 0.9301 |
| HA E2p SApNP-Wild-type vs. HA I3-01v9b SApNP-Kif-treated | ns | 0.981 |
| HA E2p SApNP-Kif-treated vs. HA I3-01v9b SApNP-Wild-type | ns | >0.9999 |
| HA E2p SApNP-Kif-treated vs. HA I3-01v9b SApNP-Kif-treated | ns | >0.9999 |
| HA I3-01v9b SApNP-Wild-type vs. HA I3-01v9b SApNP-Kif-treated | ns | >0.9999 |

### Fig. S11

#### e H3N2 HK68 HA-induced sera HAI (at week 8) against non-matched H3N2 BR07 and HK14 viruses

**A/BR/10/2007**

| Antigen | HAI Titer ($Log_2$) (week 8) | | | | | | | |
|---|---|---|---|---|---|---|---|---|
| | M1 | M2 | M3 | M4 | M5 | M6 | M7 | M8 |
| H3N2 HK68 HA-N95L trimer (Wild-type) | 20 | 20 | 20 | 20 | 20 | 20 | 20 | 20 |
| H3N2 HK68 HA-N95L trimer (Wild-type) | 20 | 20 | 20 | 20 | 20 | 20 | 20 | 20 |
| H3N2 HK68 HA-N95L trimer (Kif-treated) | 20 | 20 | 20 | 20 | 20 | 20 | 20 | 20 |
| H3N2 HK68 HA-N95L trimer (Kif-treated) | 20 | 20 | 20 | 20 | 20 | 20 | 20 | 20 |
| H3N2 HK68 HA-N95L trimer (Kif/Endo H) | 20 | 20 | 20 | 20 | 20 | 20 | 20 | 20 |
| H3N2 HK68 HA-N95L trimer (Kif/Endo H) | 20 | 20 | 20 | 20 | 20 | 20 | 20 | 20 |
| H3N2 HK68 HA-N95L FR SApNP (Wild-type) | 20 | 20 | 20 | 20 | 20 | 20 | 20 | 20 |
| H3N2 HK68 HA-N95L FR SApNP (Wild-type) | 20 | 20 | 20 | 20 | 20 | 20 | 20 | 20 |
| H3N2 HK68 HA-N95L FR SApNP (Kif-treated) | 20 | 20 | 20 | 20 | 20 | 20 | 20 | 20 |
| H3N2 HK68 HA-N95L FR SApNP (Kif-treated) | 20 | 20 | 20 | 20 | 20 | 20 | 20 | 20 |
| H3N2 HK68 HA-N95L E2p SApNP (Wild-type) | 20 | 20 | 20 | 20 | 20 | 20 | 20 | 20 |
| H3N2 HK68 HA-N95L E2p SApNP (Wild-type) | 20 | 20 | 20 | 20 | 20 | 20 | 20 | 20 |
| H3N2 HK68 HA-N95L E2p SApNP (Kif-treated) | 20 | 20 | 20 | 20 | 20 | 20 | 20 | 20 |
| H3N2 HK68 HA-N95L E2p SApNP (Kif-treated) | 20 | 20 | 20 | 20 | 20 | 20 | 20 | 20 |
| H3N2 HK68 HA-N95L I3-01v9b SApNP (Wild-type) | 20 | 20 | 20 | 20 | 20 | 20 | 20 | 20 |
| H3N2 HK68 HA-N95L I3-01v9b SApNP (Wild-type) | 20 | 20 | 20 | 20 | 20 | 20 | 20 | 20 |
| H3N2 HK68 HA-N95L I3-01v9b SApNP (Kif-treated) | 20 | 20 | 20 | 20 | 20 | 20 | 20 | 20 |
| H3N2 HK68 HA-N95L I3-01v9b SApNP (Kif-treated) | 20 | 20 | 20 | 20 | 20 | 20 | 20 | 20 |

**A/HK/4801/2014**

| Antigen | HAI Titer ($Log_2$) (week 8) | | | | | | | |
|---|---|---|---|---|---|---|---|---|
| | M1 | M2 | M3 | M4 | M5 | M6 | M7 | M8 |
| H3N2 HK68 HA-N95L trimer (Wild-type) | 20 | 20 | 20 | 20 | 20 | 20 | 20 | 20 |
| H3N2 HK68 HA-N95L trimer (Wild-type) | 20 | 20 | 20 | 20 | 20 | 20 | 20 | 20 |
| H3N2 HK68 HA-N95L trimer (Kif-treated) | 20 | 20 | 20 | 20 | 20 | 20 | 20 | 20 |
| H3N2 HK68 HA-N95L trimer (Kif-treated) | 20 | 20 | 20 | 20 | 20 | 20 | 20 | 20 |
| H3N2 HK68 HA-N95L trimer (Kif/Endo H) | 20 | 20 | 20 | 20 | 20 | 20 | 20 | 20 |
| H3N2 HK68 HA-N95L trimer (Kif/Endo H) | 20 | 20 | 20 | 20 | 20 | 20 | 20 | 20 |
| H3N2 HK68 HA-N95L FR SApNP (Wild-type) | 20 | 20 | 20 | 20 | 20 | 20 | 20 | 20 |
| H3N2 HK68 HA-N95L FR SApNP (Wild-type) | 20 | 20 | 20 | 20 | 20 | 20 | 20 | 20 |
| H3N2 HK68 HA-N95L FR SApNP (Kif-treated) | 20 | 20 | 20 | 20 | 20 | 20 | 20 | 20 |
| H3N2 HK68 HA-N95L FR SApNP (Kif-treated) | 20 | 20 | 20 | 20 | 20 | 20 | 20 | 20 |
| H3N2 HK68 HA-N95L E2p SApNP (Wild-type) | 20 | 20 | 20 | 20 | 20 | 20 | 20 | 20 |
| H3N2 HK68 HA-N95L E2p SApNP (Wild-type) | 20 | 20 | 20 | 20 | 20 | 20 | 20 | 20 |
| H3N2 HK68 HA-N95L E2p SApNP (Kif-treated) | 20 | 20 | 20 | 20 | 20 | 20 | 20 | 20 |
| H3N2 HK68 HA-N95L E2p SApNP (Kif-treated) | 20 | 20 | 20 | 20 | 20 | 20 | 20 | 20 |
| H3N2 HK68 HA-N95L I3-01v9b SApNP (Wild-type) | 20 | 20 | 20 | 20 | 20 | 20 | 20 | 20 |
| H3N2 HK68 HA-N95L I3-01v9b SApNP (Wild-type) | 20 | 20 | 20 | 20 | 20 | 20 | 20 | 20 |
| H3N2 HK68 HA-N95L I3-01v9b SApNP (Kif-treated) | 20 | 20 | 20 | 20 | 20 | 20 | 20 | 20 |
| H3N2 HK68 HA-N95L I3-01v9b SApNP (Kif-treated) | 20 | 20 | 20 | 20 | 20 | 20 | 20 | 20 |

#### f H3N2 HK68 HA-induced NAb responses against H3N2 HK68, BR07, and HK14 viruses

**A/HK/68/1968**

| Antigen | Neutralization Titer (week 8) | | | | | | | |
|---|---|---|---|---|---|---|---|---|
| | M1 | M2 | M3 | M4 | M5 | M6 | M7 | M8 |
| H3N2 HK68 HA-N95L trimer (Wild-type) | 3200 | 3200 | 800 | 400 | 3200 | 1600 | 800 | 1600 |
| H3N2 HK68 HA-N95L trimer (Wild-type) | 3200 | 3200 | 800 | 400 | 3200 | 1600 | 800 | 1600 |
| H3N2 HK68 HA-N95L trimer (Kif-treated) | 6400 | 3200 | 1600 | 1600 | 6400 | 3200 | 1600 | 6400 |
| H3N2 HK68 HA-N95L trimer (Kif-treated) | 6400 | 3200 | 1600 | 1600 | 3200 | 3200 | 1600 | 6400 |
| H3N2 HK68 HA-N95L trimer (Kif/Endo H) | 6400 | 400 | 200 | 800 | 800 | 1600 | 1600 | 200 |
| H3N2 HK68 HA-N95L trimer (Kif/Endo H) | 6400 | 200 | 200 | 800 | 800 | 1600 | 1600 | 200 |
| H3N2 HK68 HA-N95L FR SApNP (Wild-type) | 3200 | 1600 | 3200 | 1600 | 1600 | 1600 | 3200 | 1600 |
| H3N2 HK68 HA-N95L FR SApNP (Wild-type) | 3200 | 1600 | 3200 | 1600 | 1600 | 1600 | 3200 | 1600 |
| H3N2 HK68 HA-N95L FR SApNP (Kif-treated) | 1600 | 3200 | 1600 | 3200 | 1600 | 1600 | 3200 | 800 |
| H3N2 HK68 HA-N95L FR SApNP (Kif-treated) | 1600 | 3200 | 3200 | 3200 | 1600 | 1600 | 3200 | 800 |
| H3N2 HK68 HA-N95L E2p SApNP (Wild-type) | 3200 | 3200 | 1600 | 800 | 3200 | 1600 | 3200 | 3200 |
| H3N2 HK68 HA-N95L E2p SApNP (Wild-type) | 3200 | 3200 | 1600 | 800 | 3200 | 1600 | 3200 | 3200 |
| H3N2 HK68 HA-N95L E2p SApNP (Kif-treated) | 1600 | 3200 | 3200 | 3200 | 3200 | 3200 | 1600 | 6400 |
| H3N2 HK68 HA-N95L E2p SApNP (Kif-treated) | 1600 | 3200 | 3200 | 3200 | 3200 | 3200 | 1600 | 6400 |
| H3N2 HK68 HA-N95L I3-01v9b SApNP (Wild-type) | 3200 | 3200 | 6400 | 3200 | 1600 | 3200 | 1600 | 1600 |
| H3N2 HK68 HA-N95L I3-01v9b SApNP (Wild-type) | 3200 | 3200 | 3200 | 3200 | 1600 | 3200 | 1600 | 1600 |
| H3N2 HK68 HA-N95L I3-01v9b SApNP (Kif-treated) | 1600 | 3200 | 3200 | 1600 | 3200 | 1600 | 3200 | 3200 |
| H3N2 HK68 HA-N95L I3-01v9b SApNP (Kif-treated) | 1600 | 3200 | 3200 | 1600 | 3200 | 1600 | 3200 | 3200 |

**A/HK/68/1968**

##### Statistical analysis

| One-way ANOVA with Tukey's multiple comparisons test (w8) | Statistics | Adjusted P Value |
|---|---|---|
| HA trimer-Wild-type vs. HA trimer-Kif-treated | ns | 0.2168 |
| HA trimer-Wild-type vs. HA trimer-Kif/Endo H | ns | 0.9998 |
| HA trimer-Wild-type vs. HA FR SApNP-Wild-type | ns | 0.9999 |
| HA trimer-Wild-type vs. HA FR SApNP-Kif-treated | ns | 0.9999 |
| HA trimer-Wild-type vs. HA E2p SApNP-Wild-type | ns | 0.9883 |
| HA trimer-Wild-type vs. HA E2p SApNP-Kif-treated | ns | 0.5568 |
| HA trimer-Wild-type vs. HA I3-01v9b SApNP-Wild-type | ns | 0.8939 |
| HA trimer-Wild-type vs. HA I3-01v9b SApNP-Kif-treated | ns | 0.9714 |
| HA trimer-Kif-treated vs. HA trimer-Kif/Endo H | ns | 0.0647 |
| HA trimer-Kif-treated vs. HA FR SApNP-Wild-type | ns | 0.5075 |
| HA trimer-Kif-treated vs. HA FR SApNP-Kif-treated | ns | 0.5075 |
| HA trimer-Kif-treated vs. HA E2p SApNP-Wild-type | ns | 0.7901 |
| HA trimer-Kif-treated vs. HA E2p SApNP-Kif-treated | ns | 0.9996 |
| HA trimer-Kif-treated vs. HA I3-01v9b SApNP-Wild-type | ns | 0.9582 |
| HA trimer-Kif-treated vs. HA I3-01v9b SApNP-Kif-treated | ns | 0.8636 |
| HA trimer-Kif/Endo H vs. HA FR SApNP-Wild-type | ns | 0.9791 |
| HA trimer-Kif/Endo H vs. HA FR SApNP-Kif-treated | ns | 0.9791 |
| HA trimer-Kif/Endo H vs. HA E2p SApNP-Wild-type | ns | 0.8553 |
| HA trimer-Kif/Endo H vs. HA E2p SApNP-Kif-treated | ns | 0.2413 |
| HA trimer-Kif/Endo H vs. HA I3-01v9b SApNP-Wild-type | ns | 0.5939 |
| HA trimer-Kif/Endo H vs. HA I3-01v9b SApNP-Kif-treated | ns | 0.7798 |
| HA FR SApNP-Wild-type vs. HA FR SApNP-Kif-treated | ns | >0.9999 |
| HA FR SApNP-Wild-type vs. HA E2p SApNP-Wild-type | ns | >0.9999 |
| HA FR SApNP-Wild-type vs. HA E2p SApNP-Kif-treated | ns | 0.8636 |
| HA FR SApNP-Wild-type vs. HA I3-01v9b SApNP-Wild-type | ns | 0.9931 |
| HA FR SApNP-Wild-type vs. HA I3-01v9b SApNP-Kif-treated | ns | 0.9996 |
| HA FR SApNP-Kif-treated vs. HA E2p SApNP-Wild-type | ns | >0.9999 |
| HA FR SApNP-Kif-treated vs. HA E2p SApNP-Kif-treated | ns | 0.8636 |
| HA FR SApNP-Kif-treated vs. HA I3-01v9b SApNP-Wild-type | ns | 0.9931 |
| HA FR SApNP-Kif-treated vs. HA I3-01v9b SApNP-Kif-treated | ns | 0.9996 |
| HA E2p SApNP-Wild-type vs. HA E2p SApNP-Kif-treated | ns | 0.9813 |
| HA E2p SApNP-Wild-type vs. HA I3-01v9b SApNP-Wild-type | ns | >0.9999 |
| HA E2p SApNP-Wild-type vs. HA I3-01v9b SApNP-Kif-treated | ns | >0.9999 |
| HA E2p SApNP-Kif-treated vs. HA I3-01v9b SApNP-Wild-type | ns | 0.9996 |
| HA E2p SApNP-Kif-treated vs. HA I3-01v9b SApNP-Kif-treated | ns | 0.9931 |
| HA I3-01v9b SApNP-Wild-type vs. HA I3-01v9b SApNP-Kif-treated | ns | >0.9999 |

**A/BR/10/2007**

| Antigen | Neutralization Titer (week 8) | | | | | | | |
|---|---|---|---|---|---|---|---|---|
| | M1 | M2 | M3 | M4 | M5 | M6 | M7 | M8 |
| H3N2 HK68 HA-N95L trimer (Wild-type) | 50 | 50 | 50 | 50 | 50 | 50 | 50 | 50 |
| H3N2 HK68 HA-N95L trimer (Wild-type) | 50 | 50 | 50 | 50 | 50 | 50 | 50 | 50 |
| H3N2 HK68 HA-N95L trimer (Kif-treated) | 50 | 50 | 50 | 50 | 50 | 50 | 50 | 50 |
| H3N2 HK68 HA-N95L trimer (Kif-treated) | 50 | 50 | 50 | 50 | 50 | 50 | 50 | 50 |
| H3N2 HK68 HA-N95L trimer (Kif/Endo H) | 50 | 50 | 50 | 50 | 50 | 50 | 50 | 50 |
| H3N2 HK68 HA-N95L trimer (Kif/Endo H) | 50 | 50 | 50 | 50 | 50 | 50 | 50 | 50 |
| H3N2 HK68 HA-N95L FR SApNP (Wild-type) | 50 | 50 | 50 | 50 | 50 | 50 | 50 | 50 |
| H3N2 HK68 HA-N95L FR SApNP (Wild-type) | 50 | 50 | 50 | 50 | 50 | 50 | 50 | 50 |
| H3N2 HK68 HA-N95L FR SApNP (Kif-treated) | 50 | 50 | 50 | 50 | 50 | 50 | 50 | 50 |
| H3N2 HK68 HA-N95L FR SApNP (Kif-treated) | 50 | 50 | 50 | 50 | 50 | 50 | 50 | 50 |
| H3N2 HK68 HA-N95L E2p SApNP (Wild-type) | 50 | 50 | 50 | 50 | 50 | 50 | 50 | 50 |
| H3N2 HK68 HA-N95L E2p SApNP (Wild-type) | 50 | 50 | 50 | 50 | 50 | 50 | 50 | 50 |
| H3N2 HK68 HA-N95L E2p SApNP (Kif-treated) | 50 | 50 | 50 | 50 | 50 | 50 | 50 | 50 |
| H3N2 HK68 HA-N95L E2p SApNP (Kif-treated) | 50 | 50 | 50 | 50 | 50 | 50 | 50 | 50 |
| H3N2 HK68 HA-N95L I3-01v9b SApNP (Wild-type) | 50 | 50 | 50 | 50 | 50 | 50 | 50 | 50 |
| H3N2 HK68 HA-N95L I3-01v9b SApNP (Wild-type) | 50 | 50 | 50 | 50 | 50 | 50 | 50 | 50 |
| H3N2 HK68 HA-N95L I3-01v9b SApNP (Kif-treated) | 50 | 50 | 50 | 50 | 50 | 50 | 50 | 50 |
| H3N2 HK68 HA-N95L I3-01v9b SApNP (Kif-treated) | 50 | 50 | 50 | 50 | 50 | 50 | 50 | 50 |

**A/HK/4801/2014**

| Antigen | Neutralization Titer (week 8) | | | | | | | |
|---|---|---|---|---|---|---|---|---|
| | M1 | M2 | M3 | M4 | M5 | M6 | M7 | M8 |
| H3N2 HK68 HA-N95L trimer (Wild-type) | 50 | 50 | 50 | 50 | 50 | 50 | 50 | 50 |
| H3N2 HK68 HA-N95L trimer (Wild-type) | 50 | 50 | 50 | 50 | 50 | 50 | 50 | 50 |
| H3N2 HK68 HA-N95L trimer (Kif-treated) | 50 | 50 | 50 | 50 | 50 | 50 | 50 | 50 |
| H3N2 HK68 HA-N95L trimer (Kif-treated) | 50 | 50 | 50 | 50 | 50 | 50 | 50 | 50 |
| H3N2 HK68 HA-N95L trimer (Kif/Endo H) | 50 | 50 | 50 | 50 | 50 | 50 | 50 | 50 |
| H3N2 HK68 HA-N95L trimer (Kif/Endo H) | 50 | 50 | 50 | 50 | 50 | 50 | 50 | 50 |
| H3N2 HK68 HA-N95L FR SApNP (Wild-type) | 50 | 50 | 50 | 50 | 50 | 50 | 50 | 50 |
| H3N2 HK68 HA-N95L FR SApNP (Wild-type) | 50 | 50 | 50 | 50 | 50 | 50 | 50 | 50 |
| H3N2 HK68 HA-N95L FR SApNP (Kif-treated) | 50 | 50 | 50 | 50 | 50 | 50 | 50 | 50 |
| H3N2 HK68 HA-N95L FR SApNP (Kif-treated) | 50 | 50 | 50 | 50 | 50 | 50 | 50 | 50 |
| H3N2 HK68 HA-N95L E2p SApNP (Wild-type) | 50 | 50 | 50 | 50 | 50 | 50 | 50 | 50 |
| H3N2 HK68 HA-N95L E2p SApNP (Wild-type) | 50 | 50 | 50 | 50 | 50 | 50 | 50 | 50 |
| H3N2 HK68 HA-N95L E2p SApNP (Kif-treated) | 50 | 50 | 50 | 50 | 50 | 50 | 50 | 50 |
| H3N2 HK68 HA-N95L E2p SApNP (Kif-treated) | 50 | 50 | 50 | 50 | 50 | 50 | 50 | 50 |
| H3N2 HK68 HA-N95L I3-01v9b SApNP (Wild-type) | 50 | 50 | 50 | 50 | 50 | 50 | 50 | 50 |
| H3N2 HK68 HA-N95L I3-01v9b SApNP (Wild-type) | 50 | 50 | 50 | 50 | 50 | 50 | 50 | 50 |
| H3N2 HK68 HA-N95L I3-01v9b SApNP (Kif-treated) | 50 | 50 | 50 | 50 | 50 | 50 | 50 | 50 |
| H3N2 HK68 HA-N95L I3-01v9b SApNP (Kif-treated) | 50 | 50 | 50 | 50 | 50 | 50 | 50 | 50 |

**Fig. S11. Immunogenicity of H3N2 HK68 HA vaccines in mice.** (**a**) ELISA curves of mouse sera from H3N2 HK68 HA-N95L trimer and SApNP vaccine groups (n = 8 mice/group) binding to the coating antigen to H3N2 HK68 HA-N95L(1TD0) trimer (**b**) (Top) Summary of geometric mean $EC_{50}$ titers measured for H3N2 HK68 HA-N95L vaccine groups against H3N2 HK68 HA-N95L(1TD0) trimer. Color coding indicates $EC_{50}$ levels (green to red: low to high binding). (Bottom) Summary of statistical analysis performed for each timepoint. (**c**) ELISA curves of mouse sera from H3N2 HK68 HA-N95L vaccine groups at week 8 after three immunizations binding to H1N1 CA09 HA-N95L(1TD0) trimer. Summary of geometric mean $EC_{50}$ titers and statistical analysis. (**d**) H3N2 HK68 HA-N95L-induced sera HAI against vaccine matched H3N2 HK68 virus. Color coding indicates HAI titers (green to red: low to high binding). Summary of statistical analysis performed for each timepoint. (**e**) H3N2 HK68 HA-N95L-induced sera HAI against non-matched H3N2 BR07 and HK14 viruses at week 8 after three immunizations. Summary of HAI titers and statistical analysis. (**f**) H3N2 HK68 HA-N95L-induced NAb responses against H3N2 HK68, BR07, and HK14 viruses at week 8 after three immunizations. Summary of neutralization titers and statistical analysis. Error bars represent the difference between duplicate measurements at each concentration for each sample. $EC_{50}$ values were calculated using GraphPad Prism version 10.3.1. Data were analyzed using one-way ANOVA, followed by Tukey's multiple comparison post hoc test for each timepoint. For significance, ns (not significant), *$p < 0.05$, **$p < 0.01$, ***$p < 0.001$, and ****$p < 0.0001$.

**Fig. S12**

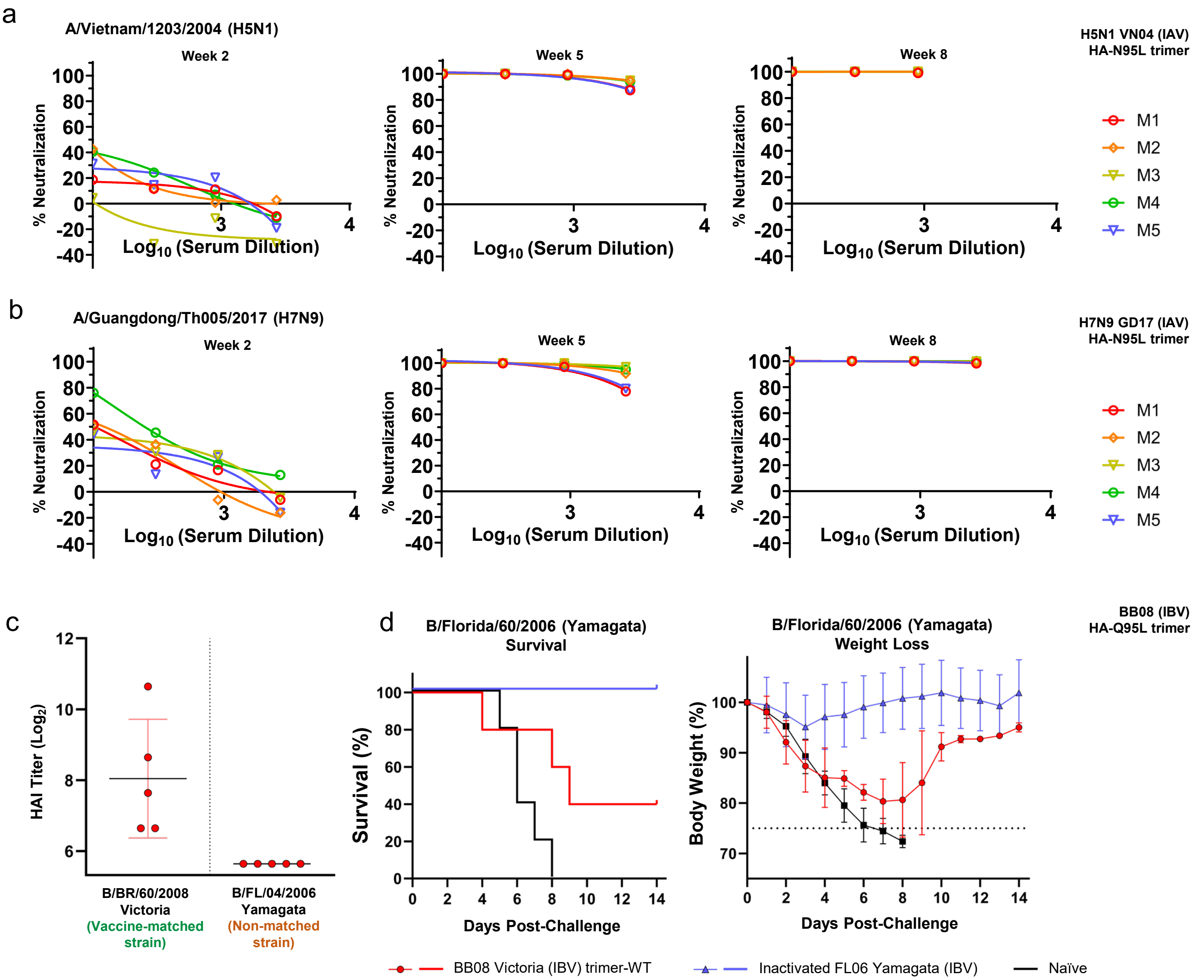

**Fig. S12. Immunogenicity and protection of IAV H5 and H7 HA-N95L trimers as well as IBV HA-Q95L trimers in mice.** (**a**) Vaccine-matched neutralization of pseudotyped A/Vietnam/1203/2004 (H5) was elicited by the sera of mice immunized with the VN04 HA-N95L trimer. (**b**) Vaccine-matched neutralization of pseudotyped A/Guangdong/Th005/2017 (H7) was elicited by the sera of mice immunized with the GD17 HA-N95L trimer. For both H5 (**a**) and H7 (**b**), two doses were sufficient to elicit potent neutralization (Week 5), which remained elevated at week 8 post-third immunization. (**c**) Hemagglutination inhibition (HAI) titers were elicited by the sera of mice immunized with the BB08 HA-Q95L trimer (Victoria lineage) against the matched virus but not against non-matched B/Florida/04/2006 (FL06; Yamagata lineage). (**d**) BB08 HA-Q95L trimer-immunized mice against a non-matched FL06 challenge. Only 40% of animals survived.

**Table S1. X-ray data collection and refinement statistics**

| Data set | **HK68 HA-N95L** | **SH13 HA-N95L** |
|---|---|---|
| **Data Collection** | | |
| X-ray source | APS 23ID-D | NSLS-II 17-ID-1 |
| Wavelength (Å) | 1.03320 | 0.92010 |
| Space group | $P2_12_12_1$ | $P2_1$ |
| Unit cell (Å) | *a* = 73.8 | *a* = 69.5 |
| | *b* = 114.2 | *b* = 115.5 |
| | *c* = 235.3 | *c* = 259.0 |
| angle (°) | 90, 90, 90 | 90, 95.9, 90 |
| Resolution (Å)[a] | 45.16-2.30 (2.34-2.30) | 49.70-2.90 (2.95-2.90) |
| Unique reflections[a] | 89,310 (4,356) | 89,227 (2,902) |
| Redundancy[a] | 11.9 (8.1) | 4.5 (3.2) |
| Average I/σ(I)[a] | 13.4 (1.0) | 4.3 (0.6) |
| Completeness (%)[a] | 100 (99.9) | 92.7 (60.8) |
| $R_{sym}$[a,b] | 0.18 (>1.0) | 0.17 (0.43) |
| $R_{pim}$[a,b] | 0.05 (0.46) | 0.08 (0.24) |
| $CC_{1/2}$[a] | 0.996 (0.798) | 0.973 (0.761) |
| No. protomers per ASU[c] | 3 | 6 |
| **Refinement** | | |
| Reflections in refinement | 88,908 | 83,805 |
| Refined residues | 1,489 | 2,966 |
| Refined waters | 112 | 77 |
| $R_{cryst}$[d] | 0.224 | 0.242 |
| $R_{free}$[e] | 0.262 | 0.281 |
| *B*-values (Å$^2$) | | |
| Protein | 70 | 71 |
| Water | 60 | 47 |
| Wilson *B*-values (Å$^2$) | 48 | 51 |
| Ramachandran values (%)[f] | 96.8, 0.3 | 93.9, 0.9 |
| R.m.s.d. bond (Å) | 0.004 | 0.006 |
| R.m.s.d. angle (deg.) | 0.66 | 1.02 |
| PDB codes | 9YOI | 9YOJ |

[a] Parentheses denote outer-shell statistics.
[b] $R_{sym} = \sum_{hkl}\sum_i |I_{hkl,i} - \langle I_{hkl}\rangle| / \sum_{hkl}\sum_i I_{hkl,i}$ and $R_{pim} = \sum_{hkl}[1/(N-1)]^{1/2}\sum_i |I_{hkl,i} - \langle I_{hkl}\rangle| / \sum_{hkl}\sum_i I_{hkl,i}$, where $I_{hkl,i}$ is the scaled intensity of the i$^{th}$ measurement of reflection *h*, *k*, *l*, $\langle I_{hkl}\rangle$ is the average intensity for that reflection, and *N* is the redundancy. $R_{pim} = \Sigma_{hkl}\ (1/(n-1))^{1/2}\ \Sigma_i\ |\ I_{hkl,i} - \langle I_{hkl}\rangle\ |\ /\ \Sigma_{hkl}\ \Sigma_i\ I_{hkl,i}$, where n is the redundancy
[c] No. protomers refers to number of HA protomers per asymmetric unit (ASU); 3 protomers per HA trimer
[d] $R_{cryst} = \sum_{hkl} |F_o - F_c| / \sum_{hkl} |F_o|$, where $F_o$ and $F_c$ are the observed and calculated structure factors.
[e] $R_{free}$ was calculated as for $R_{cryst}$, but on 5% of data excluded before refinement.
[f] Percentage of residues in the favored and outliers regions analyzed by MolProbity (Ref.165).
